# A meta-analysis of the effect of protein synthesis inhibitors on rodent fear conditioning

**DOI:** 10.1101/2022.10.11.509645

**Authors:** Clarissa F. D. Carneiro, Felippe E. Amorim, Olavo B. Amaral

## Abstract

Systematic reviews and meta-analyses have been increasingly recognized for their potential value in pre-clinical research, but their applications have not been extensively explored in behavioral neuroscience. In this work, we studied protein synthesis inhibition, a classic intervention used to disrupt fear learning, reconsolidation, and extinction in rodents, to explore how meta-analyses can identify potential moderators of its effect. We initially performed separate meta-analyses for different injection sites and target sessions to evaluate the effect of the intervention in various scenarios. Heterogeneity was further investigated by multilevel meta-regression models aggregating various sites, with article or research group as additional levels. We detected robust effects of protein synthesis inhibitors on training and reconsolidation, but not on extinction, possibly due to the lower number of studies on the latter. Our analyses identified some well-established moderators, such as intervention timing and reexposure duration. However, other factors proposed as boundary conditions for reconsolidation, such as memory age and training strength, were not associated with effect size. While our results point to the value of meta-analyses in consolidating findings from the literature, we believe that associations suggested by data synthesis should ideally be verified by well-powered, rigorous confirmatory experiments.

## Introduction

Over the last decades, biomedical research has grown consistently; however, low reproducibility rates in some areas have shed doubt over its reliability^1–3^. Industry figures for the replicability of basic research have been as low as 11% and 21%^4,5^. In experimental neurology, only 30% of replications in spinal cord injury research were at least partly successful^6^. In basic cancer biology, a 6-year systematic effort reported that 46% of experiments from highly cited articles were reproduced, with effect sizes being 85% smaller in replications than in the original studies^2^.

Systematic replication efforts, however, are cost- and labor-intensive endeavors and may not be feasible as a routine procedure in every area of science. Meta-analyses of the existing literature can thus be a useful and accessible tool to assess the reliability of effect estimates, investigate publication bias, explore sources of heterogeneity among studies and identify knowledge gaps in the literature^7^. This latter point is of particular interest to discovery science: as data synthesis is still rare in this field, conflicting results in the literature are usually not settled properly. Scientists tend to assume protocol discrepancies are responsible for divergences between studies, without considering the possibility that they might be due to random variation around statistical thresholds^8,9^.

Methods for synthesizing data from basic biomedical research, however, are not as mature as in clinical research, where outcomes are typically fewer and more clearly defined. A single article in basic discovery science can include experiments exploring multiple questions, multiple methods addressing a single question, or multiple control experiments testing boundary conditions (i.e. changing protocol parameters in order to establish the necessary conditions for an effect to be observed). This also leads to multiple experiments published in individual articles, as well as multiple articles by the same research groups, violating the assumption of independence between different results when synthesizing data.

While greater attention is present in other areas, meta-analysis of rodent behavioral studies is still incipient. Fear conditioning is a learning paradigm characterized by the association of a neutral conditioned stimulus (CS; usually a context or a tone) to an aversive unconditioned stimulus (US; for example, a footshock)^10^. Conditioning protocols vary in many ways, but the outcome used to measure memory is usually the same – the animal’s freezing response to the CS in a test session. The paradigm has been widely used to dissect the anatomical and biochemical pathways involved in the acquisition and consolidation of fear memories. Furthermore, the same paradigm is employed to study reconsolidation, a process where a reactivated memories become labile and require protein synthesis to regain stability^11^, and extinction, which involves the replacement of the original fearful association by a new, non-aversive one^12,13^.

After a series of failed attempts to replicate findings on reconsolidation of fear conditioning in rodents^14,15^, Schroyens *et al*. found evidence of publication bias in the field by reviewing the literature and contacting relevant authors^16^. In their sample, 80% of the published findings yielded statistically significant results, in contrast to only 20% of the unpublished findings shared with them^16^. The failed replications also raise the possibility that differences in protocols can determine the magnitude of the outcome. This suggests the need to determine the robustness of the effect of amnestic agents in different conditions, as well as to systematically investigate moderators that may explain the heterogeneity in results.

In this work, we use a classic intervention in mechanistic studies – the inhibition of protein synthesis to modulate fear learning, reconsolidation and extinction in mice and rats – as a model for exploring the possibilities of data synthesis in basic biomedical science. By selecting a well-established intervention with a large body of literature, we aim to investigate whether meta-analytic methods can detect consensual findings – such as the effect of protein synthesis inhibition on consolidation – and whether they can shed light on more controversial topics concerning its boundary conditions – such as the effect of reexposure duration, memory strength or memory age on reconsolidation. For this purpose, we explore different ways of aggregating studies within meta-analyses, as well as different approaches to deal with non-independence between experimental results.

## Methods

The data collection and analysis protocol was posted prior to performing the work and is available online at https://osf.io/y8r2j. Details of the methods are presented below, highlighting additions to the prespecified protocol.

### Study search and identification

PubMed, Web of Science and Scopus were searched for peer-reviewed publications using the terms “(“protein synthesis” OR “anisomycin” OR “emetine” OR “cycloheximide” OR “protein synthesis inhibitor” OR “psi” OR “puromycin” OR “acetoxycycloheximide” OR “isocycloheximide” OR “chloramphenicol” OR “rapamycin”) AND (“fear conditioning” OR “aversive conditioning” OR “conditioned fear” OR “fear memory” OR “fear learning” OR “aversive learning” OR “pavlovian conditioning” OR “fear-motivated learning” OR “threat conditioning” OR “threat learning” OR “contextual conditioning” OR “auditory conditioning” OR “fear memory retrieval” OR “fear memory extinction” OR “fear memory reconsolidation” OR “freezing behavior”)”. The specific protein synthesis inhibitors included in the search were drawn from a pilot search using “protein synthesis” and “memory”. In all searches, publication date was restricted to articles published until December 31^st^, 2018. The Scopus search was restricted to titles and abstracts, and the Web of Science search excluded book chapters, editorials, meeting abstracts and proceedings papers. Duplicates were excluded using Mendeley (version 1.17.10, Mendeley Ltd).

### Study selection

The first screening step considered only titles and abstracts and excluded (i) articles not written in English, (ii) articles not presenting original results, such as reviews, and (iii) articles not describing fear conditioning experiments in mice or rats. Retracted articles were also excluded during this stage. This step was performed by two reviewers, and at least one of them had to include the reference for it to be taken to the next stage. If the title/abstract was not clear about the three criteria described above, the article was still included for further screening.

The second screening stage considered the full text of the article, including any supplementary material if available, and papers were included if they met the following criteria: (i) described the effects of a single dose of a protein synthesis inhibitor, administered either systemically or intracerebrally, (ii) administered protein synthesis inhibitors from 1 h before up to 24 h after a behavioral session (e.g. fear conditioning training or reexposure), (iii) assessed memory by a drug-free test using freezing behavior as a measure of fear, (iv) had a clearly defined control group to which the experimental group was compared, and (v) had available data on mean freezing and standard deviation (SD) or standard error of the mean (SEM) in the test session for each experimental group. In case multiple retrieval tests were performed, only the first drug-free test session was included in the meta-analysis. Drugs were considered as protein synthesis inhibitors only when directly affecting the process of translation itself. Therefore, drugs interfering with transcription or mRNA synthesis (such as actinomycin D) or affecting translation as a consequence of interference with an upstream signalling pathway (such as rapamycin) were excluded. Drugs interfering with the synthesis of specific proteins rather with the protein synthesis process as a whole, such as antisense oligonucleotides, were also excluded.

In the second screening stage, each article was evaluated by one of two investigators who indicated which experiments should be extracted for the meta-analysis. Extraction of the data was performed by the other reviewer, so that all included experiments were double-checked, and disagreements regarding inclusion were discussed with a third investigator until consensus was reached. After the completion of the screening phase, a random sample of 10% of excluded articles was double-checked by both investigators to obtain agreement levels for exclusions.

### Data extraction

For each included article, we extracted data regarding article- and experiment-level features, as well as the quantitative results required for the meta-analyses. The dataset used in all analyses is provided as **Supplementary File 1**. Additional data collected but not used in the analyses presented here are described in the original protocol and available at https://osf.io/agdte/.

For each article, we recorded the year of publication, impact factor (as per the respective year’s Journal Citation Reports), number of citations (obtained at the end of data collection on June 25^th^ 2020) and region of origin (defined by the corresponding author’s affiliation). As articles were published within a range of over 20 years, citations were normalized as number of citations per year (number of citations/(2020 – year of publication)).

As risk of bias indicators for each article, we assessed if the following items were reported for any of the included experiments: randomization of animals to experimental groups, blinding or automation of outcome assessment (i.e. freezing measure), sample size calculation, statement regarding compliance with ethical regulations, and statement on conflict of interest.

The following experiment-level features were extracted: type of conditioning, training and testing protocols, reexposure protocols (when applicable), habituation and handling procedures, species, sex, age and number of animals per cage. Habituation and handling protocols were transformed to categorical dichotomous variables (i.e. reported or not). When housing information was presented as a range, we recorded the highest value. When age was presented as a range, we used the mean between minimum and maximum values.

Fear conditioning protocols were classified as “contextual” or “tone” according to the type of CS used in the training session. If contextual memory was evaluated by a test in the same context after tone conditioning, this was labelled as “contextual background”. If presentation of the tone and footshock were separated by an interval, this was labelled as “tone-trace”. We also recorded the number and intensity of footshocks presented at training and the time interval between sessions (training-testing, training-reexposure and reexposure-testing).

For each intervention, we extracted the active drug, dose and site of injection. These were later categorized as systemic, intra-amygdala, intra-hippocampal, intra-cerebroventricular or other (for other brain structures). We also recorded the time interval between the intervention and the behavioral session (training, reconsolidation or extinction) at which it was targeted.

Mean and SD or SEM for freezing levels and sample size were extracted for both the experimental and control groups. Numerical values were obtained from the text or legends when available, or directly from graphs when necessary using Gsys (version 2.4.6, Hokkaido University Nuclear Reaction Data Centre).

#### Missing Data

Corresponding authors were contacted for included articles in which data on the sample size of individual groups or the meaning of error bars (SD or SEM) were missing. Missing data on other protocol features were left blank.

#### Data exclusion

If authors did not respond to contact attempts, experiments with missing data on sample size were excluded from the analyses. When error bars were not defined as SD or SEM, data were included, but were assumed to represent SEM for a conservative estimate of variance. Additional tests of the same type (tone, tone-trace or contextual) on the same cohort of animals were excluded – i.e. meta-analyses only include the first drug-free test after intervention session (training or reexposure). However, if the same cohort was tested for tone and context memory (i.e. contextual background conditioning), both tests were included and sample size was divided between them for analysis. Similarly, when a group served as a control for more than one comparison, its sample size was divided by the number of extracted comparisons to avoid overrepresentation of this group in the meta-analyses.

During data extraction, we identified one article with duplicated data (same figure presented for different experiments) and one article in which mean freezing values in the text did not match the values in the figures. In both cases, the authors were contacted but could not retrieve the original data, so these experiments were excluded. After data extraction was completed, we also decided to exclude unclear or unusual test protocols, such as re-exposure protocols including the US or those in which the response to tone conditioning was not measured during the tone.

### Analysis

All analyses were performed in R^17^ (version 4.0.4). The complete code used is provided as **Supplementary File 2**.

#### Article Features

Frequency distributions of region of origin, impact factor, number of citations and year of publication are presented for the complete dataset. The percentage of risk of bias items reported was also calculated for the whole sample of articles.

#### Identification of research groups

Research groups were identified based on co-authorship networks of included articles^18^. Briefly, the list of authors was obtained for all articles included in our dataset to create a co-authorship graph using the Louvain method^19^. Modularity classes were then added to the dataset as an identification of research group. In case an article had authors attributed to different research groups, we assigned the article to the group to which most authors belonged.

#### Meta-Analyses

The random-effects model was chosen for all meta-analyses and fitted with the restricted maximum-likelihood (REML) estimator (except for multivariable meta-regressions, where the maximum-likelihood estimator was used due to computational limitations). For three-level meta-analyses, we opted for a fully random model, where random effects were used at both the article/group and experiment level. Main results present effect sizes as absolute mean differences in freezing, but we also performed sensitivity analyses using standardized mean differences (i.e. Hedges’ *g*). All meta-analyses are based on sample sizes adjusted for the number of comparisons per control group.

As defined in the protocol, separate meta-analyses were performed for interventions in different sessions (training, reconsolidation and extinction) and sites (systemic, intra-amygdala, intra-hippocampus or intracerebroventricular). The distinction between reconsolidation and extinction in this case was based on the article’s description of the experiment. We also included a meta-analysis for prefrontal cortex interventions on training, as these were identified in 3 articles (the minimum number specified in the protocol).

Also following the protocol, we performed a meta-analysis for memory reactivation, including all studies with interventions in reexposure sessions – i.e. both reconsolidation and extinction studies. Two experiments did not clearly state whether their protocol was meant to cause reconsolidation or extinction and were included in the reactivation analyses only. Of note, as most articles had more than one experiment included, they could be included in multiple meta-analyses, or contribute multiple experiments to a single meta-analysis.

Although this was not specified in the predefined protocol, we also performed aggregated meta-analyses, combining data from all injection sites for each intervention session and from all intervention sessions in each injection site. A meta-analysis of the complete dataset, including all injection sites identified and all intervention sessions was also included. This complete dataset, as well as the training and reactivation datasets, were used in moderator analyses due to its larger sample size.

For each meta-analysis, I^2^ is presented as a measure of heterogeneity alongside the p-value for the Q-test for heterogeneity, which measures whether the variability in the observed effect sizes is larger than would be expected based on sampling variability alone. Low p values thus indicate that the observed differences in effect sizes between experiments or studies would be unlikely to occur by chance alone. Possible sources of variability were further explored through moderator analyses.

#### Moderator analyses

Univariable models were built for aggregated meta-analyses (i.e. complete, training and reactivation datasets) to investigate if individual methodological variables or study features were associated with effect size across experiments. Categorical variables used were conditioning type, reporting of habituation, reporting of handling procedures, species, sex, active drug, site of injection, randomization, blinding, sample size calculation, statement of compliance with regulatory requirements, presence of conflict of interest and region of origin. Continuous variables used were shock intensity, number of shocks, time between drug administration and training/reexposure, time between training and reexposure, reexposure duration (not applicable for interventions on initial conditioning), time between intervention session and test, mean age of animals, number of animals housed together, drug dose, impact factor and citations per year. Doses were z-scored by normalizing each value according to the mean value for the respective drug and route of administration (systemic or intra-cerebral) for each species (rat or mouse). Similarly, reexposure duration was z-scored according to conditioning type using the mean number of tones (for tone reexposure) or duration (for contextual reexposure) across studies to allow inclusion of both types of study in the same analysis.

For each univariable model, we present the R^2^ value and the p-value of the Q test of moderators. In 3-level models, R^2^ values were calculated as the difference in the sum of σ^2^ values for both levels between the model with no moderators and the univariable model, divided by the sum of σ^2^ values in the model with no moderators. In case this yielded a negative value, R^2^ was considered to be zero.

As defined in the protocol, we restricted the number of moderators included in multivariable models to one variable for every 10 experiments. The following priority order was defined *a priori* in the protocol: re-exposure duration (not applicable to training meta-analyses), drug, dose, time between intervention session and test, time to reexposure (not applicable to training meta-analyses), species, shock intensity, number of shocks, sex and age of animals. Due to low prevalence of reporting, animal age was not included among the variables tested in any of the datasets. In addition to the predefined list, we added intervention session (applicable only to the complete dataset), injection site, conditioning type and time between drug administration and training/reexposure, which were selected on the basis of univariable models.

All combinations of variables from the selected list were tested in multivariable models, and the best models were ranked by corrected Akaike Information Criteria (AICc). For each best model selected (for complete, training and reactivation datasets with 2- and 3-level analyses), we decomposed the R^2^ value for each moderator included. For this, we calculated the mean of the differences between R^2^ from models with and without the moderator in all possible orders of moderator inclusion. Additionally, we performed a Q test of moderators for each variable (including all dummy variables for each categorical moderator) to obtain p-values for individual variables.

#### Publication Bias

Due to limitations in sample size, publication bias analyses were restricted to systemic, intra-amygdala and intra-hippocampal interventions on training and reconsolidation. This was performed using Egger’s regression^20^ and trim-and-fill analyses^21,22^. As recommended by the authors of the method^23^, both L_0_ and R_0_ estimators were used to estimate the number of missing studies and to correct the effect estimate (and its p-value) after imputation of these studies.

As an additional exploratory measure of publication bias, excess significance tests^24^ were also performed for each meta-analysis. These were based on the meta-analytic estimates using sample sizes as reported in the articles, with no adjustment for control group or test session redundancies.

Finally, as planned in the protocol, statistical power for a two-sample two-sided test was calculated for each experiment, considering its own sample size (without adjustments) and standard deviation, along with the respective meta-analytical effect size estimate expressed as absolute mean difference and an alpha of 0.05. We used this to perform correlation analyses between statistical power and effect size with article-level features. For this, we used the mean power and effect size of all experiments per article and calculated Spearman’s correlation with impact factor, citations per year and risk of bias. In this case, risk of bias was used as a score calculated as 5 minus the sum of reported items.

## Results

### Study selection

Our search initially retrieved 934 articles, of which 91 met our inclusion criteria^11,13,25–113^. **Figure 1** presents a PRISMA flowchart for the inclusion and exclusion of articles^114^.

**Figure 1-.**
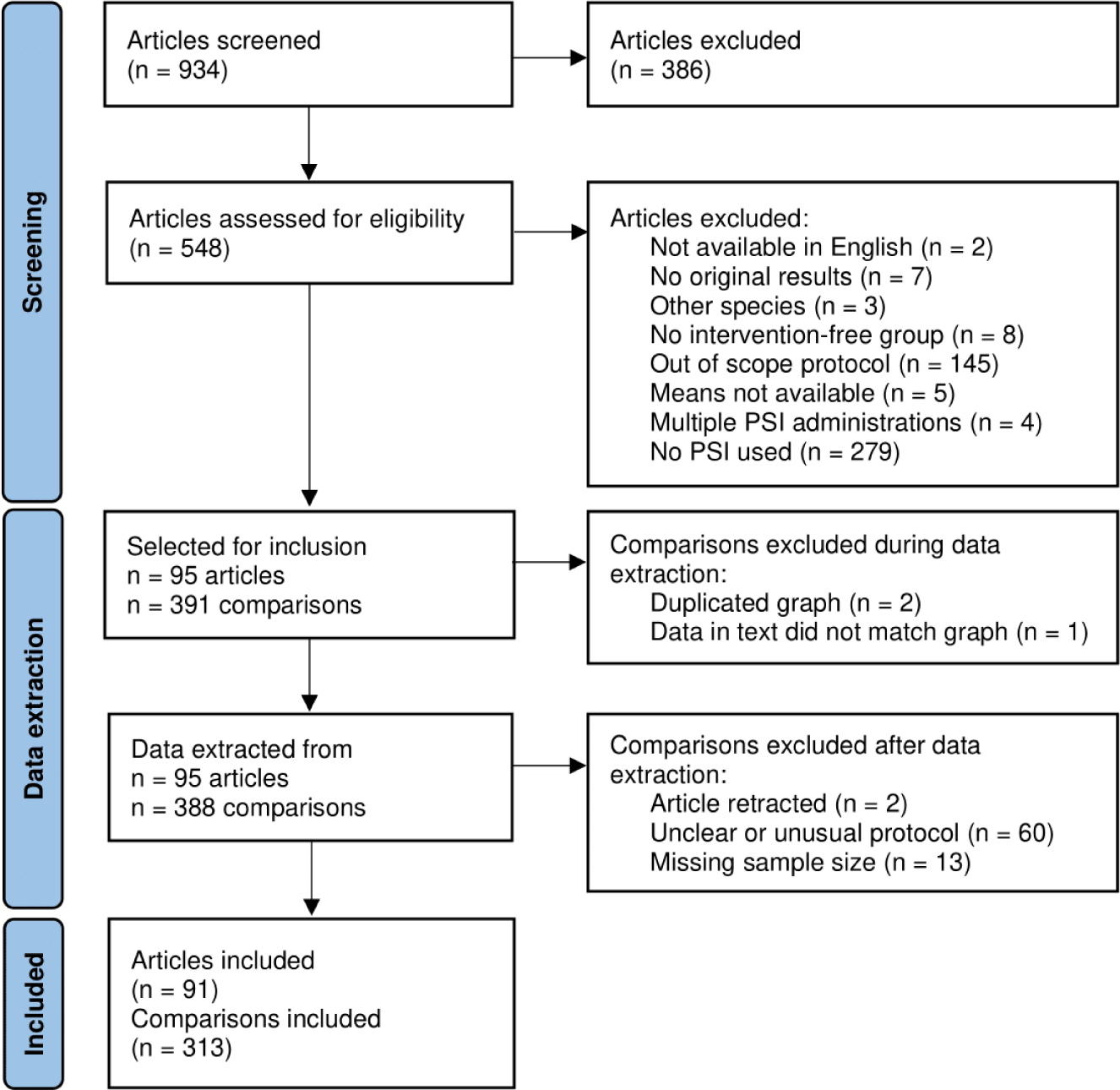
Flowchart of screening stages. In the first stage of screening (n=934), two reviewers assessed titles and abstracts. If one reviewer included the article, it was selected for full-text screening. During this stage we did not record reasons for exclusion. The second screening stage (n=548) considered the full-text and supplementary material, with each being assessed by a single reviewer (no disagreements found in a double-screened sample of 36 articles). Additional exclusion of comparisons took place during or after data extraction (see Methods for details), leaving 313 comparisons from 91 articles to be included in the analyses. PSI: protein synthesis inhibitor.

### Study description

**Suppl. Figure 1** presents the frequency distribution of region of origin, year of publication, impact factor and citations for the full set of articles. Included articles were mostly from North America, published between 2005 and 2015 in journals with a median impact factor of 4.3. Frequency of reporting of risk of bias factors is presented in **Suppl. Table 1**. A fair number of articles reported blinded/automated assessment of outcomes, while randomization and sample size calculation were reported very rarely.

A summary of protocol variables is presented in **Table 1** (full data in **Suppl. Table 2)**. Most comparisons studied the inhibition of protein synthesis in training or reconsolidation sessions, while studies of extinction were less common. Systemic injections accounted for about a third of all comparisons, while intracerebral injections were mostly directed at the amygdala and hippocampus. Distribution between species was balanced between rats and mice, with 94% of all comparisons including only male animals, and 86% of studies using anisomycin to inhibit protein synthesis.

**Table 1-.** Summary of protocol features of comparisons included in the analyses. For categorical variables, we show the number of comparisons per category, while for continuous variables we show the median and range (minimum and maximum) and the number of comparisons in which the variable was reported. Percentages refer to the total of 313 experiments, except for reexposure duration, where the total is 58 experiments using tone-cued re-exposure protocols and 87 experiments using contextual reexposure protocols. “Contextual background” refers to cases where conditioning was performed with tone cues, but testing/reexposure used the context as the CS. Complete data is presented in Suppl. Table 2.

| Protocol variable | Value | N (%) |
| --- | --- | --- |
| Intervention session | Training | 168 (53.7%) |
|  | Reconsolidation | 119 (38.0%) |
|  | Extinction | 23 (7.3%) |
|  | Reactivation | 3 (1.0%) |
| Site of injection | Systemic | 97 (31.0%) |
|  | Amygdala | 94 (30.0%) |
|  | Hippocampus | 71 (22.7%) |
|  | Cerebral ventricles | 9 (2.9%) |
|  | Prefrontal cortex | 9 (2.9%) |
|  | Other | 33 (10.5%) |
| Type of conditioning | Contextual | 138 (44.1%) |
|  | Tone | 103 (32.9%) |
|  | Contextual background | 59 (18.8%) |
|  | Tone-trace | 13 (4.2%) |
| Number of footshocks | 2 (1 – 76) | 313 (100%) |
| Footshock intensity | 0.75 mA (0.3 – 2) | 310 (99.0%) |
| Reexposure duration | Context: 3 min (0.5 – 30) | 84 (96.5%) |
|  | Tone: 1 tone (1 – 15) | 58 (100%) |
| Species | Rats | 167 (53.4%) |
|  | Mice | 146 (46.6%) |
| Sex | Male | 293 (93.6%) |
|  | Both | 18 (5.8%) |
|  | Not informed | 2 (0.6%) |
| Age | 10 weeks (4 – 22) | 122 (39.0%) |
| Animals per cage | 1 (1 – 6) | 242 (77.3%) |
| Active drug | Anisomycin | 270 (86.3%) |
|  | Cycloheximide | 41 (13.1%) |
|  | 4EGI-1 | 2 (0.6%) |

### Meta-analyses

As planned *a priori*, we initially performed meta-analyses separated by intervention session (training, reconsolidation, extinction and reactivation) and site of drug administration (systemic, amygdala, hippocampus and cerebral ventricles). Most other intracerebral sites of drug administration were reported in less than 3 articles (**Suppl. Table 3**); thus, the only additional meta-analysis performed was for interventions on training with prefrontal cortex injections. In addition to these individual meta-analyses, we performed an exploratory meta-analysis with the complete dataset in order to explore predictors with greater power. **Table 2** summarizes these analyses, and forest plots are presented as **Suppl. Figures 2-4**.

**Table 2-.** Summary of meta-analyses. Effect sizes are expressed as absolute mean differences in freezing percentages between PSI-treated and control groups in the test session. Sample size is the number of experiments. p values refer to main effect comparisons in random-effects meta-analyses (4^th^ column) and Q-tests for heterogeneity (6^th^ column). i.c.v.: intracerebroventricular, CI: confidence interval.

| Dataset | Sample size | Effect size (% freezing) [95% CI] | Meta-analysis p-value | I<sup>2</sup> | Q-test p-value |
| --- | --- | --- | --- | --- | --- |
| Training, systemic | 57 | -20.2<br>[-24.0, -16.3] | 6.4x10<sup>-25</sup> | 82.8% | 1.5x10<sup>-38</sup> |
| Training, amygdala | 38 | -20.2<br>[-26.1, -14.3] | 1.5x10<sup>-11</sup> | 66.6% | 1.5x10<sup>-10</sup> |
| Training, hippocampus | 41 | -14.1<br>[-18.9, -9.3] | 9.0x10<sup>-9</sup> | 87.3% | 1.9x10<sup>-23</sup> |
| Training, i.c.v. | 3 | -22.0<br>[-53.5, 9.6] | 0.173 | 80.9% | 0.004 |
| Training, prefrontal cortex | 6 | -10.7<br>[-20.5, -0.9] | 0.033 | 0% | 0.713 |
| Reconsolidation, systemic | 36 | -18.6<br>[-23.1, -14.1] | 4.2x10<sup>-16</sup> | 83.4% | 4.4x10<sup>-35</sup> |
| Reconsolidation, amygdala | 47 | -20.1<br>[-25.2, -15.0] | 1.4x10<sup>-14</sup> | 78.8% | 5.5x10<sup>-26</sup> |
| Reconsolidation, hippocampus | 20 | -19.3<br>[-27.8, -10.9] | 7.8x10<sup>-6</sup> | 89.6% | 1.9x10<sup>-34</sup> |
| Reconsolidation, i.c.v. | 4 | -12.5<br>[-43.9, 18.9] | 0.435 | 88.5% | 9.0x10<sup>-4</sup> |
| Extinction, systemic | 4 | 28.0<br>[21.4, 34.6] | 1.2x10<sup>-16</sup> | 0% | 0.771 |
| Extinction, amygdala | 8 | -1.1<br>[-11.6, 9.4] | 0.831 | 69.9% | 0.001 |
| Extinction, hippocampus | 8 | 6.3<br>[-7.3, 19.9] | 0.364 | 79.2% | 0.002 |
| Extinction, i.c.v. | 2 | 21.0<br>[-12.2, 54.2] | 0.214 | 84.8% | 0.010 |
| Reactivation, systemic | 40 | -14.2<br>[-20.1, -8.2] | 3.3x10<sup>-6</sup> | 91.7% | 1.3x10<sup>-66</sup> |
| Reactivation, amygdala | 56 | -18.1<br>[-23.2, -13.0] | 2.9x10<sup>-12</sup> | 82.4% | 2.3x10<sup>-43</sup> |
| Reactivation, hippocampus | 30 | -13.2<br>[-21.3, -5.2] | 0.001 | 90.7% | 1.9x10<sup>-59</sup> |
| Reactivation, i.c.v. | 6 | 0.9<br>[-21.7, 23.5] | 0.939 | 91.6% | 2.8x10<sup>-5</sup> |
| Complete | 313 | -15.4<br>[-17.4, -13.3] | 6.0x10<sup>-48</sup> | 87.0% | 6.5x10<sup>-284</sup> |

We found significant negative effects on test freezing after protein synthesis inhibition in training and reconsolidation for almost all studied structures, with roughly comparable effect sizes. For extinction, however, an effect in the opposite direction (i.e. preventing freezing decline) was found only for systemic injections. That said, these results should be interpreted with caution, as there were far fewer studies on extinction than on training or reconsolidation. Since the reactivation dataset largely overlaps with the reconsolidation one, results were similar between them, but with smaller effect sizes for reactivation due to the inclusion of extinction experiments. Most analyses showed a large amount of heterogeneity between experiments. Results of the same analyses using standardized mean differences (Hedge’s g) are shown as a robustness check in **Suppl. Table 4**.

### Publication bias

To evaluate the presence of publication bias in our sample, we performed trim-and-fill analyses and Egger’s regression for systemic, intra-amygdala and intra-hippocampal interventions on training and reconsolidation (other meta-analyses were not included due to their small sample size). **Figure 2** presents funnel plots with the results of Egger’s regression and trim-and-fill analyses using the L_0_ method (results with the R_0_ method are presented in **Suppl. Table 5**). Unlike what was described by Schroyens et al. (2021) for various drugs in contextual fear reconsolidation, we did not find consistent evidence of publication bias for protein synthesis inhibitors: a significant correlation between effect size and precision was found only for intra-hippocampal interventions on reconsolidation, but in the opposite direction as would be expected from publication bias.

**Figure 2-.**
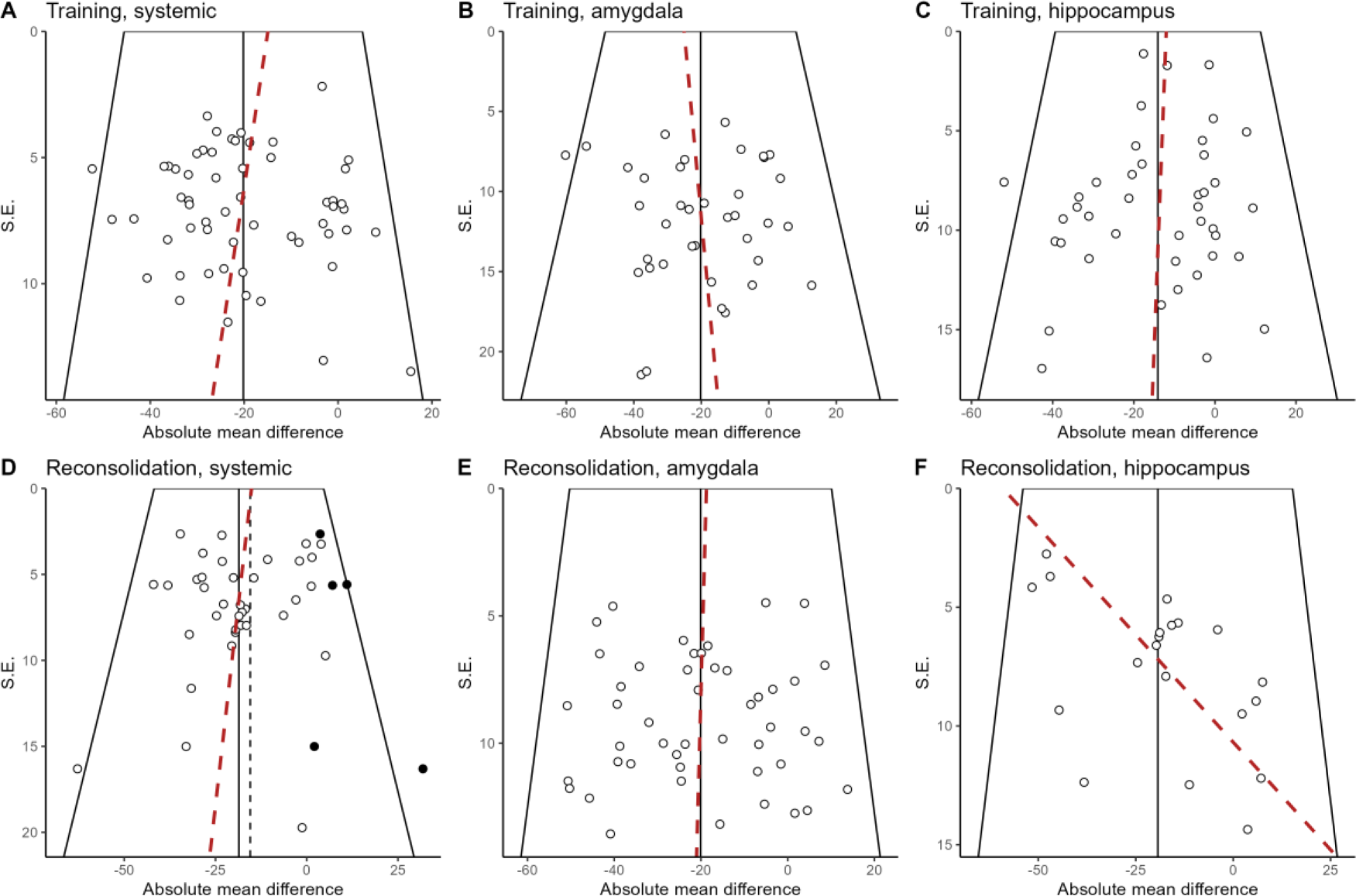
Funnel plots and Egger’s regression. Each observed study is represented by the white circles and the red dashed line represents Egger’s regression line. Black circles represent missing studies imputed by the trim-and-fill L_0_ method. **(A)** Training, systemic. Egger’s regression p-value = 0.39. **(B)** Training, amygdala. Egger’s regression p-value = 0.75. **(C)** Training, hippocampus. Egger’s regression p-value = 0.39. **(D)** Reconsolidation, systemic. Egger’s regression p-value = 0.24. Estimated number of missing studies = 5. **(E)** Reconsolidation, amygdala. Egger’s regression p-value = 0.98. **(F)** Reconsolidation, hippocampus. Egger’s regression p-value = 0.009. S.E.; Standard error.

Another potential source of bias in a meta-analysis is lack of statistical power in the original studies, which along with publication bias can lead to effect size inflation. In **Suppl. Figure 5,** we show the distribution of sample sizes and statistical power for training and reconsolidation experiments for systemic, intra-amygdala and intra-hippocampus interventions, calculated using the respective meta-analytic estimate as the reference effect size. Mean sample size (± SD) for all experiments combined was 9.1 ± 2.8 and mean statistical power for the corresponding meta-analytical effect size estimate was 0.66 ± 0.26. While differences in power for individual meta-analyses are heavily determined by differences between effect size estimates, experiments with intracerebral injections also tended to have smaller sample sizes than those with systemic interventions. No correlation was found between mean effect size or statistical power and impact factor, citations or risk of bias score for articles (**Suppl. Figure 6**). Consistent with what we found by funnel-plot analysis, excess significance test did not detect evidence of publication bias (**Suppl. Table 6**).

### Meta-regressions

A major challenge in performing meta-analyses of basic and pre-clinical research is defining the boundaries of a research question – i.e. how similar experiments should be for their data to be synthesized meaningfully. On one hand, including experiments with diverse protocols adds heterogeneity to the analyses, rendering their interpretation more difficult; on the other, this allows one to explore the impact of protocol variables on effect size.

*A priori*, we determined that our analyses would be divided by the target session of drug administration (training, reconsolidation or extinction) and by the site of drug administration (hippocampus, amygdala, i.c.v. or systemic), as presented in **Table 2** and **Suppl. Table 4**. However, to investigate whether aggregating data could provide additional information, we also combined all experiments for each intervention session (i.e. training, reconsolidation, extinction and reactivation), testing the injection site as a moderator (**Table 3, Suppl. Table 7**), and combined all experiments for each site of injection, testing the session as a moderator (**Table 3, Suppl. Table 8**). While the variance explained by injection site was relatively small (and driven mostly by structures other than the hippocampus and amygdala), the intervention session had a larger impact on effect sizes, particularly due to the influence of extinction experiments, in which the direction of effect was reversed. Nevertheless, a large amount of residual heterogeneity persisted even after adding these moderators.

**Table 3-.** Meta-regression models for testing intervention site and target session. Grey rows show the results of the models without moderators, while white lines show the results of meta-regressions. p-values refer to the Q-test for residual heterogeneity (3^rd^ column) or the test of significance for the moderator (5^th^ column). R^2^ and p-values for multivariate meta-regression are presented in order, with the first row showing the results of intervention session and the second row showing those of injection site.

| Dataset and moderators | I<sup>2</sup> | p-value | R<sup>2</sup> | p-value |
| --- | --- | --- | --- | --- |
| Training, no moderators | 81.9% | 9.7x10<sup>-79</sup> |  |  |
| Training, injection site as moderator | 80.3% | 3.3x10<sup>-69</sup> | 5.2% | 0.058 |
| Reconsolidation, no moderators | 85.9% | 2.3x10<sup>-123</sup> |  |  |
| Reconsolidation, injection site as moderator | 83.2% | 1.2x10<sup>-92</sup> | 17.6% | 2.3x10<sup>-4</sup> |
| Extinction, no moderators | 84.4% | 5.0x10<sup>-12</sup> |  |  |
| Extinction, injection site as moderator | 70.0% | 1.3x10<sup>-5</sup> | 37.1% | 0.018 |
| Systemic, no moderators | 88.2% | 3.7x10<sup>-104</sup> |  |  |
| Systemic, intervention session as moderator | 82.4% | 4.2x10<sup>-70</sup> | 36.6% | 3.8x10<sup>-10</sup> |
| Amygdala, no moderators | 77.8% | 9.2x10<sup>-51</sup> |  |  |
| Amygdala, intervention session as moderator | 74.1% | 6.4x10<sup>-36</sup> | 16.5% | 0.002 |
| Hippocampus, no moderators | 90.5% | 1.0x10<sup>-89</sup> |  |  |
| Hippocampus, intervention session as moderator | 88.3% | 1.6x10<sup>-55</sup> | 17.0% | 0.004 |
| i.c.v., no moderators | 91.7% | 1.8x10<sup>-9</sup> |  |  |
| i.c.v., intervention session as moderator | 84.9% | 6.6x10<sup>-6</sup> | 14.1% | 0.198 |
| Other, no moderators | 70.0% | 2.0x10<sup>-12</sup> |  |  |
| Other, intervention session as moderator | 57.0% | 8.1x10<sup>-6</sup> | 41.1% | 4.9x10<sup>-4</sup> |
| Complete, no moderators | 86.9% | 2.9x10<sup>-282</sup> |  |  |
|  | 81.9% | 2.4x10<sup>-195</sup> | 20.9% | 7.6x10<sup>-14</sup> |
| Complete, intervention session and injection site as moderators | | | 7.4% | $9.0 \times 10^{-5}$ |

A final possibility to increase sample size for meta-regression is by combining both intervention sessions and sites of drug administration using the complete dataset. To assess the impact of this merging, we used a multivariable meta-regression model considering both intervention session and site of injection (**Table 3, Suppl. Table 9**). Although both moderators account for a relevant amount of heterogeneity, we considered it reasonable to use the complete dataset to explore additional protocol variables in meta-regression, due to the greater sample size achieved. Thus, from here onward, we restrict our analyses to the complete dataset and to datasets with all interventions (irrespectively of site) on training and on reactivation (including reconsolidation, extinction and reexposure interventions).

### Multilevel meta-analyses

The standard random-effects models in the meta-analyses presented above consider two sources of variability: the real variance among different experiments and the sampling error of each experiment. Both of these, however, are nested within at least two additional sources of variability: that between articles from which experiments are obtained and that between research groups publishing the articles. As multiple experiments within a paper and/or by a research group are not completely independent of each other, not taking these levels into account may bias the analyses towards results of particular articles or research groups that are overrepresented in the sample.

To test the importance of including these additional levels of variability, we applied an authorship network-based method^18^ to identify research groups in the complete set of articles included (see Methods for details). Forty-two groups were identified by modularity analyses, as shown in **Suppl. Fig. 7**. The distribution of experiments within articles and research groups can be visualized in **Suppl. Fig. 8**. Although most articles and research groups contribute only a few experiments to the sample, a few are distinctly overrepresented in the sample (with a single research group contributing 58 experiments).

We then built additional models assuming random effects both within and between either articles or research groups (**Table 4**). This led to minor changes in estimates, and variance was larger between experiments than between either articles or research groups for all models. Nevertheless, in the reactivation dataset, these additional levels accounted for a reasonable fraction of the observed heterogeneity. The amount of variance explained by articles and research groups can also be observed by including articles and research groups as moderators (i.e. fixed factors) in meta-regressions (**Suppl. Table 10**), although this does not allow correction of effect estimates, as occurs when they are included as levels (i.e. random factors).

**Table 4-.** Three-level meta-analyses. Two 3-level models were built, considering nesting within clusters that represented either articles or research groups, which are modeled as additional random factors. The two-level (i.e. standard random-effects) model is also included for comparison. Effect sizes are in absolute mean differences. CI, confidence interval; Exp., experiments; cl., clusters.

| Model | Sample size |  | Effect size<br>(% freezing)<br>[95% CI] | p-value | Variance<br>[95% CI] |  | I<sup>2</sup> |  |
| --- | --- | --- | --- | --- | --- | --- | --- | --- |
|  | Exp. | cl |  |  | Between experiments | Between clusters | Exp. | Cluster |
| Complete dataset |  |  |  |  |  |  |  |  |
| 2-level | 313 | 0 | -15.35<br>[-17.4, -13.3] | 7.9x10<sup>-48</sup> | 265.8<br>[215.7, 325.2] | - | 85.8% | - |
| 3-level<br>(article) | 313 | 91 | -15.14<br>[-17.9, -12.4] | 8.2x10<sup>-28</sup> | 202.3<br>[157.0, 260.5] | 68.3<br>[25.2, 135.1] | 65.2% | 22.0% |
| 3-level<br>(group) | 313 | 42 | -15.27<br>[-18.6, -12.0] | 1.2x10<sup>-19</sup> | 237.2<br>[188.9, 296.4] | 40.4<br>[3.7, 125.8] | 74.7% | 12.7% |
| Training dataset |  |  |  |  |  |  |  |  |
| 2-level | 168 | 0 | -17.35<br>[-19.8, -14.9] | 2.9x10<sup>-44</sup> | 174.6<br>[122.8, 230.2] | - | 81.9% | - |
| 3-level<br>(article) | 168 | 57 | -17.19<br>[-19.8, -14.5] | 4.5x10<sup>-37</sup> | 163.3<br>[114.8, 228.9] | 11.4<br>[0, 58.1] | 76.6% | 5.4% |
| 3-level<br>(group) | 168 | 29 | -17.31<br>[-19.8, -14.9] | 4.4x10<sup>-44</sup> | 174.6<br>[128.5, 236.9] | 1.1x10<sup>-6</sup><br>[0, >10.0] | 81.9% | 0% |
| Reactivation dataset |  |  |  |  |  |  |  |  |
| 2-level | 145 | 0 | -13.24<br>[-16.7, -9.8] | 3.1x10<sup>-14</sup> | 361.6<br>[280.8, 497.8] | - | 89.5% | - |
| 3-level<br>(article) | 145 | 56 | -13.06<br>[-17.5, -8.6] | 6.8x10<sup>-9</sup> | 249.1<br>[171.2, 363.6] | 119.8<br>[32.2, 265.5] | 60.6% | 29.1% |
| 3-level<br>(group) | 145 | 28 | -12.21<br>[-17.9, -6.5] | 2.5x10<sup>-5</sup> | 296.9<br>[215.4, 413.6] | 96.9<br>[6.6, 309.2] | 68.0% | 22.2% |

We also built a four-level model with articles and research groups as different levels (**Suppl. Table 11**). However, variance estimations were much less precise in this case, indicating a lack of clear distinction between each level’s contribution to total heterogeneity.

### Univariable meta-regressions

To explore the impact that individual features in the experiments might have on effect sizes, we initially performed univariable meta-regressions using the two-level model for all protocol and article variables collected, including risk of bias measures. Results are summarized in **Figure 3A**, with a more extensive description available in Suppl. Tables 12-14.

**Figure 3-.**
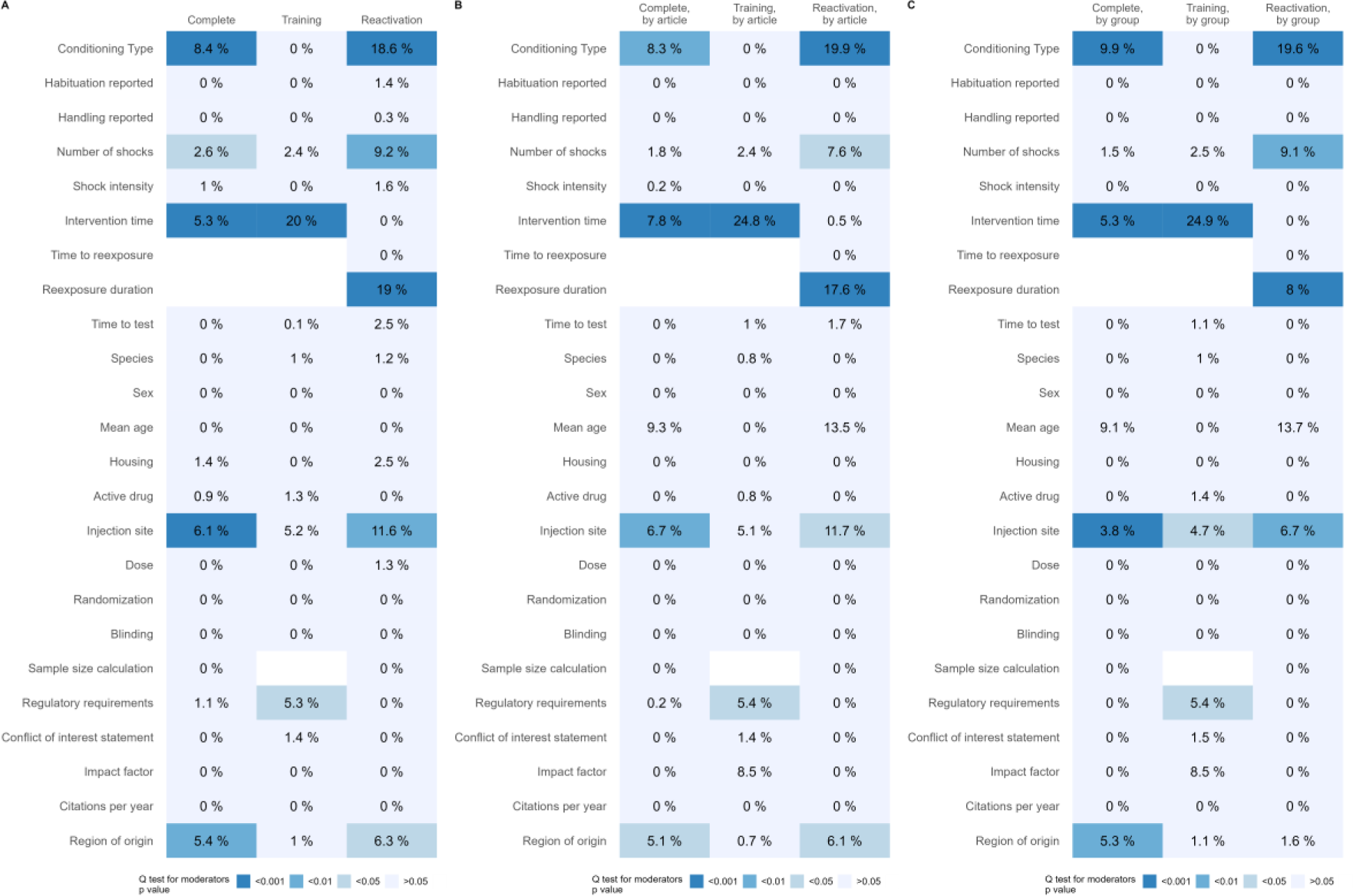
Summary of univariable meta-regression models. Each column represents a dataset, while lines correspond to each variable. Cell colors represent p-values for Q tests of the respective variable as a moderator and numbers within cells are the R^2^ value for the moderator in a univariable model. White cells indicate that the variable was not applicable or unfeasible to include in the model due to lack of variability. Reexposure duration is based on z-scored values for reexposure to tone (measured in number of tones) and to context (measured in minutes), based on the mean and standard deviation for each type of conditioning. Doses were also z-scored based on the mean and standard deviation of each drug and route of administration for each species. **(A)** Two-level models. **(B)** Three-level models, accounting for nesting of experiments within articles. **(C)** Three-level models, accounting for nesting of experiments within research groups. Full details for each model, including betas and directionality of effects, are shown on **Suppl. Tables S12-S21**.

For interventions in the training session, the time between drug administration and training (i.e. intervention time) was found to be a significant moderator of effect size. As can be observed in **Suppl. Figure 9**, this relationship is largely driven by injections performed 24 hours after the training session, which yield near-null effects.

In the reactivation sample, reexposure duration and number of tones (which were z-scored and included as a single variable) predictably influenced effect size. As there was little variation in the number of tones across experiments, the moderator effect can be attributed mainly to contextual fear conditioning experiments, in which effects on freezing were negative with under 10 minutes of reexposure (suggesting interference with reconsolidation), variable between 10 and 20 minutes, and mostly positive with 30 minutes of reexposure (suggesting effects on extinction) (**Suppl. Figure 10A-B**). This is confirmed by performing separate meta-regressions based on the duration of reexposure to context or number of tones (**Suppl. Table 15**), which show that the effect of reexposure duration on effect sizes is much more robust for contextual conditioning than for tone conditioning, in which experiments exploring larger number of tones were less common.

Additionally, conditioning type by itself also impacted effect sizes, with tone conditioning leading to larger effects on reconsolidation. As can be observed in **Suppl. Figure 10C**, the impact of conditioning type is probably due to the large number of near-null effects for contextual fear reconsolidation, as well as to a greater number of extinction experiments using contextual conditioning. A higher number of shocks was also associated with larger effects on reconsolidation. That said, as can be seen from **Suppl. Figure 11A**, this association is mostly driven by a few experiments with 60 shocks and large effects, while an opposite trend is observed for experiments with up to 10 shocks.

Other proposed boundary conditions for reconsolidation, such as time between training and reexposure, were not detected as moderators in the model. That said, **Suppl. Figure 11B** shows that, for the few experiments with more than 20 days between training and reexposure, effects seem to be consistently smaller than those with shorter intervals, where there is greater variability in results. It is also worth noting that extinction experiments generally used short intervals, which might have obscured an effect of this variable on reconsolidation. Surprisingly, drug dose was also not found to be a significant moderator, although this could be due to the relatively narrow range of doses used in most studies when specific drugs, administration routes and species were taken into account (**Suppl. Figure 12**).

Next, we performed the same meta-regressions for three-level models, accounting for the nesting of experiments within articles (**Figure 3B** and **Suppl. Tables 16-18**) and within research groups (**Figure 3C** and **Suppl. Tables 19-21**). In these models, the amount of heterogeneity accounted for most moderators is lower, suggesting that some of the moderator effects might be explained by the article/research group of origin instead. However, the main findings persist – as in the case of injection time in training, and of conditioning type and reexposure duration in reactivation.

### Multivariable meta-regressions

Lastly, we explored multivariable models to investigate whether the moderator effects detected in univariable regressions could be due to confounding by other variables. We started with the complete dataset and the original variables that separated the analyses (injection site and intervention session), the two variables with the largest effects in univariable models (conditioning type and intervention time) and 7 other variables selected *a priori* (active drug, dose, time to test, species, number of shocks, shock intensity and sex). Missing data for some of these variables led to the exclusion of 36 experiments, leaving 277 to be used in these analyses.

In the training dataset, including the same variables listed above (except for intervention session) led to the exclusion of 11 experiments, leaving 157 available for the analyses. In the reactivation dataset, the predetermined list of variables included two additional variables: reexposure duration and time between test and reexposure. This led to exclusion of 19 experiments, leaving 126 available.

After testing all possible variable combinations in the 2-level and 3-level models (2,048 in the complete dataset, 1,024 in the training and reactivation datasets), we ranked the resulting models by the Akaike Information Criteria (AICc). For each dataset and clustering option, we present a summary of the best models in **Table 7** and effect sizes are presented in **Suppl. Tables 22-30**.

**Table 7-.** Best multivariable meta-regression models for each dataset and clustering option. Only the model with lowest AICc is presented for each scenario. Weight refers to the Akaike weights. Q-test p-values refer to the test for all included moderators. Effect sizes and p values for individual moderators in each model are presented in **Suppl.** Tables 22-30.

| Dataset | Clusters | Included variables | AICc | Weight | I<sup>2</sup> | R<sup>2</sup> | Q-test p-value |
| --- | --- | --- | --- | --- | --- | --- | --- |
| Complete | 2-level | Intervention session<br>Site of injection<br>Conditioning type<br>Active drug<br>Number of shocks<br>Intervention time | 2328.1 | 0.060 | 76.4% | 45.1% | 9.6x10<sup>-24</sup> |
| Complete | 3-level (article) | Intervention session<br>Site of injection<br>Active drug<br>Number of shocks<br>Intervention time | 2328.8 | 0.056 | 79.2% | 44.5% | 4.6x10<sup>-22</sup> |
| Complete | 3-level (group) | Intervention session<br>Site of injection<br>Active drug<br>Number of shocks<br>Intervention time | 2326.7 | 0.081 | 79.7% | 44.0% | 2.1x10<sup>-22</sup> |
| Training | 2-level | Site of injection<br>Intervention time | 1299.2 | 0.037 | 73.1% | 27.7% | 1.9x10<sup>-6</sup> |
| Training | 3-level (article) | Site of injection<br>Intervention time | 1301.4 | 0.036 | 75.4% | 30.7% | 1.9x10<sup>-6</sup> |
| Training | 3-level (group) | Site of injection<br>Intervention time | 1301.4 | 0.037 | 75.4% | 30.1% | 1.2x10<sup>-6</sup> |
| Reactivation | 2-level | Site of injection<br>Reexposure duration<br>Number of shocks | 1090.9 | 0.117 | 81.1% | 45.2% | 5.1x10<sup>-14</sup> |
| Reactivation | 3-level (article) | Site of injection<br>Reexposure duration<br>Number of shocks | 1092.4 | 0.124 | 82.6% | 43.6% | 7.3x10<sup>-13</sup> |
| Reactivation | 3-level (group) | Site of injection<br>Reexposure duration<br>Number of shocks | 1093.2 | 0.122 | 82.3% | 47.4% | 5.1x10<sup>-14</sup> |

Notably, multivariable models did not include some article-level variables detected as significant by univariable models, such as presence of a regulatory requirements statement and region of origin, suggesting that these effects were likely due to confounding. Nevertheless, most variables related to the experimental protocol itself were preserved. These results can also be visualized by the importance of all variables in each set of models (**Suppl. Figure 13**).

For each of these models we decomposed the R^2^ value for each of the variables selected to investigate the contribution of each one (**Figure 4**). Once more, it is notable that R^2^ values in the reactivation set are lower for most variables in multilevel models, suggesting that some of the correlation between these variables and effect sizes is driven by confounding from the use of specific reexposure protocols by particular articles or groups. Conditioning type in particular was not detected as a moderator when additional levels are considered, suggesting that this effect might have been driven by research groups who preferentially use tone or context conditioning. This does not seem to be the case in the training set, where including articles or groups as levels actually increased the effect of intervention time and site of injection.

**Figure 4-.**
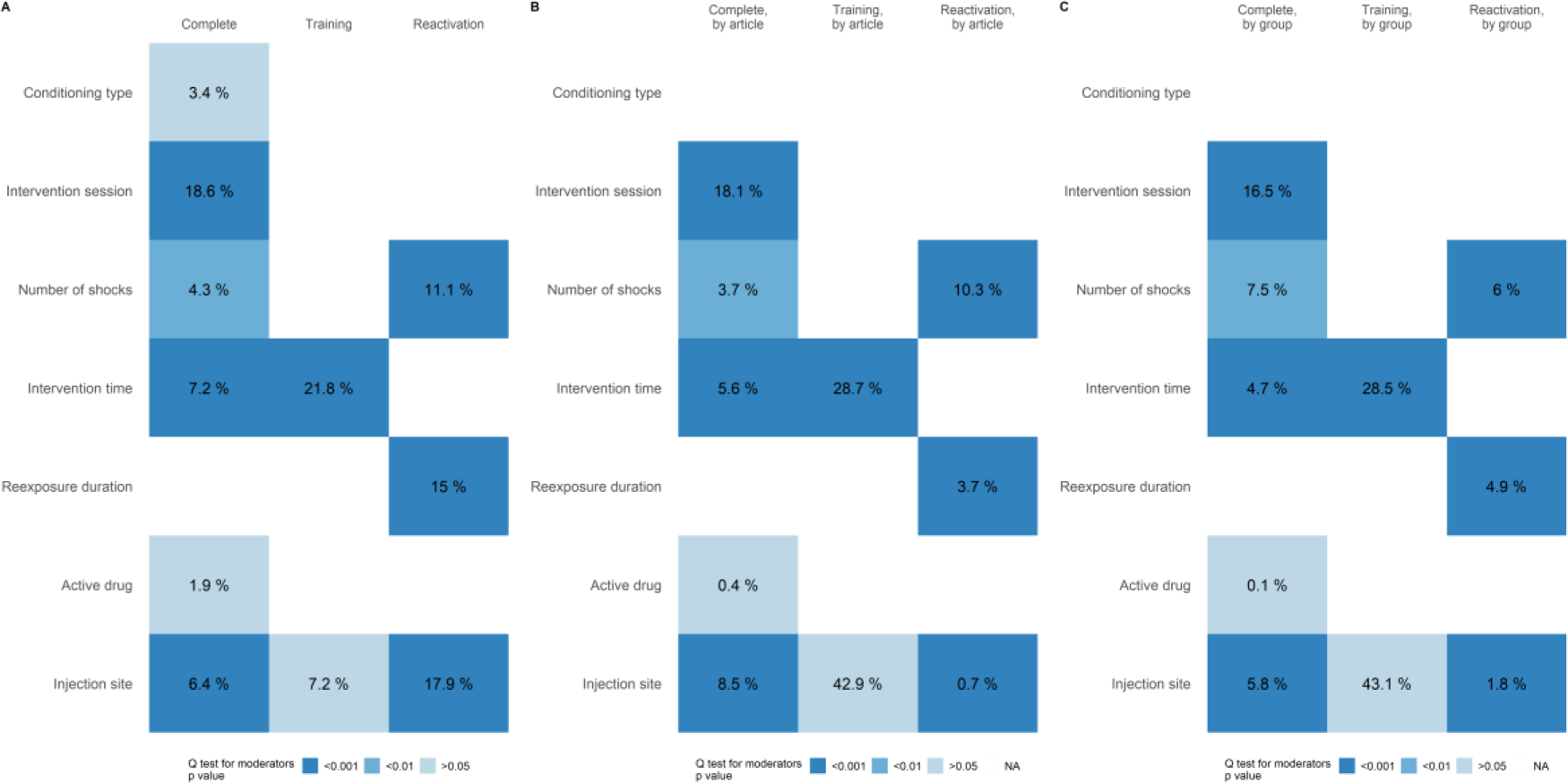
Summary of multivariable meta-regression models. Each column represents a dataset and lines are variables selected in the best model for each scenario. Cell colors represent the p value for the Q test for each moderator in the meta-regression and numbers within are the decomposed R^2^ value for the moderator (see Methods). Reexposure duration is based on z-scored values for reexposure to tone (measured in numbers) and to context (measured in minutes), based on the mean and standard deviation for each type of conditioning. **(A)** Two-level models. **(B)** Three-level models, accounting for nesting of experiments within articles. **(C)** Three-level models, accounting for nesting of experiments within research groups.

## Discussion

In this study, we explored different meta-analysis methods to analyze the effect of protein synthesis inhibitors on fear conditioning. Beyond the results, this effort also leads to some insights on how to apply existing meta-analysis methods to discovery science, which presents challenges to data synthesis.

### Effect of protein synthesis inhibitors on consolidation, reconsolidation and extinction

One particular focus was to investigate different ways to aggregate studies. Combining them by intervention session and by injection site showed a very robust effect for interventions in the three most studied sites (systemic injections, hippocampus and amygdala) on training and reconsolidation. This is unsurprising, as the requirement of protein synthesis for consolidation has been well accepted for decades^115^. More recently, protein synthesis inhibitors also became the paradigmatic intervention to demonstrate the existence of reconsolidation^11,116^.

On the other hand, the effect of protein synthesis inhibition on fear extinction was not as robust. Findings on the role of protein synthesis in extinction, in fear conditioning and beyond^117,118^, were not consensual from the start^112^. Moreover, when compared to reconsolidation, much less evidence on the topic is available. Thus, our meta-analysis found a significant effect only for systemic interventions. That said, most null effects on extinction from amygdala interventions come from a single study^90^, whereas the hippocampus analysis is heavily influenced by a study which, although investigating extinction according to the authors, would be considered a reconsolidation experiment by most standards^106^. Our meta-analysis thus suggests that it would be worthwhile to conduct further investigations into the role of protein synthesis in extinction in both brain structures.

Even though the training and reconsolidation data are robust on aggregate, there is great heterogeneity among individual experiments for virtually all meta-analyses performed. When articles were included as an additional level, most of this heterogeneity was observed between different experiments within studies, perhaps because behavioral neuroscience articles typically include experiments as tests of boundary conditions or negative controls. This presents a challenge for meta-analysis, as selection criteria will commonly lead to inclusion of many of these experiments, which are likely to bias effect size estimates downwards.

Aggregating studies from all injection sites or from interventions in training and reactivation did not cause much of an increase in heterogeneity, perhaps because it was already large to start with. As the increased sample sizes obtained provided greater power for identifying moderators, we used this approach for our meta-regressions, in a decision that was not prespecified in the original protocol.

### Moderators and boundary conditions

Some of the included moderators provide sanity checks of whether meta-regression captures consensual knowledge. It is widely accepted, for instance, that the effect of protein synthesis inhibitors is time-dependent, occurring when they are administered up to a few hours after learning^110,119^. Accordingly, in the training dataset, we found the time between training and drug administration to be a significant moderator in all models tested.

Similarly, it is well accepted that the occurrence of reconsolidation or extinction depends on the degree of mismatch between the training session and the reactivation, which is classically modulated by varying the number of CS presentations^120,121^ or the duration of reexposure^13,122^. Our measure of reexposure duration, which incorporated both of these manipulations, was indeed found to be a significant moderator in the reactivation dataset. Interestingly, however, it had a much smaller effect size when research group was included as a level in the analysis.

Other postulated boundary conditions for reconsolidation, however, were not observed as clearly. Many studies suggest that training strength can determine the occurrence of reconsolidation, extinction or neither^13,123,124^. This relation, however, is not linear: while very low training strength favors extinction rather than reconsolidation^123^, very strong memories have been suggested to be more resistant to destabilization^124^. Contrary to that, we found that a higher number of shocks led to a greater effect of protein synthesis inhibitors; that said, this effect was largely driven by a few experiments with a very large number of shocks. In contrast, shock intensity did not appear to be a relevant moderator, contradicting studies where administration of glucocorticoids after reactivation impaired retrieval only when stronger shocks were used^125^.

Another proposed boundary condition for reconsolidation is memory age^126^, but this has not been consistent across studies^49^ and may depend on reexposure conditions^13^. Our meta-analysis did not find an influence of memory age on the effect of protein synthesis inhibitors on reactivation; This runs counter to widely cited studies showing that older memories are resistant both to protein synthesis inhibitors^13,85,126^ and other drugs^127^. Nevertheless, it is worth pointing out that our model does not include interactions between moderators; thus, an effect could have been obscured by confounding. It is also possible that this effect is limited to very remote memories: studies with memories over 20 days-old had smaller effects in our sample, but this was offset by large effects for those between 1- and 20 days-old.

Type of conditioning (tone vs. contextual) was consistently detected as a significant moderator for reconsolidation effects in our univariable models, mostly driven by frequent null results in contextual conditioning experiments. Importantly, however, the effect we found was nearly absent from multivariable models, perhaps due to its interaction with site of injection, as both types of conditioning tend to be studied with interventions in different brain structures.

Although one would expect that drug dose would be a relevant moderator for a pharmacological intervention, this was not the case in our meta-analysis, probably due to the low variability in the doses used for each drug, species and route of administration (which were used as normalizers in the analysis). This is likely due to the fact that, as protein synthesis inhibitors have been in study for many years, effective doses are well established and rarely varied in more recent studies.

The changes in the relative importance of different moderators when multivariable and multilevel models are used underscore the importance of these methods to minimize confounding. Interestingly, the effect of intervention time, perhaps our best-established moderator, became stronger when multiple levels were added. Conversely, seemingly spurious factors such as region of origin or presence of a conflict-of-interest statement disappeared when research group was used as a level.

### Heterogeneity and replication failures

Even after moderators and additional levels were included, however, much heterogeneity remained, suggesting that classic boundary conditions are not sufficient to explain discrepancies in the literature. This echoes recent failures in replicating post-retrieval amnesia studies, despite the use of similar protocols^14,128^. Minor differences in reactivation protocols, animal traits^129^, housing conditions^130^ or interactions between moderators could explain this additional heterogeneity. Nevertheless, it is also possible that heterogeneity and replication failures are due to biases in the literature.

In this sense, the prevalence of randomization and sample size calculations in our sample was low, in accordance with previous surveys of preclinical studies^131^. Encouragingly, however, we found a higher level of blinding than reported in other areas^131^, mostly due to the use of automated freezing detection. Another interesting finding is the near absence of studies using female animals, which has also been pointed out in previous studies of fear conditioning^132^ and deviates markedly from current recommendations^133,134^.

Another explanation for divergences in the literature is random error, which can lead to effect size vibration across statistical thresholds, particularly when power is low. We found the average power of individual studies to detect the respective meta-analytical effect size estimate to be 63%, which is in line with a previous estimate for fear conditioning^132^. This means that around one third of studies should fail to find significant differences between groups even when a real effect exists, suggesting that lack of replication based on statistical significance criteria is expected to occur frequently.

Interestingly, we found little evidence of publication bias, in contrast to what was found by Schroyens et al (2021) for anisomycin, propranolol, MK-801 and midazolam in contextual fear reconsolidation. The evidence for publication bias in their study is weaker for anisomycin, however, suggesting that it may be a smaller problem in the literature on protein synthesis inhibitors. Nevertheless, our ability to detect publication bias may have been limited by experiments testing boundary conditions or negative controls, as bias may act on the opposite direction in these cases (i.e. suppressing statistically significant results). Publication bias assessments can also be limited by true heterogeneity^23^, which is quite large in our sample.

### Conclusion and future remarks

In conclusion, we have found that meta-analysis and meta-regression can capture some well-established findings in the fear conditioning literature. This includes a robust effect of protein synthesis inhibitors on consolidation and reconsolidation and its dependence on moderators such as timing of post-training interventions and the degree of reexposure in reactivation experiments. Although our analysis only includes studies published up to 2018, these main effects are robust and would be unlikely to change much if more recent studies were included.

Nevertheless, we also found some knowledge gaps, particularly on the effects of protein synthesis inhibition in specific brain structures on extinction. Moreover, not every boundary condition commonly mentioned in the literature was confirmed, including factors such as number of shocks, shock intensity and memory age. This may indicate that the evidence for some of these is more controversial than suggested in the literature, or alternatively that meta-regression might fail to capture real effects due to noise, non-linearity or interactions between variables (which were not assessed in our analyses). This leads to a conundrum – when meta-regression doesn’t agree with individual studies that test a particular moderator, which of them should we trust?

Unfortunately, there is no easy answer. On one hand, individual studies are typically underpowered and subject to many biases. On the other, meta-regression findings are observational and inevitably prone to confounding. The use of multilevel models and multivariable regression can mitigate this problem but will not fully eliminate it. Moderators found by meta-regression should thus be considered tentative and ideally tested in empirical studies designed for this purpose. Although data synthesis can be useful for evaluating the primary literature, it incorporates its biases, and cannot fully substitute for confirmatory, high-powered replication studies^135^ – which should be regarded as the preferred option for solving controversies between meta-analyses and primary studies.

A final advantage of empirically testing findings from meta-analysis is to compare the predictions of different types of models in terms of their accuracy. Although we argue in favor of using multilevel and multivariable models, these can be unfeasible or inaccurate when the number of studies is low. It is thus important to evaluate how much is lost by traditional two-level models, which in our work were in general agreement with multilevel ones.

All of these issues will only be resolved when systematic iteration between data synthesis and experimentation becomes more common in discovery science. This requires engaging researchers in systematic reviews, as well as making these more efficient through automated tools^136^ and better structuring of primary data. It also involves building up infrastructure for confirmatory research that can validate meta-analytical findings empirically^135^. Ultimately, such interplay may produce a virtuous cycle that can improve the robustness of basic science beyond what can be achieved with the current model of isolated experiments and narrative synthesis.

## Acknowledgments

We’d like to thank Daniel Menezes for assistance in the early stages of screening, Thiago Moulin for assistance with co-authorship graphs, Kleber Neves for general assistance with analyses in R and Jonathan Lee for opinions on the manuscript.

Furthermore we thank the recommender and reviewers from Peer Community In Neuroscience for their assessments and suggestions to improve this manuscript.

