## Supplementary Figures for "A meta-analysis of the effect of protein synthesis inhibitors on rodent fear conditioning"

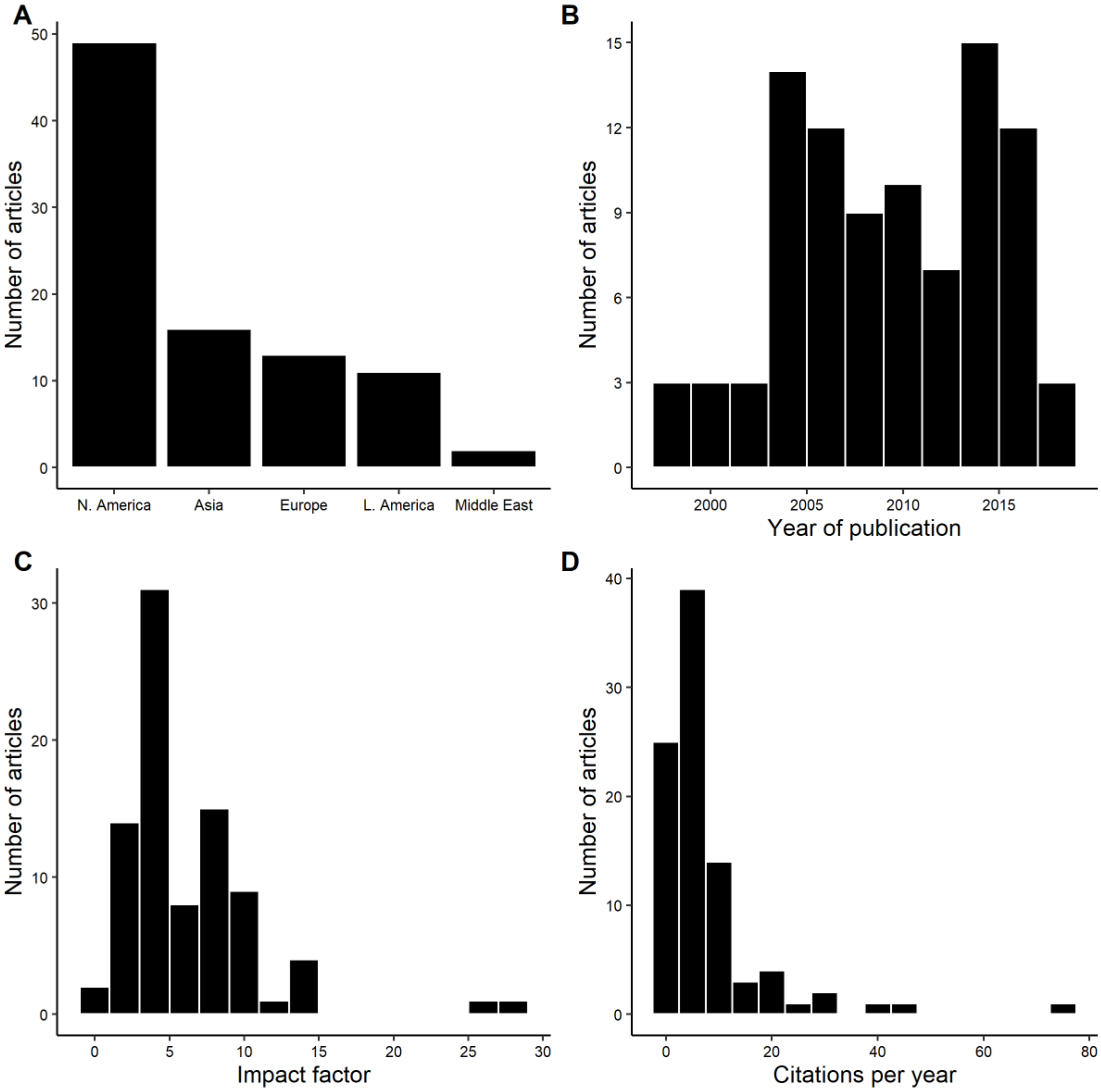


**Suppl. Figure 1 - General features of included articles.** (A) Distribution of articles by region of origin. (B) Distribution of publication year. For articles with different years for online and print publication, the former was considered. (C) Distribution of impact factors. (D) Distribution of citations per year. N. America, North America; L. America, Latin America.

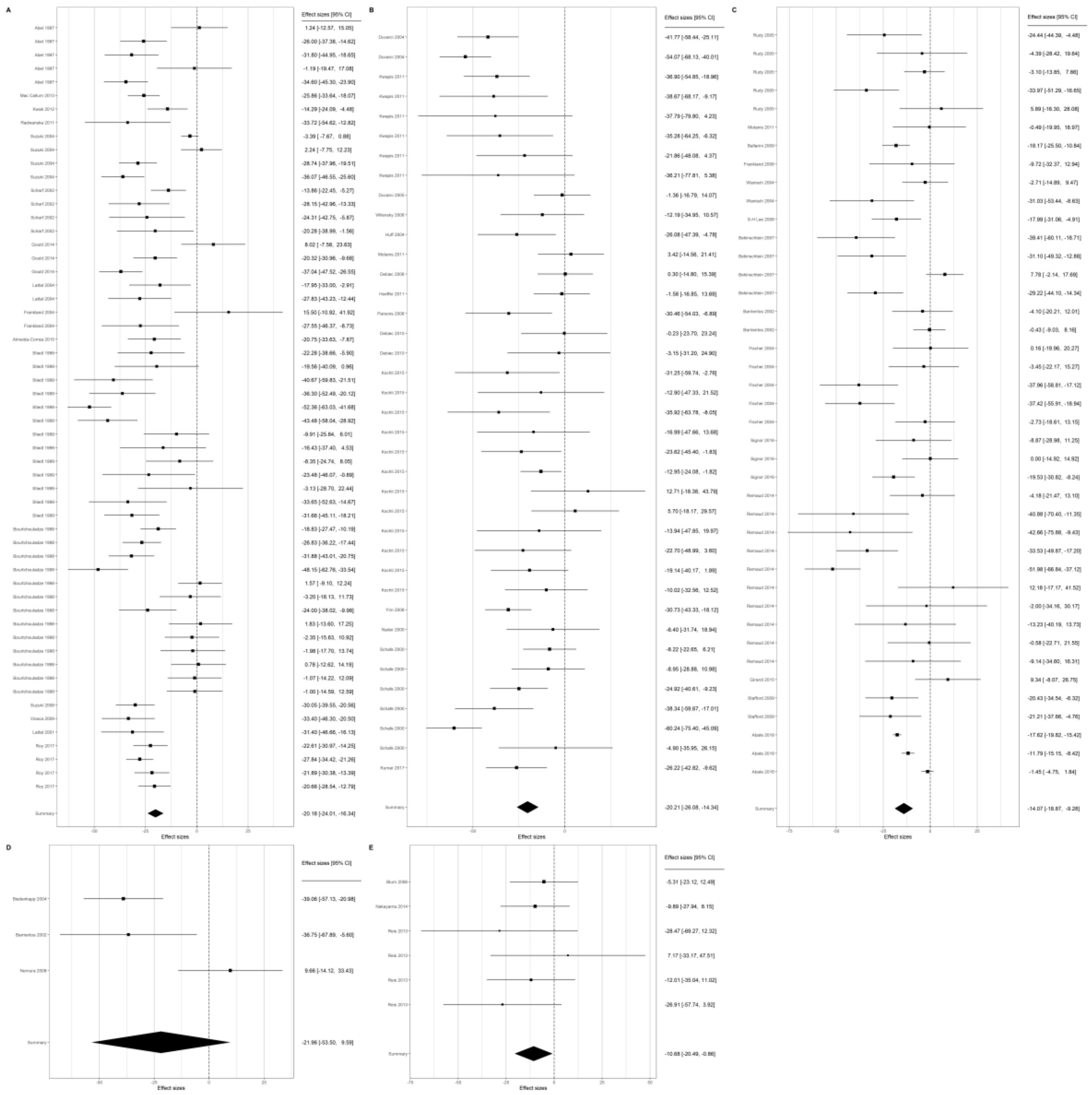

**Suppl. Figure 2 – Forest plots of interventions in training.** Forest plots show study source, effect size and confidence intervals of included studies. Effect size is expressed in the x axis as absolute mean difference in freezing, with the dashed line representing absence of effect. Square size is proportional to the weight of individual studies, while diamonds show summary estimates along with their confidence intervals. (A) Systemic interventions. (B) Intra-amygdala interventions. (C) Intra-hippocampus interventions. (D) Intracerebroventricular interventions. (E) Intra-prefrontal cortex interventions.

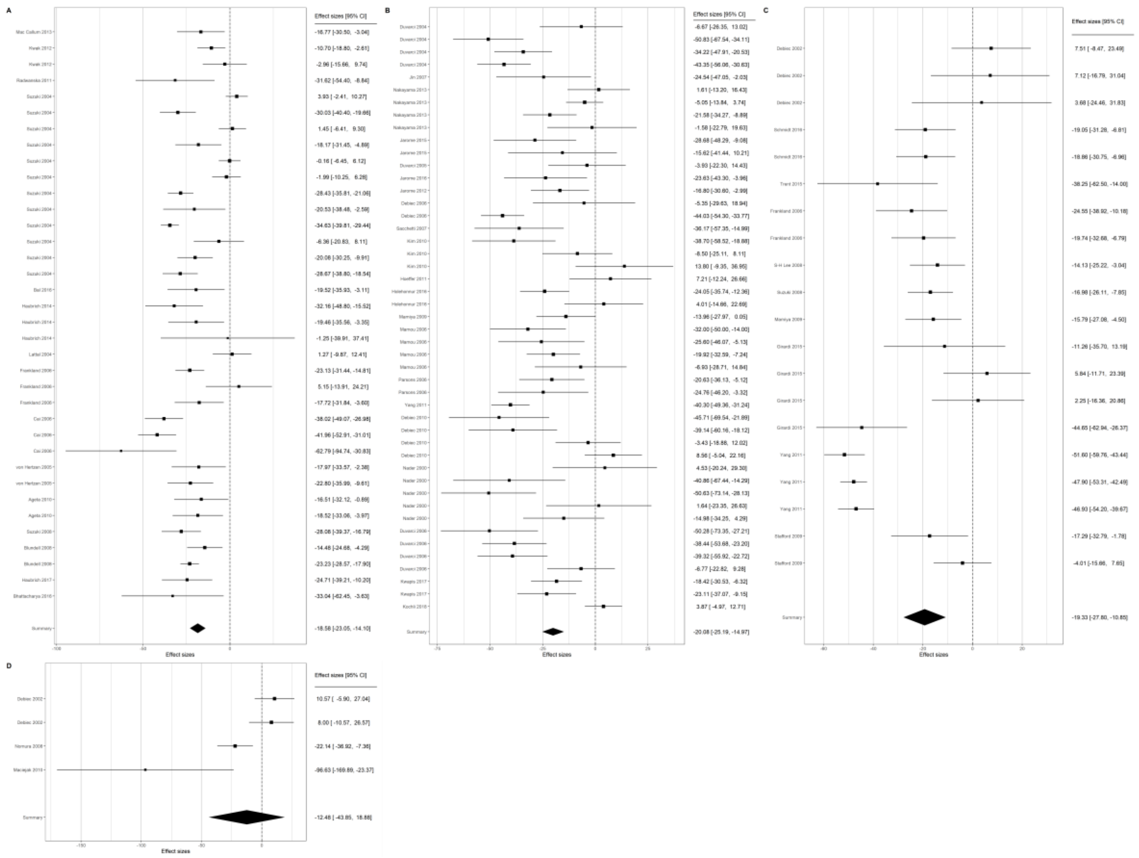

**Suppl. Figure 3 – Forest plots of interventions in reconsolidation.** Forest plots show study source, effect size and confidence intervals of included studies. Effect size is expressed in the x axis as absolute mean difference in freezing, with the dashed line representing absence of effect. Square size is proportional to the weight of individual studies, while diamonds show summary estimates along with their confidence intervals. (A) Systemic interventions. (B) Intra-amygdala interventions. (C) Intra-hippocampus interventions. (D) Intracerebroventricular interventions.

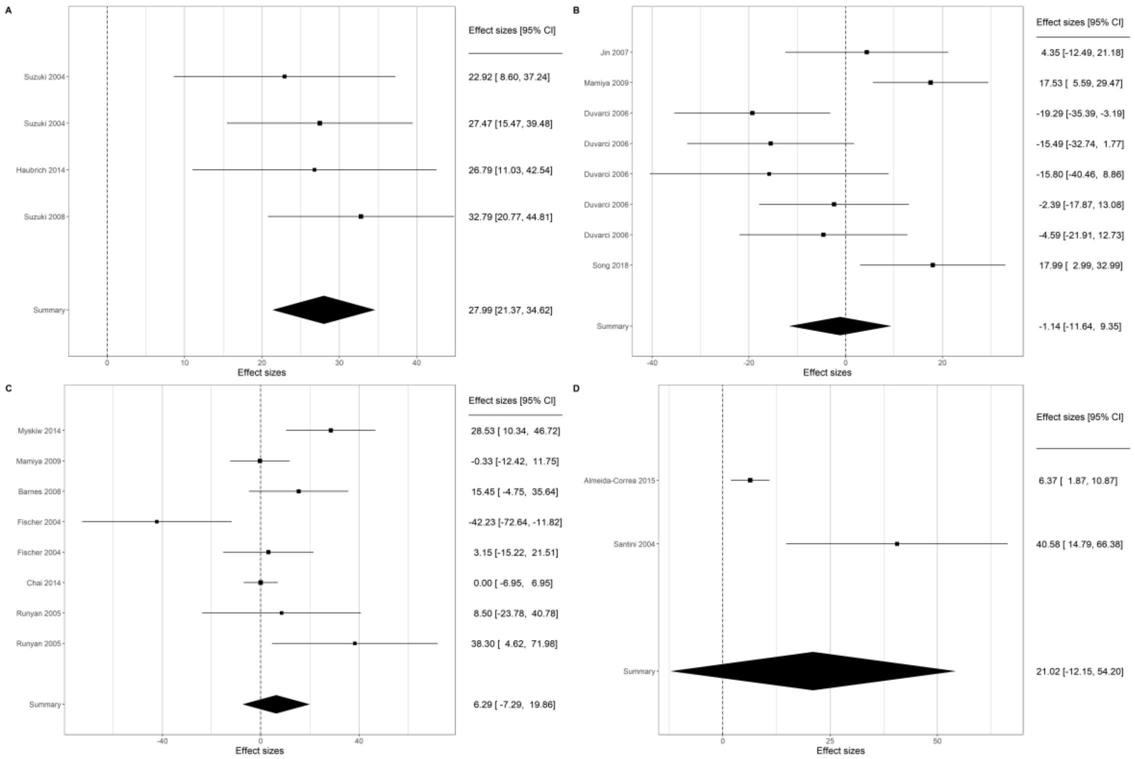


**Suppl. Figure 4 – Forest plots of interventions on extinction.** Forest plots show study source, effect size and confidence intervals of included studies. Effect size is expressed in the x axis as absolute mean difference in freezing, with the dashed line representing absence of effect. Square size is proportional to the weight of individual studies, while diamonds show summary estimates along with their confidence intervals. (A) Systemic interventions. (B) Intra-amygdala interventions. (C) Intra-hippocampus interventions. (D) Intracerebroventricular interventions.

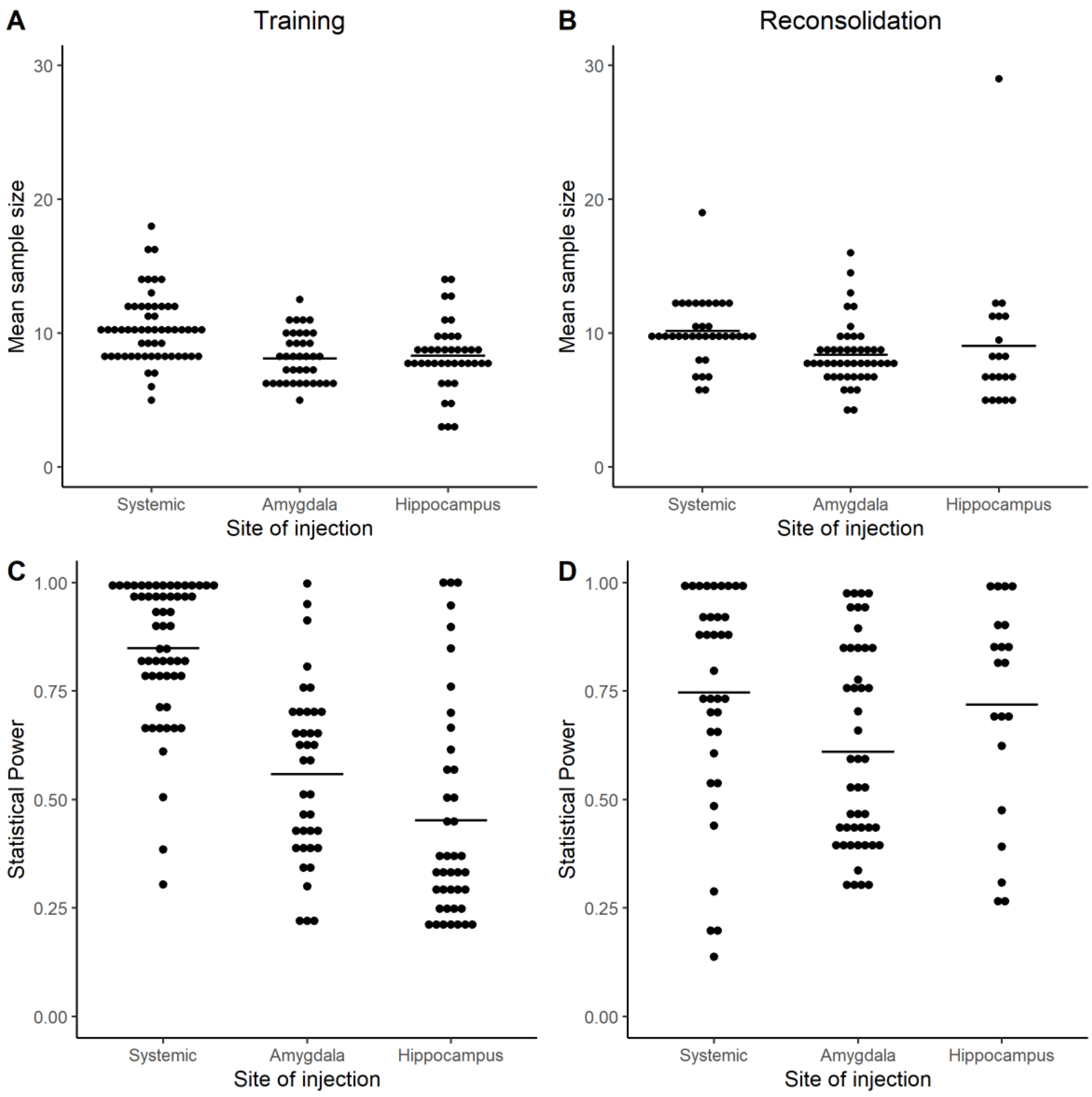


**Suppl. Figure 5 – Statistical power and sample size distributions. (A)** Sample sizes of experiments with interventions on training as originally reported. All: 9.1 ± 2.6, Systemic: 10.2 ± 2.6, Amygdala: 8.1 ± 1.9, Hippocampus: 8.3 ± 2.5. **(B)** Sample sizes of experiments with interventions on reconsolidation as originally reported. All: 9.1 ± 3.2, Systemic: 10.2 ± 2.3, Amygdala: 8.4± 2.3, Hippocampus: 9.1 ± 5.3. **(C)** Statistical power of experiments with interventions on training, calculated based on the sample sizes described in panel A and effect sizes presented in Table 2 All: 0.65 ± 0.27, Systemic: 0.85 ± 0.16, Amygdala: 0.56 ± 0.20, Hippocampus: 0.45 ± 0.25. **(D)** Statistical power of experiments with interventions on reconsolidation. All: 0.68 ± 0.25, Systemic: 0.75 ± 0.25, Amygdala: 0.61 ± 0.23, Hippocampus: 0.72 ± 0.25. All values are mean ± standard deviation. Lines in figures represent mean values for each group.

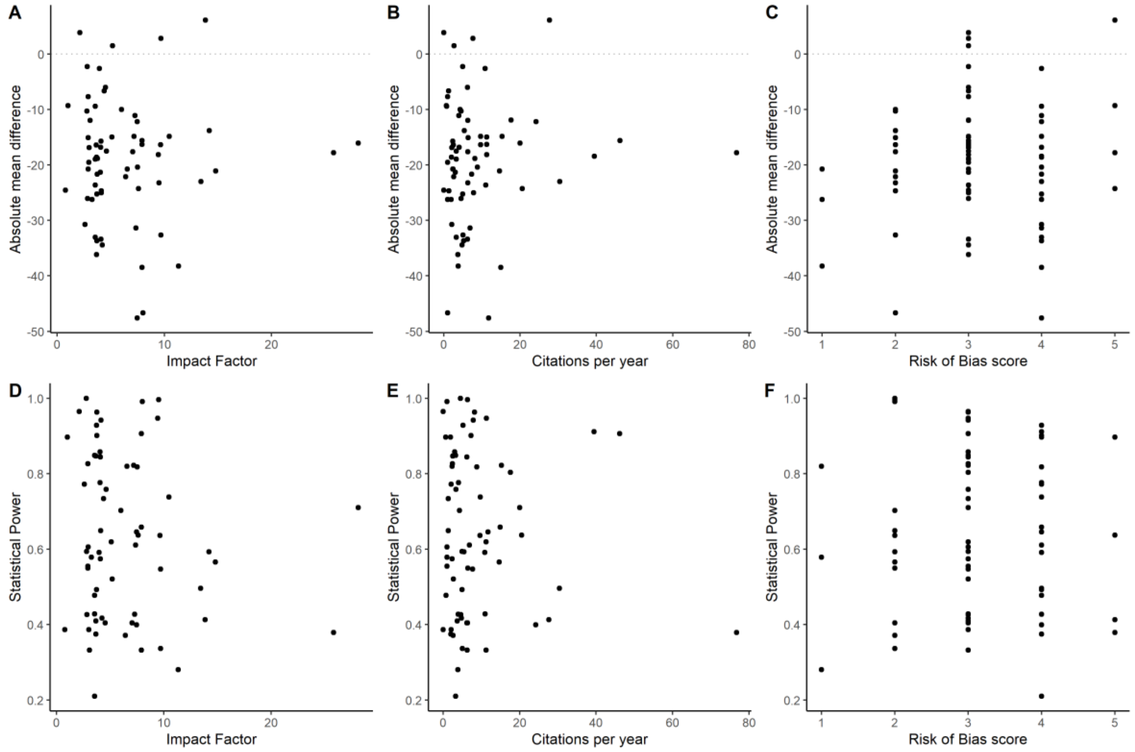


**Suppl. Figure 6 – Correlations between effect size and statistical power and article-level variables.** X axes show impact factor, citations/year and risk of bias for individual articles, while Y axes show the mean effect size or statistical power for the mean of all comparisons within individual articles. Risk of bias scores are calculated as described in the Methods section. **(A)** Correlation between mean effect size and impact factor. Spearman's ρ = -0.04, p = 0.72, n = 66. **(B)** Correlation between mean effect size and citations per year. Spearman's ρ = 0.05, p = 0.66, n = 69. **(C)** Correlation between mean effect size and risk of bias score. Spearman's ρ = 0.03, p = 0.80, n = 69. **(D)** Correlation between mean statistical power and impact factor. Spearman's ρ = -0.07, p = 0.56, n = 66. **(E)** Correlation between mean statistical power and citations per year. Spearman's ρ = 0.002, p = 0.99, n = 69. **(F)** Correlation between mean statistical power and risk of bias score. Spearman's ρ = -0.008, p = 0.95, n = 69.

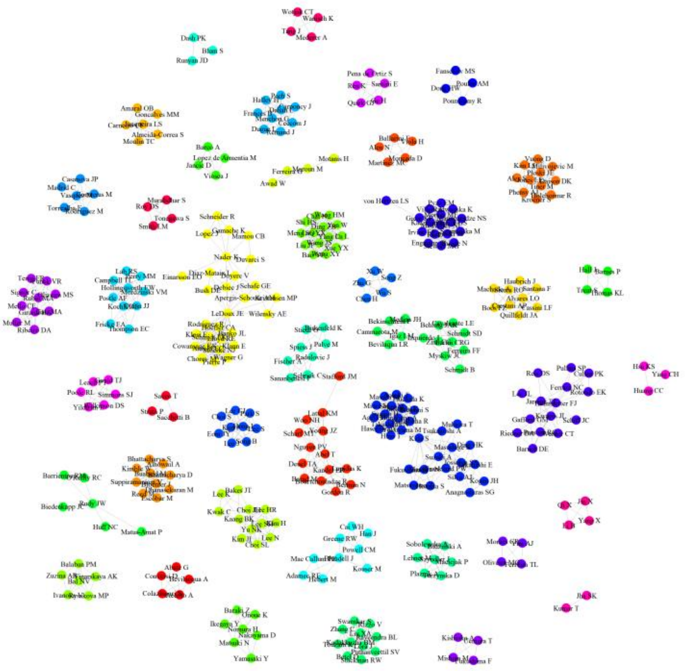

**Suppl. Figure 7 – Co-authorship network of articles selected for inclusion.** Each node in the graph represents and author, while edges represent co-authorships in the papers included in the meta-analysis. Colors and positions on the map are arbitrary. Forty-two groups were identified, with modularity = 0.94.

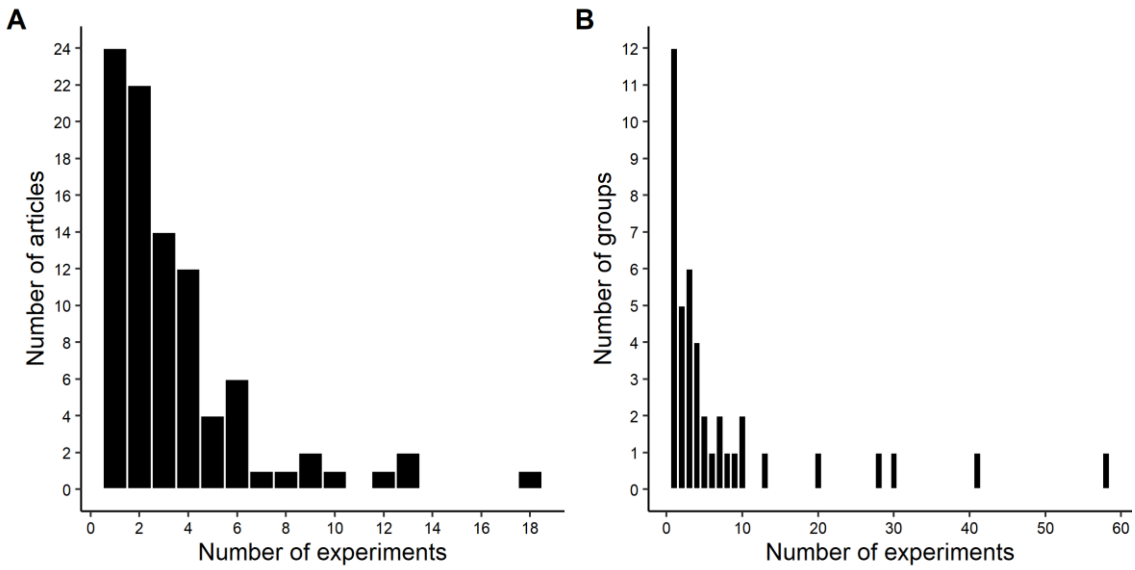


**Suppl. Figure 8 – Distribution of experiments among clusters for each level of analysis.** (A) Histogram showing the distribution of number of experiments per article. (B) Histogram showing the distribution of number of experiments per research group.

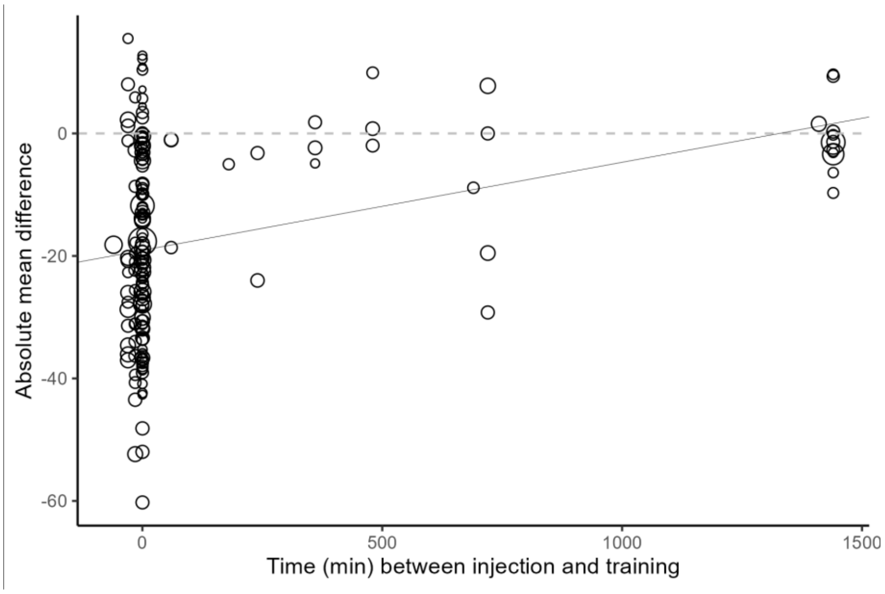


**Suppl. Figure 9 – Impact of time between injection and training on effect size.** Only interventions targeting the training session are included. Each circle represents a comparison between two groups and its size is inversely proportional to the comparison’s variance. Injections described as immediately before or immediately after the training session were coded as -0.1 and 0.1 min, respectively. Black line represents the meta-regression model. N = 166 comparisons.

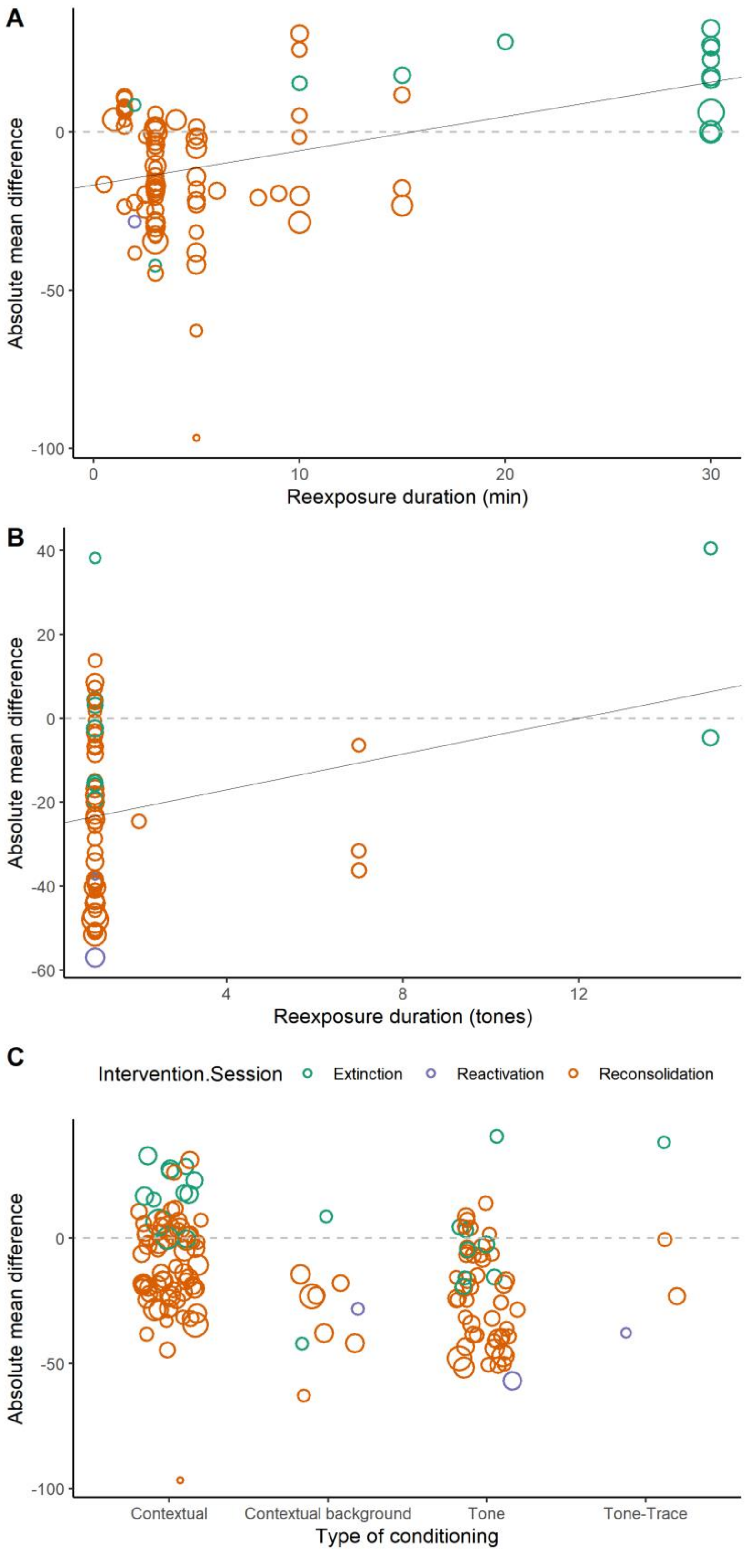


**Suppl. Figure 10 – Impact of conditioning type and reexposure duration on effect size.** Only interventions targeting the reactivation session (i.e. extinction, reconsolidation or reexposure) are included. In all panels, each circle represents a comparison between two groups and its size is inversely proportional to the comparison's variance. Black lines represent the meta-regression models. **(A)** Duration of context reexposure (in minutes). N = 14 (Extinction), 1 (Reactivation), 70 (Reconsolidation). **(B)** Duration of reexposure to the conditioned tones (in number of tones. N = 9 (Extinction), 2 (Reactivation), 47 (Reconsolidation). **(C)** Type of conditioning. Contextual: N = 12 (Extinction), 65 (Reconsolidation).

Contextual background: N = 2 (Extinction), 1 (Reactivation), 7 (Reconsolidation). Tone: N = 8 (Extinction), 1 (Reactivation), 45 (Reconsolidation). Tone-trace: N = 1 (Extinction), 1 (Reactivation), 2 (Reconsolidation).

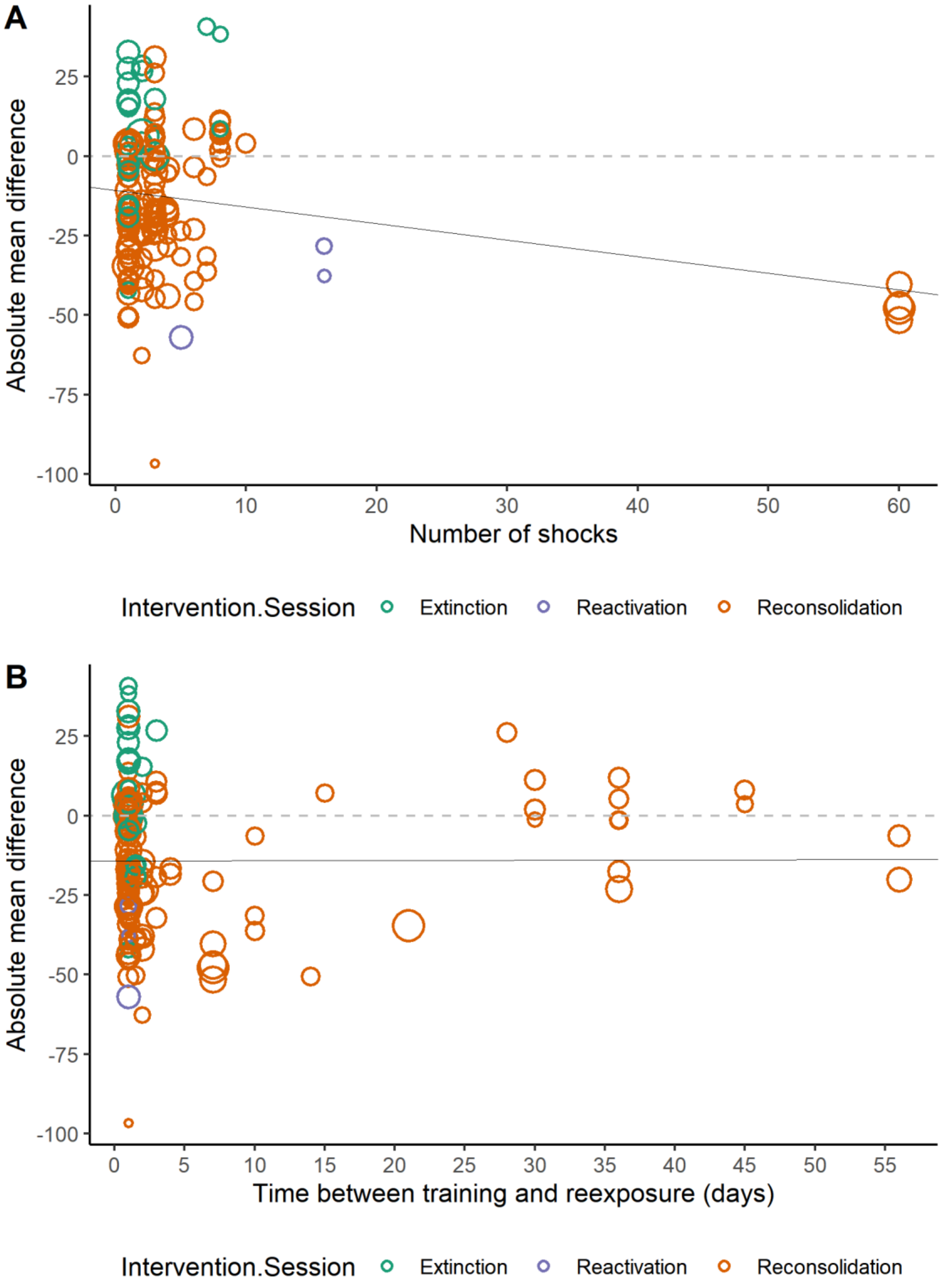


**Suppl. Figure 11 – Impact of number of shocks and memory age on effect size.** Only interventions targeting the reactivation session (i.e. extinction, reconsolidation or reexposure) are included. In all panels, each circle represents a comparison between two groups, and its size is inversely proportional to the comparison's variance. The black line represents results from meta-regression models. **(A)** Number of shocks presented during training. N = 23 (Extinction), 3 (Reactivation), 119 (Reconsolidation). **(B)** Time between training and reactivation session. N = 23 (Extinction), 3 (Reactivation), 119 (Reconsolidation).

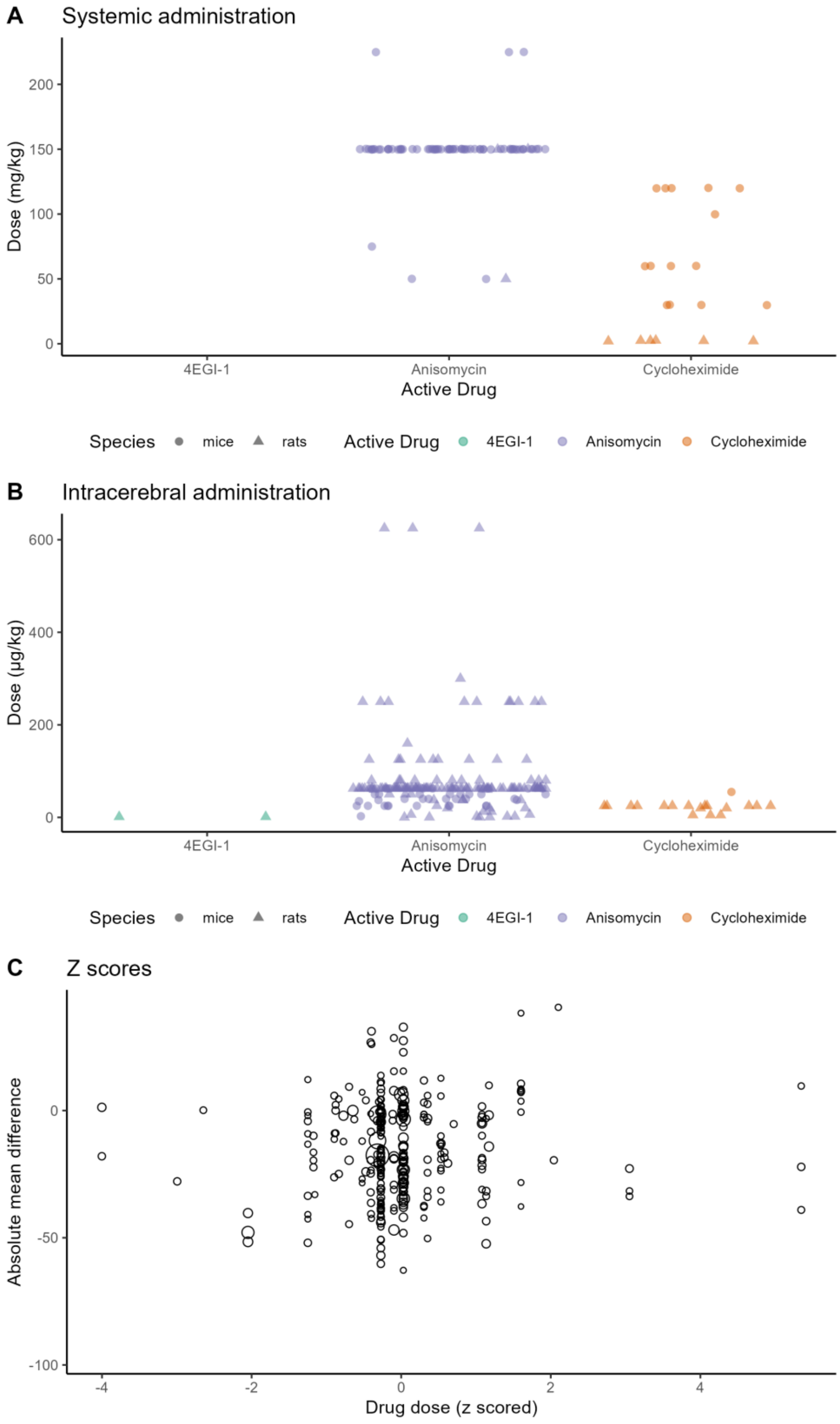


**Suppl. Figure 12 – Impact of drug dose on effect size.** All experiments are included, regardless of the targeted session (i.e. training or reexposure). **(A)** Distribution of drug doses for systemic interventions. **(B)** Distribution of drug doses for intracerebral interventions. **(C)** Correlation between z-scored doses (considering drug, route of administration and species) and effect sizes. Each circle represents a comparison between two groups, and its size is inversely proportional to the comparison's variance.

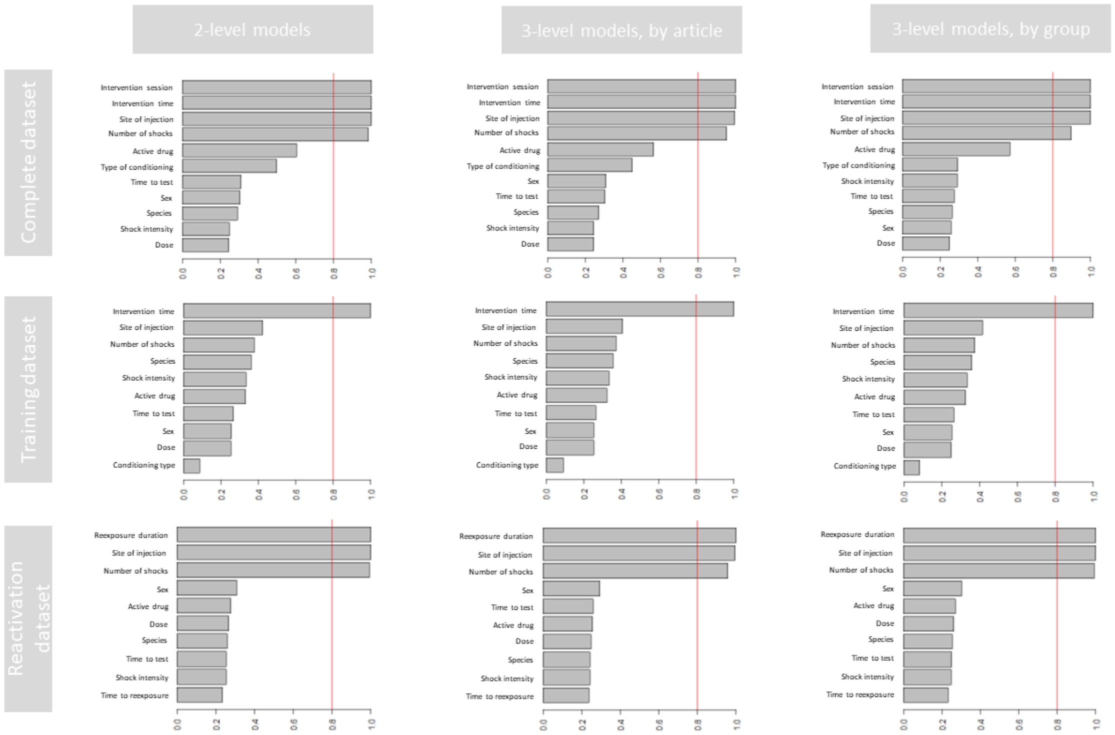


**Suppl. Figure 13 – Variable importance in multivariate models.** Each panel shows the model-averaged importance of terms across all tested models, as the sum of the weights for the models in which each variable appears. For the complete dataset (top row), 2,048 models were tested. For the training and reactivation datasets (middle and bottom rows, respectively), 1,024 models were tested. Red lines mark the 0.8 threshold, commonly used to distinguish variables of high importance. Reexposure duration is based on z-scored values for reexposure to tone (measured in number of tones) and to context (measured in minutes), based on the mean and standard deviation for each type of conditioning. Doses were also z-scored based on the mean and standard deviation of each drug and route of administration.
