## Supplementary Tables for "A meta-analysis of the effect of protein synthesis inhibitors on rodent fear conditioning"

**Suppl. Table 1 - Risk of bias measures.** Counts and percentages of articles reporting each of the items presented are based on the complete dataset (91 articles).

| | Randomization of allocation | Blinded or automated assessment | Sample size calculation | Statement of compliance with regulatory requirements | Statement on conflict of interest |
|---|---|---|---|---|---|
| **Number of articles (%)** | 3 (3.3%) | 61* (67%) | 2 (2.2%) | 84 (92.3%) | 28** (30.8%) |

*Of these, 29 reported manual assessment.

**Of these, 1 declared an existing conflict of interest.

**Suppl. Table 2 – Protocol features of comparisons included in the analyses.** For categorical variables, we show the count of comparisons per category, while for continuous variables we show the median and range (minimum and maximum). Percentages refer to the number of applicable protocols. "Contextual background" refers to cases where conditioning was performed with tone cues, but testing (and reexposure, where applicable) was performed for the context (without tone). Interventions described as immediately before or immediately after training or reexposure were coded as -0.1 min and 0.1 min, respectively.

| Protocol variable | Value | N (%) |
|---|---|---|
| Intervention session | Training | 168 (53.7%) |
| | Reconsolidation | 119 (38.0%) |
| | Extinction | 23 (7.3%) |
| | Reactivation | 3 (1.0%) |
| Site of injection | Systemic | 97 (31.0%) |
| | Amygdala | 94 (30.0%) |
| | Hippocampus | 71 (22.7%) |
| | Cerebral ventricles | 9 (2.9%) |
| | Prefrontal cortex | 9 (2.9%) |
| | Retrosplenial cortex | 8 (2.6%) |
| | Anterior cingulate cortex | 7 (2.2%) |
| | Striatum/Nucleus accumbes | 3 (1.0%) |
| | Auditory thalamus | 2 (0.6%) |
| | Frontal association cortex | 2 (0.6%) |
| | Infralimbic cortex | 2 (0.6%) |
| | Perirhinal cortex | 2 (0.6%) |
| | Prelimbic cortex | 2 (0.6%) |
| | Bed nuclei of stria terminalis | 1 (0.3%) |
| | Cerebellar vermis | 1 (0.3%) |
| | Medial geniculate nucleus | 1 (0.3%) |
| | Posterior insular cortex | 1 (0.3%) |
| | Primary/Secondary motor cortex | 1 (0.3%) |
| Type of conditioning | Contextual | 138 (44.1%) |
| | Tone | 103 (32.9%) |
| | Contextual background | 59 (18.8%) |
| | Tone-trace | 13 (4.2%) |
| Habituation | Reported | 76 (24.3%) |
| | Not reported | 237 (75.7%) |
| Handling | Reported | 83 (26.5%) |
| | Not reported | 230 (73.5%) |
| Number of footshocks | 2 (1 – 76) | 313 (100%) |
| Footshock intensity | 0.75 (0.3 – 2) mA | 310 (99.0%) |
| Time between drug administration and training/reexposure | 0.1 h* (1 h before – 24 h after) | 303 (96.8%) |
| Time between training and reexposure | 24 h (24 h – 56 days) | 145 (100%) |
| Reexposure duration | Context: 3 min (30 sec – 30 min) | 84 (96.5%) |

| | | |
|---|---|---|
| | Tone: 1 tone (1 – 15) | 58 (100%) |
| Time between intervention session and test | 24 h (30 min – 36 days) | 312 (99.7%) |
| Species | Rats<br>Mice | 167 (53.4%)<br>146 (46.6%) |
| Sex | Male<br>Both<br>Not informed | 293 (93.6%)<br>18 (5.8%)<br>2 (0.6%) |
| Mean age | 10 (4 – 22) weeks | 122 (39.0%) |
| Housing | 1 (1 – 6) animals per cage | 242 (77.3%) |
| Active drug | Anisomycin<br>Cycloheximide<br>4EGI-1 | 270 (86.3%)<br>41 (13.1%)<br>2 (0.6%) |
| Dose | Anisomycin (systemic): 150 (50 – 225) mg/kg<br>Anisomycin (intracerebral): 62.5 (0.08 – 625) µg<br>Cycloheximide (systemic): 45 (2.2 – 120) mg/kg<br>Cycloheximide (intracerebral): 25 (5 – 55) µg<br>4EGI-1 (intracerebral): 1.25 µg | 73 (94.8%)<br>180 (93.3%)<br>20 (100%)<br>18 (85.7%)<br>2 (100%) |

**Suppl. Table 3 – Intracerebral sites of injection with less than 3 articles per intervention session.** As defined in our analysis protocol, at least 3 articles had to be available for meta-analyses to be performed.

| Site of injection | Intervention session | Nº articles |
|---|---|---|
| Anterior cingulate cortex | Training<br>Reconsolidation | 2<br>2 |
| Auditory thalamus | Training | 1 |
| Bed nuclei of the stria terminalis | Training | 1 |
| Cerebellar vermis | Reconsolidation | 1 |
| Frontal association cortex | Training | 1 |
| Infralimbic cortex | Reconsolidation | 1 |
| Medial geniculate nucleus | Training | 1 |
| Perirhinal cortex | Training<br>Reconsolidation | 1<br>1 |
| Posterior insular cortex | Training | 1 |
| Prefrontal cortex | Training<br>Reconsolidation<br>Extinction | 3<br>2<br>1 |
| Prelimbic cortex | Training | 1 |
| Primary/secondary motor cortex | Reconsolidation | 1 |
| Retrosplenial cortex | Training | 1 |
| Striatum/nucleus accumbens | Training | 1 |

**Suppl. Table 4 - Summary of meta-analyses using standardized mean differences.** Effect sizes are expressed as standardized mean differences (Hedges' g) between PSI-treated and control groups in the test session. Sample size is the number of experiments. p values refer to main effect comparisons in random-effects meta-analyses (4th column) and Q-tests for heterogeneity (6th column). i.c.v.: intracerebroventricular, CI: confidence interval.

| **Dataset** | **Sample size** | **Effect size [95% CI]** | **Meta-analysis p-value** | **$I^2$** | **Q-test p-value** |
|---|---|---|---|---|---|
| Training, systemic | 57 | -1.3 [-1.6, -1.1] | $5.6x10^{-20}$ | 70.3% | $8.1x10^{-16}$ |
| Training, amygdala | 36 | -0.9 [-1.2, -0.6] | $7.2x10^{-11}$ | 36.9% | 0.007 |
| Training, hippocampus | 41 | -0.76 [-1.0, -0.5] | $1.9x10^{-8}$ | 48.0% | $1.7x10^{-5}$ |
| Training, i.c.v. | 3 | -0.9 [-2.3, 0.5] | 0.211 | 78.6% | 0.009 |
| Training, prefrontal cortex | 6 | -0.5 [-0.9, 0.03] | 0.065 | 0% | 0.826 |
| Reconsolidation, systemic | 36 | -1.3 [-1.6, -0.9] | $4.2x10^{-12}$ | 77.8% | $1.7x10^{-14}$ |
| Reconsolidation, amygdala | 47 | -1.0 [-1.3, -0.7] | $5.1x10^{-12}$ | 69.3% | $1.1x10^{-10}$ |
| Reconsolidation, hippocampus | 20 | -1.5 [-2.4, -0.6] | $8.5x10^{-4}$ | 92.1% | $1.6x10^{-11}$ |
| Reconsolidation, i.c.v. | 4 | -0.3 [-1.4, 0.7] | 0.501 | 72.6% | 0.012 |
| Extinction, systemic | 4 | 1.8 [1.2, 2.3] | $5.3x10^{-11}$ | 0% | 0.733 |
| Extinction, amygdala | 8 | -0.1 [-0.9, 0.7] | 0.825 | 62.5% | 0.119 |
| Extinction, hippocampus | 8 | 0.3 [-0.2, 0.8] | 0.264 | 46.1% | 0.063 |
| Extinction, i.c.v. | 2 | 1.4 [0.6, 2.1] | $3.1x10^{-4}$ | 0% | 0.934 |
| Reactivation, systemic | 40 | -1.0 [-1.4, -0.6] | $9.1x10^{-6}$ | 86.7% | $3.7x10^{-61}$ |
| Reactivation, amygdala | 56 | -1.0 [-1.3, -0.7] | $2.8x10^{-10}$ | 74.6% | $3.5x10^{-16}$ |
| Reactivation, hippocampus | 30 | -0.9 [-1.5, -0.3] | 0.003 | 87.9% | $2.3x10^{-15}$ |
| Reactivation, i.c.v. | 6 | 0.2 [-0.7, 1.2] | 0.651 | 79.5% | $2.0x10^{-4}$ |
| Complete | 311 | -0.9 [-1.0, -0.7] | $1.2x10^{-40}$ | 74.5% | $8.1x10^{-93}$ |

**Suppl. Table 5 – Trim-and-fill analyses.** Columns 2 and 3 include the same results presented in Table 3. p values refer to main effects in meta-analyses (5th and 7th columns) after correction for publication bias and to the estimated number of missing studies for the $R_0$ method (8th column) against a null hypothesis of 0.

| Dataset | Original estimate (p-value) | Sample size | $L_0$ Method | | $R_0$ Method | | |
|---|---|---|---|---|---|---|---|
| | | | Estimated # of missing studies | Bias-corrected estimate (p-value) | Estimated # of missing studies | Bias-corrected estimate (p-value) | p-value |
| Training, systemic | -20.18 ($6.4 \times 10^{-25}$) | 57 | 0 | NA | 0 | NA | 0.50 |
| Training, amygdala | -20.37 ($8.9 \times 10^{-12}$) | 38 | 0 | NA | 1 | -18.82 ($2.7 \times 10^{-9}$) | 0.25 |
| Training, hippocampus | -14.07 ($8.9 \times 10^{-9}$) | 41 | 0 | NA | 5 | -10.88 ($5.1 \times 10^{-5}$) | 0.02 |
| Reconsolidation, systemic | -18.58 ($4.2 \times 10^{-16}$) | 36 | 5 | -15.58 ($1.6 \times 10^{-10}$) | 0 | NA | 0.50 |
| Reconsolidation, amygdala | -19.93 ($2.0 \times 10^{-14}$) | 47 | 0 | NA | 0 | NA | 0.50 |
| Reconsolidation, hippocampus | -19.33 ($7.8 \times 10^{-6}$) | 20 | 0 | NA | 4 | -11.45 (0.022) | 0.03 |

**Suppl. Table 6 – Excess significance tests for meta-analyses.** Mean statistical power used for each meta-analysis is presented to provide a comparison to that displayed in Suppl. Fig. 5, as it was calculated for meta-analytical estimates using unadjusted sample sizes. The p-value for excess of significance is from a chi-squared test.

| Dataset | Sample size | Mean statistical power (± S.D.) | Expected # of positive studies | Observed # of positive studies | p-value |
|---|---|---|---|---|---|
| Training, systemic | 57 | 0.73 ± 0.10 | 41.8 | 41 | 0.819 |
| Training, amygdala | 38 | 0.56 ± 0.12 | 21.3 | 21 | 0.919 |
| Training, hippocampus | 41 | 0.52 ± 0.16 | 21.3 | 19 | 0.466 |
| Reconsolidation, systemic | 36 | 0.67 ± 0.17 | 24.0 | 27 | 0.290 |
| Reconsolidation, amygdala | 47 | 0.60 ± 0.11 | 28.1 | 27 | 0.734 |
| Reconsolidation, hippocampus | 20 | 0.65 ± 0.13 | 13.1 | 13 | 0.974 |

**Suppl. Table 7 – Meta-regression models for testing intervention site.** Grey lines show the results of the models without moderators, while white lines show the results of meta-regressions. In meta-regression models, systemic injections are used as the reference (i.e. intercept), and injection sites not included in the predefined list were combined in a single category ("Other"). Effect sizes are presented as absolute mean differences in freezing between groups for the intercept (i.e. systemic injections), and between the effect on each site and that of systemic injections for the remaining groups. p-values refer to the estimated effect size (4th column), the Q-test for residual heterogeneity (6th column) or the significance test for moderators (8th column).

| Dataset and moderators | Sample size | Effect size (% freezing) [95% C.I.] | p-value | $I^2$ | p-value | $R^2$ | p-value |
|---|---|---|---|---|---|---|---|
| Training | 168 | -17.3 [-19.8, -14.9] | $4.4 \times 10^{-44}$ | 81.9% | $9.7 \times 10^{-79}$ | - | |
| Intercept | 57 | -20.2 [-24.0, -16.4] | $3.7 \times 10^{-25}$ | 80.3% | $3.3 \times 10^{-69}$ | 5.2% | 0.058 |
| Amygdala | 38 | -0.1 [-6.7, 6.6] | 0.982 | | | | |
| Hippocampus | 41 | 6.1 [-0.02, 12.2] | 0.050 | | | | |
| i.c.v. | 3 | -2.0 [-22.4, 18.3] | 0.844 | | | | |
| Other | 29 | 9.0 [1.8, 16.2] | 0.014 | | | | |
| Reconsolidation | 119 | -16.9 [-20.2, -13.7] | $3.6 \times 10^{-24}$ | 85.9% | $2.3 \times 10^{-123}$ | - | |
| Intercept | 36 | -18.7 [-24.0, -13.4] | $4.6 \times 10^{-12}$ | 83.2% | $1.2 \times 10^{-92}$ | 17.6% | $2.3 \times 10^{-4}$ |
| Amygdala | 47 | -1.4 [-8.6, 5.9] | 0.714 | | | | |
| Hippocampus | 20 | -1.0 [-10.0, 8.0] | 0.830 | | | | |
| i.c.v. | 4 | 11.7 [-7.6, 31.0] | 0.235 | | | | |
| Other | 12 | 23.6 [12.2, 35.0] | $4.8 \times 10^{-5}$ | | | | |
| Extinction | 23 | 9.3 [2.0, 16.7] | 0.012 | 84.4% | $5.0 \times 10^{-12}$ | - | |
| Intercept | 4 | 27.6 [13.8, 41.4] | $8.9 \times 10^{-5}$ | 70.0% | $1.3 \times 10^{-5}$ | 37.1% | 0.018 |
| Amygdala | 8 | -28.7 [-46.0, -11.5] | 0.001 | | | | |
| Hippocampus | 8 | -21.5 [-39.2, -3.7] | 0.018 | | | | |
| i.c.v. | 2 | -10.1 [-34.5, 14.3] | 0.417 | | | | |
| Other | 1 | -10.8 [-41.0, 19.4] | 0.483 | | | | |

**Suppl. Table 8 – Meta-regression models for testing intervention session.** Grey lines show the results of the models without moderators, while white lines show the results of meta-regressions. Reexposure refers to comparisons with interventions in protocols that were not defined as reconsolidation or extinction in the original article. In all meta-regression models, interventions on training are used as the reference (i.e. intercept). Effect sizes are presented as absolute mean differences in freezing between groups for the intercept (i.e. systemic injections), and between the effect in each session and that observed in the training session for the remaining groups. p-values refer to the estimated effect size (4th column), the Q-test for residual heterogeneity (6th column) or the significance test for moderators (8th column).

| Dataset and moderators | Sample size | Effect size (% freezing) [95% C.I.] | p-value | $I^2$ | p-value | $R^2$ | p-value |
|---|---|---|---|---|---|---|---|
| Systemic | 97 | -17.6 [-21.0, -14.2] | $1.7x10^{-24}$ | 88.2% | $3.7x10^{-104}$ | | |
| Intercept | 57 | -20.2 [-23.9, -16.5] | $5.3x10^{-27}$ | 82.4% | $4.2x10^{-70}$ | 36.6% | $3.8x10^{-10}$ |
| Extinction | 4 | 47.8 [33.5, 62.1] | $5.2x10^{-11}$ | | | | |
| Reconsolidation | 36 | 1.6 [-4.3, 7.5] | 0.596 | | | | |
| Amygdala | 94 | -18.88 [-22.7, -15.0] | $7.1x10^{-22}$ | 77.8% | $9.2x10^{-51}$ | | |
| Intercept | 38 | -20.2 [-26.2, -14.2] | $3.2x10^{-11}$ | 74.1% | $6.4x10^{-36}$ | 16.5% | 0.002 |
| Extinction | 8 | 19.5 [6.1, 32.9] | 0.005 | | | | |
| Reconsolidation | 47 | 0.4 [-7.3, 8.2] | 0.976 | | | | |
| Reexposure | 1 | -36.6 [-68.2, -5.0] | 0.023 | | | | |
| Hippocampus | 71 | -14.0 [-18.3, -9.6] | $3.5x10^{-10}$ | 90.5% | $1.0x10^{-89}$ | | |
| Intercept | 41 | -14.2 [-19.5, -8.8] | $1.9x10^{-7}$ | 88.3% | $1.6x10^{-55}$ | 17.0% | 0.004 |
| Extinction | 8 | 20.4 [6.6, 34.2] | 0.004 | | | | |
| Reconsolidation | 20 | -5.5 [-14.6, 3.6] | 0.234 | | | | |
| Reexposure | 2 | -17.7 [-52.5, 17.1] | 0.319 | | | | |
| i.c.v. | 9 | -7.1 [-26.6, 12.5] | 0.477 | 91.7% | $1.8x10^{-9}$ | | |
| Intercept | 3 | -22.0 [-53.6, 9.7] | 0.174 | 84.9% | $6.6x10^{-6}$ | 14.1% | 0.198 |
| Extinction | 2 | 43.4 [-5.2, 92.0] | 0.080 | | | | |
| Reconsolidation | 4 | 10.7 [-31.8, 53.2] | 0.622 | | | | |
| Other | 42 | -5.5 [-10.6, -0.4] | 0.036 | 70.0% | $2.0x10^{-12}$ | | |
| Intercept | 29 | -11.2 [-16.5, -5.9] | $3.5x10^{-5}$ | 57.0% | $8.1x10^{-6}$ | 41.1% | $4.9x10^{-4}$ |
| Extinction | 1 | 28.0 [4.2, 51.8] | 0.021 | | | | |
| Reconsolidation | 12 | 16.8 [7.2, 26.5] | $6.4x10^{-4}$ | | | | |

**Suppl. Table 9 – Multivariable meta-regression model for testing intervention session and injection site.** Grey line shows the results of the model without moderators, while white lines show the results of the meta-regression. Reexposure refers to comparisons with interventions in protocols that were not defined as reconsolidation or extinction in the original article. Systemic interventions on training are used as a reference. Injection sites not included in the predefined list were combined in a single category ("Other"). Effect sizes are presented as absolute mean differences in effects between groups for the intercept), and between sites or sessions, using systemic injections or training as a reference. p-values refer to the estimated effect size (4th column), the Q-test for residual heterogeneity (6th column) or the significance test for moderators (8th column).

<table>
<tr><th>Dataset and moderators</th><th>Sample size</th><th>Effect size (% freezing) [95% C.I.]</th><th>p-value</th><th>$I^2$</th><th>p-value</th><th>$R^2$</th><th>p-value</th></tr>
<tr><td>Complete</td><td>313</td><td>-15.4 [-17.4, -13.3]</td><td>$6.0x10^{-48}$</td><td>87.0%</td><td>$6.5x10^{-284}$</td><td></td><td></td></tr>
<tr><td>Intercept</td><td>168</td><td>-19.41 [-22.9, -16.0]</td><td>$3.2x10^{-28}$</td><td rowspan="8">81.9%</td><td rowspan="8">$2.4x10^{-195}$</td><td></td><td></td></tr>
<tr><td>Extinction</td><td>23</td><td>27.78 [20.8, 35.4]</td><td>$6.4x10^{-14}$</td><td rowspan="3">20.9%</td><td rowspan="3">$7.6x10^{-14}$</td></tr>
<tr><td>Reexposure</td><td>3</td><td>-26.14 [-48.4, -3.9]</td><td>0.021</td></tr>
<tr><td>Reconsolidation</td><td>119</td><td>1.72 [-2.1, 5.7]</td><td>0.391</td></tr>
<tr><td>Amygdala</td><td>94</td><td>-2.62 [-7.2, 2.1]</td><td>0.275</td><td rowspan="4">7.4%</td><td rowspan="4">$9.0x10^{-5}$</td></tr>
<tr><td>Hippocampus</td><td>71</td><td>2.37 [-2.6, 7.3]</td><td>0.349</td></tr>
<tr><td>i.c.v.</td><td>9</td><td>6.48 [-5.6, 18.4]</td><td>0.292</td></tr>
<tr><td>Other</td><td>42</td><td>12.52 [6.4, 18.6]</td><td>$5.7x10^{-5}$</td></tr>
</table>

**Suppl. Table 10 – Meta-regression models for testing article and group as moderators.** In these analyses, articles are included as moderators (i.e. fixed factors) rather than random factors (i.e. additional levels) as in Table 6. Q-test p-values refer to the test for moderators.

| Model | $I^2$ | $R^2$ | Q-test p-value |
|---|---|---|---|
| Complete dataset, article as moderator | 81.79% | 27.38% | $1.9x10^{-7}$ |
| Complete dataset, group as moderator | 83.95% | 13.63% | $2.9x10^{-4}$ |
| Training dataset, article as moderator | 79.91% | 4.77% | 0.265 |
| Training dataset, group as moderator | 80.88% | 0% | 0.790 |
| Reactivation dataset, article as moderator | 84.39% | 32.71% | $1.9x10^{-5}$ |
| Reactivation dataset, groups a moderator | 86.29% | 20.00% | $6.7x10^{-4}$ |

**Suppl. Table 11 – Four-level meta-analyses.** In these models, experiments are nested within articles, which are in turn nested within research groups, with all of these levels modeled as random factors. Effect sizes are in absolute mean differences. CI: confidence interval.

| Dataset | Four-level full-random model | | | | |
|---|---|---|---|---|---|
| | Estimate [95% CI] | p-value | $\sigma^2$ experiments [95% CI] | $\sigma^2$ articles [95% CI] | $\sigma^2$ groups [95% CI] |
| Complete | -15.11 [-18.0, -12.2] | $1.6x10^{-24}$ | 202.3 [157.0, 260.4] | 63.7 [13.6, 134.6] | 5.5 [0, 88.3] |
| Training | -17.19 [-19.8, -14.5] | $4.5x10^{-37}$ | 163.3 [114.9, 228.9] | 11.4 [0, 58.1] | 0 [0, >10.0] |
| Reactivation | -12.75 [-17.5, -8.0] | $1.7x10^{-7}$ | 249.5 [171.4, 364.5] | 106.7 [0, 263.9] | 14.0 [0, 233.7] |

**Suppl. Table 12 – Meta-regression models for testing all protocol variables on the complete dataset.** All variables listed in Suppl. Table 2 are included, except for intervention site and session (presented in Table 3 and Suppl. Tables 6 and 7) and the variables specific to reactivation protocols. Doses from the different drugs and routes of administration from each species were z-scored to allow cross-drug comparisons. Grey lines contain sample sizes, intercept effect sizes as absolute mean differences, p-values and $I^2$ for each model, as well as $R^2$ values and Q-test p-values for the moderator. White lines contain betas indicating the additional contribution of each unit/category to the effect size (as well as sample sizes and p-values for individual categories). For categorical variables, reference groups are described in the first column, with the sample size for these groups indicated in parentheses in the second column.

| Moderators and categories | Sample size | Effect (% freezing) [95% C.I.] | p-value | $I^2$ | $R^2$ | Moderator p-value |
|---|---|---|---|---|---|---|
| Conditioning type (intercept=contextual) | 313 (138) | -10.3 [-13.3, -7.4] | $3.8 \times 10^{-12}$ | 85.8% | 8.4% | $4.2 \times 10^{-5}$ |
| Contextual background | 59 | -11.00 [-16.5, -5.5] | | | | $9.3 \times 10^{-5}$ |
| Tone | 103 | -9.18 [-13.8, -4.5] | | | | $9.2 \times 10^{-5}$ |
| Tone-trace | 13 | -3.80 [-15.4, 7.8] | | | | 0.521 |
| Habituation (intercept=unreported) | 313 (237) | -15.14 [-17.5, -12.7] | $2.4 \times 10^{-35}$ | 86.9% | 0% | 0.701 |
| Reported | 76 | -0.92 [-5.7, 3.9] | | | | |
| Handling (intercept=unreported) | 313 (230) | -15.95 [-18.4, -13.5] | $9.7 \times 10^{-39}$ | 86.9% | 0% | 0.347 |
| Reported | 83 | 2.28 [-2.5, 7.0] | | | | |
| # of shocks | 313 | -14.33 [-16.6, -12.1] | $2.1 \times 10^{-35}$ | 86.6% | 2.6% | 0.033 |
| (number) | | -0.26 [-0.51, -0.02] | | | | |
| Shock intensity | 310 | -11.18 [-16.1, -6.3] | $7.8 \times 10^{-6}$ | 87.0% | 1.0% | 0.070 |
| (mA) | | -4.48 [-9.3, 0.4] | | | | |
| Time between drug administration and intervention session | 303 | -15.98 [-18.1, -13.9] | $4.7 \times 10^{-49}$ | 85.9% | 5.3% | $2.5 \times 10^{-4}$ |
| (min) | | 0.01 [0.006, 0.02] | | | | |
| Time between intervention sessions and test | 312 | -15.06 [-17.3, -12.8] | $5.7 \times 10^{-39}$ | 86.6% | 0% | 0.626 |
| (hour) | | -0.005 [-0.02, 0.01] | | | | |
| Species (intercept=mice) | 313 | -15.31 [-18.2, -12.4] | $1.0 \times 10^{-24}$ | 86.9% | 0% | 0.958 |
| Rats | 167 | -0.11 [-4.3, 4.0] | | | | |
| Sex (intercept=both) | 311 (18) | -16.03 [-24.6, -7.5] | $2.3 \times 10^{-4}$ | 87.1% | 0% | 0.856 |
| Male | 293 | 0.81 [-8.0, 9.6] | | | | |
| Age | 122 | -10.63 [-23.6, 2.3] | 0.107 | 87.7% | 0% | 0.430 |
| (weeks) | | -0.50 [-1.7, 0.7] | | | | |
| Housing | 242 | -18.44 [-22.6, -14.2] | $1.2 \times 10^{-17}$ | 85.5% | 1.45% | 0.058 |
| (animals/cage) | | 1.28 [-0.04, 2.6] | | | | |
| Active drug (intercept=anisomycin) | 313 (270) | -14.91 [-17.1, -12.7] | $8.7 \times 10^{-40}$ | 86.8% | 0.9% | 0.138 |

| | | | | | | |
|---|---|---|---|---|---|---|
| 4EGI-1 | 2 | 17.48 [-8.2, 43.2] | | | | 0.183 |
| Cycloheximide | 41 | -4.58 [-10.8, 1.7] | | | | 0.150 |
| Site of injection (intercept=systemic) | 313 (97) | -17.62 [-21.1, -14.2] | $1.5x10^{-23}$ | 85.7% | 6.1% | $8.3x10^{-4}$ |
| Amygdala | 94 | -1.26 [-6.4, 3.9] | | | | 0.630 |
| Hippocampus | 71 | 3.65 [-1.8, 9.1] | | | | 0.191 |
| i.c.v. | 9 | 12.61 [-0.6, 25.8] | | | | 0.061 |
| Other | 42 | 12.02 [5.3, 18.7] | | | | $4.6x10^{-4}$ |
| Dose | 293 | -15.75 [-17.9, -13.6] | $1.9x10^{-46}$ | 86.9% | 0% | 0.325 |
| (z-score) | | 1.11 [-1.1, 3.3] | | | | |
| Randomization (intercept=unreported) | 313 (298) | -15.29 [-17.4, -13.2] | $1.5x10^{-45}$ | 87.0% | 0% | 0.745 |
| Reported | 15 | -1.73 [-12.2, 8.7] | | | | |
| Blinding (intercept=unreported) | 313 (99) | -16.22 [-19.9, -12.5] | $5.6x10^{-18}$ | 87.0% | 0% | 0.581 |
| Reported | 214 | 1.25 [-3.2, 5.7] | | | | |
| Sample size calculation (intercept=unreported) | 313 (311) | -15.42 [-17.5, -13.3] | $5.9x10^{-48}$ | 87.0% | 0% | 0.514 |
| Reported | 2 | 8.86 [-17.7, 35.4] | | | | |
| Regulatory requirements statement (intercept=unreported) | 313 (38) | -20.54 [-26.5, -14.6] | $1.4x10^{-11}$ | 86.8% | 1.1% | 0.070 |
| Reported | 275 | 5.88 [-0.5, 12.2] | | | | |
| Conflict of interest (Intercept=unreported) | 313 (249) | -15.60 [-17.9, -13.3] | $4.5x10^{-39}$ | 86.8% | 0% | 0.814 |
| No | 62 | 0.93 [-4.2, 6.1] | | | | 0.722 |
| Yes | 2 | 7.42 [-19.1, 34.0] | | | | 0.584 |
| Impact factor | 278 | -14.63 [-18.5, -10.7] | $1.7x10^{-13}$ | 87.2% | 0% | 0.745 |
| (impact factor) | | -0.08 [-0.6, 0.4] | | | | |
| Citations | 313 | -16.06 [-18.7, -13.4] | $1.7x10^{-32}$ | 86.9% | 0% | 0.414 |
| (citations/year) | | 0.06 [-0.08, 0.2] | | | | |
| Region of origin (intercept=N. America) | 313 (178) | -15.88 [-18.6, -13.1] | $4.8x10^{-30}$ | 85.7% | 5.4% | 0.001 |
| Asia | 59 | 2.94 [-2.2, 8.1] | | | | 0.265 |
| Europe | 47 | -5.54 [-11.6, 0.5] | | | | 0.072 |
| Latin America | 25 | 3.17 [-4.4, 10.7] | | | | 0.414 |
| Middle East | 4 | 31.46 [13.4, 49.5] | | | | $6.4x10^{-4}$ |

**Suppl. Table 13 – Meta-regression models for testing all protocol variables on the training dataset.** All variables collected, listed in Suppl. Table 2, are included except the session of intervention, site of drug administration (presented in Table 3), sample size calculation (as none of the experiments reported it) and variables specific to reexposure protocols. Doses from the different drugs and routes of administration were z-scored to allow cross-drug comparisons. Grey lines contain sample sizes, intercept effect sizes as absolute mean differences, p-values and $I^2$ for each model, as well as $R^2$ values and Q-test p-values for the moderator. White lines contain betas indicating the additional contribution of each unit/category to the effect size (as well as sample sizes and p-values for individual categories). For categorical variables, reference groups are described in the first column, with the sample size for these groups indicated in parentheses in the second column.

| Moderators and categories | Sample size | Effect (% freezing) [95% C.I.] | p-value | $I^2$ | $R^2$ | Moderator p-value |
|---|---|---|---|---|---|---|
| Conditioning type (intercept=contextual) | 168 (61) | -15.67 [-19.5, -11.8] | $1.7x10^{-15}$ | 81.8% | 0% | 0.575 |
| Contextual background | 49 | -4.26 [-10.2, 1.7] | | | | 0.162 |
| Tone | 49 | -1.28 [-7.4, 4.8] | | | | 0.681 |
| Tone-trace | 9 | -1.75 [-14.5, 11.0] | | | | 0.788 |
| Habituation (intercept=unreported) | 168 (136) | -17.89 [-20.6, -15.2] | $7.7x10^{-38}$ | 81.8% | 0% | 0.354 |
| Reported | 32 | 2.89 [-3.2, 9.0] | | | | |
| Handling (intercept=unreported) | 168 (123) | -17.15 [-19.9, -14.4] | $1.6x10^{-33}$ | 81.9% | 0% | 0.812 |
| Reported | 45 | -0.70 [-6.5, 5.1] | | | | |
| # of shocks | 168 | -18.66 [-21.4, -15.9] | $5.1x10^{-40}$ | 81.4% | 2.4% | 0.050 |
| (number) | - | 0.43 [-0.002, 0.9] | | | | |
| Shock intensity | 168 | -14.61 [-20.5, -8.7] | $1.1x10^{-6}$ | 81.9% | 0.0% | 0.322 |
| (mA) | - | -3.04 [-9.1, 3.0] | | | | |
| Time between drug administration and training session | 166 | -19.10 [-21.5, -16.7] | $6.6 x10^{-56}$ | 76.5% | 20.0% | $4.2x10^{-7}$ |
| (min) | - | 0.01 [0.009, 0.02] | | | | |
| Time between training and test | 167 | -17.66 [-20.3, -15.0] | $5.2x10^{-40}$ | 80.6% | 0.1% | 0.307 |
| (hour) | - | 0.01 [-0.009, 0.03] | | | | |
| Species (intercept=mice) | 168 (90) | -18.64 [-21.8, -15.5] | $6.4x10^{-31}$ | 81.6% | 1.0% | 0.200 |
| Rats | 78 | 3.23 [-1.7, 8.2] | | | | |
| Sex (intercept=both) | 167 (12) | -19.71 [-28.5, -11.0] | $1.0x10^{-5}$ | 82.0% | 0% | 0.563 |
| Male | 155 | 2.69 [-6.4, 11.8] | | | | |
| Age | 75 | -10.68 [-31.2, 9.9] | 0.308 | 86.2% | 0% | 0.486 |
| (weeks) | - | -0.70 | | | | |

| | | | | | | |
|---|---|---|---|---|---|---|
| | | [-2.7, 1.3] | | | | |
| Housing | 122 | -17.96 [-23.1, -12.8] | 1.0x10$^{-11}$ | 76.5% | 0% | 0.870 |
| (animals/cage) | - | 0.14 [-1.6, 1.9] | | | | |
| Active drug (intercept=anisomycin) | 168 (140) | -16.74 [-19.4, -14.1] | 1.5x10$^{-35}$ | 81.7% | 1.3% | 0.242 |
| 4EGI-1 | 1 | 15.16 [-14.9, 45.2] | | | | 0.323 |
| Cycloheximide | 27 | -4.66 [-11.5, 2.2] | | | | 0.184 |
| Dose | 161 | -17.31 [-19.8, -14.8] | 1.3x10$^{-40}$ | 82.1% | 0% | 0.623 |
| (z-score) | - | -0.66 [-3.3, 2.0] | | | | |
| Randomization (intercept=unreported) | 168 (155) | -17.38 [-19.9, -14.9] | 1.2x10$^{-41}$ | 82.0% | 0% | 0.837 |
| Reported | 13 | 1.08 [-9.2, 11.3] | | | | |
| Blinding (intercept=unreported) | 168 (56) | -18.61 [-22.8, -14.5] | 1.8x10$^{-18}$ | 81.8% | 0% | 0.451 |
| Reported | 112 | 1.97 [-3.2, 7.1] | | | | |
| Regulatory requirements statement (intercept=unreported) | 168 (21) | -25.42 [-32.2, -18.7] | 1.7x10$^{-13}$ | 81.1% | 5.3% | 0.012 |
| Reported | 147 | 9.26 [2.0, 16.5] | | | | |
| Conflict of interest (Intercept=unreported) | 168 (142) | -18.26 [-20.9, -15.6] | 2.6x10$^{-41}$ | 80.9% | 1.4% | 0.108 |
| No | 25 | 4.94 [-1.6, 11.5] | | | | 0.141 |
| Yes | 1 | 27.91 [-7.2, 63.0] | | | | 0.119 |
| Impact factor | 134 | -16.87 [-21.6, -12.1] | 4.5x10$^{-12}$ | 80.6% | 0% | 0.868 |
| (impact factor) | - | -0.06 [-0.7, 0.6] | | | | |
| Citations | 168 | -17.49 [-20.7, -14.3] | 3.3x10$^{-26}$ | 81.7% | 0% | 0.872 |
| (citations/year) | - | 0.02 [-0.2, 0.2] | | | | |
| Region of origin (intercept=N. America) | 168 (104) | -16.16 [-19.3, -13.0] | 5.1x10$^{-24}$ | 80.5% | 1.0% | 0.237 |
| Asia | 15 | -1.75 [-9.9, 6.5] | | | | 0.674 |
| Europe | 35 | -5.30 [-11.5, 0.9] | | | | 0.094 |
| Latin America | 12 | 1.18 [-10.4, 8.1] | | | | 0.802 |
| Middle East | 2 | 17.68 [-5.1, 40.4] | | | | 0.127 |

**Suppl. Table 14 – Meta-regression models for testing all protocol variables on the reactivation dataset.** All variables collected, listed in Suppl. Table 1, are included. Reexposure duration is based on z-scored values for reexposure to tone (measured in number of tones) and to context (measured in minutes), based on the mean and standard deviation for each type of conditioning. Doses from the different drugs and routes of administration were also z-scored to allow cross-drug comparisons. Grey lines contain sample sizes, intercept effect sizes as absolute mean differences, p-values and $I^2$ for each model, as well as $R^2$ values and Q-test p-values for the moderator. White lines contain betas indicating the additional contribution of each unit/category to the effect size (as well as sample sizes and p-values for individual categories). For categorical variables, reference groups are described in the first column, with the sample size for these groups indicated in parentheses in the second column.

| Moderators and categories | Sample size | Effect size (% freezing) [95% C.I.] | p-value | $I^2$ | $R^2$ | Moderator p-value |
|---|---|---|---|---|---|---|
| Conditioning type (intercept=contextual) | 145 (77) | -6.16 [-10.4, -1.9] | 0.004 | 87.1% | 18.6% | $6.3x10^{-6}$ |
| Contextual background | 10 | -21.72 [-34.7, -8.7] | | | | 0.001 |
| Tone | 54 | -15.54 [-22.2, -8.8] | | | | $5.5x10^{-6}$ |
| Tone-Trace | 4 | 0.40 [-22.0, 22.8] | | | | 0.972 |
| Habituation (intercept=unreported) | 145 (101) | -11.7 [-15.8, -7.6] | $1.8x10^{-8}$ | 89.3% | 1.4% | 0.187 |
| Reported | 44 | -4.96 [-12.3, 2.4] | | | | |
| Handling (intercept=unreported) | 145 (107) | -14.52 [-18.5, -10.5] | $1.8x10^{-12}$ | 89.3% | 0.3% | 0.230 |
| Reported | 38 | 4.68 [-3.0, 12.3] | 0.230 | | | |
| # of shocks | 145 | -10.76 [-14.4, -7.2] | $4.9x10^{-9}$ | 88.3% | 9.2% | 0.001 |
| (number) | - | -0.52 [-0.8, -0.2] | | | | |
| Shock intensity | 142 | -7.02 [-15.0, 0.9] | 0.084 | 89.5% | 1.4% | 0.102 |
| (mA) | - | -6.33 [-13.9, 1.3] | | | | |
| Time between training and re-exposure session | 145 | -14.28 [-18.1, -10.5] | $1.6x10^{-13}$ | 89.4% | 0% | 0.216 |
| (hours) | - | 0.008 [-0.005, 0.02] | | | | |
| Time between drug administration and re-exposure session | 137 | -12.79 [-16.4, -9.2] | $2.7x10^{-12}$ | 89.6% | 0% | 0.376 |
| (min) | - | 0.01 [-0.01, 0.03] | | | | |
| Re-exposure duration | 143 | -13.26 [-16.4, -10.1] | $3.0x10^{-16}$ | 86.9% | 19.0% | $2.8x10^{-7}$ |
| (z-score) | - | 8.31 [5.1, 11.5] | | | | |
| Time between re-exposure and test | 145 | -11.46 [-15.3, -7.6] | $4.7x10^{-39}$ | 89.0% | 2.5% | 0.055 |
| (hour) | - | -0.04 [-0.08, 0.0008] | | | | |
| Species (intercept=mice) | 145 (56) | -10.32 [-15.6, -5.0] | $1.3x10^{-4}$ | 89.3% | 1.2% | 0.159 |

| | | | | | | |
|---|---|---|---|---|---|---|
| Rats | 89 | -4.96 [-11.9, 1.9] | | | | |
| Sex (intercept=both) | 144 (6) | -8.43 [-25.7, 8.8] | 0.337 | 89.6% | 0% | 0.586 |
| Male | 138 | -4.89 [-22.5, 12.7] | | | | |
| Age | 38 | -19.67 [-32.3, -7.0] | 0.002 | 68.5% | 0% | 0.785 |
| (weeks) | - | 0.17 [-1.0, 1.4] | | | | |
| Housing | 120 | -17.92 [-24.6, -11.3] | $1.3x10^{-7}$ | 89.7% | 2.5% | 0.062 |
| (animals/cage) | - | 1.86 [-0.1, 3.8] | | | | |
| Active drug (intercept=anisomycin) | 145 (130) | -13.01 [-16.6, -9.4] | $1.7x10^{-12}$ | 89.4% | 0% | 0.521 |
| 4EGI-1 | 1 | 20.22 [-22.1, 62.5] | | | | 0.350 |
| Cycloheximide | 14 | -3.72 [-15.4, 7.9] | | | | 0.531 |
| Site of injection (intercept=systemic) | 145 (40) | -14.16 [-20.1, -8.2] | $3.5x10^{-6}$ | 87.7% | 11.6% | 0.001 |
| Amygdala | 56 | -3.97 [-11.9, 4.0] | | | | 0.328 |
| Hippocampus | 30 | 0.73 [-8.7, 10.1] | | | | 0.879 |
| i.c.v. | 6 | 16.67 [-1.3, 34.7] | | | | 0.069 |
| Other | 13 | 19.81 [7.2, 32.4] | | | | 0.002 |
| Dose | 132 | -14.01 [-17.6, -10.4] | $2.3x10^{-14}$ | 89.4% | 1.3% | 0.107 |
| (z-score) | - | 3.03 [-0.6, 6.7] | | | | |
| Randomization (intercept=unreported) | 145 (143) | -13.12 [-16.6, -9.7] | $9.1x10^{-14}$ | 89.5% | 0% | 0.597 |
| Reported | 2 | -7.61 [-35.8, 20.6] | | | | |
| Blinding (intercept=unreported) | 145 (43) | -13.16 [-19.5, -6.8] | $4.9x10^{-5}$ | 89.5% | 0% | 0.977 |
| Reported | 102 | -0.11 [-7.7, 7.4] | | | | |
| Sample size calculation (intercept=unreported) | 145 (143) | -13.31 [-16.8, -9.9] | $3.7x10^{-14}$ | 89.6% | 0% | 0.695 |
| Reported | 2 | 6.01 [-24.1, 36.1] | | | | |
| Regulatory requirements statement (intercept=unreported) | 145 (17) | -14.86 [-25.0, -4.7] | 0.004 | 89.4% | 0% | 0.740 |
| Reported | 128 | 1.83 [-8.9, 12.7] | | | | |
| Conflict of interest (Intercept=unreported) | 145 (107) | -12.32 [-16.3, -8.3] | $1.5x10^{-9}$ | 89.5% | 0% | 0.651 |
| No | 37 | -3.27 [-11.1, 4.6] | | | | 0.413 |
| Yes | 1 | -9.81 [-50.2, 30.6] | | | | 0.634 |

| | | | | | | |
|---|---|---|---|---|---|---|
| Impact factor | 144 | -11.95 [-18.0, -5.9] | $1.1x10^{-4}$ | 89.5% | 0% | 0.562 |
| (impact factor) | - | -0.20 [-0.9, 0.5] | | | | |
| Citations | 145 | -14.00 [-18.4, -9.7] | $2.7x10^{-10}$ | 89.4% | 0% | 0.574 |
| (citations/year) | - | 0.06 [-0.1, 0.3] | | | | |
| Region of origin (intercept=N. America) | 145 (74) | -15.48 [-20.2, -10.8] | $1.4x10^{-10}$ | 88.6% | 6.3% | 0.013 |
| Asia | 44 | 4.25 [-3.2, 11.7] | | | | 0.264 |
| Europe | 12 | -6.30 [-19.6, 7.0] | | | | 0.353 |
| Latin America | 13 | 7.48 [-4.6, 19.6] | | | | 0.226 |
| Middle East | 2 | 44.28 [15.8, 72.7] | | | | 0.002 |

**Suppl. Table 15 – Meta-regression models for testing reexposure duration separately in context and tone conditioning.** Grey lines contain sample sizes, intercept effect sizes as absolute mean differences, p-values and $I^2$ for each model, as well as $R^2$ values and Q-test p-values for the moderator. White lines contain betas indicating the additional contribution of each unit to the effect size.

| Dataset and moderators | Sample size | Effect (% freezing) [95%C.I.] | p-value | $I^2$ | $R^2$ | Q-test p-value |
|---|---|---|---|---|---|---|
| Reexposure to context | 85 | -16.57 [-21.3, -11.8] | $6.1x10^{-12}$ | 84.5% | 30.7% | $9.0x10^{-8}$ |
| (minutes) | - | 1.08 [0.7, 1.5] | | | | |
| Reexposure to tone | 54 | -25.45 [-31.8, -19.1] | $3.5x10^{-15}$ | 84.7% | 7.1% | 0.029 |
| (number) | - | 2.12 [0.2, 4.0] | 0.029 | | | |

**Suppl. Table 16 – Three-level meta-regression models for testing all protocol variables on the complete dataset, accounting for nesting of experiments within articles.** All variables collected, listed in Suppl. Table 2, are included except the variables specific to reexposure protocols. Doses from the different drugs and routes of administration were z-scored to allow cross-drug comparisons. Grey lines contain sample sizes, intercept effect sizes as absolute mean differences, p-values and $I^2$ for each model, as well as $R^2$ values and Q-test p-values for the moderator. White lines contain betas indicating the additional contribution of each unit/category to the effect size (as well as sample sizes and p-values for individual categories). For categorical variables, reference groups are described in the first column, with the sample size for these groups indicated in parentheses in the second column. $R^2$ values are calculated as the difference between total variances in the model with no moderators and in the tested model, divided by the total variance in the model with no moderators.

| Moderators and categories | Sample size | Effect (% freezing) [95%C.I.] | p-value | $I^2$ | $R^2$ | Moderator p-value |
|---|---|---|---|---|---|---|
| Intervention session (intercept=training) | 313 (168) | -17.08 [-20.5, -13.7] | $2.8x10^{-23}$ | 84.4% | 20.7% | $1.8x10^{-15}$ |
| Extinction | 23 | 29.30 [21.6, 37.0] | | | | $8.1x10^{-14}$ |
| Reactivation | 3 | -29.97 [-53.4, -6.5] | | | | 0.012 |
| Reconsolidation | 119 | 0.006 [-4.5, 4.5] | | | | 0.998 |
| Conditioning type (intercept=contextual) | 313 (138) | -10.51 [-14.1, -7.0] | $7.1x10^{-9}$ | 86.2% | 8.3% | 0.001 |
| Contextual background | 59 | -12.75 [-19.4, -6.1] | | | | $1.9x10^{-4}$ |
| Tone | 103 | -7.74 [-13.2, -2.3] | | | | 0.005 |
| Tone-trace | 13 | -5.05 [-17.0, 6.9] | | | | 0.410 |
| Habituation (intercept=unreported) | 313 (237) | -15.38 [-18.6, -12.2] | $3.9x10^{-21}$ | 87.3% | 0% | 0.765 |
| Reported | 76 | 0.91 [-5.1, 6.9] | | | | |
| Handling (intercept=unreported) | 313 (230) | -15.67 [-18.9, -12.5] | $9.1x10^{-22}$ | 87.2% | 0% | 0.538 |
| Reported | 83 | 1.89 [-4.1, 7.9] | | | | |
| # of shocks | 313 | -14.32 [-17.3, -11.4] | $1.2x10^{-21}$ | 87.0% | 1.8% | 0.194 |
| (number) | - | -0.20 [-0.5, 0.1] | | | | |
| Shock intensity | 310 | -11.10 [-17.4, -4.8] | $6.1x10^{-4}$ | 87.2% | 0.2% | 0.179 |
| (mA) | - | -4.28 [-10.5, 2.0] | | | | |
| Time between drug administration and intervention session | 303 | -15.92 [-18.6, -13.3] | $7.3x10^{-32}$ | 86.4% | 7.8% | $1.7x10^{-4}$ |
| (min) | - | 0.01 [0.006, 0.019] | | | | |
| Time between intervention sessions and test | 312 | -14.90 [-17.8, -12.0] | $1.6x10^{-23}$ | 87.2% | 0% | 0.819 |
| (hour) | - | -0.002 [-0.023, 0.018] | | | | |
| Species (intercept=mice) | 313 (146) | -15.74 [-19.9, -11.6] | $1.1x10^{-13}$ | 87.3% | 0% | 0.701 |

| | | | | | | |
|---|---|---|---|---|---|---|
| Rats | 167 | 1.08<br>[-4.4, 6.6] | | | | |
| Sex<br>(intercept=both) | 311 (18) | -16.01<br>[-26.5, -5.5] | 0.003 | 87.3% | 0% | 0.844 |
| Male | 293 | 1.10<br>[-9.8, 12.0] | | | | |
| Age | 122 | -10.48<br>[-24.4, 3.5] | 0.140 | 88.2% | 9.2% | 0.459 |
| (weeks) | - | -0.51<br>[-1.9, 0.8] | | | | |
| Housing | 242 | -16.89<br>[-22.6, -11.1] | $8.2x10^{-9}$ | 85.8% | 0% | 0.318 |
| (animals/cage) | - | 0.97<br>[-0.9, 2.9] | | | | |
| Active drug<br>(intercept=anisomycin) | 313 (270) | -15.14<br>[-18.0, -12.3] | $8.7x10^{-25}$ | 87.2% | 0% | 0.445 |
| 4EGI-1 | 2 | 17.66<br>[-10.8, 46.1] | | | | 0.224 |
| Cycloheximide | 41 | -1.46<br>[-10.0, 7.0] | | | | 0.736 |
| Site of injection<br>(intercept=systemic) | 313 (97) | -18.87<br>[-23.5, -14.2] | $2.4x10^{-15}$ | 86.4% | 6.7% | 0.006 |
| Amygdala | 94 | 0.77<br>[-5.8, 7.3] | | | | 0.816 |
| Hippocampus | 71 | 5.09<br>[-1.7, 11.8] | | | | 0.139 |
| i.c.v. | 9 | 11.76<br>[-2.0, 25.5] | | | | 0.095 |
| Other | 42 | 13.36<br>[5.4, 21.3] | | | | 0.001 |
| Dose | 293 | -15.57<br>[-18.4, -12.8] | $9.4x10^{-24}$ | 87.2% | 0% | 0.940 |
| (z-score) | - | 0.09<br>[-2.4, 2.5] | | | | |
| Randomization<br>(intercept=unreported) | 313 (298) | -15.01<br>[-17.8, -12.2] | $4.7x10^{-26}$ | 87.3% | 0% | 0.641 |
| Reported | 15 | -3.55<br>[-18.5, 11.4] | | | | |
| Blinding<br>(intercept=unreported) | 313 (99) | -15.34<br>[-20.0, -10.7] | $1.3x10^{-10}$ | 87.3% | 0% | 0.915 |
| Reported | 214 | 0.31<br>[-5.5, 6.1] | | | | |
| Sample size calculation<br>(intercept=unreported) | 313 (311) | -15.23<br>[-18.0, -12.5] | $8.2x10^{-28}$ | 87.2% | 0% | 0.529 |
| Reported | 2 | 8.62<br>[-18.2, 35.5] | | | | |
| Regulatory requirements statement<br>(intercept=unreported) | 313 (38) | -19.53<br>[-28.0, -11.1] | $5.6x10^{-6}$ | 87.1% | 0.2% | 0.281 |
| Reported | 275 | 4.90<br>[-4.0, 13.8] | | | | |
| Conflict of interest<br>(Intercept=unreported) | 313 (249) | -13.45<br>[-18.9, -8.0] | $1.1x10^{-6}$ | 87.3% | 0% | 0.696 |
| No | 62 | -2.34<br>[-8.6, 4.0] | | | | 0.466 |
| Yes | 2 | 4.85 | | | | 0.748 |

| | | | | | | |
|---|---|---|---|---|---|---|
| | | [-24.8, 34.5] | | | | |
| Impact factor | 278 | -14.81 [-19.7, -9.9] | $3.2x10^{-9}$ | 87.6% | 0% | 0.853 |
| (impact factor) | - | -0.06 [-0.7, 0.6] | | | | |
| Citations | 313 | -15.31 [-18.7, -11.9] | $1.6x10^{-18}$ | 87.3% | 0% | 0.861 |
| (citations/year) | - | 0.02 [-0.2, 0.2] | | | | |
| Region of origin (intercept=N. America) | 313 (178) | -15.84 [-19.3, -12.4] | $4.3x10^{-19}$ | 86.6% | 5.1% | 0.012 |
| Asia | 59 | 2.58 [-4.3, 9.5] | | | | 0.463 |
| Europe | 47 | -5.10 [-13.1, 2.9] | | | | 0.209 |
| Latin America | 25 | 2.86 [-6.0, 11.7] | | | | 0.526 |
| Middle East | 4 | 31.39 [11.6, 51.1] | | | | 0.002 |

**Suppl. Table 17 – Three-level meta-regression models for testing all protocol variables on the training dataset, accounting for nesting of experiments within articles.** All variables collected, listed in Suppl. Table 2, are included except the variables specific to reexposure protocols and sample size calculation due to lack of reporting in all included articles. Doses from the different drugs and routes of administration were z-scored to allow cross-drug comparisons. Grey lines contain sample sizes, intercept effect sizes as absolute mean differences, p-values and $I^2$ for each model, as well as $R^2$ values and Q-test p-values for the moderator. White lines contain betas indicating the additional contribution of each unit/category to the effect size (as well as sample sizes and p-values for individual categories). For categorical variables, reference groups are described in the first column, with the sample size for these groups indicated in parentheses in the second column. $R^2$ values are calculated as the difference between total variances in the model with no moderators and in the tested model, divided by the total variance in the model with no moderators.

| Moderators and categories | Sample size | Effect (% freezing) [95%C.I.] | p-value | $I^2$ | $R^2$ | Moderator p-value |
|---|---|---|---|---|---|---|
| Conditioning type (intercept=contextual) | 168 (61) | -15.62 [-19.7, -11.5] | $9.2x10^{-14}$ | 82.2% | 0% | 0.410 |
| Contextual background | 49 | -5.40 [-12.1, 1.3] | | | | 0.114 |
| Tone | 49 | -0.51 [-7.0, 5.9] | | | | 0.877 |
| Tone-trace | 9 | -2.28 [-15.2, 10.6] | | | | 0.730 |
| Habituation (intercept=unreported) | 168 (136) | -17.90 [-20.9, -14.9] | $5.7x10^{-31}$ | 82.0% | 0% | 0.315 |
| Reported | 32 | 3.28 [-3.1, 9.7] | | | | |
| Handling (intercept=unreported) | 168 (123) | -16.92 [-20.0, -13.8] | $4.3x10^{-27}$ | 82.0% | 0% | 0.737 |
| Reported | 45 | -1.08 [-7.4, 5.2] | | | | |
| # of shocks | 168 | -18.56 [-21.5, -15.6] | $4.5x10^{-35}$ | 81.6% | 2.4% | 0.058 |
| (number) | - | 0.42 [-0.01, 0.85] | | | | |
| Shock intensity | 168 | -14.49 [-20.8, -8.8] | $7.2x10^{-6}$ | 81.9% | 0% | 0.356 |

| | | | | | | |
|---|---|---|---|---|---|---|
| (mA) | - | -3.01 [-9.4, 3.4] | | | | |
| Time between drug administration and intervention session | 166 | -19.04 [-21.5, -16.6] | $3.9x10^{-53}$ | 77.4% | 24.8% | $5.2x10^{-7}$ |
| (min) | - | 0.01 [0.009, 0.02] | | | | |
| Time between intervention sessions and test | 167 | -17.50 [-20.4, -14.6] | $4.1x10^{-33}$ | 81.7% | 1.0% | 0.340 |
| (hour) | - | 0.009 [-0.01, 0.03] | | | | |
| Species (intercept=mice) | 168 (90) | -18.70 [-22.2, -15.2] | $3.7x10^{-25}$ | 81.8% | 0.8% | 0.201 |
| Rats | 78 | 3.50 [-1.9, 8.9] | | | | |
| Sex (intercept=both) | 167 (12) | -19.74 [-29.2, -10.3] | $4.0x10^{-5}$ | 82.1% | 0% | 0.562 |
| Male | 155 | 2.91 [-6.9, 12.7] | | | | |
| Age | 75 | -7.09 [-29.7, 15.5] | 0.539 | 87.1% | 0% | 0.345 |
| (weeks) | - | -1.06 [-3.3, 1.1] | | | | |
| Housing | 122 | -16.32 [-22.4, -10.3] | $1.2x10^{-7}$ | 77.1% | 0% | 0.812 |
| (animals/cage) | - | -0.24 [-2.2, 1.7] | | | | |
| Active drug (intercept=anisomycin) | 168 (140) | -16.82 [-19.6, -14.0] | $2.2x10^{-32}$ | 81.8% | 0.8% | 0.347 |
| 4EGI-1 | 1 | 15.24 [-14.9, 45.4] | | | | 0.321 |
| Cycloheximide | 27 | -4.06 [-11.8, 3.7] | | | | 0.306 |
| Site of injection (intercept=systemic) | 168 (57) | -20.52 [-24.8, -16.2] | $1.0x10^{-20}$ | 81.1% | 5.1% | 0.066 |
| Amygdala | 38 | 0.28 [-7.0, 7.5] | | | | 0.940 |
| Hippocampus | 41 | 6.73 [0.06, 13.4] | | | | 0.048 |
| i.c.v. | 3 | -2.00 [-22.5, 18.5] | | | | 0.848 |
| Other | 29 | 9.39 [1.7, 17.1] | | | | 0.017 |
| Dose | 161 | -17.19 [-20.0, -14.4] | $1.4x10^{-33}$ | 82.2% | 0% | 0.515 |
| (z-score) | - | -0.89 [-3.6, 1.8] | | | | |
| Randomization (intercept=unreported) | 168 (155) | -17.20 [-20.0, -14.5] | $4.3x10^{-34}$ | 82.1% | 0% | 0.940 |
| Reported | 13 | 0.46 [-11.5, 12.5] | | | | |
| Blinding (intercept=unreported) | 168 (56) | -18.42 [-22.8, -14.0] | $2.9x10^{-16}$ | 82.0% | 0% | 0.494 |
| Reported | 112 | 1.93 [-3.6, 7.5] | | | | |

| | | | | | | |
|---|---|---|---|---|---|---|
| Regulatory requirements statement (intercept=unreported) | 168 (21) | -25.42 [-32.2, -18.7] | $1.7x10^{-13}$ | 81.1% | 5.4% | 0.012 |
| Reported | 147 | 9.26 [2.0, 16.5] | | | | |
| Conflict of interest (Intercept=unreported) | 168 (142) | -13.15 [-19.4, -6.9] | $4.0x10^{-5}$ | 81.7% | 1.4% | 0.113 |
| No | 25 | -5.11 [-12.0, 1.8] | | | | 0.148 |
| Yes | 1 | 22.80 [-12.8, 58.4] | | | | 0.209 |
| Impact factor | 134 | -16.87 [-21.6, -12.1] | $4.5x10^{-12}$ | 81.1% | 8.5% | 0.868 |
| (impact factor) | - | -0.06 [-0.7, 0.6] | | | | |
| Citations | 168 | -17.17 [-20.7, -13.7] | $8.2x10^{-22}$ | 82.1% | 0% | 0.995 |
| (citations/year) | - | 0.0007 [-0.2, 0.2] | | | | |
| Region of origin (intercept=N. America) | 168 (104) | -16.24 [-19.5, -12.9] | $5.9x10^{-22}$ | 81.8% | 0.7% | 0.318 |
| Asia | 15 | -1.69 [-10.1, 6.7] | | | | 0.694 |
| Europe | 35 | -4.89 [-11.6, 1.8] | | | | 0.151 |
| Latin America | 12 | -1.11 [-10.6, 8.4] | | | | 0.819 |
| Middle East | 2 | 17.76 [-5.3, 40.8] | | | | 0.131 |

**Suppl. Table 18 – Three-level meta-regression models for testing all protocol variables on the reactivation dataset, accounting for nesting of experiments within articles.** All variables collected, listed in Suppl. Table 2, are included. Reexposure duration is based on z-scored values for reexposure to tone (measured in number of tones) and to context (measured in minutes), based on the mean and standard deviation for each type of conditioning. Doses from the different drugs and routes of administration were also z-scored to allow cross-drug comparisons. Grey lines contain sample sizes, intercept effect sizes as absolute mean differences, p-values and $I^2$ for each model, as well as $R^2$ values and Q-test p-values for the moderator. White lines contain betas indicating the additional contribution of each unit/category to the effect size (as well as sample sizes and p-values for individual categories). For categorical variables, reference groups are described in the first column, with the sample size for these groups indicated in parentheses in the second column. $R^2$ values are calculated as the difference between total variances in the model with no moderators and in the tested model, divided by the total variance in the model with no moderators.

| **Moderators and categories** | **Sample size** | **Effect (% freezing) [95%C.I.]** | **p-value** | **$I^2$** | **$R^2$** | **Moderator p-value** |
|---|---|---|---|---|---|---|
| Intervention session (intercept=reexposure) | 145 (3) | -46.98 [-71.4, -22.6] | $1.6x10^{-4}$ | 85.6% | 31.5% | $5.0x10^{-16}$ |
| Extinction | 23 | 60.26 [35.1, 85.4] | | | | $2.7x10^{-6}$ |
| Reconsolidation | 119 | 29.80 [5.1, 54.5] | | | | 0.018 |
| Conditioning type (intercept=contextual) | 145 (77) | -5.96 [-10.7, -1.2] | 0.014 | 87.4% | 19.9% | $6.4x10^{-5}$ |
| Contextual background | 10 | -21.91 [-35.5, -8.3] | | | | 0.002 |
| Tone | 54 | -15.20 [-22.6, -7.8] | | | | $5.3x10^{-5}$ |
| Tone-trace | 4 | -0.47 [-23.1, 22.2] | | | | 0.967 |
| Habituation (intercept=unreported) | 145 (101) | -12.26 [-17.6, -6.9] | $7.0x10^{-6}$ | 89.7% | 0% | 0.599 |
| Reported | 44 | -2.53 [-12.0, 6.9] | | | | |
| Handling (intercept=unreported) | 145 (107) | -14.35 [-19.6, -9.0] | $1.1x10^{-7}$ | 89.7% | 0.007% | 0.396 |
| Reported | 38 | 4.04 [-5.3, 13.4] | | | | |
| # of shocks | 145 | -10.69 [-15.3, -6.1] | $4.7x10^{-6}$ | 88.9% | 7.6% | 0.016 |
| (number) | - | -0.52 [-0.9, -0.1] | | | | |
| Shock intensity | 142 | -7.76 [-18.1, 2.5] | 0.140 | 89.8% | 0% | 0.286 |
| (mA) | - | -5.47 [-15.5, 4.6] | | | | |
| Time between training and re-exposure session | 145 | -13.35 [-17.9, -8.8] | $1.0x10^{-8}$ | 89.7% | 0% | 0.668 |
| (hours) | - | 0.003 [-0.01, 0.02] | | | | |
| Time between drug administration and re-exposure session | 137 | -13.14 [-17.6, -8.7] | $8.6x10^{-9}$ | 89.8% | 0.5% | 0.317 |
| (min) | - | 0.01 [-0.01, 0.03] | | | | |
| Re-exposure duration | 143 | -12.90 [-17.2, -8.6] | $4.1x10^{-9}$ | 87.3% | 17.6% | $2.0x10^{-9}$ |

| | | | | | | |
|---|---|---|---|---|---|---|
| (z-score) | - | 8.94 [6.0, 11.9] | | | | |
| Time between re-exposure and test | 145 | -11.93 [-16.6, -7.2] | 7.0x10$^{-7}$ | 89.5% | 1.7% | 0.222 |
| (hour) | - | -0.03 [-0.07, 0.02] | | | | |
| Species (intercept=mice) | 145 (56) | -10.95 [-18.4, -3.5] | 0.004 | 89.7% | 0% | 0.487 |
| Rats | 89 | -3.28 [12.5, 6.0] | | | | |
| Sex (intercept=both) | 144 (6) | -8.90 [-30.6, 12.8] | 0.421 | 89.8% | 0% | 0.714 |
| Male | 138 | -4.14 [-26.3, 18.0] | | | | |
| Age | 47 | -11.59 [-31.0, 7.8] | 0.243 | 87.8% | 13.5% | 0.889 |
| (weeks) | - | -0.13 [-2.0, 1.7] | | | | |
| Housing | 120 | -17.75 [-26.4, -9.1] | 5.6x10$^{-5}$ | 89.9% | 0% | 0.141 |
| (animals/cage) | - | 2.13 [-0.7, 5.0] | | | | |
| Active drug (intercept=anisomycin) | 145 (130) | -13.27 [-18.0, -8.5] | 4.6x10$^{-8}$ | 89.8% | 0% | 0.644 |
| 4EGI-1 | 1 | 20.48 [-22.4, 63.3] | | | | 0.349 |
| Cycloheximide | 14 | -0.10 [-13.2, 13.0] | | | | 0.988 |
| Site of injection (intercept=systemic) | 145 (40) | -15.28 [-22.9, -7.7] | 8.5x10$^{-5}$ | 88.4% | 11.7% | 0.011 |
| Amygdala | 56 | -1.69 [-11.5, 8.1] | | | | 0.735 |
| Hippocampus | 30 | 1.90 [-8.8, 12.6] | | | | 0.728 |
| i.c.v. | 6 | 14.91 [-3.9, 33.8] | | | | 0.121 |
| Other | 13 | 20.22 [6.4, 34.1] | | | | 0.004 |
| Dose | 132 | -13.72 [-18.3, -9.1] | 4.7x10$^{-9}$ | 89.7% | 0% | 0.518 |
| (z-score) | - | 1.32 [-2.7, 5.3] | | | | |
| Randomization (intercept=unreported) | 145 (143) | -12.91 [-17.4, -8.4] | 1.9x10$^{-8}$ | 89.8% | 0% | 0.638 |
| Reported | 2 | -7.81 [-40.4, 24.7] | | | | |
| Blinding (intercept=unreported) | 145 (43) | -11.86 [-20.0, -3.7] | 0.004 | 89.8% | 0% | 0.731 |
| Reported | 102 | -1.71 [-11.5, 8.0] | | | | |
| Sample size calculation (intercept=unreported) | 145 (143) | -13.19 [-17.7, -8.7] | 8.3x10$^{-9}$ | 89.7% | 0% | 0.707 |
| Reported | 2 | 5.84 [-24.6, 36.3] | | | | |
| Regulatory requirements statement | 145 (17) | -16.03 [-29.3, -2.8] | 0.018 | 89.8% | 0% | 0.641 |

| | | | | | | |
|---|---|---|---|---|---|---|
| (intercept=unreported) | | | | | | |
| Reported | 128 | 3.36 [-10.7, 17.4] | | | | |
| Conflict of interest (Intercept=unreported) | 145 (107) | -12.76 [-18.2, -7.3] | 4.6x10$^{-6}$ | 89.9% | 0% | 0.902 |
| No | 37 | -0.59 [-10.3, 9.1] | | | | 0.905 |
| Yes | 1 | -9.38 [-50.6, 31.8] | | | | 0.655 |
| Impact factor | 144 | -11.80 [-19.4, -4.2] | 0.002 | 89.7% | 0% | 0.617 |
| (impact factor) | - | -0.22 [-1.1, 0.7] | | | | |
| Citations | 145 | -13.25 [-18.7, -7.8] | 1.9x10$^{-6}$ | 89.8% | 0% | 0.897 |
| (citations/year) | - | 0.02 [-0.3, 0.3] | | | | |
| Region of origin (intercept=N. America) | 145 (74) | -15.20 [-21.1, -9.3] | 5.2x10$^{-7}$ | 89.1% | 6.1% | 0.043 |
| Asia | 44 | 3.98 [-6.0, 14.0] | | | | 0.436 |
| Europe | 12 | -6.45 [-21.2, 8.3] | | | | 0.392 |
| Latin America | 13 | 8.19 [-6.2, 22.6] | | | | 0.264 |
| Middle East | 2 | 44.04 [12.0, 76.0] | | | | 0.007 |

**Suppl. Table 19 – Three-level meta-regression models for testing all protocol variables on the complete dataset, accounting for nesting of experiments within research groups.** All variables collected, listed in Suppl. Table 2, are included except the variables specific to reexposure protocols. Doses from the different drugs and routes of administration were z-scored to allow cross-drug comparisons. Grey lines contain sample sizes, intercept effect sizes as absolute mean differences, p-values and $I^2$ for each model, as well as $R^2$ values and Q-test p-values for the moderator. White lines contain betas indicating the additional contribution of each unit/category to the effect size (as well as sample sizes and p-values for individual categories). For categorical variables, reference groups are described in the first column, with the sample size for these groups indicated in parentheses in the second column. $R^2$ values are calculated as the difference between total variances in the model with no moderators and in the tested model, divided by the total variance in the model with no moderators.

| Moderators and categories | Sample size | Effect (% freezing) [95%C.I.] | p-value | $I^2$ | $R^2$ | Moderator p-value |
|---|---|---|---|---|---|---|
| Intervention session (intercept=training) | 313 (168) | -17.36 [-21.1, -13.6] | 9.2x10$^{-20}$ | 85.1% | 16.1% | 6.8x10$^{-13}$ |
| Extinction | 23 | 26.12 [18.6, 33.7] | | | | 1.1x10$^{-11}$ |
| Reactivation | 3 | -30.91 [-53.5, -8.3] | | | | 0.007 |
| Reconsolidation | 119 | 0.27 [-4.1, 4.7] | | | | 0.904 |
| Conditioning type (intercept=contextual) | 313 (138) | -10.48 [-13.6, -7.3] | 5.6x10$^{-11}$ | 86.0% | 9.9% | 1.2x10$^{-4}$ |
| Contextual background | 59 | -10.91 [-16.7, -5.2] | | | | 2.0x10$^{-4}$ |
| Tone | 103 | -9.36 [-14.3, -4.4] | | | | 1.9x10$^{-4}$ |
| Tone-trace | 13 | -3.64 [-15.4, 8.1] | | | | 0.543 |

| | | | | | | |
|---|---|---|---|---|---|---|
| Habituation (intercept=unreported) | 313 (237) | -15.33 [-18.9, -11.8] | $2.0x10^{-17}$ | 87.6% | 0% | 0.909 |
| Reported | 76 | 0.35 [-5.6, 6.3] | | | | |
| Handling (intercept=unreported) | 313 (230) | -15.94 [-19.5, -12.4] | $2.2x10^{-18}$ | 87.5% | 0% | 0.319 |
| Reported | 83 | 2.64 [-2.6, 7.8] | | | | |
| # of shocks | 313 | -14.35 [-17.6, -11.1] | $1.1x10^{-17}$ | 87.0% | 1.4% | 0.139 |
| (number) | - | -0.21 [-0.5, 0.07] | | | | |
| Shock intensity | 310 | -9.41 [-16.1, -2.7] | 0.006 | 87.6% | 0% | 0.054 |
| (mA) | - | -7.05 [-14.2, 0.11] | | | | |
| Time between drug administration and intervention session | 303 | -16.12 [-19.4, -12.8] | $1.5x10^{-21}$ | 86.7% | 5.3% | $3.0x10^{-4}$ |
| (min) | - | 0.01 [0.005, 0.02] | | | | |
| Time between intervention sessions and test | 312 | -14.95 [-18.6, -11.4] | $3.8x10^{-16}$ | 87.5% | 0% | 0.852 |
| (hour) | - | -0.002 [-0.02, 0.02] | | | | |
| Species (intercept=mice) | 313 (146) | -15.84 [-20.5, -11.2] | $3.3x10^{-11}$ | 87.6% | 0% | 0.722 |
| Rats | 167 | 1.17 [-5.3, 7.6] | | | | |
| Sex (intercept=both) | 311 (18) | -17.04 [-26.6, -7.5] | $4.7x10^{-4}$ | 87.6% | 0% | 0.670 |
| Male | 293 | 2.06 [-7.4, 11.6] | | | | |
| Age | 122 | -8.90 [-22.3, 4.5] | 0.192 | 88.2% | 9.1% | 0.303 |
| (weeks) | - | -0.67 [-2.0, 0.6] | | | | |
| Housing | 242 | -16.73 [-23.2, -10.3] | $3.9x10^{-7}$ | 86.0% | 0% | 0.475 |
| (animals/cage) | - | 0.70 [-1.2, 2.6] | | | | |
| Active drug (intercept=anisomycin) | 313 (270) | -14.76 [-18.3, -11.3] | $1.6x10^{-16}$ | 87.4% | 0% | 0.220 |
| 4EGI-1 | 2 | 18.91 [-6.1, 43.9] | | | | 0.138 |
| Cycloheximide | 41 | -3.53 [-11.3, 4.3] | | | | 0.374 |
| Site of injection (intercept=systemic) | 313 (97) | -19.47 [-24.5, -14.4] | $3.0x10^{-14}$ | 86.7% | 3.8% | $3.4x10^{-4}$ |
| Amygdala | 94 | 0.79 [-6.4, 8.0] | | | | 0.830 |
| Hippocampus | 71 | 5.55 [-0.7, 11.8] | | | | 0.082 |
| i.c.v. | 9 | 14.52 [0.9, 28.2] | | | | 0.037 |
| Other | 42 | 14.23 | | | | $3.6x10^{-4}$ |

| | | | | | | |
|---|---|---|---|---|---|---|
| | | [6.4, 22.0] | | | | |
| Dose | 293 | -15.59 [-18.9, -12.3] | $8.8x10^{-21}$ | 87.3% | 0% | 0.576 |
| (z-score) | - | 0.67 [-1.7, 3.0] | | | | |
| Randomization (intercept=unreported) | 313 (298) | -15.07 [-18.5, -11.6] | $8.8x10^{-18}$ | 87.6% | 0% | 0.600 |
| Reported | 15 | -3.50 [-16.6, 9.6] | | | | |
| Blinding (intercept=unreported) | 313 (99) | -15.96 [-21.0, -11.0] | $3.9x10^{-10}$ | 87.5% | 0% | 0.713 |
| Reported | 214 | 0.99 [-4.3, 6.3] | | | | |
| Sample size calculation (intercept=unreported) | 313 (311) | -15.38 [-18.7, -12.1] | $7.4x10^{-20}$ | 87.5% | 0% | 0.602 |
| Reported | 2 | 7.19 [-19.9, 34.2] | | | | |
| Regulatory requirements statement (intercept=unreported) | 313 (38) | -18.49 [-25.7, -11.3] | $4.5x10^{-7}$ | 87.3% | 0% | 0.333 |
| Reported | 275 | 3.51 [-3.6, 10.6] | | | | |
| Conflict of interest (Intercept=unreported) | 313 (249) | -16.26 [-20.1, -12.4] | $1.8x10^{-16}$ | 87.7% | 0% | 0.488 |
| No | 62 | 3.26 [-2.5, 9.0] | | | | 0.266 |
| Yes | 2 | 6.55 [-20.4, 33.5] | | | | 0.633 |
| Impact factor | 278 | -15.00 [-19.6, -10.4] | $1.2x10^{-10}$ | 87.7% | 0% | 0.756 |
| (impact factor) | - | -0.08 [-0.6, 0.4] | | | | |
| Citations | 313 | -15.19 [-18.7, -11.7] | $3.2x10^{-17}$ | 87.5% | 0% | 0.908 |
| (citations/year) | - | -0.009 [-0.2, 0.1] | | | | |
| Region of origin (intercept=N. America) | 313 (178) | -15.54 [-19.4, -11.6] | $5.5x10^{-15}$ | 86.6% | 5.3% | 0.010 |
| Asia | 59 | 0.91 [-5.5, 7.3] | | | | 0.782 |
| Europe | 47 | -5.55 [-13.1, 2.0] | | | | 0.148 |
| Latin America | 25 | 2.43 [-6.3, 11.2] | | | | 0.587 |
| Middle East | 4 | 31.15 [11.5, 50.8] | | | | 0.002 |

**Suppl. Table 20 – Three-level meta-regression models for testing all protocol variables on the training dataset, accounting for nesting of experiments within research group.** All variables collected, listed in Suppl. Table 2, are included except the variables specific to re-exposure protocols and sample size calculation due to lack of reporting in all included articles. Doses from the different drugs and routes of administration were z-scored to allow cross-drug comparisons. Grey lines contain sample sizes, intercept effect sizes as absolute mean differences, p-values and $I^2$ for each model, as well as $R^2$ values and Q-test p-values for the moderator. White lines contain betas indicating the additional contribution of each unit/category to the effect size (as well as sample sizes and p-values for individual categories). For categorical variables, reference groups are described in the first column, with the sample size for these groups indicated in parentheses in the second column. $R^2$ values are calculated as the difference between total variance in the model with no moderators and in the tested model, divided by the total variance in the model with no moderators.

| Moderators and categories | Sample size | Effect (% freezing) [95%C.I.] | p-value | $I^2$ | $R^2$ | Moderator p-value |
|---|---|---|---|---|---|---|
| Conditioning type (intercept=contextual) | 168 (61) | -15.67 [-19.5, -11.8] | $1.7x10^{-15}$ | 82.1% | 0% | 0.575 |
| Contextual background | 49 | -4.26 [-10.2, 1.7] | | | | 0.162 |
| Tone | 49 | -1.28 [-7.4, 4.8] | | | | 0.681 |
| Tone-trace | 9 | -1.75 [-14.5, 11.0] | | | | 0.788 |
| Habituation (intercept=unreported) | 168 (136) | -17.89 [-20.6, -15.2] | $7.7x10^{-38}$ | 81.9% | 0% | 0.354 |
| Reported | 32 | 2.89 [-3.2, 9.0] | | | | |
| Handling (intercept=unreported) | 168 (123) | -17.15 [-19.9, -14.4] | $1.6x10^{-33}$ | 82.0% | 0% | 0.812 |
| Reported | 45 | -0.70 [-6.5, 5.1] | | | | |
| # of shocks | 168 | -18.66 [-21.4, -15.9] | $5.1x10^{-40}$ | 81.5% | 2.5% | 0.050 |
| (number) | - | 0.43 [-0.002, 0.9] | | | | |
| Shock intensity | 168 | -14.27 [-20.3, -8.3] | $3.3x10^{-6}$ | 81.9% | 0% | 0.278 |
| (mA) | - | -3.42 [-9.6, 2.8] | | | | |
| Time between drug administration and intervention session | 166 | -19.08 [-21.5, -16.7] | $6.6x10^{-56}$ | 77.4% | 24.9% | $4.3x10^{-7}$ |
| (min) | - | 0.01 [0.009, 0.02] | | | | |
| Time between intervention sessions and test | 167 | -17.66 [-20.3, -15.0] | $5.2x10^{-40}$ | 81.7% | 1.1% | 0.307 |
| (hour) | - | 0.01 [-0.009, 0.03] | | | | |
| Species (intercept=mice) | 168 (90) | -18.64 [-21.8, -15.5] | $6.4x10^{-31}$ | 81.8% | 1.0% | 0.200 |
| Rats | 78 | 3.23 [-1.7, 8.2] | | | | |
| Sex (intercept=both) | 167 (12) | -19.71 [-28.5, -11.0] | $1.0x10^{-5}$ | 82.0% | 0% | 0.563 |
| Male | 155 | 2.69 [-6.4, 11.8] | | | | |
| Age | 75 | -7.95 | 0.474 | 87.2% | 0% | 0.372 |

| | | | | | | |
|---|---|---|---|---|---|---|
| | | [-29.7, 13.8] | | | | |
| (weeks) | - | -0.95<br>[-3.0, 1.1] | | | | |
| Housing | 122 | -17.92<br>[-23.2, -12.7] | $1.9x10^{-11}$ | 76.9% | 0% | 0.881 |
| (animals/cage) | - | 0.13<br>[-1.6, 1.9] | | | | |
| Active drug (intercept=anisomycin) | 168 (140) | -16.74<br>[-19.4, -14.1] | $1.5x10^{-35}$ | 81.7% | 1.4% | 0.242 |
| 4EGI-1 | 1 | 15.16<br>[-14.9, 45.2] | | | | 0.323 |
| Cycloheximide | 27 | -4.66<br>[-11.5, 2.2] | | | | 0.184 |
| Site of injection (intercept=systemic) | 168 (57) | -20.45<br>[-24.7, -16.2] | $5.8x10^{-21}$ | 81.2% | 4.7% | 0.045 |
| Amygdala | 38 | -0.25<br>[-7.5, 7.0] | | | | 0.946 |
| Hippocampus | 41 | 6.54<br>[0.1, 13.0] | | | | 0.046 |
| i.c.v. | 3 | -1.76<br>[-22.3, 18.8] | | | | 0.867 |
| Other | 29 | 9.51<br>[1.9, 17.1] | | | | 0.015 |
| Dose | 161 | -17.31<br>[-19.8, -14.8] | $1.3x10^{-40}$ | 82.1% | 0% | 0.623 |
| (z-score) | - | -0.66<br>[-3.3, 2.0] | | | | |
| Randomization (intercept=unreported) | 168 (155) | -17.38<br>[-19.9, -14.9] | $1.2x10^{-42}$ | 82.0% | 0% | 0.837 |
| Reported | 13 | 1.08<br>[-9.2, 11.3] | | | | |
| Blinding (intercept=unreported) | 168 (56) | -18.61<br>[-22.8, -14.5] | $1.8x10^{-18}$ | 82.0% | 0% | 0.451 |
| Reported | 112 | 1.97<br>[-3.2, 7.1] | | | | |
| Regulatory requirements statement (intercept=unreported) | 168 (21) | -25.42<br>[-32.2, -18.7] | $1.7x10^{-13}$ | 81.1% | 5.4% | 0.012 |
| Reported | 147 | 9.26<br>[2.0, 16.5] | | | | |
| Conflict of interest (Intercept=unreported) | 168 (142) | -13.32<br>[-19.3, -7.3] | $1.4x10^{-5}$ | 81.7% | 1.5% | 0.108 |
| No | 25 | -4.94<br>[-11.5, 1.6] | | | | 0.141 |
| Yes | 1 | 22.97<br>[-12.6, 58.5] | | | | 0.205 |
| Impact factor | 134 | -16.87<br>[-21.6, -12.1] | $4.5x10^{-12}$ | 81.1% | 8.5% | 0.868 |
| (impact factor) | - | -0.06<br>[-0.7, 0.6] | | | | |
| Citations | 168 | -17.49<br>[-20.7, -14.3] | $3.3x10^{-26}$ | 82.0% | 0% | 0.872 |
| (citations/year) | - | 0.02<br>[-0.2, 0.2] | | | | |
| Region of origin (intercept=N. America) | 168 (104) | -16.16<br>[-19.3, -13.0] | $5.1x10^{-24}$ | 81.8% | 1.1% | 0.237 |

| | | | | | | |
|---|---|---|---|---|---|---|
| Asia | 15 | -1.75 [-9.9, 6.4] | | | | 0.674 |
| Europe | 35 | -5.30 [-11.5, 0.9] | | | | 0.094 |
| Latin America | 12 | -1.18 [-10.4, 8.1] | | | | 0.802 |
| Middle East | 2 | 17.68 [-5.1, 40.4] | | | | 0.128 |

**Suppl. Table 21 – Three-level meta-regression models for testing all protocol variables on the reactivation dataset, accounting for nesting of experiments within research group.** All variables collected, listed in Suppl. Table 2, are included. Reexposure duration is based on z-scored values for reexposure to tone (measured in number of tones) and to context (measured in minutes), based on the mean and standard deviation for each type of conditioning. Doses from the different drugs and routes of administration were also z-scored to allow cross-drug comparisons. Grey lines contain sample sizes, intercept effect sizes as absolute mean differences, p-values and $I^2$ for each model, as well as $R^2$ values and Q-test p-values for the moderator. White lines contain betas indicating the additional contribution of each unit/category to the effect size (as well as sample sizes and p-values for individual categories). For categorical variables, reference groups are described in the first column, with the sample size for these groups indicated in parentheses in the second column. $R^2$ values are calculated as the difference between total variance in the model with no moderators and in the tested model, divided by the total variance in the model with no moderators.

| Moderators and categories | Sample size | Effect (% freezing) [95%C.I.] | p-value | $I^2$ | $R^2$ | Moderator p-value |
|---|---|---|---|---|---|---|
| Intervention session (intercept=reexposure) | 145 (3) | -48.13 [-71.7, -24.6] | $6.2x10^{-5}$ | 87.4% | 19.8% | $1.0x10^{-11}$ |
| Extinction | 23 | 57.22 [33.2, 81.3] | | | | $3.1x10^{-6}$ |
| Reconsolidation | 119 | 31.61 [8.3, 54.9] | | | | 0.008 |
| Conditioning type (intercept=contextual) | 145 (77) | -5.71 [-10.6, -0.8] | 0.021 | 87.4% | 19.6% | $2.2x10^{-5}$ |
| Contextual background | 10 | -22.00 [-35.6, -8.4] | | | | 0.001 |
| Tone | 54 | -16.35 [-23.8, -8.9] | | | | $1.6x10^{-5}$ |
| Tone-trace | 4 | -0.36 [-23.1, 22.4] | | | | 0.975 |
| Habituation (intercept=unreported) | 145 (101) | -12.15 [-18.3, -6.0] | $1.1x10^{-4}$ | 90.3% | 0% | 0.966 |
| Reported | 44 | -0.21 [-9.7, 9.3] | | | | |
| Handling (intercept=unreported) | 145 (107) | -13.46 [-19.7, -7.2] | $2.2x10^{-5}$ | 90.2% | 0% | 0.365 |
| Reported | 38 | 3.77 [-4.4, 11.9] | | | | |
| # of shocks | 145 | -10.20 [-15.0, -5.4] | $3.3x10^{-5}$ | 88.7% | 9.1% | 0.006 |
| (number) | - | -0.50 [-0.9, -0.1] | | | | |
| Shock intensity | 142 | -7.38 [-19.3, 4.5] | 0.223 | 90.5% | 0% | 0.410 |
| (mA) | - | -5.41 [-18.3, 7.4] | | | | |

| | | | | | | |
|---|---|---|---|---|---|---|
| Time between training and re-exposure session | 145 | -12.73 [-18.5, -7.0] | $1.5x10^{-5}$ | 90.3% | 0% | 0.343 |
| (hours) | - | 0.006 [-0.006, 0.02] | | | | |
| Time between drug administration and re-exposure session | 137 | -12.64 [-18.5, -6.8] | $2.1x10^{-5}$ | 90.4% | 0% | 0.330 |
| (min) | - | 0.01 [-0.01, 0.03] | | | | |
| Re-exposure duration | 143 | -12.91 [-18.5, -7.3] | $6.1x10^{-6}$ | 88.5% | 8.0% | $2.3x10^{-7}$ |
| (z-score) | - | 8.25 [5.1, 11.4] | | | | |
| Time between re-exposure and test | 145 | -11.08 [-16.8, -5.4] | $1.5x10^{-4}$ | 89.8% | 0% | 0.221 |
| (hour) | - | -0.03 [-0.07, 0.02] | | | | |
| Species (intercept=mice) | 145 (56) | -10.02 [-18.9, -1.2] | 0.026 | 90.4% | 0% | 0.536 |
| Rats | 89 | -3.40 [-14.2, 7.4] | | | | |
| Sex (intercept=both) | 144 (6) | -6.97 [-27.6, 13.7] | 0.509 | 90.4% | 0% | 0.619 |
| Male | 138 | -5.40 [-26.7, 15.9] | | | | |
| Age | 47 | -8.66 [-26.4, 9.0] | 0.337 | 87.8% | 13.7% | 0.584 |
| (weeks) | - | -0.47 [-2.2, 1.2] | | | | |
| Housing | 120 | -15.25 [-26.1, -4.4] | 0.006 | 90.5% | 0% | 0.411 |
| (animals/cage) | - | 1.29 [-1.8, 4.4] | | | | |
| Active drug (intercept=anisomycin) | 145 (130) | -12.08 [-18.3, -5.8] | $1.5x10^{-4}$ | 90.4% | 0% | 0.489 |
| 4EGI-1 | 1 | 23.93 [-15.5, 63.4] | | | | 0.234 |
| Cycloheximide | 14 | -0.75 [-14.6, 13.2] | | | | 0.916 |
| Site of injection (intercept=systemic) | 145 (40) | -14.98 [-23.3, -6.7] | $4.2x10^{-4}$ | 89.0% | 6.7% | 0.004 |
| Amygdala | 56 | -1.29 [-11.9, 9.4] | | | | 0.812 |
| Hippocampus | 30 | 1.94 [-8.4, 12.3] | | | | 0.714 |
| i.c.v. | 6 | 17.78 [-1.1, 36.6] | | | | 0.065 |
| Other | 13 | 19.80 [6.9, 32.7] | | | | 0.003 |
| Dose | 132 | -13.04 [-18.6, -7.5] | $4.1x10^{-6}$ | 90.1% | 0% | 0.215 |
| (z-score) | - | 2.51 [-1.5, 6.5] | | | | |
| Randomization (intercept=unreported) | 145 (143) | -12.19 [-17.9, -6.5] | $2.8x10^{-5}$ | 90.3% | 0% | 0.841 |
| Reported | 2 | -2.92 | | | | |

| | | | | | | |
|---|---|---|---|---|---|---|
| | | [-31.5, 25.7] | | | | |
| Blinding (intercept=unreported) | 145 (43) | -8.29 [-17.3, 0.7] | 0.071 | 90.4% | 0% | 0.274 |
| Reported | 102 | -5.03 [-14.1, 4.0] | | | | |
| Sample size calculation (intercept=unreported) | 145 (143) | -12.26 [-18.1, -6.5] | $3.4x10^{-5}$ | 90.3% | 0% | 0.912 |
| Reported | 2 | 1.76 [-29.4, 32.9] | | | | |
| Regulatory requirements statement (intercept=unreported) | 145 (17) | -7.36 [-19.5, 4.8] | 0.235 | 90.6% | 0% | 0.387 |
| Reported | 128 | -5.13 [-16.8, 6.5] | | | | |
| Conflict of interest (Intercept=unreported) | 145 (107) | -12.26 [-18.8, -5.8] | $2.2x10^{-4}$ | 90.5% | 0% | 0.806 |
| No | 37 | 0.69 [-8.0, 9.3] | | | | 0.875 |
| Yes | 1 | -12.75 [-52.3, 26.8] | | | | 0.527 |
| Impact factor | 144 | -12.35 [-19.5, -4.9] | $8.5x10^{-4}$ | 90.4% | 0% | 0.900 |
| (impact factor) | - | -0.05 [-0.8, 0.7] | | | | |
| Citations | 145 | -12.29 [-18.2, -6.4] | $4.4x10^{-5}$ | 90.3% | 0% | 0.933 |
| (citations/year) | - | 0.009 [-0.2, 0.2] | | | | |
| Region of origin (intercept=N. America) | 145 (74) | -13.99 [-21.7, -6.3] | $3.8x10^{-4}$ | 89.5% | 1.6% | 0.056 |
| Asia | 44 | 1.31 [-8.7, 11.3] | | | | 0.797 |
| Europe | 12 | -7.82 [-23.7, 8.1] | | | | 0.335 |
| Latin America | 13 | 6.64 [-9.0, 22.2] | | | | 0.404 |
| Middle East | 2 | 42.80 [10.8, 74.8] | | | | 0.009 |

**Suppl. Table 22 – Two-level multivariate meta-regression models for testing selected protocol variables on the complete dataset.** The complete list of covariates tested is intervention session, site of injection, active drug, dose, time to test, species, number and intensity of shock, sex, conditioning type and intervention time. Out of 2,048 tested models, we present the best one as selected by AICc. Grey lines contain sample sizes, AICc, Akaike weights, intercept effect sizes as absolute mean differences, p-values and $I^2$ for the model, as well as $R^2$ values and Q-test p-values for the full range of moderators. White lines contain betas indicating the additional contribution of each unit/category to the effect size (as well as sample sizes and p-values for individual variables). Reference categories in the intercept are training session, systemic injection, contextual conditioning and 4EGI-1.

| Model and categories | Sample size | AICc | Weight | Effect size [95% C.I.] | p-value | $I^2$ | $R^2$ | Moderator p-value |
|---|---|---|---|---|---|---|---|---|
| Intervention session + Site of injection + Conditioning type + Active drug + Number of shocks + Intervention time | 277 | 2328.07 | 0.060 | 8.23 [-13.1, 29.6] | 0.450 | 76.4% | 45.1% | $9.6x10^{-24}$ |
| Extinction | 16 | | | 30.06 [22.0, 38.1] | | | | $2.1x10^{-13}$ |
| Reactivation | 3 | | | -21.33 [-42.0, -0.7] | | | | 0.043 |
| Reconsolidation | 101 | | | 3.65 [-0.7, 8.0] | | | | 0.098 |
| Amygdala | 79 | | | -1.77 [-7.3, 3.7] | | | | 0.528 |
| Hippocampus | 63 | | | 1.52 [-3.7, 6.8] | | | | 0.572 |
| i.c.v. | 8 | | | 4.11 [-7.2, 15.4] | | | | 0.476 |
| Other | 37 | | | 13.34 [6.9, 19.8] | | | | $4.6x10^{-5}$ |
| Contextual background | 59 | | | -6.34 [-11.5, -1.2] | | | | 0.015 |
| Tone | 90 | | | -4.52 [-9.7, 0.6] | | | | 0.085 |
| Tone-trace | 13 | | | -0.75 [-11.4, 9.9] | | | | 0.891 |
| Anisomycin | 237 | | | -25.06 [-46.1, -4.0] | | | | 0.019 |
| Cycloheximide | 38 | | | -25.50 [-47.0, -4.0] | | | | 0.020 |
| (number of shocks) | - | | | -0.39 [-0.6, -0.15] | | | | 0.002 |
| (time in min) | - | | | 0.01 [0.009, 0.02] | | | | $6.4x10^{-7}$ |

**Suppl. Table 23 – Three-level multivariate meta-regression models for testing selected protocol variables on the complete dataset, considering the nesting of experiments within articles.** The complete list of covariates tested is: intervention session, site of injection, active drug, dose, time to test, species, number and intensity of shock, sex, conditioning type and intervention time. Out of 2,048 tested models, we present the best one as selected by AICc. Grey lines contain sample sizes, AICc, Akaike weights, intercept effect sizes as absolute mean differences, p-values and $I^2$ for the model, as well as $R^2$ values and Q-test p-values for the full range of moderators. White lines contain betas indicating the additional contribution of each unit/category to the effect size (as well as sample sizes and p-values for individual variables). Reference categories in the intercept are training session, systemic injection, and 4EGI-1.

| Model and categories | Sample size | AICc | Weight | Effect size [95% C.I.] | p-value | $I^2$ | $R^2$ | Moderator p-value |
|---|---|---|---|---|---|---|---|---|
| Intervention session + Site of injection + Active drug + Number of shocks + Intervention time | 277 | 2328.77 | 0.056 | 3.85 [-18.3, 26.0] | 0.734 | 79.0% | 44.9% | $2.8x10^{-22}$ |
| Extinction | 16 | | | 32.44 [24.3, 40.6] | | | | $7.6x10^{-15}$ |
| Reactivation | 3 | | | -21.77 [-42.8, -0.8] | | | | 0.042 |
| Reconsolidation | 101 | | | 5.46 [1.1, 9.8] | | | | 0.014 |
| Amygdala | 79 | | | -2.60 [-8.0, 2.8] | | | | 0.343 |
| Hippocampus | 63 | | | 3.47 [-2.4, 9.3] | | | | 0.248 |
| i.c.v. | 8 | | | 5.29 [-6.3, 16.9] | | | | 0.372 |
| Other | 37 | | | 15.26 [8.5, 22.1] | | | | $1.1x10^{-5}$ |
| Anisomycin | 237 | | | -24.80 [-46.7, -2.9] | | | | 0.026 |
| Cycloheximide | 38 | | | -24.94 [-47.6, -2.3] | | | | 0.031 |
| (number of shocks) | - | | | -0.44 [-0.7, -0.2] | | | | 0.001 |
| (time in min) | - | | | 0.015 [0.009, 0.02] | | | | $3.5x10^{-7}$ |

**Suppl. Table 24 – Three-level multivariate meta-regression models for testing selected protocol variables on the complete dataset, considering the nesting of experiments within groups.** The complete list of covariates tested is: intervention session, site of injection, active drug, dose, time to test, species, number and intensity of shock, sex, conditioning type and intervention time. Out of 2,048 tested models, we present the best one as selected by AICc. Grey lines contain sample sizes, AICc, Akaike weights, intercept effect sizes as absolute mean differences, p-values and $I^2$ for the model, as well as $R^2$ values and Q-test p-values for the full range of moderators. White lines contain betas indicating the additional contribution of each unit/category to the effect size (as well as sample sizes and p-values for individual variables). Reference categories in the intercept are training session, systemic injection, and 4EGI-1.

| Model and categories | Sample size | AICc | Weight | Effect size [95% C.I.] | p-value | $I^2$ | $R^2$ | Moderator p-value |
|---|---|---|---|---|---|---|---|---|
| Intervention session + Site of injection + Active drug + Number of shocks + Intervention time | 277 | 2326.69 | 0.081 | 2.68 [-18.5, 23.8] | 0.804 | 79.7% | 44.0% | $8.4 \times 10^{-23}$ |
| Extinction | 16 | | | 31.95 [23.8, 40.1] | | | | $1.8 \times 10^{-14}$ |
| Reactivation | 3 | | | -22.77 [-43.4, -2.1] | | | | 0.031 |
| Reconsolidation | 101 | | | 6.03 [1.6, 10.8] | | | | 0.008 |
| Amygdala | 79 | | | -3.23 [-9.8, 3.4] | | | | 0.337 |
| Hippocampus | 63 | | | 4.32 [-2.1, 10.8] | | | | 0.189 |
| i.c.v. | 8 | | | 6.80 [-5.3, 18.7] | | | | 0.264 |
| Other | 37 | | | 15.88 [8.5, 23.3] | | | | $2.7 \times 10^{-5}$ |
| Anisomycin | 237 | | | -24.45 [-45.2, -3.7] | | | | 0.021 |
| Cycloheximide | 38 | | | -23.17 [-44.8, -1.5] | | | | 0.036 |
| (number of shocks) | - | | | -0.44 [-0.7, -0.2] | | | | 0.003 |
| (time in min) | - | | | 0.016 [0.01, 0.02] | | | | $8.6 \times 10^{-8}$ |

**Suppl. Table 25 – Two-level multivariate meta-regression models for testing selected protocol variables on the training dataset.** The complete list of covariates tested is: site of injection, active drug, dose, time to test, species, number and intensity of shock, sex, conditioning type and intervention time. Out of 1,024 tested models, we present the best one as selected by AICc. Grey lines contain sample sizes, AICc, Akaike weights, intercept effect sizes as absolute mean differences, p-values and $I^2$ for the model, as well as $R^2$ values and Q-test p-values for the full range of moderators. White lines contain betas indicating the additional contribution of each unit/category to the effect size (as well as sample sizes and p-values for individual variables). Reference category in the intercept is systemic injection.

| Model and categories | Sample size | AICc | Weight | Effect size [95% C.I.] | p-value | $I^2$ | $R^2$ | Moderator p-value |
|---|---|---|---|---|---|---|---|---|
| Site of injection + Intervention time | 157 | 1299.22 | 0.037 | -21.00 [-24.7, -17.3] | $2.9x10^{-29}$ | 73.1% | 27.7 % | $1.2x10^{-6}$ |
| Amygdala | 36 | | | -0.63 [-7.0, 5.7] | | | | 0.845 |
| Hippocampus | 39 | | | 3.44 [-2.3, 9.2] | | | | 0.241 |
| i.c.v. | 3 | | | -8.80 [-27.7, 10.1] | | | | 0.363 |
| Other | 28 | | | 8.78 [2.1, 15.5] | | | | 0.010 |
| (time in min) | - | | | 0.016 [0.01, 0.02] | | | | $9.8x10^{-8}$ |

**Suppl. Table 26 – Three-level multivariate meta-regression models for testing selected protocol variables on the training dataset, considering nesting of experiments within articles.** The complete list of covariates tested is: site of injection, active drug, dose, time to test, species, number and intensity of shock, sex, conditioning type and intervention time. Out of 1,024 tested models, we present the best one as selected by AICc. Grey lines contain sample sizes, AICc, Akaike weights, intercept effect sizes as absolute mean differences, p-values and $I^2$ for the model, as well as $R^2$ values and Q-test p-values for the full range of moderators. White lines contain betas indicating the additional contribution of each unit/category to the effect size (as well as sample sizes and p-values for individual variables). Reference category in the intercept is systemic injection.

| Model and categories | Sample size | AICc | Weight | Effect size [95% C.I.] | p-value | $I^2$ | $R^2$ | Moderator p-value |
|---|---|---|---|---|---|---|---|---|
| Site of injection + Intervention time | 157 | 1301.44 | 0.036 | -21.00 [-24.7, -17.3] | $2.9x10^{-29}$ | 75.4% | 30.7% | $1.2x10^{-6}$ |
| Amygdala | 36 | | | -0.63 [-7.0, 5.7] | | | | 0.845 |
| Hippocampus | 39 | | | 3.44 [-2.3, 9.2] | | | | 0.241 |
| i.c.v. | 3 | | | -8.80 [-27.7, 10.1] | | | | 0.363 |
| Other | 28 | | | 8.78 [2.1, 15.5] | | | | 0.010 |
| (time in min) | - | | | 0.016 [0.01, 0.02] | | | | $9.8x10^{-8}$ |

**Suppl. Table 27 – Three-level multivariate meta-regression models for testing selected protocol variables on the training dataset, considering nesting of experiments within groups.** The complete list of covariates tested is: site of injection, active drug, dose, time to test, species, number and intensity of shock, sex, conditioning type and intervention time. Out of 1,024 tested models, we present the best one as selected by AICc. Grey lines contain sample sizes, AICc, Akaike weights, intercept effect sizes as absolute mean differences, p-values and $I^2$ for the model, as well as $R^2$ values and Q-test p-values for the full range of moderators. White lines contain betas indicating the additional contribution of each unit/category to the effect size (as well as sample sizes and p-values for individual variables). Reference category in the intercept is systemic injection.

| Model and categories | Sample size | AICc | Weight | Effect size [95% C.I.] | p-value | $I^2$ | $R^2$ | Moderator p-value |
|---|---|---|---|---|---|---|---|---|
| Site of injection + Intervention time | 157 | 1301.44 | 0.037 | -21.01 [-24.7, -17.3] | $5.9\times10^{-29}$ | 75.4% | 30.7% | $1.2\times10^{-6}$ |
| Amygdala | 36 | | | -0.65 [-7.0, 5.7] | | | | 0.843 |
| Hippocampus | 39 | | | 3.45 [-2.3, 9.2] | | | | 0.241 |
| i.c.v. | 3 | | | -8.81 [-27.8, 10.1] | | | | 0.362 |
| Other | 28 | | | 8.79 [2.1, 15.5] | | | | 0.011 |
| (time in min) | - | | | 0.016 [0.01, 0.02] | | | | $9.9\times10^{-8}$ |

**Suppl. Table 28 – Two-level multivariate meta-regression models for testing selected protocol variables on the reactivation dataset.** The complete list of covariates tested is: site of injection, re-exposure duration, active drug, dose, time to test, time to re-exposure, species, number and intensity of shock and sex. Out of 1,024 tested models, we present the best one as selected by AICc. Grey lines contain sample sizes, AICc, Akaike weights, intercept effect sizes as absolute mean differences, p-values and $I^2$ for the model, as well as $R^2$ values and Q-test p-values for the full range of moderators. White lines contain betas indicating the additional contribution of each unit/category to the effect size (as well as sample sizes and p-values for individual variables). Reference category in the intercept is systemic injection.

| Model and categories | Sample size | AICc | Weight | Effect size [95% C.I.] | p-value | $I^2$ | $R^2$ | Moderator p-value |
|---|---|---|---|---|---|---|---|---|
| Site of injection + Re-exposure duration + Number of shocks | 126 | 1090.89 | 0.117 | -12.71 [-17.9, -7.5] | $1.7\times10^{-6}$ | 81.3% | 45.2% | $5.1\times10^{-14}$ |
| Amygdala | 51 | | | -3.50 [-10.5, 3.5] | | | | 0.330 |
| Hippocampus | 24 | | | 8.83 [-0.5, 18.2] | | | | 0.064 |
| i.c.v. | 5 | | | 14.29 [-1.5, 30.1] | | | | 0.076 |
| Other | 9 | | | 27.16 [14.7, 39.6] | | | | $2.0\times10^{-5}$ |
| (duration, z-score) | - | | | 8.05 [4.8, 11.3] | | | | $1.4\times10^{-6}$ |
| (number of shocks) | - | | | -0.58 [-0.9, -0.3] | | | | $4.1\times10^{-5}$ |

**Suppl. Table 29 – Three-level multivariate meta-regression models for testing selected protocol variables on the reactivation dataset, considering nesting of experiments within articles.** The complete list of covariates tested is: site of injection, re-exposure duration, active drug, dose, time to test, time to re-exposure, species, number and intensity of shock and sex. Out of 1,024 tested models, we present the best one as selected by AICc. Grey lines contain sample sizes, AICc, Akaike weights, intercept effect sizes as absolute mean differences, p-values and $I^2$ for the model, as well as $R^2$ values and Q-test p-values for the full range of moderators. White lines contain betas indicating the additional contribution of each unit/category to the effect size (as well as sample sizes and p-values for individual variables). Reference category in the intercept is systemic injection.

| Model and categories | Sample size | AICc | Weight | Effect size [95% C.I.] | p-value | $I^2$ | $R^2$ | Moderator p-value |
|---|---|---|---|---|---|---|---|---|
| Site of injection + Re-exposure duration + Number of shocks | 126 | 1092.45 | 0.124 | -13.36 [-19.5, -7.2] | $1.9 \times 10^{-5}$ | 82.6% | 43.6% | $7.3 \times 10^{-13}$ |
| Amygdala | 51 | | | -2.33 [-10.4, 5.7] | | | | 0.569 |
| Hippocampus | 24 | | | 8.99 [-1.1, 19.1] | | | | 0.082 |
| i.c.v. | 5 | | | 12.84 [-3.5, 29.2] | | | | 0.123 |
| Other | 9 | | | 28.01 [14.8, 41.2] | | | | $3.3 \times 10^{-5}$ |
| (duration, z score) | - | | | 8.43 [5.2, 11.7] | | | | $3.6 \times 10^{-7}$ |
| (number of shocks) | - | | | -0.57 [-0.9, -0.3] | | | | $3.3 \times 10^{-4}$ |

**Suppl. Table 30 – Three-level multivariate meta-regression models for testing selected protocol variables on the reactivation dataset, considering nesting of experiments within groups.** The complete list of covariates tested is: site of injection, re-exposure duration, active drug, dose, time to test, time to re-exposure, species, number and intensity of shock and sex. Out of 2,048 tested models, we present the best one as selected by AICc. Grey lines contain sample sizes, AICc, Akaike weights, intercept effect sizes as absolute mean differences, p-values and $I^2$ for the model, as well as $R^2$ values and Q-test p-values for the full range of moderators. White lines contain betas indicating the additional contribution of each unit/category to the effect size (as well as sample sizes and p-values for individual variables). Reference category in the intercept is systemic injection.

| Model and categories | Sample size | AICc | Weight | Effect size [95% C.I.] | p-value | $I^2$ | $R^2$ | Moderator p-value |
|---|---|---|---|---|---|---|---|---|
| Site of injection + Re-exposure duration + Number of shocks | 126 | 1093.21 | 0.122 | -12.71 [-17.9, -7.5] | $1.7 \times 10^{-6}$ | 82.6% | 47.4% | $5.1 \times 10^{-14}$ |
| Amygdala | 51 | | | -3.50 [-10.5, 3.5] | | | | 0.330 |
| Hippocampus | 24 | | | 8.83 [-0.5, 18.2] | | | | 0.064 |
| i.c.v. | 5 | | | 14.29 [-1.5, 30.1] | | | | 0.076 |
| Other | 9 | | | 27.16 [14.7, 39.6] | | | | $2.0 \times 10^{-5}$ |
| (duration, z score) | - | | | 8.05 [4.8, 11.3] | | | | $1.4 \times 10^{-6}$ |
| (number of shocks) | - | | | -0.58 [-0.9, -0.3] | | | | $4.1 \times 10^{-5}$ |
