## Supplementary File 2 for "A meta-analysis of the effect of protein synthesis inhibitors on rodent fear conditioning": Carneiro-Amorim---Amaral-2022---bioRxiv-v2---Supplementary-File-2.Rmd

````
---
title: "Meta-analyses of interventions on protein synthesis in fear conditioning experiments in rodents"
author: "Clarissa F. D. Carneiro"
date: "10/08/2022"
output: html_document
---
```{r setup, include=FALSE}
memory.limit(50000)
knitr::opts_chunk$set(echo = FALSE)
```

Packages used:
```{r echo=TRUE, warning=FALSE, message=FALSE}
library(readxl)
library(tidyr)
library(dplyr)
library(ggplot2)
library(cowplot)
library(metafor)
library(metaviz)
library(ggforce)
library(viridis)
library(igraph)
library(glmulti)
library(purrr)
```

Importing dataset and excluding:

- article retracted after data collection

- studies with missing sample size 

- protocols out of our criteria: test freezing not measured during tone or unclear test protocol OR reexposure protocol includes the footshock (coded in the dataset with *) 

- protocols out of our criteria: repeated testing

Plus, additional data cleaning of studies with missing protocol data.
```{r warning=FALSE}
full_data = read_excel("Supplementary File 1 - revisado.xlsx", sheet = 1)
full_data = full_data %>% filter(full_data$`Article code`!=83)
full_data = full_data %>% filter(full_data$`Test number`==1)
full_data = full_data %>% filter(full_data$ConditioningType!="Tone*")
full_data = full_data %>% filter(full_data$ConditioningType!="Tone-Trace*")
full_data = full_data %>% filter(full_data$`Intervention Session`!="Reconsolidation*")
full_data = full_data %>% filter(full_data$nControl>0)
full_data$`Time to reexposure` = ifelse(full_data$`Time to reexposure`==0, NA, full_data$`Time to reexposure`)
full_data$InterventionTime = as.numeric(full_data$InterventionTime)
full_data$Dose_mgkg = as.numeric(full_data$Dose_mgkg)
full_data$Dose_ugside = as.numeric(full_data$Dose_ugside)
```

**Final inclusion numbers:**
```{r}
comparisons_by_articles = full_data %>% count(full_data$`Article code`)
```

`r nrow(comparisons_by_articles)` articles and `r sum(comparisons_by_articles$n)` comparisons

**Research groups:**

The list of doi associated with their comparison codes was exported and the list of authors was obtained using Matlab code from Moulin & Amaral (2020), doi: 10.1002/jrsm.1430. 
```{r}
#input_author_network = full_data %>% select(c("Comparison.Code","Doi"))
#write.csv2(input_author_network, file = "input_author_network.csv", row.names = FALSE)

authors = read_excel("Supplementary File 1 - revisado.xlsx", sheet = 2)
authors_article = authors %>% select(-c("Comparison.Code"))
authors_article = distinct(authors_article)
authors_article = authors_article %>% arrange(PMID,Authors)
```

Then, we got the pairs of authors (code from https://towardsdatascience.com/simple-iterative-programming-and-error-handling-in-r-a68ebccdf9b9)
```{r}
unique_pairs <- function(char_vector = NULL) {
  
  vector <- as.character(unique(char_vector))
  
  df <- data.frame(from = character(), to = character(), stringsAsFactors = FALSE)
  
  if (length(vector) > 1) {
    for (i in 1:(length(vector) - 1)) {
      from <- rep(vector[i], length(vector) - i) 
      to <- vector[(i + 1): length(vector)]
      
      df <- df %>% 
        dplyr::bind_rows(
          data.frame(from = from, to = to, stringsAsFactors = FALSE) 
        )
    }
  }

  df
  
}

author_pairs = data.frame(from = character(), to = character(), stringsAsFactors = FALSE)
PMID_list = unique(authors_article$PMID)
for (i in 1:91) {
  author_pairs_new = authors_article %>% filter(PMID == PMID_list[i]) %>% pull(Authors) %>% unique_pairs()
  
  author_pairs = author_pairs %>% bind_rows(author_pairs_new)
}

# add up number of shared comparisons to get weights
edges_comparisons = author_pairs %>% count(from, to, name = "weight")
```
and prepared the network matrix.
```{r}
edgelist_matrix = as.matrix(edges_comparisons[ ,c("from", "to")])
authors_graph = graph_from_edgelist(edgelist_matrix, directed = FALSE) %>% set.edge.attribute("weight", value = edges_comparisons$weight)
```

Next, we detected communities
```{r}
finding_groups = cluster_louvain(authors_graph, weights = E(authors_graph)$weight)

V(authors_graph)$community = finding_groups$membership
authors_groups = data.frame(Authors = V(authors_graph)$name, group = V(authors_graph)$community)
```
and added the results of modularity analyses to the full dataset. Comparisons with authors from more than one group are assigned to the group of the majority of authors.
```{r}
modularity_results = full_join(authors, authors_groups, by = "Authors")

getmode = function(v) {
   uniqv = unique(v)
   uniqv[which.max(tabulate(match(v, uniqv)))]
}

modularity_results2 = modularity_results %>% group_by(Comparison.Code) %>% summarise(group = getmode(group))

full_data = full_join(full_data, modularity_results2, by = "Comparison.Code")
```

Alternative citation metrics (https://journals.plos.org/plosbiology/article?id=10.1371/journal.pbio.1002541)
```{r}
pmid_by_comp_code = modularity_results %>% select(PMID, Comparison.Code)
pmid_by_comp_code = distinct(pmid_by_comp_code)

icite_data = read_excel("Supplementary File 1 - revisado.xlsx", sheet = 3)
icite_data_comp_code = full_join(pmid_by_comp_code, icite_data, by = "PMID")

full_data = full_join(full_data, icite_data_comp_code, by = "Comparison.Code")
```
```{r}
a_data_cit_old = full_data %>% group_by(`Article code`) %>% summarise(CPY=mean(`Citations per year`), ANO=mean(`Year of publication`))
a_data_cit_old$CPY = as.numeric(a_data_cit_old$CPY)

ggplot(a_data_cit_old, aes(x=CPY)) + geom_histogram(binwidth = 5, fill = "black", color="white") + labs(x = "Citations per year", y = "Number of articles") + theme_classic() + theme(axis.text = element_text(color = "black", size = 8), axis.title = element_text(color = "black", size = 12), plot.margin = margin(0.5,0,0,0.5, "cm"))
ggsave("citations_old_histo.png")

ggplot(a_data_cit_old, aes(x=ANO, y=CPY)) + 
  geom_point() +
  theme_classic()
ggsave("citations_old.png")

a_data_cit23 = full_data %>% group_by(`Article code`) %>% summarise(CPY=mean(CitationsPerYear_23), ANO=mean(`Year of publication`))
a_data_cit23$CPY = as.numeric(a_data_cit23$CPY)

ggplot(a_data_cit23, aes(x=CPY)) + geom_histogram(binwidth = 5, fill = "black", color="white") + labs(x = "Citations per year", y = "Number of articles") + theme_classic() + theme(axis.text = element_text(color = "black", size = 8), axis.title = element_text(color = "black", size = 12), plot.margin = margin(0.5,0,0,0.5, "cm"))
ggsave("citations_icite_histo.png")

ggplot(a_data_cit23, aes(x=ANO, y=CPY)) + 
  geom_point() +
  theme_classic()
ggsave("citations_icite.png")

a_data_rcr = full_data %>% group_by(`Article code`) %>% summarise(RCR=mean(RCR), ANO=mean(`Year of publication`))
a_data_rcr$RCR = as.numeric(a_data_rcr$RCR)

ggplot(a_data_rcr, aes(x=RCR)) + geom_histogram(binwidth = 5, fill = "black", color="white") + labs(x = "RCR", y = "Number of articles") + theme_classic() + theme(axis.text = element_text(color = "black", size = 8), axis.title = element_text(color = "black", size = 12), plot.margin = margin(0.5,0,0,0.5, "cm"))
ggsave("citations_rcr_histo.png")

ggplot(a_data_rcr, aes(x=ANO, y=RCR)) + 
  geom_point() +
  theme_classic()
ggsave("citations_rcr.png")

a_data_rcr_cit = full_data %>% group_by(`Article code`) %>% summarise(RCR=mean(RCR), ANO=mean(`Year of publication`), CPY=mean(CitationsPerYear_23))

ggplot(a_data_rcr_cit, aes(x=as.numeric(CPY), y=as.numeric(RCR))) + 
  geom_point() +
  theme_classic()
ggsave("correlation_rcr.png")

ggplot(a_data_rcr_cit %>% filter(CPY<30), aes(x=as.numeric(CPY), y=as.numeric(RCR))) + 
  geom_point() +
  theme_classic()
ggsave("correlation_rcr_menor30.png")

ggplot(a_data_rcr_cit %>% filter(CPY<20), aes(x=as.numeric(CPY), y=as.numeric(RCR))) + 
  geom_point() +
  theme_classic()
ggsave("correlation_rcr_menor20.png")

ggplot(a_data_rcr_cit %>% filter(CPY<10), aes(x=as.numeric(CPY), y=as.numeric(RCR))) + 
  geom_point() +
  theme_classic()
ggsave("correlation_rcr_menor10.png")
```

**Article features**

Region of origin
```{r}
a_data_region = full_data %>% group_by(`Article code`) %>% count(`Region of origin`)
a_data_region %>% group_by(`Region of origin`) %>% count(`Region of origin`)
a_data_region[a_data_region=="North America"] = "N. America"
a_data_region[a_data_region=="Latin America"] = "L. America"

region = ggplot(a_data_region, aes(x = factor(`Region of origin`, level = c("N. America", "Asia", "Europe", "L. America", "Middle East")))) + geom_bar(color="white", fill="black") + labs(x = "", y = "Number of articles") + theme_classic() + theme(axis.text = element_text(color = "black", size = 8), axis.title = element_text(color = "black", size = 12), plot.margin = margin(0.5,0,0,0, "cm"))
```
Year of publication
```{r}
a_data_yr = full_data %>% group_by(`Article code`) %>% summarise(YR=mean(`Year of publication`))
a_data_yr$YR = as.numeric(a_data_yr$YR)

year_publ = ggplot(a_data_yr, aes(x=YR)) + geom_histogram(binwidth = 2, fill = "black", color="white") + labs(x = "Year of publication", y = "Number of articles") + theme_classic() + theme(axis.text = element_text(color = "black", size = 8), axis.title = element_text(color = "black", size = 12), plot.margin = margin(0.5,0,0,0.5, "cm")) + scale_y_continuous(breaks = c(0,3,6,9,12,15)) + scale_x_continuous(breaks = c(2000,2005,2010,2015))

a_data_yr %>% summarise(median = median(a_data_yr$YR, na.rm=TRUE), min = min(a_data_yr$YR, na.rm=TRUE), max = max(a_data_yr$YR, na.rm=TRUE))
```
Impact Factor
```{r}
a_data_if = full_data %>% group_by(`Article code`) %>% summarise(IF=mean(`Impact Factor`))
a_data_if$IF = as.numeric(a_data_if$IF)

impact_factor = ggplot(a_data_if, aes(x=IF)) + geom_histogram(binwidth = 2, fill = "black", color="white") + labs(x = "Impact factor", y = "Number of articles") + theme_classic() + theme(axis.text = element_text(color = "black", size = 8), axis.title = element_text(color = "black", size = 12), plot.margin = margin(0.5,0,0,0, "cm")) + scale_x_continuous(breaks=c(0, 5, 10, 15, 20, 25, 30, 35))

a_data_if %>% summarise(median = median(a_data_if$IF, na.rm=TRUE), min = min(a_data_if$IF, na.rm=TRUE), max = max(a_data_if$IF, na.rm=TRUE))
```
Citations per year
```{r}
a_data_cit = full_data %>% group_by(`Article code`) %>% summarise(CIT=mean(`Citations per year`))
a_data_cit$CIT = as.numeric(a_data_cit$CIT)

cit_per_year = ggplot(a_data_cit, aes(x=CIT)) + geom_histogram(binwidth = 5, fill = "black", color="white") + labs(x = "Citations per year", y = "Number of articles") + theme_classic() + theme(axis.text = element_text(color = "black", size = 8), axis.title = element_text(color = "black", size = 12), plot.margin = margin(0.5,0,0,0.5, "cm"))

a_data_cit %>% summarise(median = median(a_data_cit$CIT, na.rm=TRUE), min = min(a_data_cit$CIT, na.rm=TRUE), max = max(a_data_cit$CIT, na.rm=TRUE))
```
Suppl. Fig. 1
```{r, warning=FALSE}
plot_grid(region, year_publ, impact_factor, cit_per_year, nrow = 2, labels = LETTERS)
ggsave("FIG_S1.png", width = 20, height = 20, units = "cm")
```

Risk of bias (Suppl. Table 1)
```{r}
a_data = full_data %>% distinct(full_data$`Article code`, .keep_all = TRUE) 
a_data %>% count(Randomization)
a_data %>% count(Blinding)
a_data %>% filter(Blinding==TRUE) %>% count(FreezingSoftware) 
a_data %>% count(SampleSizeCalc)
a_data %>% count(RegulatoryReq)
a_data %>% count(ConflictInterest)
```

**Experiment features**
(Suppl. Table 2 and Table 1)
```{r, warning=FALSE}
full_data %>% group_by(`Intervention Session`) %>% count(`Intervention Session`)
full_data %>% group_by(`Site of injection`) %>% count(`Site of injection`)
full_data %>% group_by(ConditioningType) %>% count(ConditioningType)
full_data %>% group_by(Habituation) %>% count(Habituation)
full_data %>% group_by(Handling) %>% count(Handling)
full_data %>% summarise(median(NumberShock, na.rm = TRUE), min(NumberShock, na.rm=TRUE), max(NumberShock, na.rm=TRUE))
full_data %>% summarise(median(ShockIntensity, na.rm=TRUE), min(ShockIntensity, na.rm=TRUE), max(ShockIntensity, na.rm=TRUE))
full_data %>% summarise(median(InterventionTime, na.rm = TRUE), min(InterventionTime, na.rm = TRUE), max(InterventionTime, na.rm = TRUE))
full_data %>% filter(`Intervention Session`!="Training") %>% summarise(median(`Time to reexposure`, na.rm=TRUE), min(`Time to reexposure`, na.rm=TRUE), max(`Time to reexposure`, na.rm=TRUE))
full_data_context = full_data %>% filter(ConditioningType=="Contextual"|ConditioningType=="Contextual background")
full_data_context %>% filter(`Intervention Session`!="Training") %>% summarise(median(ReexposureDuration_min, na.rm = TRUE), min(ReexposureDuration_min, na.rm = TRUE), max(ReexposureDuration_min, na.rm = TRUE))
full_data_tone = full_data %>% filter(ConditioningType=="Tone"|ConditioningType=="Tone-Trace")
full_data_tone %>% filter(`Intervention Session`!="Training") %>% summarise(median(ReexposureDuration_number, na.rm = TRUE), min(ReexposureDuration_number, na.rm = TRUE), max(ReexposureDuration_number, na.rm = TRUE))
full_data %>% summarise(median(TimetoTest, na.rm=TRUE), min(TimetoTest, na.rm=TRUE), max(TimetoTest, na.rm=TRUE))
full_data %>% group_by(Species) %>% count(Species)
full_data %>% group_by(Sex) %>% count(Sex)
full_data %>% summarise(median(Age_mean, na.rm=TRUE), min(Age_mean, na.rm=TRUE), max(Age_mean, na.rm=TRUE))
full_data %>% summarise(median(Housing, na.rm=TRUE), min(Housing, na.rm=TRUE), max(Housing, na.rm=TRUE))
full_data %>% group_by(`Active Drug`) %>% count(`Active Drug`)
full_data %>% group_by(`Active Drug`) %>% summarise(median(Dose_mgkg, na.rm = TRUE), min(Dose_mgkg, na.rm = TRUE), max(Dose_mgkg, na.rm = TRUE))
full_data %>% group_by(`Active Drug`) %>% summarise(median(Dose_ugside, na.rm = TRUE), min(Dose_ugside, na.rm = TRUE), max(Dose_ugside, na.rm = TRUE))
full_data %>% group_by(`Active Drug`, `Route of administration`) %>% count(is.na(Dose_mgkg))
full_data %>% group_by(`Active Drug`, `Route of administration`) %>% count(is.na(Dose_ugside))
```

```{r}
ggplot(full_data, aes(x = `Active Drug`, y = as.numeric(Dose_mgkg), color = `Active Drug`)) + 
  geom_jitter(  ) + 
  theme_classic()

ggplot(full_data, aes(x = `Active Drug`, y = as.numeric(Dose_ugside), color = `Active Drug`)) + 
  geom_jitter(  ) + 
  theme_classic()
```

**General adjustments for meta-analyses**

Given that we are not considering the distinction between the different sites for systemic injections, we combined them.
```{r}
full_data[full_data =="i.p."] = "Systemic"
full_data[full_data =="s.c."] = "Systemic"
full_data[full_data =="i.p. or s.c."] = "Systemic"
```
In some analyses we want to combine sites of injections that don't appear often as a single "other" category, so we created a new variable.
```{r}
full_data = full_data %>% mutate(Site.injection.other = if_else(`Site of injection`!="Systemic" & `Site of injection`!="Amygdala" & `Site of injection`!="Hippocampus" & `Site of injection`!="i.c.v.", "Other", `Site of injection`))
```
We are combining contextual and tone tests performed on the same animals. To account for this duplication of the same cohort, we divided the sample size in both comparisons. 
```{r}
full_data = full_data %>% mutate(adj_nControl = nControl) %>% mutate(adj_nTreated = nTreated) %>% mutate(repeated_cohort = 0)
full_data = full_data %>% group_by(full_data$`Cohort Code`)
full_data = full_data %>% mutate(repeated_cohort = (duplicated(`Cohort Code`)|duplicated(`Cohort Code`, fromLast = TRUE)))
full_data$adj_nControl = ifelse(full_data$repeated_cohort==TRUE, full_data$nControl/2, full_data$nControl)
full_data$adj_nTreated = ifelse(full_data$repeated_cohort==TRUE, full_data$nTreated/2, full_data$nTreated)
```
Some variables are present in different scales depending on other characteristics of the experiment (dose and reexposure duration), so we normalized these values to a single scale.
```{r, warning=FALSE}
#doses = full_data %>% select(c(Comparison.Code, `Site of injection`, `Active Drug`, Dose_mgkg, Dose_ugside))

#doses_a_s = doses %>% filter(`Active Drug`=="Anisomycin" & `Site of injection`=="Systemic" & is.na(Dose_mgkg)==FALSE)
#doses_a_s = doses_a_s %>% mutate(dose_zscore = ((Dose_mgkg-mean(doses_a_s$Dose_mgkg))/sd(doses_a_s$Dose_mgkg)))

#doses_a_i = doses %>% filter(`Active Drug`=="Anisomycin" & `Site of injection`!="Systemic" & is.na(Dose_ugside)==FALSE)
#doses_a_i = doses_a_i %>% mutate(dose_zscore = ((Dose_ugside-mean(doses_a_i$Dose_ugside))/sd(doses_a_i$Dose_ugside)))

#doses_c_s = doses %>% filter(`Active Drug`=="Cycloheximide" & `Site of injection`=="Systemic" & is.na(Dose_mgkg)==FALSE)
#doses_c_s = doses_c_s %>% mutate(dose_zscore = ((Dose_mgkg-mean(doses_c_s$Dose_mgkg))/sd(doses_c_s$Dose_mgkg)))

#doses_c_i = doses %>% filter(`Active Drug`=="Cycloheximide" & `Site of injection`!="Systemic" & is.na(Dose_ugside)==FALSE)
#doses_c_i = doses_c_i %>% mutate(dose_zscore = ((Dose_ugside-mean(doses_c_i$Dose_ugside))/sd(doses_c_i$Dose_ugside)))

#doses_4_i = doses %>% filter(`Active Drug`=="4EGI-1" & `Site of injection`!="Systemic" & is.na(Dose_ugside)==FALSE)
#doses_4_i = doses_4_i %>% mutate(dose_zscore = 0)

#doses_z = rbind(doses_a_s, doses_a_i, doses_c_s, doses_c_i, doses_4_i)
#doses_z = doses_z %>% select(c(Comparison.Code, dose_zscore))

#full_data = left_join(full_data, doses_z, by = "Comparison.Code")
```

```{r}
#z score doses por especie
doses_species = full_data %>% select(c(Comparison.Code, `Site of injection`, `Active Drug`, Dose_mgkg, Dose_ugside, Species))

doses_a_s_rats = doses_species %>% filter(`Active Drug`=="Anisomycin" & `Site of injection`=="Systemic" & is.na(Dose_mgkg)==FALSE & Species=="rats")
doses_a_s_rats = doses_a_s_rats %>% mutate(dose_zscore_s = ((Dose_mgkg-mean(doses_a_s_rats$Dose_mgkg))/sd(doses_a_s_rats$Dose_mgkg)))

doses_a_s_mice = doses_species %>% filter(`Active Drug`=="Anisomycin" & `Site of injection`=="Systemic" & is.na(Dose_mgkg)==FALSE & Species=="mice")
doses_a_s_mice = doses_a_s_mice %>% mutate(dose_zscore_s = ((Dose_mgkg-mean(doses_a_s_mice$Dose_mgkg))/sd(doses_a_s_mice$Dose_mgkg)))

doses_a_i_rats = doses_species %>% filter(`Active Drug`=="Anisomycin" & `Site of injection`!="Systemic" & is.na(Dose_ugside)==FALSE & Species=="rats")
doses_a_i_rats = doses_a_i_rats %>% mutate(dose_zscore_s = ((Dose_ugside-mean(doses_a_i_rats$Dose_ugside))/sd(doses_a_i_rats$Dose_ugside)))

doses_a_i_mice = doses_species %>% filter(`Active Drug`=="Anisomycin" & `Site of injection`!="Systemic" & is.na(Dose_ugside)==FALSE & Species=="mice")
doses_a_i_mice = doses_a_i_mice %>% mutate(dose_zscore_s = ((Dose_ugside-mean(doses_a_i_mice$Dose_ugside))/sd(doses_a_i_mice$Dose_ugside)))

doses_c_s_rats = doses_species %>% filter(`Active Drug`=="Cycloheximide" & `Site of injection`=="Systemic" & is.na(Dose_mgkg)==FALSE & Species=="rats")
doses_c_s_rats = doses_c_s_rats %>% mutate(dose_zscore_s = ((Dose_mgkg-mean(doses_c_s_rats$Dose_mgkg))/sd(doses_c_s_rats$Dose_mgkg)))

doses_c_s_mice = doses_species %>% filter(`Active Drug`=="Cycloheximide" & `Site of injection`=="Systemic" & is.na(Dose_mgkg)==FALSE & Species=="mice")
doses_c_s_mice = doses_c_s_mice %>% mutate(dose_zscore_s = ((Dose_mgkg-mean(doses_c_s_mice$Dose_mgkg))/sd(doses_c_s_mice$Dose_mgkg)))

doses_c_i_rats = doses_species %>% filter(`Active Drug`=="Cycloheximide" & `Site of injection`!="Systemic" & is.na(Dose_ugside)==FALSE & Species=="rats")
doses_c_i_rats = doses_c_i_rats %>% mutate(dose_zscore_s = ((Dose_ugside-mean(doses_c_i_rats$Dose_ugside))/sd(doses_c_i_rats$Dose_ugside)))

doses_c_i_mice = doses_species %>% filter(`Active Drug`=="Cycloheximide" & `Site of injection`!="Systemic" & is.na(Dose_ugside)==FALSE & Species=="mice")
doses_c_i_mice = doses_c_i_mice %>% mutate(dose_zscore_s = 0)

doses_4_i_rats = doses_species %>% filter(`Active Drug`=="4EGI-1" & `Site of injection`!="Systemic" & is.na(Dose_ugside)==FALSE & Species=="rats")
doses_4_i_rats = doses_4_i_rats %>% mutate(dose_zscore_s = 0)

doses_z_species = rbind(doses_a_s_rats, doses_a_s_mice, doses_a_i_rats, doses_a_i_mice, doses_c_s_rats, doses_c_s_mice, doses_c_i_rats, doses_c_i_mice, doses_4_i_rats)
doses_z_species = doses_z_species %>% select(c(Comparison.Code, dose_zscore_s))

full_data = left_join(full_data, doses_z_species, by = "Comparison.Code")
```

```{r, warning=FALSE}
reexposure = full_data %>% select(c(Comparison.Code, ConditioningType, ReexposureDuration_min, ReexposureDuration_number))

reexposure_cont = reexposure %>% filter((ConditioningType=="Contextual"|ConditioningType=="Contextual background") & is.na(ReexposureDuration_min)==FALSE) 
reexposure_cont = reexposure_cont %>% mutate(duration_zscore = ((reexposure_cont$ReexposureDuration_min-mean(reexposure_cont$ReexposureDuration_min))/sd(reexposure_cont$ReexposureDuration_min)))

reexposure_tone = reexposure %>% filter((ConditioningType=="Tone"|ConditioningType=="Tone-Trace") & is.na(ReexposureDuration_number)==FALSE)
reexposure_tone = reexposure_tone %>% mutate(duration_zscore = ((reexposure_tone$ReexposureDuration_number-mean(reexposure_tone$ReexposureDuration_number))/sd(reexposure_tone$ReexposureDuration_number)))

reexposure_z = rbind(reexposure_cont, reexposure_tone)
reexposure_z = reexposure_z %>% select(c(Comparison.Code, duration_zscore))

full_data = left_join(full_data, reexposure_z, by = "Comparison.Code")
```
We added other variables that will be useful for power calculations (mean sample size and pooled standard deviation).
```{r}
full_data = full_data %>% mutate(mean_n = (nControl_original+nTreated_original)/2)
full_data = full_data %>% mutate(pooled_sd = sqrt(((SDControl_calc^2)+(SDTreated_calc^2))/2))
```

Finally, we looked for the number of articles per intervention session and site of injection: (Suppl Table 3)
```{r}
sites_injection = full_data %>% group_by(`Article code`,`Intervention Session`) %>% count(`Site of injection`)
sites_injection %>% group_by(`Site of injection`,`Intervention Session`,) %>% count(`Site of injection`)
```

**Meta-analyses (absolute mean differences)**

- Calculating effect sizes
```{r}
full_data_md = escalc(measure = "MD", m1i = as.numeric(MeanTreated), m2i = as.numeric(MeanControl), sd1i = as.numeric(SDTreated_calc), sd2i = as.numeric(SDControl_calc), n1i = as.numeric(adj_nTreated), n2i = as.numeric(adj_nControl), data = full_data)
```

```{r}
Figs12A = ggplot(full_data, aes(x = `Active Drug`, y = as.numeric(Dose_mgkg), color = `Active Drug`, shape = Species)) + 
  geom_jitter( alpha = 0.5, size = 2 ) + 
  scale_color_manual(values = c("#1b9e77", "#7570b3", "#d95f02")) + 
  theme_classic() + 
  labs(title = "Systemic administration") + ylab("Dose (mg/kg)")  + 
  theme(legend.position = "bottom")

Figs12B = ggplot(full_data, aes(x = `Active Drug`, y = as.numeric(Dose_ugside), color = `Active Drug`, shape = Species)) + 
  geom_jitter( alpha = 0.5, size = 2 ) + 
  scale_color_manual(values = c("#1b9e77", "#7570b3", "#d95f02")) + 
  theme_classic()  + 
  labs(title = "Intracerebral administration") + ylab("Dose (\u03BCg/kg)")   + 
  theme(legend.position = "bottom")

Figs12C = ggplot(full_data_md, aes(x = as.numeric(dose_zscore_s), y = yi, size = 1/vi)) + 
  geom_point(shape = 1, stroke = 0.5) + 
  theme_classic() + 
  theme(legend.position = "none") + 
  labs(title = "Z scores") + ylab("Absolute mean difference") + xlab("Drug dose (z scored)")

FigS12 = plot_grid(Figs12A, Figs12B, Figs12C, labels = LETTERS, nrow = 3, rel_widths = c(1.25, 1.25, 1))
ggsave("Figure S12.png", units = "cm", height = 30)
```

- Breaking datasets
```{r}
training_md = full_data_md %>% filter(Intervention.Session=="Training")
training_syst_md = training_md %>% filter(Site.of.injection=="Systemic")
training_amy_md = training_md %>% filter(Site.of.injection=="Amygdala")
training_hip_md = training_md %>% filter(Site.of.injection=="Hippocampus")
training_icv_md = training_md %>% filter(Site.of.injection=="i.c.v.")
training_pre_md = training_md %>% filter(Site.of.injection=="Prefrontal Cortex")

reconsolidation_md = full_data_md %>% filter(Intervention.Session=="Reconsolidation")
reconsolidation_syst_md = reconsolidation_md %>% filter(Site.of.injection=="Systemic")
reconsolidation_amy_md = reconsolidation_md %>% filter(Site.of.injection=="Amygdala")
reconsolidation_hip_md = reconsolidation_md %>% filter(Site.of.injection=="Hippocampus")
reconsolidation_icv_md = reconsolidation_md %>% filter(Site.of.injection=="i.c.v.")

extinction_md = full_data_md %>% filter(Intervention.Session=="Extinction")
extinction_syst_md = extinction_md %>% filter(Site.of.injection=="Systemic")
extinction_amy_md = extinction_md %>% filter(Site.of.injection=="Amygdala")
extinction_hip_md = extinction_md %>% filter(Site.of.injection=="Hippocampus")
extinction_icv_md = extinction_md %>% filter(Site.of.injection=="i.c.v.")

reactivation_md = full_data_md %>% filter(Intervention.Session!="Training")
reactivation_syst_md = reactivation_md %>% filter(Site.of.injection=="Systemic")
reactivation_amy_md = reactivation_md %>% filter(Site.of.injection=="Amygdala")
reactivation_hip_md = reactivation_md %>% filter(Site.of.injection=="Hippocampus")
reactivation_icv_md = reactivation_md %>% filter(Site.of.injection=="i.c.v.")
```

(Table 2)

Complete dataset:
```{r}
full_data_meta_md = rma(yi=yi, vi=vi, data = full_data_md, measure = "MD", method = "REML", slab = ID)
summary(full_data_meta_md)
confint(full_data_meta_md)
```
Exact model p value: `r full_data_meta_md$pval` | 
Exact Q-test p value: `r full_data_meta_md$QEp`

Training, systemic:
```{r}
training_syst_meta_md = rma(yi=yi, vi=vi, data = training_syst_md, measure = "MD", method = "REML", slab = ID)
summary(training_syst_meta_md)
```
Exact model p value: `r training_syst_meta_md$pval` |
Exact Q-test p value: `r training_syst_meta_md$QEp`

Training, amygdala:
```{r}
training_amy_meta_md = rma(yi=yi, vi=vi, data = training_amy_md, measure = "MD", method = "REML", slab = ID)
summary(training_amy_meta_md)
```
Exact model p value: `r training_amy_meta_md$pval` |
Exact Q-test p value: `r training_amy_meta_md$QEp`

Training, hippocampus:
```{r}
training_hip_meta_md = rma(yi=yi, vi=vi, data = training_hip_md, measure = "MD", method = "REML", slab = ID)
summary(training_hip_meta_md)
```
Exact model p value: `r training_hip_meta_md$pval` |
Exact Q-test p value: `r training_hip_meta_md$QEp`

Training, whole brain:
```{r}
training_icv_meta_md = rma(yi=yi, vi=vi, data = training_icv_md, measure = "MD", method = "REML", slab = ID)
summary(training_icv_meta_md)
```

Training, prefrontal cortex:
```{r}
training_pre_meta_md = rma(yi=yi, vi=vi, data = training_pre_md, measure = "MD", method = "REML", slab = ID)
summary(training_pre_meta_md)
```

All training forest plots (Suppl Fig 2)
```{r}
forest_t_s = viz_forest(training_syst_meta_md, study_labels = training_syst_md$ID, col = "black", annotate_CI = TRUE, xlab = "Effect sizes", text_size = 4)
forest_t_a = viz_forest(training_amy_meta_md, study_labels = training_amy_md$ID, col = "black", annotate_CI = TRUE, xlab = "Effect sizes", text_size = 4)
forest_t_h = viz_forest(training_hip_meta_md, study_labels = training_hip_md$ID, col = "black", annotate_CI = TRUE, xlab = "Effect sizes", text_size = 4)
forest_t_i = viz_forest(training_icv_meta_md, study_labels = training_icv_md$ID, col = "black", annotate_CI = TRUE, xlab = "Effect sizes", text_size = 4)
forest_t_p = viz_forest(training_pre_meta_md, study_labels = training_pre_md$ID, col = "black", annotate_CI = TRUE, xlab = "Effect sizes", text_size = 4)
plot_grid(forest_t_s, forest_t_a, forest_t_h, forest_t_i, forest_t_p, ncol = 3, labels = LETTERS, rel_heights = c(1, 0.3))
ggsave("FIG_S2.png", height = 80, width = 80, units = "cm")
```

Reconsolidation, systemic:
```{r}
reconsolidation_syst_meta_md = rma(yi=yi, vi=vi, data = reconsolidation_syst_md, measure = "MD", method = "REML", slab = ID)
summary(reconsolidation_syst_meta_md)
```
Exact model p value: `r reconsolidation_syst_meta_md$pval` |
Exact Q-test p value: `r reconsolidation_syst_meta_md$QEp`

Reconsolidation, amygdala:
```{r}
reconsolidation_amy_meta_md = rma(yi=yi, vi=vi, data = reconsolidation_amy_md, measure = "MD", method = "REML", slab = ID)
summary(reconsolidation_amy_meta_md)
```
Exact model p value: `r reconsolidation_amy_meta_md$pval` |
Exact Q-test p value: `r reconsolidation_amy_meta_md$QEp`

Reconsolidation, hippocampus:
```{r}
reconsolidation_hip_meta_md = rma(yi=yi, vi=vi, data = reconsolidation_hip_md, measure = "MD", method = "REML", slab = ID)
summary(reconsolidation_hip_meta_md)
```
Exact model p value: `r reconsolidation_hip_meta_md$pval` |
Exact Q-test p value: `r reconsolidation_hip_meta_md$QEp`

Reconsolidation, whole brain:
```{r}
reconsolidation_icv_meta_md = rma(yi=yi, vi=vi, data = reconsolidation_icv_md, measure = "MD", method = "REML", slab = ID)
summary(reconsolidation_icv_meta_md)
```
Exact Q-test p value: `r reconsolidation_icv_meta_md$QEp`

All reconsolidation forest plots (Suppl Fig 3)
```{r}
forest_r_s = viz_forest(reconsolidation_syst_meta_md, study_labels = reconsolidation_syst_md$ID, col = "black", annotate_CI = TRUE, xlab = "Effect sizes", text_size = 4)
forest_r_a = viz_forest(reconsolidation_amy_meta_md, study_labels = reconsolidation_amy_md$ID, col = "black", annotate_CI = TRUE, xlab = "Effect sizes", text_size = 4)
forest_r_h = viz_forest(reconsolidation_hip_meta_md, study_labels = reconsolidation_hip_md$ID, col = "black", annotate_CI = TRUE, xlab = "Effect sizes", text_size = 4)
forest_r_i = viz_forest(reconsolidation_icv_meta_md, study_labels = reconsolidation_icv_md$ID, col = "black", annotate_CI = TRUE, xlab = "Effect sizes", text_size = 4)
plot_grid(forest_r_s, forest_r_a, forest_r_h, forest_r_i, ncol = 3, labels = LETTERS, rel_heights = c(1, 0.3))
ggsave("FIG_S3.png", height = 60, width = 80, units = "cm")
```

Extinction, systemic:
```{r}
extinction_syst_meta_md = rma(yi=yi, vi=vi, data = extinction_syst_md, measure = "MD", method = "REML", slab = ID)
summary(extinction_syst_meta_md)
```
Exact model p value: `r extinction_syst_meta_md$pval` |
Exact Q-test p value: `r extinction_syst_meta_md$QEp`

Extinction, amygdala:
```{r}
extinction_amy_meta_md = rma(yi=yi, vi=vi, data = extinction_amy_md, measure = "MD", method = "REML", slab = ID)
summary(extinction_amy_meta_md)
```

Extinction, hippocampus:
```{r}
extinction_hip_meta_md = rma(yi=yi, vi=vi, data = extinction_hip_md, measure = "MD", method = "REML", slab = ID)
summary(extinction_hip_meta_md)
```

Extinction, whole brain:
```{r}
extinction_icv_meta_md = rma(yi=yi, vi=vi, data = extinction_icv_md, measure = "MD", method = "REML", slab = ID)
summary(extinction_icv_meta_md)
```
Exact model p value: `r extinction_icv_meta_md$pval` |
Exact Q-test p value: `r extinction_icv_meta_md$QEp`

All extinction forest plots (Suppl Fig 4)
```{r}
forest_e_s = viz_forest(extinction_syst_meta_md, study_labels = extinction_syst_md$ID, col = "black", annotate_CI = TRUE, xlab = "Effect sizes", x_limit = c(-5, 45), text_size = 5)
forest_e_a = viz_forest(extinction_amy_meta_md, study_labels = extinction_amy_md$ID, col = "black", annotate_CI = TRUE, xlab = "Effect sizes", text_size = 5)
forest_e_h = viz_forest(extinction_hip_meta_md, study_labels = extinction_hip_md$ID, col = "black", annotate_CI = TRUE, xlab = "Effect sizes", text_size = 5)
forest_e_i = viz_forest(extinction_icv_meta_md, study_labels = extinction_icv_md$ID, col = "black", annotate_CI = TRUE, xlab = "Effect sizes", x_limit = c(-15, 70), text_size = 5)
plot_grid(forest_e_s, forest_e_a, forest_e_h, forest_e_i, ncol = 2, labels = LETTERS)
ggsave("FIG_S4.png", height = 40, width = 60, units = "cm")
```

Reactivation, systemic:
```{r}
reactivation_syst_meta_md = rma(yi=yi, vi=vi, data = reactivation_syst_md, measure = "MD", method = "REML", slab = ID)
summary(reactivation_syst_meta_md)
```
Exact model p value: `r reactivation_syst_meta_md$pval` |
Exact Q-test p value: `r reactivation_syst_meta_md$QEp`

Reactivation, amygdala:
```{r}
reactivation_amy_meta_md = rma(yi=yi, vi=vi, data = reactivation_amy_md, measure = "MD", method = "REML", slab = ID)
summary(reactivation_amy_meta_md)
```
Exact model p value: `r reactivation_amy_meta_md$pval` |
Exact Q-test p value: `r reactivation_amy_meta_md$QEp`

Reactivation, hippocampus:
```{r}
reactivation_hip_meta_md = rma(yi=yi, vi=vi, data = reactivation_hip_md, measure = "MD", method = "REML", slab = ID)
summary(reactivation_hip_meta_md)
```
Exact Q-test p value: `r reactivation_hip_meta_md$QEp`

Reactivation, whole brain:
```{r}
reactivation_icv_meta_md = rma(yi=yi, vi=vi, data = reactivation_icv_md, measure = "MD", method = "REML", slab = ID)
summary(reactivation_icv_meta_md)
```
Exact Q-test p value: `r reactivation_icv_meta_md$QEp`

**Meta-analyses (Hedge's g)**

- Calculating effect sizes
```{r}
full_data = arrange(full_data, full_data$`Year of publication`)
full_data_g = full_data %>% filter(adj_nControl!=1.5)

full_data_g = escalc(measure = "SMD", m1i = as.numeric(MeanTreated), m2i = as.numeric(MeanControl), sd1i = as.numeric(SDTreated_calc), sd2i = as.numeric(SDControl_calc), n1i = as.numeric(adj_nTreated), n2i = as.numeric(adj_nControl), data = full_data_g)
```
- Breaking datasets
```{r}
training_g = full_data_g %>% filter(Intervention.Session=="Training")
training_syst_g = training_g %>% filter(Site.of.injection=="Systemic")
training_amy_g = training_g %>% filter(Site.of.injection=="Amygdala")
training_hip_g = training_g %>% filter(Site.of.injection=="Hippocampus")
training_icv_g = training_g %>% filter(Site.of.injection=="i.c.v.")
training_pre_g = training_g %>% filter(Site.of.injection=="Prefrontal Cortex")

reconsolidation_g = full_data_g %>% filter(Intervention.Session=="Reconsolidation")
reconsolidation_syst_g = reconsolidation_g %>% filter(Site.of.injection=="Systemic")
reconsolidation_amy_g = reconsolidation_g %>% filter(Site.of.injection=="Amygdala")
reconsolidation_hip_g = reconsolidation_g %>% filter(Site.of.injection=="Hippocampus")
reconsolidation_icv_g = reconsolidation_g %>% filter(Site.of.injection=="i.c.v.")

extinction_g = full_data_g %>% filter(Intervention.Session=="Extinction")
extinction_syst_g = extinction_g %>% filter(Site.of.injection=="Systemic")
extinction_amy_g = extinction_g %>% filter(Site.of.injection=="Amygdala")
extinction_hip_g = extinction_g %>% filter(Site.of.injection=="Hippocampus")
extinction_icv_g = extinction_g %>% filter(Site.of.injection=="i.c.v.")

reactivation_g = full_data_g %>% filter(Intervention.Session!="Training")
reactivation_syst_g = reactivation_g %>% filter(Site.of.injection=="Systemic")
reactivation_amy_g = reactivation_g %>% filter(Site.of.injection=="Amygdala")
reactivation_hip_g = reactivation_g %>% filter(Site.of.injection=="Hippocampus")
reactivation_icv_g = reactivation_g %>% filter(Site.of.injection=="i.c.v.")
```

(Suppl. Table 4)

Complete dataset:
```{r}
full_data_meta_g = rma(yi=yi, vi=vi, data = full_data_g, measure = "SMD", method = "REML")
summary(full_data_meta_g)
```
Exact model p value: `r full_data_meta_g$pval` |
Exact Q-test p value: `r full_data_meta_g$QEp`

Training, systemic:
```{r}
training_syst_meta_g = rma(yi=yi, vi=vi, data = training_syst_g, measure = "SMD", method = "REML", slab = training_syst_g$ID)
summary(training_syst_meta_g)
```
Exact model p value: `r training_syst_meta_g$pval` |
Exact Q-test p value: `r training_syst_meta_g$QEp`

Training, amygdala:
```{r}
training_amy_meta_g = rma(yi=yi, vi=vi, data = training_amy_g, measure = "SMD", method = "REML", slab = ID)
summary(training_amy_meta_g)
```
Exact model p value: `r training_amy_meta_g$pval` |
Exact Q-test p value: `r training_amy_meta_g$QEp`

Training, hippocampus:
```{r}
training_hip_meta_g = rma(yi=yi, vi=vi, data = training_hip_g, measure = "SMD", method = "REML", slab = training_hip_g$ID)
summary(training_hip_meta_g)
```
Exact model p value: `r training_hip_meta_g$pval` |
Exact Q-test p value: `r training_hip_meta_g$QEp`

Training, whole brain:
```{r}
training_icv_meta_g = rma(yi=yi, vi=vi, data = training_icv_g, measure = "SMD", method = "REML", slab = ID)
summary(training_icv_meta_g)
```

Training, Prefrontal cortex:
```{r}
training_pre_meta_g = rma(yi=yi, vi=vi, data = training_pre_g, measure = "SMD", method = "REML", slab = ID)
summary(training_pre_meta_g)
```

Reconsolidation, systemic:
```{r}
reconsolidation_syst_meta_g = rma(yi=yi, vi=vi, data = reconsolidation_syst_g, measure = "SMD", method = "REML", slab = reconsolidation_syst_g$ID)
summary(reconsolidation_syst_meta_g)
```
Exact model p value: `r reconsolidation_syst_meta_g$pval` |
Exact Q-test p value: `r reconsolidation_syst_meta_g$QEp`

Reconsolidation, amygdala:
```{r}
reconsolidation_amy_meta_g = rma(yi=yi, vi=vi, data = reconsolidation_amy_g, measure = "SMD", method = "REML", slab = ID)
summary(reconsolidation_amy_meta_g)
```
Exact model p value: `r reconsolidation_amy_meta_g$pval` |
Exact Q-test p value: `r reconsolidation_amy_meta_g$QEp`

Reconsolidation, hippocampus:
```{r}
reconsolidation_hip_meta_g = rma(yi=yi, vi=vi, data = reconsolidation_hip_g, measure = "SMD", method = "REML", slab = reconsolidation_hip_g$ID)
summary(reconsolidation_hip_meta_g)
```
Exact model p value: `r reconsolidation_hip_meta_g$pval` |
Exact Q-test p value: `r reconsolidation_hip_meta_g$QEp`

Reconsolidation, whole brain:
```{r}
reconsolidation_icv_meta_g = rma(yi=yi, vi=vi, data = reconsolidation_icv_g, measure = "SMD", method = "REML", slab = ID)
summary(reconsolidation_icv_meta_g)
```

Extinction, systemic:
```{r}
extinction_syst_meta_g = rma(yi=yi, vi=vi, data = extinction_syst_g, measure = "SMD", method = "REML", slab = ID)
summary(extinction_syst_meta_g)
```
Exact model p value: `r extinction_syst_meta_g$pval` 

Extinction, amygdala:
```{r}
extinction_amy_meta_g = rma(yi=yi, vi=vi, data = extinction_amy_g, measure = "SMD", method = "REML", slab = ID)
summary(extinction_amy_meta_g)
```

Extinction, hippocampus:
```{r}
extinction_hip_meta_g = rma(yi=yi, vi=vi, data = extinction_hip_g, measure = "SMD", method = "REML", slab = ID)
summary(extinction_hip_meta_g)
```

Extinction, whole brain:
```{r}
extinction_icv_meta_g = rma(yi=yi, vi=vi, data = extinction_icv_g, measure = "SMD", method = "REML")
summary(extinction_icv_meta_g)
```
Exact model p value: `r extinction_icv_meta_g$pval` 

Reactivation, systemic:
```{r}
reactivation_syst_meta_g = rma(yi=yi, vi=vi, data = reactivation_syst_g, measure = "SMD", method = "REML", slab = ID)
summary(reactivation_syst_meta_g)
```
Exact model p value: `r reactivation_syst_meta_g$pval` |
Exact Q-test p value: `r reactivation_syst_meta_g$QEp`

Reactivation, amygdala:
```{r}
reactivation_amy_meta_g = rma(yi=yi, vi=vi, data = reactivation_amy_g, measure = "SMD", method = "REML", slab = ID)
summary(reactivation_amy_meta_g)
```
Exact model p value: `r reactivation_amy_meta_g$pval` |
Exact Q-test p value: `r reactivation_amy_meta_g$QEp`

Reactivation, hippocampus:
```{r}
reactivation_hip_meta_g = rma(yi=yi, vi=vi, data = reactivation_hip_g, measure = "SMD", method = "REML", slab = reactivation_hip_g$ID)
summary(reactivation_hip_meta_g)
```
Exact model p value: `r reactivation_hip_meta_g$pval` |
Exact Q-test p value: `r reactivation_hip_meta_g$QEp`

Reactivation, whole brain:
```{r}
reactivation_icv_meta_g = rma(yi=yi, vi=vi, data = reactivation_icv_g, measure = "SMD", method = "REML", slab = ID)
summary(reactivation_icv_meta_g)
```
Exact Q-test p value: `r reactivation_icv_meta_g$QEp`

**Publication Bias (trim-and-fill)**
(Suppl. Table 5)
Training, systemic:
```{r}
training_syst_meta_md_fillR = trimfill(training_syst_meta_md, estimator = "R0", side = "right")
training_syst_meta_md_fillL = trimfill(training_syst_meta_md, estimator = "L0", side = "right")
```
Number of missing studies (R0): `r training_syst_meta_md_fillR$k0` |
p value (R0): `r training_syst_meta_md_fillR$p.k0` |
Number of missing studies (L0): `r training_syst_meta_md_fillL$k0`

Training, amygdala:
```{r}
training_amy_meta_md_fillR = trimfill(training_amy_meta_md, estimator = "R0", side = "right")
training_amy_meta_md_fillL = trimfill(training_amy_meta_md, estimator = "L0", side = "right")
```
Number of missing studies (R0): `r training_amy_meta_md_fillR$k0` |
p value (R0): `r training_amy_meta_md_fillR$p.k0` |
Bias-corrected ES (R0): `r training_amy_meta_md_fillR$beta` | 
Bias-corrected p value (R0): `r training_amy_meta_md_fillR$pval` | 
Number of missing studies (L0): `r training_amy_meta_md_fillL$k0` 

Training, hippocampus:
```{r}
training_hip_meta_md_fillR = trimfill(training_hip_meta_md, estimator = "R0", side = "right")
training_hip_meta_md_fillL = trimfill(training_hip_meta_md, estimator = "L0", side = "right")
```
Number of missing studies (R0): `r training_hip_meta_md_fillR$k0` |
p value (R0): `r training_hip_meta_md_fillR$p.k0` |
Bias-corrected ES (R0): `r training_hip_meta_md_fillR$beta` | 
Bias-corrected p value (R0): `r training_hip_meta_md_fillR$pval` | 
Number of missing studies (L0): `r training_hip_meta_md_fillL$k0` 

Reconsolidation, systemic:
```{r}
reconsolidation_syst_meta_md_fillR = trimfill(reconsolidation_syst_meta_md, estimator = "R0", side = "right")
reconsolidation_syst_meta_md_fillL = trimfill(reconsolidation_syst_meta_md, estimator = "L0", side = "right")
```
Number of missing studies (R0): `r reconsolidation_syst_meta_md_fillR$k0` |
p value (R0): `r reconsolidation_syst_meta_md_fillR$p.k0` |
Number of missing studies (L0): `r reconsolidation_syst_meta_md_fillL$k0` |
Bias-corrected ES (L0): `r reconsolidation_syst_meta_md_fillL$beta` | 
Bias-corrected p value (L0): `r reconsolidation_syst_meta_md_fillL$pval` | 

Reconsolidation, amygdala:
```{r}
reconsolidation_amy_meta_md_fillR = trimfill(reconsolidation_amy_meta_md, estimator = "R0", side = "right")
reconsolidation_amy_meta_md_fillL = trimfill(reconsolidation_amy_meta_md, estimator = "L0", side = "right")
```
Number of missing studies (R0): `r reconsolidation_amy_meta_md_fillR$k0` |
p value (R0): `r reconsolidation_amy_meta_md_fillR$p.k0` |
Number of missing studies (L0): `r reconsolidation_amy_meta_md_fillL$k0` |

Reconsolidation, hippocampus:
```{r}
reconsolidation_hip_meta_md_fillR = trimfill(reconsolidation_hip_meta_md, estimator = "R0", side = "right")
reconsolidation_hip_meta_md_fillL = trimfill(reconsolidation_hip_meta_md, estimator = "L0", side = "right")
```
Number of missing studies (R0): `r reconsolidation_hip_meta_md_fillR$k0` |
p value (R0): `r reconsolidation_hip_meta_md_fillR$p.k0` |
Bias-corrected ES (R0): `r reconsolidation_hip_meta_md_fillR$beta` | 
Bias-corrected p value (R0): `r reconsolidation_hip_meta_md_fillR$pval` | 
Number of missing studies (L0): `r reconsolidation_hip_meta_md_fillL$k0` |

**Publication Bias (Egger's regression)**
(Fig 2 legend)
Training, systemic:
```{r}
training_syst_funnel_md = viz_funnel(x = training_syst_meta_md, trim_and_fill = TRUE, trim_and_fill_side = "right", xlab = "Absolute mean difference", y_axis = "se", ylab = "S.E.", contours = TRUE, sig_contours = FALSE, contours_type = "REM", egger = TRUE) + theme_classic() + 
  labs(title = "Training, systemic")


regtest(training_syst_meta_md)
```

Training, amygdala:
```{r}
training_amy_funnel_md = viz_funnel(x = training_amy_meta_md, trim_and_fill = TRUE, trim_and_fill_side = "right", xlab = "Absolute mean difference", y_axis = "se", ylab = "S.E.", contours = TRUE, sig_contours = FALSE, contours_type = "REM", egger = TRUE) + theme_classic() + 
  labs(title = "Training, amygdala")


regtest(training_amy_meta_md)
```

Training, hippocampus:
```{r}
training_hip_funnel_md = viz_funnel(x = training_hip_meta_md, trim_and_fill = TRUE, trim_and_fill_side = "right", xlab = "Absolute mean difference", y_axis = "se", ylab = "S.E.", contours = TRUE, sig_contours = FALSE, contours_type = "REM", egger = TRUE) + theme_classic() + 
  labs(title = "Training, hippocampus")


regtest(training_hip_meta_md)
```

Reconsolidation, systemic:
```{r}
reconsolidation_syst_funnel_md = viz_funnel(x = reconsolidation_syst_meta_md, trim_and_fill = TRUE, trim_and_fill_side = "right", xlab = "Absolute mean difference", y_axis = "se", ylab = "S.E.", contours = TRUE, sig_contours = FALSE, contours_type = "REM", egger = TRUE) + theme_classic() + 
  labs(title = "Reconsolidation, systemic")


regtest(reconsolidation_syst_meta_md)
```

Reconsolidation, amygdala:
```{r}
reconsolidation_amy_funnel_md = viz_funnel(x = reconsolidation_amy_meta_md, trim_and_fill = TRUE, trim_and_fill_side = "right", xlab = "Absolute mean difference", y_axis = "se", ylab = "S.E.", contours = TRUE, sig_contours = FALSE, contours_type = "REM", egger = TRUE) + theme_classic() + 
  labs(title = "Reconsolidation, amygdala")


regtest(reconsolidation_amy_meta_md)
```

Reconsolidation, hippocampus:
```{r}
reconsolidation_hip_funnel_md = viz_funnel(x = reconsolidation_hip_meta_md, trim_and_fill = TRUE, trim_and_fill_side = "right", xlab = "Absolute mean difference", y_axis = "se", ylab = "S.E.", contours = TRUE, sig_contours = FALSE, contours_type = "REM", egger = TRUE) + theme_classic() + 
  labs(title = "Reconsolidation, hippocampus")

regtest(reconsolidation_hip_meta_md)
```

**Publication bias (funnel plots)**

Fig 2:
```{r}
plot_grid(training_syst_funnel_md, training_amy_funnel_md, training_hip_funnel_md, reconsolidation_syst_funnel_md, reconsolidation_amy_funnel_md, reconsolidation_hip_funnel_md, ncol = 3, labels = LETTERS)

ggsave("FIG_2.png", width = 30, height = 20, units = "cm")
```

**Power analyses**
```{r}
power_t_s = power.t.test(n = training_syst_md$mean_n, sd = training_syst_md$pooled_sd, delta = as.vector(training_syst_meta_md$beta), sig.level = 0.05, type = "two.sample", alternative = "two.sided")

training_syst_md = training_syst_md %>% mutate(power = power_t_s$power)
```
```{r}
power_t_a = power.t.test(n = training_amy_md$mean_n, sd = training_amy_md$pooled_sd, delta = as.vector(training_amy_meta_md$beta), sig.level = 0.05, type = "two.sample", alternative = "two.sided")

training_amy_md = training_amy_md %>% mutate(power = power_t_a$power)
```
```{r}
power_t_h = power.t.test(n = training_hip_md$mean_n, sd = training_hip_md$pooled_sd, delta = as.vector(training_hip_meta_md$beta), sig.level = 0.05, type = "two.sample", alternative = "two.sided")

training_hip_md = training_hip_md %>% mutate(power = power_t_h$power)
```
```{r}
power_r_s = power.t.test(n = reconsolidation_syst_md$mean_n, sd = reconsolidation_syst_md$pooled_sd, delta = as.vector(reconsolidation_syst_meta_md$beta), sig.level = 0.05, type = "two.sample", alternative = "two.sided")

reconsolidation_syst_md = reconsolidation_syst_md %>% mutate(power = power_r_s$power)
```
```{r}
power_r_a = power.t.test(n = reconsolidation_amy_md$mean_n, sd = reconsolidation_amy_md$pooled_sd, delta = as.vector(reconsolidation_amy_meta_md$beta), sig.level = 0.05, type = "two.sample", alternative = "two.sided")

reconsolidation_amy_md = reconsolidation_amy_md %>% mutate(power = power_r_a$power)
```
```{r}
power_r_h = power.t.test(n = reconsolidation_hip_md$mean_n, sd = reconsolidation_hip_md$pooled_sd, delta = as.vector(reconsolidation_hip_meta_md$beta), sig.level = 0.05, type = "two.sample", alternative = "two.sided")

reconsolidation_hip_md = reconsolidation_hip_md %>% mutate(power = power_r_h$power)
```

Training dataset:
```{r}
combine_training = bind_rows(training_syst_md, training_amy_md, training_hip_md)
combine_training = combine_training %>% unite(col = "dataset", c("Route.of.administration","Site.of.injection"), na.rm = TRUE)
combine_training %>% summarise(mean(power))
combine_training %>% summarise(sd(power))
combine_training %>% group_by(dataset) %>% summarise(mean(power))
combine_training %>% group_by(dataset) %>% summarise(sd(power))

combine_training %>% summarise(mean(mean_n))
combine_training %>% summarise(sd(mean_n))
combine_training %>% group_by(dataset) %>% summarise(mean(mean_n))
combine_training %>% group_by(dataset) %>% summarise(sd(mean_n))
```

Reconsolidation dataset:
```{r}
combine_reconsolidation = bind_rows(reconsolidation_syst_md, reconsolidation_amy_md, reconsolidation_hip_md)
combine_reconsolidation = combine_reconsolidation %>% unite(col = "dataset", c("Route.of.administration","Site.of.injection"), na.rm = TRUE)
combine_reconsolidation %>% summarise(mean(power))
combine_reconsolidation %>% summarise(sd(power))
combine_reconsolidation %>% group_by(dataset) %>% summarise(mean(power))
combine_reconsolidation %>% group_by(dataset) %>% summarise(sd(power))

combine_reconsolidation %>% summarise(mean(mean_n))
combine_reconsolidation %>% summarise(sd(mean_n))
combine_reconsolidation %>% group_by(dataset) %>% summarise(mean(mean_n))
combine_reconsolidation %>% group_by(dataset) %>% summarise(sd(mean_n))
```

Combining these datasets is a filtered version of full_data_md, excluding interventions on extinction and some intracerebral sites of drug administration:
```{r}
partial_data_md = bind_rows(combine_training, combine_reconsolidation)
partial_data_md %>% summarise(mean(power))
partial_data_md %>% summarise(sd(power))

partial_data_md %>% summarise(mean(mean_n))
partial_data_md %>% summarise(sd(mean_n))
```

Suppl. Fig. 5:
```{r, warning=FALSE}
power_t = ggplot(combine_training, aes(x = dataset, y = power)) + geom_dotplot(binaxis = "y", stackdir = "center", dotsize = 0.5) + labs(y = "Statistical Power", x = "Site of injection") + theme_classic() + scale_y_continuous(limits = c(0, 1))  + scale_x_discrete(limits = c("systemic_Systemic", "intracerebral_Amygdala", "intracerebral_Hippocampus"), labels = c("Systemic", "Amygdala", "Hippocampus")) + stat_summary(fun = "mean", fun.min = "mean", fun.max = "mean", geom = "crossbar", width = 0.5, size = 0.25)

power_r = ggplot(combine_reconsolidation, aes(x = dataset, y = power)) + geom_dotplot(binaxis = "y", stackdir = "center", dotsize = 0.5) + labs(y = "Statistical Power", x = "Site of injection") + theme_classic() + scale_y_continuous(limits = c(0, 1))  + scale_x_discrete(limits = c("systemic_Systemic", "intracerebral_Amygdala", "intracerebral_Hippocampus"), labels = c("Systemic", "Amygdala", "Hippocampus")) + stat_summary(fun = "mean", fun.min = "mean", fun.max = "mean", geom = "crossbar", width = 0.5, size = 0.25)

samplesize_t = ggplot(combine_training, aes(x = dataset, y = mean_n)) + geom_dotplot(binaxis = "y", stackdir = "center", dotsize = 0.5) + labs(y = "Mean sample size", x = "Site of injection", title = "Training") + theme_classic() + scale_x_discrete(limits = c("systemic_Systemic", "intracerebral_Amygdala", "intracerebral_Hippocampus"), labels = c("Systemic", "Amygdala", "Hippocampus")) + stat_summary(fun = "mean", fun.min = "mean", fun.max = "mean", geom = "crossbar", width = 0.5, size = 0.25) + scale_y_continuous(limits = c(0,30)) + theme(plot.title = element_text(hjust = 0.5))

samplesize_r = ggplot(combine_reconsolidation, aes(x = dataset, y = mean_n)) + geom_dotplot(binaxis = "y", stackdir = "center", dotsize = 0.5) + labs(y = "Mean sample size", x = "Site of injection", title = "Reconsolidation") + theme_classic() + scale_x_discrete(limits = c("systemic_Systemic", "intracerebral_Amygdala", "intracerebral_Hippocampus"), labels = c("Systemic", "Amygdala", "Hippocampus")) + stat_summary(fun = "mean", fun.min = "mean", fun.max = "mean", geom = "crossbar", width = 0.5, size = 0.25) + scale_y_continuous(limits = c(0,30)) + theme(plot.title = element_text(hjust = 0.5))

plot_grid(samplesize_t, samplesize_r, power_t, power_r, ncol = 2, labels = LETTERS)
ggsave("FIG_S5.png", width = 20, height = 20, units = "cm")
```

**Additional correlations**

For correlations with article-level variables, we need to get mean effect size and power for each article (that includes at least one experiment with systemic, intra-amygdala or intra-hippocampus interventions on training or reconsolidation):
```{r}
partial_data_md$ConflictInterest[partial_data_md$ConflictInterest=="Not informed"] = FALSE
partial_data_md$ConflictInterest[partial_data_md$ConflictInterest=="No"] = TRUE
partial_data_md$ConflictInterest[partial_data_md$ConflictInterest=="Yes"] = TRUE

partial_data_md$ConflictInterest = as.logical(partial_data_md$ConflictInterest)

partial_data_md_article = partial_data_md %>% group_by(Article.code) %>% summarise(ES=mean(yi), Power=mean(power), IF=mean(Impact.Factor), Cit=mean(Citations.per.year), Rand=mean(Randomization), Blind = mean(Blinding), SSC=mean(SampleSizeCalc), Reg=mean(RegulatoryReq), COI=mean(ConflictInterest))

partial_data_md_article = partial_data_md_article %>% mutate(Bias = (5-(Rand+Blind+SSC+Reg+COI)))
```

Effect size vs. Impact factor
```{r}
es_if = ggplot(partial_data_md_article, aes(x = IF, y = ES)) + geom_point() + theme_classic() + geom_hline(aes(yintercept=0), color="grey", linetype="dotted") + scale_y_continuous(name = "Absolute mean difference") + scale_x_continuous(name = "Impact Factor")
es_if_cor = cor.test(x = partial_data_md_article$IF, y = partial_data_md_article$ES, method = "spearman", alternative = "two.sided", exact = F)
es_if_cor
partial_data_md_article %>% count(!is.na(IF))
```

Effect size vs. Citations
```{r}
es_cit = ggplot(partial_data_md_article, aes(x = Cit, y = ES)) + geom_point() + theme_classic() + geom_hline(aes(yintercept=0), color="grey", linetype="dotted") + scale_y_continuous(name = "Absolute mean difference") + scale_x_continuous(name = "Citations per year")
es_cit_cor = cor.test(x = partial_data_md_article$Cit, y = partial_data_md_article$ES, method = "spearman", alternative = "two.sided", exact = F)
es_cit_cor
partial_data_md_article %>% count(!is.na(Cit))
```

Effect size vs. Risk of Bias
```{r}
es_bias = ggplot(partial_data_md_article, aes(x = Bias, y = ES)) + geom_point() + theme_classic() + geom_hline(aes(yintercept=0), color="grey", linetype="dotted") + scale_y_continuous(name = "Absolute mean difference") + scale_x_continuous(name = "Risk of Bias score")
es_bias_cor = cor.test(x = partial_data_md_article$Bias, y = partial_data_md_article$ES, method = "spearman", alternative = "two.sided", exact = F)
es_bias_cor
partial_data_md_article %>% count(!is.na(Bias))
```

Power vs. Impact factor
```{r}
power_if = ggplot(partial_data_md_article, aes(x = IF, y = Power)) + geom_point() + theme_classic() + scale_y_continuous(name = "Statistical Power") + scale_x_continuous(name = "Impact Factor")
power_if_cor = cor.test(x = partial_data_md_article$IF, y = partial_data_md_article$Power, method = "spearman", alternative = "two.sided", exact = F)
power_if_cor
```

Power vs. Citations
```{r}
power_cit = ggplot(partial_data_md_article, aes(x = Cit, y = Power)) + geom_point() + theme_classic() + scale_y_continuous(name = "Statistical Power") + scale_x_continuous(name = "Citations per year")
power_cit_cor = cor.test(x = partial_data_md_article$Cit, y = partial_data_md_article$Power, method = "spearman", alternative = "two.sided", exact = F)
power_cit_cor
```

Power vs. Risk of Bias
```{r}
power_bias = ggplot(partial_data_md_article, aes(x = Bias, y = Power)) + geom_point() + theme_classic() + scale_y_continuous(name = "Statistical Power") + scale_x_continuous(name = "Risk of Bias score")
power_bias_cor = cor.test(x = partial_data_md_article$Bias, y = partial_data_md_article$Power, method = "spearman", alternative = "two.sided", exact = F)
power_bias_cor
```

Suppl. Fig. 6
```{r, warning=FALSE}
plot_grid(es_if, es_cit, es_bias, power_if, power_cit, power_bias, ncol = 3, labels = LETTERS)
ggsave("FIG_S6.png", width = 30, height = 20, units = "cm")
```

**Excess significance**
(Suppl Table 6)
For these analyses, we considered the original (unadjusted) sample sizes, so we had to:

- Recalculate effect sizes
```{r}
full_data_md_n = escalc(measure = "MD", m1i = as.numeric(MeanTreated), m2i = as.numeric(MeanControl), sd1i = as.numeric(SDTreated_calc), sd2i = as.numeric(SDControl_calc), n1i = as.numeric(nTreated_original), n2i = as.numeric(nControl_original), data = full_data)
```
- Break the datasets with the new effect sizes
```{r}
training_md_n = full_data_md_n %>% filter(Intervention.Session=="Training")
training_syst_md_n = training_md_n %>% filter(Site.of.injection=="Systemic")
training_amy_md_n = training_md_n %>% filter(Site.of.injection=="Amygdala")
training_hip_md_n = training_md_n %>% filter(Site.of.injection=="Hippocampus")

reconsolidation_md_n = full_data_md_n %>% filter(Intervention.Session=="Reconsolidation")
reconsolidation_syst_md_n = reconsolidation_md_n %>% filter(Site.of.injection=="Systemic")
reconsolidation_amy_md_n = reconsolidation_md_n %>% filter(Site.of.injection=="Amygdala")
reconsolidation_hip_md_n = reconsolidation_md_n %>% filter(Site.of.injection=="Hippocampus")
```

Training, systemic:
```{r}
training_syst_meta_md_n = rma(yi=yi, vi=vi, data = training_syst_md_n, measure = "MD", method = "REML", slab = ID)
excess_t_s_md_n = tes(training_syst_meta_md_n, test = "chi2", tes.alternative = "two.sided")
```
Mean power: `r mean(excess_t_s_md_n$power)` | 
SD power: `r sd(excess_t_s_md_n$power)` | 
Expected: `r excess_t_s_md_n$E` | 
Observed: `r excess_t_s_md_n$O` | 
p value: `r excess_t_s_md_n$pval`

Training, amygdala:
```{r}
training_amy_meta_md_n = rma(yi=yi, vi=vi, data = training_amy_md_n, measure = "MD", method = "REML", slab = ID)
excess_t_a_md_n = tes(training_amy_meta_md_n, test = "chi2", tes.alternative = "two.sided")
```
Mean power: `r mean(excess_t_a_md_n$power)` | 
SD power: `r sd(excess_t_a_md_n$power)` | 
Expected: `r excess_t_a_md_n$E` | 
Observed: `r excess_t_a_md_n$O` | 
p value: `r excess_t_a_md_n$pval`

Training, hippocampus 
```{r}
training_hip_meta_md_n = rma(yi=yi, vi=vi, data = training_hip_md_n, measure = "MD", method = "REML", slab = ID)
excess_t_h_md_n = tes(training_hip_meta_md_n, test = "chi2", tes.alternative = "two.sided")
```
Mean power: `r mean(excess_t_h_md_n$power)` | 
SD power: `r sd(excess_t_h_md_n$power)` | 
Expected: `r excess_t_h_md_n$E` | 
Observed: `r excess_t_h_md_n$O` | 
p value: `r excess_t_h_md_n$pval`

Reconsolidation, systemic:
```{r}
reconsolidation_syst_meta_md_n = rma(yi=yi, vi=vi, data = reconsolidation_syst_md_n, measure = "MD", method = "REML", slab = ID)
excess_r_s_md_n = tes(reconsolidation_syst_meta_md_n, test = "chi2", tes.alternative = "two.sided")
```
Mean power: `r mean(excess_r_s_md_n$power)` | 
SD power: `r sd(excess_r_s_md_n$power)` | 
Expected: `r excess_r_s_md_n$E` | 
Observed: `r excess_r_s_md_n$O` | 
p value: `r excess_r_s_md_n$pval`

Reconsolidation, amygdala:
```{r}
reconsolidation_amy_meta_md_n = rma(yi=yi, vi=vi, data = reconsolidation_amy_md_n, measure = "MD", method = "REML", slab = ID)
excess_r_a_md_n = tes(reconsolidation_amy_meta_md_n, test = "chi2", tes.alternative = "two.sided")
```
Mean power: `r mean(excess_r_a_md_n$power)` | 
SD power: `r sd(excess_r_a_md_n$power)` | 
Expected: `r excess_r_a_md_n$E` | 
Observed: `r excess_r_a_md_n$O` | 
p value: `r excess_r_a_md_n$pval`

Reconsolidation, hippocampus: 
```{r}
reconsolidation_hip_meta_md_n = rma(yi=yi, vi=vi, data = reconsolidation_hip_md_n, measure = "MD", method = "REML", slab = ID)

excess_r_h_md_n = tes(reconsolidation_hip_meta_md_n, test = "chi2", tes.alternative = "two.sided")
```
Mean power: `r mean(excess_r_h_md_n$power)` | 
SD power: `r sd(excess_r_h_md_n$power)` | 
Expected: `r excess_r_h_md_n$E` | 
Observed: `r excess_r_h_md_n$O` | 
p value: `r excess_r_h_md_n$pval`

**Meta-regressions - intervention site**
(Table 3 + Suppl Table 7)
Training dataset, no moderators:
```{r}
training_md_meta = rma(yi=yi, vi=vi, data = training_md, measure = "MD", method = "REML")
summary(training_md_meta)
confint(training_md_meta)
```
Exact model p value: `r training_md_meta$pval` | 
Exact Q test p value: `r training_md_meta$QEp`

Training dataset, intervention site:
```{r}
training_md %>% group_by(Site.injection.other) %>% summarise(n())

training_md_sites = rma(yi=yi, vi=vi, data = training_md, measure = "MD", method = "REML", mods = ~relevel(factor(training_md$Site.injection.other), ref = "Systemic"))
summary(training_md_sites)
```
Exact model p values: `r training_md_sites$pval`

Reconsolidation dataset, no moderators:
```{r}
reconsolidation_md_meta = rma(yi=yi, vi=vi, data = reconsolidation_md, measure = "MD", method = "REML")
summary(reconsolidation_md_meta)
```
Exact model p value: `r reconsolidation_md_meta$pval` | 
Exact Q test p value: `r reconsolidation_md_meta$QEp`

Reconsolidation dataset, intervention site:
```{r}
reconsolidation_md %>% group_by(Site.injection.other) %>% summarise(n())

reconsolidation_md_sites = rma(yi=yi, vi=vi, data = reconsolidation_md, measure = "MD", method = "REML", mods = ~relevel(factor(reconsolidation_md$Site.injection.other), ref = "Systemic"))
summary(reconsolidation_md_sites)
```
Exact model p values: `r reconsolidation_md_sites$pval` | 
Exact Q test of moderators p value: `r reconsolidation_md_sites$QMp`

Extinction dataset, no moderators:
```{r}
extinction_md_meta = rma(yi=yi, vi=vi, data = extinction_md, measure = "MD", method = "REML")
summary(extinction_md_meta)
```
Exact Q test p value: `r extinction_md_meta$QEp`

Extinction dataset, intervention site:
```{r}
extinction_md %>% group_by(Site.injection.other) %>% summarise(n())

extinction_md_sites = rma(yi=yi, vi=vi, data = extinction_md, measure = "MD", method = "REML", mods = ~relevel(factor(extinction_md$Site.injection.other), ref = "Systemic"))
summary(extinction_md_sites)
```
Exact model p values: `r extinction_md_sites$pval`

**Meta-regressions - intervention session**
(Table 3 + Suppl Table 8)
As we are now grouping by injection site, we had to break the datasets again:
```{r}
systemic_md = full_data_md %>% filter(Site.injection.other=="Systemic")
amygdala_md = full_data_md %>% filter(Site.injection.other=="Amygdala")
hippocampus_md = full_data_md %>% filter(Site.injection.other=="Hippocampus")
icv_md = full_data_md %>% filter(Site.injection.other=="i.c.v.")
other_md = full_data_md %>% filter(Site.injection.other=="Other")
```

Systemic dataset, no moderators:
```{r}
systemic_md_meta = rma(yi=yi, vi=vi, data = systemic_md, measure = "MD", method = "REML")
summary(systemic_md_meta)
```
Exact model p value: `r systemic_md_meta$pval` | 
Exact Q test p value: `r systemic_md_meta$QEp`

Systemic dataset, intervention session:
```{r}
systemic_md %>% group_by(Intervention.Session) %>% summarise(n())

systemic_md_session = rma(yi=yi, vi=vi, data = systemic_md, measure = "MD", method = "REML", mods = ~relevel(factor(systemic_md$Intervention.Session), ref = "Training"))
summary(systemic_md_session)
```
Exact model p values: `r systemic_md_session$pval` | 
Exact Q test of moderators p value: `r systemic_md_session$QMp`

Amygdala dataset, no moderators:
```{r}
amygdala_md_meta = rma(yi=yi, vi=vi, data = amygdala_md, measure = "MD", method = "REML")
summary(amygdala_md_meta)
```
Exact model p value: `r amygdala_md_meta$pval` | 
Exact Q test p value: `r amygdala_md_meta$QEp`

Amygdala dataset, intervention session:
```{r}
amygdala_md %>% group_by(Intervention.Session) %>% summarise(n())

amygdala_md_session = rma(yi=yi, vi=vi, data = amygdala_md, measure = "MD", method = "REML", mods = ~relevel(factor(amygdala_md$Intervention.Session), ref = "Training"))
summary(amygdala_md_session)
```
Exact model p values: `r amygdala_md_session$pval` | 
Exact Q test of moderators p value: `r amygdala_md_session$QMp`

Hippocampus dataset, no moderators:
```{r}
hippocampus_md_meta = rma(yi=yi, vi=vi, data = hippocampus_md, measure = "MD", method = "REML")
summary(hippocampus_md_meta)
```
Exact model p value: `r hippocampus_md_meta$pval` | 
Exact Q test p value: `r hippocampus_md_meta$QEp`

Hippocampus dataset, intervention session:
```{r}
hippocampus_md %>% group_by(Intervention.Session) %>% summarise(n())

hippocampus_md_session = rma(yi=yi, vi=vi, data = hippocampus_md, measure = "MD", method = "REML", mods = ~relevel(factor(hippocampus_md$Intervention.Session), ref = "Training"))
summary(hippocampus_md_session)
```
Exact model p values: `r hippocampus_md_session$pval` | 
Exact Q test of moderators p value: `r hippocampus_md_session$QMp`

i.c.v. dataset, no moderators:
```{r}
icv_md_meta = rma(yi=yi, vi=vi, data = icv_md, measure = "MD", method = "REML")
summary(icv_md_meta)
```
Exact model p value: `r icv_md_meta$pval` | 
Exact Q test p value: `r icv_md_meta$QEp`

i.c.v. dataset, intervention session:
```{r}
icv_md %>% group_by(Intervention.Session) %>% summarise(n())

icv_md_session = rma(yi=yi, vi=vi, data = icv_md, measure = "MD", method = "REML", mods = ~relevel(factor(icv_md$Intervention.Session), ref = "Training"))
summary(icv_md_session)
```
Exact model p values: `r icv_md_session$pval` | 
Exact Q test of moderators p value: `r icv_md_session$QMp`

Other dataset, no moderators:
```{r}
other_md_meta = rma(yi=yi, vi=vi, data = other_md, measure = "MD", method = "REML")
summary(other_md_meta)
```
Exact model p value: `r other_md_meta$pval` | 
Exact Q test p value: `r other_md_meta$QEp`

Other dataset, intervention session:
```{r}
other_md %>% group_by(Intervention.Session) %>% summarise(n())

other_md_session = rma(yi=yi, vi=vi, data = other_md, measure = "MD", method = "REML", mods = ~relevel(factor(other_md$Intervention.Session), ref = "Training"))
summary(other_md_session)
```
Exact model p values: `r other_md_session$pval` | 
Exact Q test of moderators p value: `r other_md_session$QMp`

**Meta-regression - site+session**
(Table 3 + Suppl Table 9)
Complete dataset, multivariate model with injection site and intervention session as moderators:
```{r}
full_data_md %>% group_by(Site.injection.other) %>% summarise(n())

full_data_md_session_site = rma(yi=yi, vi=vi, data = full_data_md, measure = "MD", method = "ML", mods = ~relevel(factor(Intervention.Session), ref = "Training")+relevel(factor(Site.injection.other), ref = "Systemic"))
summary(full_data_md_session_site)
```
Exact model p value: `r full_data_md_session_site$pval` | 
Exact Q test of moderators p value: `r full_data_md_session_site$QMp`

Functions to decompose R2 (2-level)
```{r}
# Gera a lista de combinaÃ§Ãµes de preditores, a partir da lista de preditores e um de interesse, pra qual vocÃª quer calcular o R2 - retorna o texto das fÃ³rmulas, e.g. "~ a + b"
build_model_combinations = function (todos_preditores, preditor_interesse){
  outros_preditores = todos_preditores[todos_preditores != preditor_interesse]
  
  reduced_models = c("", map(1:length(outros_preditores), function (i) {
    # Aqui Ã© onde as combinaÃ§Ãµes sÃ£o montadas de fato
    paste0("~ ", apply(combn(outros_preditores, i), MARGIN = 2, paste0, collapse = " + "))
  }) |> unlist())
  
  # full model Ã© sÃ³ adicionar a variÃ¡vel de interesse
  full_models = map_chr(reduced_models, function (f) {
    if (f == "") {
      return (paste0("~ ", preditor_interesse) )
    } else {
      return (paste0(f, " + ", preditor_interesse) )
    }
  })
  
  list(full_models = full_models, reduced_models = reduced_models)
}

# Roda todos os modelos, par a par, full x reduced, a partir das fÃ³rmulas, e faz a diferença de R2
all_model_comparisons = function (comparison_list) {
  map2_dfr(comparison_list$reduced_models, comparison_list$full_models, function (m0s, m1s) {
    if (m0s == "") {
      # Se nÃ£o tem preditor, Ã© o modelo sem moderadores
      m0 = rma(yi, vi, data=dat, method="ML", control = list(maxiter = 3000))
    } else {
      # Se tem preditores, usa a fÃ³rmula que vem da outra funÃ§Ã£o
      m0f = as.formula(m0s)
      m0 = rma(yi, vi, mods = m0f, data=dat, method="ML", control = list(maxiter = 3000))
    }
    
    m1f = as.formula(m1s)
    m1 = rma(yi, vi, mods = m1f, data=dat, method="ML", control = list(maxiter = 3000))
    
    reduced_model_size = length(all.vars(m1f)) - 1
    
    tibble(reduced = m0s, full = m1s, R2 = (m1$R2-m0$R2), permutations = factorial(reduced_model_size))
  })
}

# Faz todo o processo pra um Ãºnico preditor e resume na mÃ©dia dos R2
get_R2_for_single_predictor = function (todos_preditores, preditor_interesse) {
  comparison_list = build_model_combinations(todos_preditores, preditor_interesse)
  
  r = all_model_comparisons(comparison_list)
  s = mean(r$R2, na.rm = T)
  # Pra essa, o peso Ã© o nÃºmero de permutaÃ§Ãµes do modelo reduzido (se a variÃ¡vel de interesse Ã© 4a, tem 3! maneiras possÃ­veis dela ser a 4a, porque tem 3 variÃ¡veis antes dela pra permutar ... faz sentido?)
  ws = weighted.mean(r$R2, r$permutations, na.rm = T)
  
  list(all_predictors = todos_preditores, predictor = preditor_interesse, results = r, mean = s, wmean = ws)
}

# Faz todo o processo em separado pra cada um dos preditores da lista
get_R2_for_all_predictors = function (todos_preditores) {
  df = map(todos_preditores, function (preditor_interesse) {
    print(paste("Running models for", preditor_interesse))
    get_R2_for_single_predictor(todos_preditores, preditor_interesse)
  })
  s = tibble(mean_R2 = map_dbl(df, "mean"), weighted_mean_R2 = map_dbl(df, "wmean"), predictor = map_chr(df, "predictor"))
  list(full_results = df, summary = s)
}
```

Decomposing R2 for the model with complete dataset and site+session as moderators: (Table 3)
```{r}
dat = full_data_md
meus_preditores = c("relevel(factor(Intervention.Session), ref = 'Training')", "relevel(factor(Site.injection.other), ref = 'Systemic')")
R2_full_data_session_site = get_R2_for_all_predictors(meus_preditores)
R2_full_data_session_site$summary
#p value intervention session
anova(full_data_md_session_site, btt = 2:4)$QMp
#p value injection site
anova(full_data_md_session_site, btt = 5:8)$QMp
```

**Multi-level model - research groups**

We found `r nrow(full_data_md %>% count(group))` groups, with modularity value of `r `modularity(finding_groups)`.

Suppl. Fig. 7 - Groups identified
```{r}
colrs = rainbow(n = nrow(full_data_md %>% count(group)))

V(authors_graph)$color = colrs[V(authors_graph)$community]

plot(authors_graph, vertex.size = 3, vertex.label.cex = 2, vertex.label.color = "black", vertex.label.dist = 0, vertex.frame.color = element_blank(), layout = layout_with_graphopt(authors_graph, charge = 0.01))
```
ps: the position and colors of the groups in the graph change every time the code is run. for the figure included in the manuscript (with good font proportion), it was manually (in RStudio) exported as a figure with width = 5000, height = 3307.

Visualization of the distribution of experiments per article and per research group:
```{r}
dist_articles = ggplot(data = full_data %>% count(ID), aes(x = n)) + 
  geom_histogram(binwidth = 1, fill = "black", color = "white") + 
  theme_classic() + 
  scale_x_continuous(name = "Number of experiments", n.breaks = 10) + 
  scale_y_continuous(name = "Number of articles", n.breaks = 15) +
  theme(axis.text = element_text(color = "black", size = 8), axis.title = element_text(color = "black", size = 12), plot.margin = margin(0.5,0,0,0.5, "cm"))
```
```{r}
dist_group = ggplot(data = full_data %>% count(group), aes(x = n)) + 
  geom_histogram(binwidth = 1, fill = "black", color = "white") + 
  theme_classic() + 
  scale_x_continuous(name = "Number of experiments", n.breaks = 10) + 
  scale_y_continuous(name = "Number of groups", n.breaks = 15) +
  theme(axis.text = element_text(color = "black", size = 8), axis.title = element_text(color = "black", size = 12), plot.margin = margin(0.5,0,0,0.5, "cm"))
```

Suppl. Fig. 8
```{r}
plot_grid(dist_articles, dist_group, labels = LETTERS)
ggsave("FIG_S8.png", width = 20, height = 10, units = "cm")
```
```{r}
n_exp_per_group = full_data %>% count(group)
n_exp_per_group %>% count(n) %>% arrange(-n)
```

(Table 4 - 3-level (group))

Complete dataset, random within and random between research groups:
```{r}
full_data_meta_group_md = rma.mv(yi=yi, V=vi, data = full_data_md, method = "REML", random = ~1|group/Comparison.Code)
summary(full_data_meta_group_md)
```
Exact model p value: `r full_data_meta_group_md$pval` |
95% C.I. for sigma values: 
```{r}
confint(full_data_meta_group_md)
```

Training dataset, random within and random between research groups
```{r}
training_meta_group_md = rma.mv(yi=yi, V=vi, data = training_md, method = "REML", random = ~1|group/Comparison.Code)
summary(training_meta_group_md)
```
Exact model p value: `r training_meta_group_md$pval` |
95% C.I. for sigma values: 
```{r} 
confint(training_meta_group_md, digits = 7)
```

Reactivation dataset, random-effects model:
```{r}
reactivation_md_meta = rma(yi=yi, vi=vi, data = reactivation_md, measure = "MD", method = "REML")
summary(reactivation_md_meta)
confint(reactivation_md_meta)
```
Exact model p value: `r reactivation_md_meta$pval` 

Reactivation dataset, random within and random between research groups
```{r}
reactivation_meta_group_md = rma.mv(yi=yi, V=vi, data = reactivation_md, method = "REML", random = ~1|group/Comparison.Code)
summary(reactivation_meta_group_md)
```
Exact model p value: `r reactivation_meta_group_md$pval` |
95% C.I. for sigma values: 
```{r} 
confint(reactivation_meta_group_md)
```

**Multi-level model - article**
(Table 4 - 3-level (article))

Complete dataset, random within and random between articles:
```{r}
full_data_meta_article_md = rma.mv(yi=yi, V=vi, data = full_data_md, method = "REML", random = ~1|ID/Comparison.Code)
summary(full_data_meta_article_md)
```
Exact model p value: `r full_data_meta_article_md$pval` |
95% C.I. for sigma values: 
```{r}
confint(full_data_meta_article_md)
```

Training dataset, random within and random between articles
```{r}
training_meta_article_md = rma.mv(yi=yi, V=vi, data = training_md, method = "REML", random = ~1|ID/Comparison.Code)
summary(training_meta_article_md)
```
Exact model p value: `r training_meta_article_md$pval` |
95% C.I. for sigma values: 
```{r}
confint(training_meta_article_md)
```

Reactivation dataset, random within and random between articles
```{r}
reactivation_meta_article_md = rma.mv(yi=yi, V=vi, data = reactivation_md, method = "REML", random = ~1|ID/Comparison.Code)
summary(reactivation_meta_article_md)
```
Exact model p value: `r reactivation_meta_article_md$pval` |
95% C.I. for sigma values: 
```{r}
confint(reactivation_meta_article_md)
```

**Distribution of total variance**

As there is not function in the package metafor for estimating I^2 values for 3-level models, we imported the function from Mathias Harrer & David Daniel Ebert (https://raw.githubusercontent.com/MathiasHarrer/dmetar/master/R/mlm.variance.distribution.R)
```{r}
mlm.variance.distribution = var.comp = function(x){

  m = x

  # Check class
  if (!(class(m)[1] %in% c("rma.mv", "rma"))){
    stop("x must be of class 'rma.mv'.")
  }

  # Check for three level model
  if (m$sigma2s != 2){
    stop("The model you provided does not seem to be a three-level model. This function can only be used for three-level models.")
  }

  # Check for right specification (nested model)
  if (sum(grepl("/", as.character(m$random[[1]]))) < 1){
    stop("Model must contain nested random effects. Did you use the '~ 1 | cluster/effect-within-cluster' notation in 'random'? See ?metafor::rma.mv for more details.")
  }

  # Get variance diagonal and calculate total variance
  n = m$k.eff
  vector.inv.var = 1/(diag(m$V))
  sum.inv.var = sum(vector.inv.var)
  sum.sq.inv.var = (sum.inv.var)^2
  vector.inv.var.sq = 1/(diag(m$V)^2)
  sum.inv.var.sq = sum(vector.inv.var.sq)
  num = (n-1)*sum.inv.var
  den = sum.sq.inv.var - sum.inv.var.sq
  est.samp.var = num/den

  # Calculate variance proportions
  level1=((est.samp.var)/(m$sigma2[1]+m$sigma2[2]+est.samp.var)*100)
  level2=((m$sigma2[2])/(m$sigma2[1]+m$sigma2[2]+est.samp.var)*100)
  level3=((m$sigma2[1])/(m$sigma2[1]+m$sigma2[2]+est.samp.var)*100)

  # Prepare df for return
  Level=c("Level 1", "Level 2", "Level 3")
  Variance=c(level1, level2, level3)
  df.res=data.frame(Variance)
  colnames(df.res) = c("% of total variance")
  rownames(df.res) = Level
  I2 = c("---", round(Variance[2:3], 2))
  df.res = as.data.frame(cbind(df.res, I2))

  totalI2 = Variance[2] + Variance[3]


  # Generate plot
  df1 = data.frame("Level" = c("Sampling Error", "Total Heterogeneity"),
                  "Variance" = c(df.res[1,1], df.res[2,1]+df.res[3,1]),
                  "Type" = rep(1,2))

  df2 = data.frame("Level" = rownames(df.res),
                   "Variance" = df.res[,1],
                   "Type" = rep(2,3))

  df = as.data.frame(rbind(df1, df2))


  g = ggplot(df, aes(fill=Level, y=Variance, x=as.factor(Type))) +
    coord_cartesian(ylim = c(0,1), clip = "off") +
    geom_bar(stat="identity", position="fill", width = 1, color="black") +
    scale_y_continuous(labels = scales::percent)+
    theme(axis.title.x=element_blank(),
          axis.text.y = element_text(color="black"),
          axis.line.y = element_blank(),
          axis.title.y=element_blank(),
          axis.line.x = element_blank(),
          axis.ticks.x = element_blank(),
          axis.text.x = element_blank(),
          axis.ticks.y = element_line(lineend = "round"),
          legend.position = "none",
          panel.grid.major = element_blank(),
          panel.grid.minor = element_blank(),
          panel.background = element_blank(),
          legend.background = element_rect(linetype="solid",
                                           colour ="black"),
          legend.title = element_blank(),
          legend.key.size = unit(0.75,"cm"),
          axis.ticks.length=unit(.25, "cm"),
          plot.margin = unit(c(1,3,1,1), "lines")) +
    scale_fill_manual(values = c("darkseagreen3", "deepskyblue3", "darkseagreen2",
                                 "deepskyblue1", "deepskyblue2")) +

    # Add Annotation

    # Total Variance
    annotate("text", x = 1.5, y = 1.05,
             label = paste("Total Variance:",
                           round(m$sigma2[1]+m$sigma2[2]+est.samp.var, 3))) +

    # Sampling Error
    annotate("text", x = 1, y = (df[1,2]/2+df[2,2])/100,
             label = paste("Sampling Error Variance: \n", round(est.samp.var, 3)), size = 3) +

    # Total I2
    annotate("text", x = 1, y = ((df[2,2])/100)/2-0.02,
             label = bquote("Total"~italic(I)^2*":"~.(round(df[2,2],2))*"%"), size = 3) +
    annotate("text", x = 1, y = ((df[2,2])/100)/2+0.05,
             label = paste("Variance not attributable \n to sampling error: \n", round(m$sigma2[1]+m$sigma2[2],3)), size = 3) +

    # Level 1
    annotate("text", x = 2, y = (df[1,2]/2+df[2,2])/100, label = paste("Level 1: \n",
                                                                       round(df$Variance[3],2), "%", sep=""), size = 3) +

    # Level 2
    annotate("text", x = 2, y = (df[5,2]+(df[4,2]/2))/100,
             label = bquote(italic(I)[Level2]^2*":"~.(round(df[4,2],2))*"%"), size = 3) +

    # Level 3
    annotate("text", x = 2, y = (df[5,2]/2)/100,
             label = bquote(italic(I)[Level3]^2*":"~.(round(df[5,2],2))*"%"), size = 3)

  returnlist = list(results = df.res,
                    totalI2 = totalI2,
                    plot = g)
  class(returnlist) = c("mlm.variance.distribution", "list")

  invisible(returnlist)

  returnlist

}
```

(Table 4 - I2)

Then, we can get the values for models with research groups as clusters:
Complete dataset
```{r fig.keep='none', warning=FALSE}
mlm.variance.distribution(x = full_data_meta_group_md)
```
Training dataset
```{r fig.keep='none', warning=FALSE}
mlm.variance.distribution(x = training_meta_group_md)
```
Reactivation dataset
```{r fig.keep='none', warning=FALSE}
mlm.variance.distribution(x = reactivation_meta_group_md)
```

And with articles as clusters:
Complete dataset
```{r fig.keep='none', warning=FALSE}
mlm.variance.distribution(x = full_data_meta_article_md)
```
Training dataset
```{r fig.keep='none', warning=FALSE}
mlm.variance.distribution(x = training_meta_article_md)
```
Reactivation dataset
```{r fig.keep='none', warning=FALSE}
mlm.variance.distribution(x = reactivation_meta_article_md)
```

(Suppl table 10 - meta-regression for articles and groups as moderators)
Complete, articles as moderators:
```{r}
full_data_meta_article_md_mod = rma(yi=yi, vi=vi, data = full_data_md, measure = "MD", method = "REML", mods = ~factor(full_data_md$ID))
summary(full_data_meta_article_md_mod)
full_data_meta_article_md_mod$QMp
```

Complete, groups as moderators:
```{r}
full_data_meta_group_md_mod = rma(yi=yi, vi=vi, data = full_data_md, measure = "MD", method = "REML", mods = ~factor(full_data_md$group))
summary(full_data_meta_group_md_mod)
full_data_meta_group_md_mod$QMp
```

Training, articles as moderators:
```{r}
training_meta_article_md_mod = rma(yi=yi, vi=vi, data = training_md, measure = "MD", method = "REML", mods = ~factor(training_md$ID))
summary(training_meta_article_md_mod)
training_meta_article_md_mod$QMp
```

Training, groups as moderators:
```{r}
training_meta_group_md_mod = rma(yi=yi, vi=vi, data = training_md, measure = "MD", method = "REML", mods = ~factor(training_md$group))
summary(training_meta_group_md_mod)
training_meta_group_md_mod$QMp
```

Reactivation, articles as moderators:
```{r}
reactivation_meta_article_md_mod = rma(yi=yi, vi=vi, data = reactivation_md, measure = "MD", method = "REML", mods = ~factor(reactivation_md$ID))
summary(reactivation_meta_article_md_mod)
reactivation_meta_article_md_mod$QMp
```

Reactivation, groups as moderators:
```{r}
reactivation_meta_group_md_mod = rma(yi=yi, vi=vi, data = reactivation_md, measure = "MD", method = "REML", mods = ~factor(reactivation_md$group))
summary(reactivation_meta_group_md_mod)
reactivation_meta_group_md_mod$QMp
```

(Suppl table 11 - four-level model with both articles and groups)
Complete dataset:
```{r}
full_data_meta_md_art_group = rma.mv(yi=yi, V=vi, data = full_data_md, method = "REML", random = ~1|group/ID/Comparison.Code)
summary(full_data_meta_md_art_group)
full_data_meta_md_art_group$pval
```
```{r}
confint(full_data_meta_md_art_group)
```

Training dataset:
```{r}
training_meta_md_art_group = rma.mv(yi=yi, V=vi, data = training_md, method = "REML", random = ~1|group/ID/Comparison.Code)
summary(training_meta_md_art_group)
training_meta_md_art_group$pval
```
```{r}
confint(training_meta_md_art_group)
```

Reactivation dataset:
```{r}
reactivation_meta_md_art_group = rma.mv(yi=yi, V=vi, data = reactivation_md, method = "REML", random = ~1|group/ID/Comparison.Code)
summary(reactivation_meta_md_art_group)
reactivation_meta_md_art_group$pval
```
```{r}
confint(reactivation_meta_md_art_group)
```

**Univariate 2-level meta-regressions**

- First, we got the complete set of results to fill in the detailed tables for each dataset.

Complete dataset:
(Suppl Table 12)
```{r}
full_data_md %>% count(ConditioningType)
conditioning = rma(yi=yi, vi=vi, data = full_data_md, measure = "MD", method = "REML", mods = ~factor(full_data_md$ConditioningType))
summary(conditioning)
```
Exact model p values: `r conditioning$pval` | 
Exact Q test of moderators p value: `r conditioning$QMp`
```{r}
full_data_md %>% count(Habituation)
habituation = rma(yi=yi, vi=vi, data = full_data_md, measure = "MD", method = "REML", mods = ~factor(full_data_md$Habituation))
summary(habituation)
```
Exact model p values: `r habituation$pval` | 
Exact Q test of moderators p value: `r habituation$QMp`
```{r}
full_data_md %>% count(Handling)
handling = rma(yi=yi, vi=vi, data = full_data_md, measure = "MD", method = "REML", mods = ~factor(full_data_md$Handling))
summary(handling)
```
Exact model p values: `r handling$pval` | 
Exact Q test of moderators p value: `r handling$QMp`
```{r}
shocks = rma(yi=yi, vi=vi, data = full_data_md, measure = "MD", method = "REML", mods = full_data_md$NumberShock)
summary(shocks)
```
Exact model p values: `r shocks$pval` | 
Exact Q test of moderators p value: `r shocks$QMp`
```{r}
intensity = rma(yi=yi, vi=vi, data = full_data_md, measure = "MD", method = "REML", mods = full_data_md$ShockIntensity)
summary(intensity)
```
Exact model p values: `r intensity$pval` | 
Exact Q test of moderators p value: `r intensity$QMp`
```{r}
time_intervention = rma(yi=yi, vi=vi, data = full_data_md, measure = "MD", method = "REML", mods = as.numeric(full_data_md$InterventionTime))
summary(time_intervention)
```
Exact model p values: `r time_intervention$pval` | 
Exact Q test of moderators p value: `r time_intervention$QMp`
```{r}
time_test = rma(yi=yi, vi=vi, data = full_data_md, measure = "MD", method = "REML", mods = as.numeric(full_data_md$TimetoTest))
summary(time_test)
```
Exact model p values: `r time_test$pval` | 
Exact Q test of moderators p value: `r time_test$QMp`
```{r}
full_data_md %>% count(Species)
species = rma(yi=yi, vi=vi, data = full_data_md, measure = "MD", method = "REML", mods = ~factor(full_data_md$Species))
summary(species)
```
Exact model p values: `r species$pval` | 
Exact Q test of moderators p value: `r species$QMp`
```{r}
full_data_md %>% count(Sex)
sex = rma(yi=yi, vi=vi, data = full_data_md, measure = "MD", method = "REML", mods = ~factor(full_data_md$Sex))
summary(sex)
```
Exact model p values: `r sex$pval` | 
Exact Q test of moderators p value: `r sex$QMp`
```{r}
age = rma(yi=yi, vi=vi, data = full_data_md, measure = "MD", method = "REML", mods = full_data_md$Age_mean)
summary(age)
```
Exact model p values: `r age$pval` | 
Exact Q test of moderators p value: `r age$QMp`
```{r}
housing = rma(yi=yi, vi=vi, data = full_data_md, measure = "MD", method = "REML", mods = full_data_md$Housing)
summary(housing)
```
Exact model p values: `r housing$pval` | 
Exact Q test of moderators p value: `r housing$QMp`
```{r}
full_data_md %>% count(Active.Drug)
drug = rma(yi=yi, vi=vi, data = full_data_md, measure = "MD", method = "REML", mods = ~relevel(factor(full_data_md$Active.Drug), ref = "Anisomycin"))
summary(drug)
```
Exact model p values: `r drug$pval` | 
Exact Q test of moderators p value: `r drug$QMp`
```{r}
full_data_md %>% count(Site.injection.other)
injection = rma(yi=yi, vi=vi, data = full_data_md, measure = "MD", method = "REML", mods = ~relevel(factor(full_data_md$Site.injection.other), ref = "Systemic"))
summary(injection)
```
Exact model p values: `r injection$pval` | 
Exact Q test of moderators p value: `r injection$QMp`
```{r}
#dose = rma(yi=yi, vi=vi, data = full_data_md, measure = "MD", method = "REML", mods = full_data_md$dose_zscore)
#summary(dose)
```

```{r}
dose_s = rma(yi=yi, vi=vi, data = full_data_md, measure = "MD", method = "REML", mods = full_data_md$dose_zscore_s)
summary(dose_s)
```
Exact model p values: `r dose_s$pval` | 
Exact Q test of moderators p value: `r dose_s$QMp`

```{r}
full_data_md %>% count(Randomization)
randomization = rma(yi=yi, vi=vi, data = full_data_md, measure = "MD", method = "REML", mods = ~factor(full_data_md$Randomization))
summary(randomization)
```
Exact model p values: `r randomization$pval` | 
Exact Q test of moderators p value: `r randomization$QMp`
```{r}
full_data_md %>% count(Blinding)
blinding = rma(yi=yi, vi=vi, data = full_data_md, measure = "MD", method = "REML", mods = ~factor(full_data_md$Blinding))
summary(blinding)
```
Exact model p values: `r blinding$pval` | 
Exact Q test of moderators p value: `r blinding$QMp`
```{r}
full_data_md %>% count(SampleSizeCalc)
samplesize = rma(yi=yi, vi=vi, data = full_data_md, measure = "MD", method = "REML", mods = ~factor(full_data_md$SampleSizeCalc))
summary(samplesize)
```
Exact model p values: `r samplesize$pval` | 
Exact Q test of moderators p value: `r samplesize$QMp`
```{r}
full_data_md %>% count(RegulatoryReq)
requirements = rma(yi=yi, vi=vi, data = full_data_md, measure = "MD", method = "REML", mods = ~factor(full_data_md$RegulatoryReq))
summary(requirements)
```
Exact model p values: `r requirements$pval` | 
Exact Q test of moderators p value: `r requirements$QMp`
```{r}
full_data_md %>% count(ConflictInterest)
conflict = rma(yi=yi, vi=vi, data = full_data_md, measure = "MD", method = "REML", mods = ~relevel(factor(full_data_md$ConflictInterest), ref = "Not informed"))
summary(conflict)
```
Exact model p values: `r conflict$pval` | 
Exact Q test of moderators p value: `r conflict$QMp`
```{r}
impact = rma(yi=yi, vi=vi, data = full_data_md, measure = "MD", method = "REML", mods = full_data_md$Impact.Factor)
summary(impact)
```
Exact model p values: `r impact$pval` | 
Exact Q test of moderators p value: `r impact$QMp`
```{r}
citations = rma(yi=yi, vi=vi, data = full_data_md, measure = "MD", method = "REML", mods = full_data_md$Citations.per.year)
summary(citations)
```
Exact model p values: `r citations$pval` | 
Exact Q test of moderators p value: `r citations$QMp`
```{r}
full_data_md %>% count(Region.of.origin)
region = rma(yi=yi, vi=vi, data = full_data_md, measure = "MD", method = "REML", mods = ~relevel(factor(full_data_md$Region.of.origin), ref = "North America"))
summary(region)
```
Exact model p values: `r region$pval` | 
Exact Q test of moderators p value: `r region$QMp`

Training dataset:
(Suppl Table 13)
```{r}
training_md %>% count(ConditioningType)
conditioning_t = rma(yi=yi, vi=vi, data = training_md, measure = "MD", method = "REML", mods = ~factor(training_md$ConditioningType))
summary(conditioning_t)
```
Exact model p values: `r conditioning_t$pval` | 
Exact Q test of moderators p value: `r conditioning_t$QMp`
```{r}
training_md %>% count(Habituation)
habituation_t = rma(yi=yi, vi=vi, data = training_md, measure = "MD", method = "REML", mods = ~factor(training_md$Habituation))
summary(habituation_t)
```
Exact model p values: `r habituation_t$pval` | 
Exact Q test of moderators p value: `r habituation_t$QMp`
```{r}
training_md %>% count(Handling)
handling_t = rma(yi=yi, vi=vi, data = training_md, measure = "MD", method = "REML", mods = ~factor(training_md$Handling))
summary(handling_t)
```
Exact model p values: `r handling_t$pval` | 
Exact Q test of moderators p value: `r handling_t$QMp`
```{r}
shocks_t = rma(yi=yi, vi=vi, data = training_md, measure = "MD", method = "REML", mods = training_md$NumberShock)
summary(shocks_t)
```
Exact model p values: `r shocks_t$pval` | 
Exact Q test of moderators p value: `r shocks_t$QMp`
```{r}
intensity_t = rma(yi=yi, vi=vi, data = training_md, measure = "MD", method = "REML", mods = training_md$ShockIntensity)
summary(intensity_t)
```
Exact model p values: `r intensity_t$pval` | 
Exact Q test of moderators p value: `r intensity_t$QMp`
```{r}
time_intervention_t = rma(yi=yi, vi=vi, data = training_md, measure = "MD", method = "REML", mods = as.numeric(training_md$InterventionTime))
summary(time_intervention_t)
```
Exact model p values: `r time_intervention_t$pval` | 
Exact Q test of moderators p value: `r time_intervention_t$QMp`
```{r}
time_test_t = rma(yi=yi, vi=vi, data = training_md, measure = "MD", method = "REML", mods = as.numeric(training_md$TimetoTest))
summary(time_test_t)
```
Exact model p values: `r time_test_t$pval` | 
Exact Q test of moderators p value: `r time_test_t$QMp`
```{r}
training_md %>% count(Species)
species_t = rma(yi=yi, vi=vi, data = training_md, measure = "MD", method = "REML", mods = ~factor(training_md$Species))
summary(species_t)
```
Exact model p values: `r species_t$pval` | 
Exact Q test of moderators p value: `r species_t$QMp`
```{r}
training_md %>% count(Sex)
sex_t = rma(yi=yi, vi=vi, data = training_md, measure = "MD", method = "REML", mods = ~factor(training_md$Sex))
summary(sex_t)
```
Exact model p values: `r sex_t$pval` | 
Exact Q test of moderators p value: `r sex_t$QMp`
```{r}
age_t = rma(yi=yi, vi=vi, data = training_md, measure = "MD", method = "REML", mods = training_md$Age_mean)
summary(age_t)
```
Exact model p values: `r age_t$pval` | 
Exact Q test of moderators p value: `r age_t$QMp`
```{r}
housing_t = rma(yi=yi, vi=vi, data = training_md, measure = "MD", method = "REML", mods = training_md$Housing)
summary(housing_t)
```
Exact model p values: `r housing_t$pval` | 
Exact Q test of moderators p value: `r housing_t$QMp`
```{r}
training_md %>% count(Active.Drug)
drug_t = rma(yi=yi, vi=vi, data = training_md, measure = "MD", method = "REML", mods = ~relevel(factor(training_md$Active.Drug), ref = "Anisomycin"))
summary(drug_t)
```
Exact model p values: `r drug_t$pval` | 
Exact Q test of moderators p value: `r drug_t$QMp`
```{r}
#dose_t = rma(yi=yi, vi=vi, data = training_md, measure = "MD", method = "REML", mods = training_md$dose_zscore)
#summary(dose_t)
```

```{r}
dose_t_s = rma(yi=yi, vi=vi, data = training_md, measure = "MD", method = "REML", mods = training_md$dose_zscore_s)
summary(dose_t_s)
```
Exact model p values: `r dose_t_s$pval` | 
Exact Q test of moderators p value: `r dose_t_s$QMp`

```{r}
training_md %>% count(Randomization)
randomization_t = rma(yi=yi, vi=vi, data = training_md, measure = "MD", method = "REML", mods = ~factor(training_md$Randomization))
summary(randomization_t)
```
Exact model p values: `r randomization_t$pval` | 
Exact Q test of moderators p value: `r randomization_t$QMp`
```{r}
training_md %>% count(Blinding)
blinding_t = rma(yi=yi, vi=vi, data = training_md, measure = "MD", method = "REML", mods = ~factor(training_md$Blinding))
summary(blinding_t)
```
Exact model p values: `r blinding_t$pval` | 
Exact Q test of moderators p value: `r blinding_t$QMp`
```{r}
training_md %>% count(SampleSizeCalc)
```
```{r}
training_md %>% count(RegulatoryReq)
requirements_t = rma(yi=yi, vi=vi, data = training_md, measure = "MD", method = "REML", mods = ~factor(training_md$RegulatoryReq))
summary(requirements_t)
```
Exact model p values: `r requirements_t$pval` | 
Exact Q test of moderators p value: `r requirements_t$QMp`
```{r}
training_md %>% count(ConflictInterest)
conflict_t = rma(yi=yi, vi=vi, data = training_md, measure = "MD", method = "REML", mods = ~relevel(factor(training_md$ConflictInterest), ref = "Not informed"))
summary(conflict_t)
```
Exact model p values: `r conflict_t$pval` | 
Exact Q test of moderators p value: `r conflict_t$QMp`
```{r}
impact_t = rma(yi=yi, vi=vi, data = training_md, measure = "MD", method = "REML", mods = training_md$Impact.Factor)
summary(impact_t)
```
Exact model p values: `r impact_t$pval` | 
Exact Q test of moderators p value: `r impact_t$QMp`
```{r}
citations_t = rma(yi=yi, vi=vi, data = training_md, measure = "MD", method = "REML", mods = training_md$Citations.per.year)
summary(citations_t)
```
Exact model p values: `r citations_t$pval` | 
Exact Q test of moderators p value: `r citations_t$QMp`
```{r}
training_md %>% count(Region.of.origin)
region_t = rma(yi=yi, vi=vi, data = training_md, measure = "MD", method = "REML", mods = ~relevel(factor(training_md$Region.of.origin), ref = "North America"))
summary(region_t)
```
Exact model p values: `r region_t$pval` | 
Exact Q test of moderators p value: `r region_t$QMp`

Reactivation dataset:
(Suppl Table 14)
```{r}
reactivation_md %>% count(ConditioningType)
reactivation_md %>% group_by(Intervention.Session) %>% count(ConditioningType)
conditioning_r = rma(yi=yi, vi=vi, data = reactivation_md, measure = "MD", method = "REML", mods = ~factor(reactivation_md$ConditioningType))
summary(conditioning_r)
```
Exact model p values: `r conditioning_r$pval` | 
Exact Q test of moderators p value: `r conditioning_r$QMp`
```{r}
reactivation_md %>% count(Habituation)
habituation_r = rma(yi=yi, vi=vi, data = reactivation_md, measure = "MD", method = "REML", mods = ~factor(reactivation_md$Habituation))
summary(habituation_r)
```
Exact model p values: `r habituation_r$pval` | 
Exact Q test of moderators p value: `r habituation_r$QMp`
```{r}
reactivation_md %>% count(Handling)
handling_r = rma(yi=yi, vi=vi, data = reactivation_md, measure = "MD", method = "REML", mods = ~factor(reactivation_md$Handling))
summary(handling_r)
```
Exact model p values: `r handling_r$pval` | 
Exact Q test of moderators p value: `r handling_r$QMp`
```{r}
shocks_r = rma(yi=yi, vi=vi, data = reactivation_md, measure = "MD", method = "REML", mods = reactivation_md$NumberShock)
summary(shocks_r)
```
Exact model p values: `r shocks_r$pval` | 
Exact Q test of moderators p value: `r shocks_r$QMp`
```{r}
intensity_r = rma(yi=yi, vi=vi, data = reactivation_md, measure = "MD", method = "REML", mods = reactivation_md$ShockIntensity)
summary(intensity_r)
```
Exact model p values: `r intensity_r$pval` | 
Exact Q test of moderators p value: `r intensity_r$QMp`
```{r}
time_reexposure_r = rma(yi=yi, vi=vi, data = reactivation_md, measure = "MD", method = "REML", mods = as.numeric(reactivation_md$Time.to.reexposure))
summary(time_reexposure_r)
```
Exact model p values: `r time_reexposure_r$pval` | 
Exact Q test of moderators p value: `r time_reexposure_r$QMp`
```{r}
time_intervention_r = rma(yi=yi, vi=vi, data = reactivation_md, measure = "MD", method = "REML", mods = as.numeric(reactivation_md$InterventionTime))
summary(time_intervention_r)
```
Exact model p values: `r time_intervention_r$pval` | 
Exact Q test of moderators p value: `r time_intervention_r$QMp`
```{r}
reactivation_md %>% filter(ConditioningType=="Contextual"|ConditioningType=="Contextual background") %>% group_by(Intervention.Session) %>% count(is.na(ReexposureDuration_min))
reactivation_md %>% group_by(Intervention.Session) %>% count(is.na(ReexposureDuration_number))
duration_r = rma(yi=yi, vi=vi, data = reactivation_md, measure = "MD", method = "REML", mods = as.numeric(reactivation_md$duration_zscore))
summary(duration_r)
```
Exact model p values: `r duration_r$pval` | 
Exact Q test of moderators p value: `r duration_r$QMp`

```{r}
time_test_r = rma(yi=yi, vi=vi, data = reactivation_md, measure = "MD", method = "REML", mods = as.numeric(reactivation_md$TimetoTest))
summary(time_test_r)
```
Exact model p values: `r time_test_r$pval` | 
Exact Q test of moderators p value: `r time_test_r$QMp`
```{r}
reactivation_md %>% count(Species)
species_r = rma(yi=yi, vi=vi, data = reactivation_md, measure = "MD", method = "REML", mods = ~factor(reactivation_md$Species))
summary(species_r)
```
Exact model p values: `r species_r$pval` | 
Exact Q test of moderators p value: `r species_r$QMp`
```{r}
reactivation_md %>% count(Sex)
sex_r = rma(yi=yi, vi=vi, data = reactivation_md, measure = "MD", method = "REML", mods = ~factor(reactivation_md$Sex))
summary(sex_r)
```
Exact model p values: `r sex_r$pval` | 
Exact Q test of moderators p value: `r sex_r$QMp`
```{r}
age_r = rma(yi=yi, vi=vi, data = reconsolidation_md, measure = "MD", method = "REML", mods = reconsolidation_md$Age_mean)
summary(age_r)
```
Exact model p values: `r age_r$pval` | 
Exact Q test of moderators p value: `r age_r$QMp`
```{r}
housing_r = rma(yi=yi, vi=vi, data = reactivation_md, measure = "MD", method = "REML", mods = reactivation_md$Housing)
summary(housing_r)
```
Exact model p values: `r housing_r$pval` | 
Exact Q test of moderators p value: `r housing_r$QMp`
```{r}
reactivation_md %>% count(Active.Drug)
drug_r = rma(yi=yi, vi=vi, data = reactivation_md, measure = "MD", method = "REML", mods = ~relevel(factor(reactivation_md$Active.Drug), ref = "Anisomycin"))
summary(drug_r)
```
Exact model p values: `r drug_r$pval` | 
Exact Q test of moderators p value: `r drug_r$QMp`
```{r}
reactivation_md %>% count(Site.injection.other)
injection_r = rma(yi=yi, vi=vi, data = reactivation_md, measure = "MD", method = "REML", mods = ~relevel(factor(reactivation_md$Site.injection.other), ref = "Systemic"))
summary(injection_r)
```
Exact model p values: `r injection_r$pval` | 
Exact Q test of moderators p value: `r injection_r$QMp`
```{r}
#dose_r = rma(yi=yi, vi=vi, data = reactivation_md, measure = "MD", method = "REML", mods = reactivation_md$dose_zscore)
#summary(dose_r)
```

```{r}
dose_r_s = rma(yi=yi, vi=vi, data = reactivation_md, measure = "MD", method = "REML", mods = reactivation_md$dose_zscore_s)
summary(dose_r_s)
```
Exact model p values: `r dose_r_s$pval` | 
Exact Q test of moderators p value: `r dose_r_s$QMp`

```{r}
reactivation_md %>% count(Randomization)
randomization_r = rma(yi=yi, vi=vi, data = reactivation_md, measure = "MD", method = "REML", mods = ~factor(reactivation_md$Randomization))
summary(randomization_r)
```
Exact model p values: `r randomization_r$pval` | 
Exact Q test of moderators p value: `r randomization_r$QMp`
```{r}
reactivation_md %>% count(Blinding)
blinding_r = rma(yi=yi, vi=vi, data = reactivation_md, measure = "MD", method = "REML", mods = ~factor(reactivation_md$Blinding))
summary(blinding_r)
```
Exact model p values: `r blinding_r$pval` | 
Exact Q test of moderators p value: `r blinding_r$QMp`
```{r}
reactivation_md %>% count(SampleSizeCalc)
samplesize_r = rma(yi=yi, vi=vi, data = reactivation_md, measure = "MD", method = "REML", mods = ~factor(reactivation_md$SampleSizeCalc))
summary(samplesize_r)
```
Exact model p values: `r samplesize_r$pval` | 
Exact Q test of moderators p value: `r samplesize_r$QMp`
```{r}
reactivation_md %>% count(RegulatoryReq)
requirements_r = rma(yi=yi, vi=vi, data = reactivation_md, measure = "MD", method = "REML", mods = ~factor(reactivation_md$RegulatoryReq))
summary(requirements_r)
```
Exact model p values: `r requirements_r$pval` | 
Exact Q test of moderators p value: `r requirements_r$QMp`
```{r}
reactivation_md %>% count(ConflictInterest)
conflict_r = rma(yi=yi, vi=vi, data = reactivation_md, measure = "MD", method = "REML", mods = ~relevel(factor(reactivation_md$ConflictInterest), ref = "Not informed"))
summary(conflict_r)
```
Exact model p values: `r conflict_r$pval` | 
Exact Q test of moderators p value: `r conflict_r$QMp`
```{r}
impact_r = rma(yi=yi, vi=vi, data = reactivation_md, measure = "MD", method = "REML", mods = reactivation_md$Impact.Factor)
summary(impact_r)
```
Exact model p values: `r impact_r$pval` | 
Exact Q test of moderators p value: `r impact_r$QMp`
```{r}
citations_r = rma(yi=yi, vi=vi, data = reactivation_md, measure = "MD", method = "REML", mods = reactivation_md$Citations.per.year)
summary(citations_r)
```
Exact model p values: `r citations_r$pval` | 
Exact Q test of moderators p value: `r citations_r$QMp`
```{r}
reactivation_md %>% count(Region.of.origin)
region_r = rma(yi=yi, vi=vi, data = reactivation_md, measure = "MD", method = "REML", mods = ~relevel(factor(reactivation_md$Region.of.origin), ref = "North America"))
summary(region_r)
```
Exact model p values: `r region_r$pval` | 
Exact Q test of moderators p value: `r region_r$QMp`

- Then, we got only the necessary information to make summarized figures
```{r}
dataset = c('Complete')
moderadores = c('Conditioning Type', 'Habituation reported', 'Handling reported', 'Number of shocks', 'Shock intensity', 'Intervention time', 'Time to reexposure', 'Reexposure duration', 'Time to test', 'Species', 'Sex', 'Mean age', 'Housing', 'Active drug', 'Injection site', 'Dose', 'Randomization', 'Blinding', 'Sample size calculation', 'Regulatory requirements', 'Conflict of interest statement', 'Impact factor', 'Citations per year', 'Region of origin')
rquadrado = c(conditioning$R2, habituation$R2, handling$R2, shocks$R2, intensity$R2, time_intervention$R2, NA, NA, time_test$R2, species$R2, sex$R2, age$R2, housing$R2, drug$R2, injection$R2, dose_s$R2, randomization$R2, blinding$R2, samplesize$R2, requirements$R2, conflict$R2, impact$R2, citations$R2, region$R2)
pvalor = c(conditioning$QMp, habituation$QMp, handling$QMp, shocks$QMp, intensity$QMp, time_intervention$QMp, NA, NA, time_test$QMp, species$QMp, sex$QMp, age$QMp, housing$QMp, drug$QMp, injection$QMp, dose_s$QMp, randomization$QMp, blinding$QMp, samplesize$QMp, requirements$QMp, conflict$QMp, impact$QMp, citations$QMp, region$QMp)

moderators = data.frame(dataset, moderadores, rquadrado, pvalor)
moderators = moderators %>% mutate(pval_cat = if_else(pvalor>0.05, ">0.05", if_else(pvalor<=0.05&pvalor>0.01, "<0.05", if_else(pvalor<=0.01&pvalor>0.001, "<0.01", "<0.001"))))
```
```{r}
dataset = c('Training')
rquadrado = c(conditioning_t$R2, habituation_t$R2, handling_t$R2, shocks_t$R2, intensity_t$R2, time_intervention_t$R2, NA, NA, time_test_t$R2, species_t$R2, sex_t$R2, age_t$R2, housing_t$R2, drug_t$R2, training_md_sites$R2, dose_t_s$R2, randomization_t$R2, blinding_t$R2, NA, requirements_t$R2, conflict_t$R2, impact_t$R2, citations_t$R2, region_t$R2)
pvalor = c(conditioning_t$QMp, habituation_t$QMp, handling_t$QMp, shocks_t$QMp, intensity_t$QMp, time_intervention_t$QMp, NA, NA, time_test_t$QMp, species_t$QMp, sex_t$QMp, age_t$QMp, housing_t$QMp, drug_t$QMp, training_md_sites$QMp, dose_t_s$QMp, randomization_t$QMp, blinding_t$QMp, NA, requirements_t$QMp, conflict_t$QMp, impact_t$QMp, citations_t$QMp, region_t$QMp)

moderators_t = data.frame(dataset, moderadores, rquadrado, pvalor)
moderators_t = moderators_t %>% mutate(pval_cat = if_else(pvalor>0.05, ">0.05", if_else(pvalor<=0.05&pvalor>0.01, "<0.05", if_else(pvalor<=0.01&pvalor>0.001, "<0.01", "<0.001"))))
```
```{r}
dataset = c('Reactivation')
rquadrado = c(conditioning_r$R2, habituation_r$R2, handling_r$R2, shocks_r$R2, intensity_r$R2, time_intervention_r$R2, time_reexposure_r$R2, duration_r$R2, time_test_r$R2, species_r$R2, sex_r$R2, age_r$R2, housing_r$R2, drug_r$R2, injection_r$R2, dose_r_s$R2, randomization_r$R2, blinding_r$R2, samplesize_r$R2, requirements_r$R2, conflict_r$R2, impact_r$R2, citations_r$R2, region_r$R2)
pvalor = c(conditioning_r$QMp, habituation_r$QMp, handling_r$QMp, shocks_r$QMp, intensity_r$QMp, time_intervention_r$QMp, time_reexposure_r$QMp, duration_r$QMp, time_test_r$QMp, species_r$QMp, sex_r$QMp, age_r$QMp, housing_r$QMp, drug_r$QMp, injection_r$QMp, dose_r_s$QMp, randomization_r$QMp, blinding_r$QMp, samplesize_r$QMp, requirements_r$QMp, conflict_r$QMp, impact_r$QMp, citations_r$QMp, region_r$QMp)

moderators_r = data.frame(dataset, moderadores, rquadrado, pvalor)
moderators_r = moderators_r %>% mutate(pval_cat = if_else(pvalor>0.05, ">0.05", if_else(pvalor<=0.05&pvalor>0.01, "<0.05", if_else(pvalor<=0.01&pvalor>0.001, "<0.01", "<0.001"))))
```

Figure with summary (R2 and p value) of all moderator analyses - Fig 3A
```{r}
all_moderators = bind_rows(moderators, moderators_t, moderators_r)
all_moderators$moderadores = factor(all_moderators$moderadores, levels = rev(factor(moderadores)))
all_moderators$dataset = factor(all_moderators$dataset, levels = c("Complete", "Training", "Reactivation"))
labels_2l = c("\nComplete", "\nTraining", "\nReactivation")

all_mods = ggplot(all_moderators, aes(x=dataset, y=moderadores, fill=pval_cat)) + 
  geom_tile() + scale_fill_manual(values=c("#3182bd","#6baed6","#bdd7e7","#eff3ff"), na.value = "white", name = "Q test for moderators \np value") +
  geom_text(aes(x=dataset, y=moderadores, label = if_else(condition = rquadrado>=0, true = paste(round(rquadrado, 1), "%"), false = " ")), color = "black", size = 6) +
  scale_x_discrete(position = "top", labels = labels_2l) + 
  theme(axis.title.x = element_blank(), axis.text.x.top = element_text(angle = 0, size = 14), axis.title.y = element_blank(), panel.background = element_blank(),  axis.ticks = element_blank(), legend.position = "bottom", axis.text = element_text(size=15), legend.text = element_text(size=12), legend.title = element_text(size=12))
all_mods
```

(Suppl Table 15)
```{r}
reactivation_md_context = reactivation_md %>% filter(ConditioningType=="Contextual"|ConditioningType=="Contextual background")
duration_r_context = rma(yi=yi, vi=vi, data = reactivation_md_context, measure = "MD", method = "REML", mods = as.numeric(reactivation_md_context$ReexposureDuration_min))
summary(duration_r_context)
```
Exact model p values: `r duration_r_context$pval` | 
Exact Q test of moderators p value: `r duration_r_context$QMp`
```{r}
reactivation_md_tone = reactivation_md %>% filter(ConditioningType=="Tone"|ConditioningType=="Tone-trace")
duration_r_tone = rma(yi=yi, vi=vi, data = reactivation_md_tone, measure = "MD", method = "REML", mods = as.numeric(reactivation_md_tone$ReexposureDuration_number))
summary(duration_r_tone)
```
Exact model p values: `r duration_r_tone$pval` | 
Exact Q test of moderators p value: `r duration_r_tone$QMp`


**Univariate 3-level meta-regressions (article)**
Complete dataset:
(Suppl Table 16)
```{r}
full_data_md %>% count(Intervention.Session)
session_3a = rma.mv(yi=yi, V=vi, data = full_data_md, method = "REML", random = ~1|ID/Comparison.Code, mods = ~relevel(factor(full_data_md$Intervention.Session), ref="Training"))
summary(session_3a)
#pseudo R-squared
session_3a.R2 = (sum(full_data_meta_article_md$sigma2)-sum(session_3a$sigma2))/sum(full_data_meta_article_md$sigma2)*100
session_3a.R2 = if_else(session_3a.R2<0, 0, session_3a.R2)
```
Exact model p values: `r session_3a$pval` | 
Exact Q test of moderators p value: `r session_3a$QMp` | 
Total I2: `r mlm.variance.distribution(session_3a)$totalI2`
```{r}
full_data_md %>% count(ConditioningType)
conditioning_3a = rma.mv(yi=yi, V=vi, data = full_data_md, method = "REML", random = ~1|ID/Comparison.Code, mods = ~factor(full_data_md$ConditioningType))
summary(conditioning_3a)
#pseudo R-squared
conditioning_3a.R2 = (sum(full_data_meta_article_md$sigma2)-sum(conditioning_3a$sigma2))/sum(full_data_meta_article_md$sigma2)*100
conditioning_3a.R2 = if_else(conditioning_3a.R2<0, 0, conditioning_3a.R2)
```
Exact model p values: `r conditioning_3a$pval` | 
Exact Q test of moderators p value: `r conditioning_3a$QMp` | 
Total I2: `r mlm.variance.distribution(conditioning_3a)$totalI2`
```{r}
full_data_md %>% count(Habituation)
habituation_3a = rma.mv(yi=yi, V=vi, data = full_data_md, method = "REML", random = ~1|ID/Comparison.Code, mods = ~factor(full_data_md$Habituation))
summary(habituation_3a)
#pseudo R-squared
habituation_3a.R2 = (sum(full_data_meta_article_md$sigma2)-sum(habituation_3a$sigma2))/sum(full_data_meta_article_md$sigma2)*100
habituation_3a.R2 = if_else(habituation_3a.R2<0, 0, habituation_3a.R2)
```
Exact model p values: `r habituation_3a$pval` | 
Exact Q test of moderators p value: `r habituation_3a$QMp` | 
Total I2: `r mlm.variance.distribution(habituation_3a)$totalI2`
```{r}
full_data_md %>% count(Handling)
handling_3a = rma.mv(yi=yi, V=vi, data = full_data_md, method = "REML", random = ~1|ID/Comparison.Code, mods = ~factor(full_data_md$Handling))
summary(handling_3a)
#pseudo R-squared
handling_3a.R2 = (sum(full_data_meta_article_md$sigma2)-sum(handling_3a$sigma2))/sum(full_data_meta_article_md$sigma2)*100
handling_3a.R2 = if_else(handling_3a.R2<0, 0, handling_3a.R2)
```
Exact model p values: `r handling_3a$pval` | 
Exact Q test of moderators p value: `r handling_3a$QMp` | 
Total I2: `r mlm.variance.distribution(handling_3a)$totalI2`
```{r}
shocks_3a = rma.mv(yi=yi, V=vi, data = full_data_md, method = "REML", random = ~1|ID/Comparison.Code, mods = full_data_md$NumberShock)
summary(shocks_3a)
#pseudo R-squared
shocks_3a.R2 = (sum(full_data_meta_article_md$sigma2)-sum(shocks_3a$sigma2))/sum(full_data_meta_article_md$sigma2)*100
shocks_3a.R2 = if_else(shocks_3a.R2<0, 0, shocks_3a.R2)
```
Exact model p values: `r shocks_3a$pval` | 
Exact Q test of moderators p value: `r shocks_3a$QMp` | 
Total I2: `r mlm.variance.distribution(shocks_3a)$totalI2`
```{r}
intensity_3a = rma.mv(yi=yi, V=vi, data = full_data_md, method = "REML", random = ~1|ID/Comparison.Code, mods = full_data_md$ShockIntensity)
summary(intensity_3a)
#pseudo R-squared
intensity_3a.R2 = (sum(full_data_meta_article_md$sigma2)-sum(intensity_3a$sigma2))/sum(full_data_meta_article_md$sigma2)*100
intensity_3a.R2 = if_else(intensity_3a.R2<0, 0, intensity_3a.R2)
```
Exact model p values: `r intensity_3a$pval` | 
Exact Q test of moderators p value: `r intensity_3a$QMp` | 
Total I2: `r mlm.variance.distribution(intensity_3a)$totalI2`
```{r}
time_intervention_3a = rma.mv(yi=yi, V=vi, data = full_data_md, method = "REML", random = ~1|ID/Comparison.Code, mods = full_data_md$InterventionTime)
summary(time_intervention_3a)
#pseudo R-squared
time_intervention_3a.R2 = (sum(full_data_meta_article_md$sigma2)-sum(time_intervention_3a$sigma2))/sum(full_data_meta_article_md$sigma2)*100
time_intervention_3a.R2 = if_else(time_intervention_3a.R2<0, 0, time_intervention_3a.R2)
```
Exact model p values: `r time_intervention_3a$pval` | 
Exact Q test of moderators p value: `r time_intervention_3a$QMp` | 
Total I2: `r mlm.variance.distribution(time_intervention_3a)$totalI2`
```{r}
time_test_3a = rma.mv(yi=yi, V=vi, data = full_data_md, method = "REML", random = ~1|ID/Comparison.Code, mods = full_data_md$TimetoTest)
summary(time_test_3a)
#pseudo R-squared
time_test_3a.R2 = (sum(full_data_meta_article_md$sigma2)-sum(time_test_3a$sigma2))/sum(full_data_meta_article_md$sigma2)*100
time_test_3a.R2 = if_else(time_test_3a.R2<0, 0, time_test_3a.R2)
```
Exact model p values: `r time_test_3a$pval` | 
Exact Q test of moderators p value: `r time_test_3a$QMp` | 
Total I2: `r mlm.variance.distribution(time_test_3a)$totalI2`
```{r}
full_data_md %>% count(Species)
species_3a = rma.mv(yi=yi, V=vi, data = full_data_md, method = "REML", random = ~1|ID/Comparison.Code, mods = ~factor(full_data_md$Species))
summary(species_3a)
#pseudo R-squared
species_3a.R2 = (sum(full_data_meta_article_md$sigma2)-sum(species_3a$sigma2))/sum(full_data_meta_article_md$sigma2)*100
species_3a.R2 = if_else(species_3a.R2<0, 0, species_3a.R2)
```
Exact model p values: `r species_3a$pval` | 
Exact Q test of moderators p value: `r species_3a$QMp` | 
Total I2: `r mlm.variance.distribution(species_3a)$totalI2`
```{r}
full_data_md %>% count(Sex)
sex_3a = rma.mv(yi=yi, V=vi, data = full_data_md, method = "REML", random = ~1|ID/Comparison.Code, mods = ~factor(full_data_md$Sex))
summary(sex_3a)
#pseudo R-squared
sex_3a.R2 = (sum(full_data_meta_article_md$sigma2)-sum(sex_3a$sigma2))/sum(full_data_meta_article_md$sigma2)*100
sex_3a.R2 = if_else(sex_3a.R2<0, 0, sex_3a.R2)
```
Exact model p values: `r sex_3a$pval` | 
Exact Q test of moderators p value: `r sex_3a$QMp` | 
Total I2: `r mlm.variance.distribution(sex_3a)$totalI2`
```{r}
age_3a = rma.mv(yi=yi, V=vi, data = full_data_md, method = "REML", random = ~1|ID/Comparison.Code, mods = full_data_md$Age_mean)
summary(age_3a)
#pseudo R-squared
age_3a.R2 = (sum(full_data_meta_article_md$sigma2)-sum(age_3a$sigma2))/sum(full_data_meta_article_md$sigma2)*100
age_3a.R2 = if_else(age_3a.R2<0, 0, age_3a.R2)
```
Exact model p values: `r age_3a$pval` | 
Exact Q test of moderators p value: `r age_3a$QMp` | 
Total I2: `r mlm.variance.distribution(age_3a)$totalI2`
```{r}
housing_3a = rma.mv(yi=yi, V=vi, data = full_data_md, method = "REML", random = ~1|ID/Comparison.Code, mods = full_data_md$Housing)
summary(housing_3a)
#pseudo R-squared
housing_3a.R2 = (sum(full_data_meta_article_md$sigma2)-sum(housing_3a$sigma2))/sum(full_data_meta_article_md$sigma2)*100
housing_3a.R2 = if_else(housing_3a.R2<0, 0, housing_3a.R2)
```
Exact model p values: `r housing_3a$pval` | 
Exact Q test of moderators p value: `r housing_3a$QMp` | 
Total I2: `r mlm.variance.distribution(housing_3a)$totalI2`
```{r}
full_data_md %>% count(Active.Drug)
drug_3a = rma.mv(yi=yi, V=vi, data = full_data_md, method = "REML", random = ~1|ID/Comparison.Code, mods = ~relevel(factor(full_data_md$Active.Drug), ref = "Anisomycin"))
summary(drug_3a)
#pseudo R-squared
drug_3a.R2 = (sum(full_data_meta_article_md$sigma2)-sum(drug_3a$sigma2))/sum(full_data_meta_article_md$sigma2)*100
drug_3a.R2 = if_else(drug_3a.R2<0, 0, drug_3a.R2)
```
Exact model p values: `r drug_3a$pval` | 
Exact Q test of moderators p value: `r drug_3a$QMp` | 
Total I2: `r mlm.variance.distribution(drug_3a)$totalI2`
```{r}
full_data_md %>% count(Site.injection.other)
sites_3a = rma.mv(yi=yi, V=vi, data = full_data_md, method = "REML", random = ~1|ID/Comparison.Code, mods = ~relevel(factor(full_data_md$Site.injection.other), ref = "Systemic"))
summary(sites_3a)
#pseudo R-squared
sites_3a.R2 = (sum(full_data_meta_article_md$sigma2)-sum(sites_3a$sigma2))/sum(full_data_meta_article_md$sigma2)*100
sites_3a.R2 = if_else(sites_3a.R2<0, 0, sites_3a.R2)
```
Exact model p values: `r sites_3a$pval` | 
Exact Q test of moderators p value: `r sites_3a$QMp` | 
Total I2: `r mlm.variance.distribution(sites_3a)$totalI2`
```{r}
#dose_3a = rma.mv(yi=yi, V=vi, data = full_data_md, method = "REML", random = ~1|ID/Comparison.Code, mods = full_data_md$dose_zscore)
#summary(dose_3a)
#pseudo R-squared
#dose_3a.R2 = (sum(full_data_meta_article_md$sigma2)-sum(dose_3a$sigma2))/sum(full_data_meta_article_md$sigma2)*100
#dose_3a.R2 = if_else(dose_3a.R2<0, 0, dose_3a.R2)
```

```{r}
dose_3a_s = rma.mv(yi=yi, V=vi, data = full_data_md, method = "REML", random = ~1|ID/Comparison.Code, mods = full_data_md$dose_zscore_s)
summary(dose_3a_s)
#pseudo R-squared
dose_3a.R2_s = (sum(full_data_meta_article_md$sigma2)-sum(dose_3a_s$sigma2))/sum(full_data_meta_article_md$sigma2)*100
dose_3a.R2_s = if_else(dose_3a.R2_s<0, 0, dose_3a.R2_s)
```
Exact model p values: `r dose_3a_s$pval` | 
Exact Q test of moderators p value: `r dose_3a_s$QMp` | 
Total I2: `r mlm.variance.distribution(dose_3a_s)$totalI2`

```{r}
full_data_md %>% count(Randomization)
randomization_3a = rma.mv(yi=yi, V=vi, data = full_data_md, method = "REML", random = ~1|ID/Comparison.Code, mods = ~factor(full_data_md$Randomization))
summary(randomization_3a)
#pseudo R-squared
randomization_3a.R2 = (sum(full_data_meta_article_md$sigma2)-sum(randomization_3a$sigma2))/sum(full_data_meta_article_md$sigma2)*100
randomization_3a.R2 = if_else(randomization_3a.R2<0, 0, randomization_3a.R2)
```
Exact model p values: `r randomization_3a$pval` | 
Exact Q test of moderators p value: `r randomization_3a$QMp` | 
Total I2: `r mlm.variance.distribution(randomization_3a)$totalI2`
```{r}
full_data_md %>% count(Blinding)
blinding_3a = rma.mv(yi=yi, V=vi, data = full_data_md, method = "REML", random = ~1|ID/Comparison.Code, mods = ~factor(full_data_md$Blinding))
summary(blinding_3a)
#pseudo R-squared
blinding_3a.R2 = (sum(full_data_meta_article_md$sigma2)-sum(blinding_3a$sigma2))/sum(full_data_meta_article_md$sigma2)*100
blinding_3a.R2 = if_else(blinding_3a.R2<0, 0, blinding_3a.R2)
```
Exact model p values: `r blinding_3a$pval` | 
Exact Q test of moderators p value: `r blinding_3a$QMp` | 
Total I2: `r mlm.variance.distribution(blinding_3a)$totalI2`
```{r}
full_data_md %>% count(SampleSizeCalc)
samplesize_3a = rma.mv(yi=yi, V=vi, data = full_data_md, method = "REML", random = ~1|ID/Comparison.Code, mods = ~factor(full_data_md$SampleSizeCalc))
summary(samplesize_3a)
#pseudo R-squared
samplesize_3a.R2 = (sum(full_data_meta_article_md$sigma2)-sum(samplesize_3a$sigma2))/sum(full_data_meta_article_md$sigma2)*100
samplesize_3a.R2 = if_else(samplesize_3a.R2<0, 0, samplesize_3a.R2)
```
Exact model p values: `r samplesize_3a$pval` | 
Exact Q test of moderators p value: `r samplesize_3a$QMp` | 
Total I2: `r mlm.variance.distribution(samplesize_3a)$totalI2`
```{r}
full_data_md %>% count(RegulatoryReq)
requirements_3a = rma.mv(yi=yi, V=vi, data = full_data_md, method = "REML", random = ~1|ID/Comparison.Code, mods = ~factor(full_data_md$RegulatoryReq))
summary(requirements_3a)
#pseudo R-squared
requirements_3a.R2 = (sum(full_data_meta_article_md$sigma2)-sum(requirements_3a$sigma2))/sum(full_data_meta_article_md$sigma2)*100
requirements_3a.R2 = if_else(requirements_3a.R2<0, 0, requirements_3a.R2)
```
Exact model p values: `r requirements_3a$pval` | 
Exact Q test of moderators p value: `r requirements_3a$QMp` | 
Total I2: `r mlm.variance.distribution(requirements_3a)$totalI2`
```{r}
full_data_md %>% count(ConflictInterest)
conflict_3a = rma.mv(yi=yi, V=vi, data = full_data_md, method = "REML", random = ~1|ID/Comparison.Code, mods = ~factor(full_data_md$ConflictInterest))
summary(conflict_3a)
#pseudo R-squared
conflict_3a.R2 = (sum(full_data_meta_article_md$sigma2)-sum(conflict_3a$sigma2))/sum(full_data_meta_article_md$sigma2)*100
conflict_3a.R2 = if_else(conflict_3a.R2<0, 0, conflict_3a.R2)
```
Exact model p values: `r conflict_3a$pval` | 
Exact Q test of moderators p value: `r conflict_3a$QMp` | 
Total I2: `r mlm.variance.distribution(conflict_3a)$totalI2`
```{r}
impact_3a = rma.mv(yi=yi, V=vi, data = full_data_md, method = "REML", random = ~1|ID/Comparison.Code, mods = full_data_md$Impact.Factor)
summary(impact_3a)
#pseudo R-squared
impact_3a.R2 = (sum(full_data_meta_article_md$sigma2)-sum(impact_3a$sigma2))/sum(full_data_meta_article_md$sigma2)*100
impact_3a.R2 = if_else(impact_3a.R2<0, 0, impact_3a.R2)
```
Exact model p values: `r impact_3a$pval` | 
Exact Q test of moderators p value: `r impact_3a$QMp` | 
Total I2: `r mlm.variance.distribution(impact_3a)$totalI2`
```{r}
citations_3a = rma.mv(yi=yi, V=vi, data = full_data_md, method = "REML", random = ~1|ID/Comparison.Code, mods = full_data_md$Citations.per.year)
summary(citations_3a)
#pseudo R-squared
citations_3a.R2 = (sum(full_data_meta_article_md$sigma2)-sum(citations_3a$sigma2))/sum(full_data_meta_article_md$sigma2)*100
citations_3a.R2 = if_else(citations_3a.R2<0, 0, citations_3a.R2)
```
Exact model p values: `r citations_3a$pval` | 
Exact Q test of moderators p value: `r citations_3a$QMp` | 
Total I2: `r mlm.variance.distribution(citations_3a)$totalI2`
```{r}
full_data_md %>% count(Region.of.origin)
region_3a = rma.mv(yi=yi, V=vi, data = full_data_md, method = "REML", random = ~1|ID/Comparison.Code, mods = ~relevel(factor(full_data_md$Region.of.origin), ref="North America"))
summary(region_3a)
#pseudo R-squared
region_3a.R2 = (sum(full_data_meta_article_md$sigma2)-sum(region_3a$sigma2))/sum(full_data_meta_article_md$sigma2)*100
region_3a.R2 = if_else(region_3a.R2<0, 0, region_3a.R2)
```
Exact model p values: `r region_3a$pval` | 
Exact Q test of moderators p value: `r region_3a$QMp` | 
Total I2: `r mlm.variance.distribution(region_3a)$totalI2`

Training dataset:
(Suppl Table 17)
```{r}
training_md %>% count(ConditioningType)
conditioning_3a_t = rma.mv(yi=yi, V=vi, data = training_md, method = "REML", random = ~1|ID/Comparison.Code, mods = ~factor(training_md$ConditioningType))
summary(conditioning_3a_t)
#pseudo R-squared
conditioning_3a_t.R2 = (sum(training_meta_article_md$sigma2)-sum(conditioning_3a_t$sigma2))/sum(training_meta_article_md$sigma2)*100
conditioning_3a_t.R2 = if_else(conditioning_3a_t.R2<0, 0, conditioning_3a_t.R2)
```
Exact model p values: `r conditioning_3a_t$pval` | 
Exact Q test of moderators p value: `r conditioning_3a_t$QMp` | 
Total I2: `r mlm.variance.distribution(conditioning_3a_t)$totalI2`
```{r}
training_md %>% count(Habituation)
habituation_3a_t = rma.mv(yi=yi, V=vi, data = training_md, method = "REML", random = ~1|ID/Comparison.Code, mods = ~factor(training_md$Habituation))
summary(habituation_3a_t)
#pseudo R-squared
habituation_3a_t.R2 = (sum(training_meta_article_md$sigma2)-sum(habituation_3a_t$sigma2))/sum(training_meta_article_md$sigma2)*100
habituation_3a_t.R2 = if_else(habituation_3a_t.R2<0, 0, habituation_3a_t.R2)
```
Exact model p values: `r habituation_3a_t$pval` | 
Exact Q test of moderators p value: `r habituation_3a_t$QMp` | 
Total I2: `r mlm.variance.distribution(habituation_3a_t)$totalI2`
```{r}
training_md %>% count(Handling)
handling_3a_t = rma.mv(yi=yi, V=vi, data = training_md, method = "REML", random = ~1|ID/Comparison.Code, mods = ~factor(training_md$Handling))
summary(handling_3a_t)
#pseudo R-squared
handling_3a_t.R2 = (sum(training_meta_article_md$sigma2)-sum(handling_3a_t$sigma2))/sum(training_meta_article_md$sigma2)*100
handling_3a_t.R2 = if_else(handling_3a_t.R2<0, 0, handling_3a_t.R2)
```
Exact model p values: `r handling_3a_t$pval` | 
Exact Q test of moderators p value: `r handling_3a_t$QMp` | 
Total I2: `r mlm.variance.distribution(handling_3a_t)$totalI2`
```{r}
shocks_3a_t = rma.mv(yi=yi, V=vi, data = training_md, method = "REML", random = ~1|ID/Comparison.Code, mods = training_md$NumberShock)
summary(shocks_3a_t)
#pseudo R-squared
shocks_3a_t.R2 = (sum(training_meta_article_md$sigma2)-sum(shocks_3a_t$sigma2))/sum(training_meta_article_md$sigma2)*100
shocks_3a_t.R2 = if_else(shocks_3a_t.R2<0, 0, shocks_3a_t.R2)
```
Exact model p values: `r shocks_3a_t$pval` | 
Exact Q test of moderators p value: `r shocks_3a_t$QMp` | 
Total I2: `r mlm.variance.distribution(shocks_3a_t)$totalI2`
```{r}
intensity_3a_t = rma.mv(yi=yi, V=vi, data = training_md, method = "REML", random = ~1|ID/Comparison.Code, mods = training_md$ShockIntensity)
summary(intensity_3a_t)
#pseudo R-squared
intensity_3a_t.R2 = (sum(training_meta_article_md$sigma2)-sum(intensity_3a_t$sigma2))/sum(training_meta_article_md$sigma2)*100
intensity_3a_t.R2 = if_else(intensity_3a_t.R2<0, 0, intensity_3a_t.R2)
```
Exact model p values: `r intensity_3a_t$pval` | 
Exact Q test of moderators p value: `r intensity_3a_t$QMp` | 
Total I2: `r mlm.variance.distribution(intensity_3a_t)$totalI2`
```{r}
time_intervention_3a_t = rma.mv(yi=yi, V=vi, data = training_md, method = "REML", random = ~1|ID/Comparison.Code, mods = training_md$InterventionTime)
summary(time_intervention_3a_t)
#pseudo R-squared
time_intervention_3a_t.R2 = (sum(training_meta_article_md$sigma2)-sum(time_intervention_3a_t$sigma2))/sum(training_meta_article_md$sigma2)*100
time_intervention_3a_t.R2 = if_else(time_intervention_3a_t.R2<0, 0, time_intervention_3a_t.R2)
```
Exact model p values: `r time_intervention_3a_t$pval` | 
Exact Q test of moderators p value: `r time_intervention_3a_t$QMp` | 
Total I2: `r mlm.variance.distribution(time_intervention_3a_t)$totalI2`
```{r}
time_test_3a_t = rma.mv(yi=yi, V=vi, data = training_md, method = "REML", random = ~1|ID/Comparison.Code, mods = training_md$TimetoTest)
summary(time_test_3a_t)
#pseudo R-squared
time_test_3a_t.R2 = (sum(training_meta_article_md$sigma2)-sum(time_test_3a_t$sigma2))/sum(training_meta_article_md$sigma2)*100
time_test_3a_t.R2 = if_else(time_test_3a_t.R2<0, 0, time_test_3a_t.R2)
```
Exact model p values: `r time_test_3a_t$pval` | 
Exact Q test of moderators p value: `r time_test_3a_t$QMp` | 
Total I2: `r mlm.variance.distribution(time_test_3a_t)$totalI2`
```{r}
training_md %>% count(Species)
species_3a_t = rma.mv(yi=yi, V=vi, data = training_md, method = "REML", random = ~1|ID/Comparison.Code, mods = ~factor(training_md$Species))
summary(species_3a_t)
#pseudo R-squared
species_3a_t.R2 = (sum(training_meta_article_md$sigma2)-sum(species_3a_t$sigma2))/sum(training_meta_article_md$sigma2)*100
species_3a_t.R2 = if_else(species_3a_t.R2<0, 0, species_3a_t.R2)
```
Exact model p values: `r species_3a_t$pval` | 
Exact Q test of moderators p value: `r species_3a_t$QMp` | 
Total I2: `r mlm.variance.distribution(species_3a_t)$totalI2`
```{r}
training_md %>% count(Sex)
sex_3a_t = rma.mv(yi=yi, V=vi, data = training_md, method = "REML", random = ~1|ID/Comparison.Code, mods = ~factor(training_md$Sex))
summary(sex_3a_t)
#pseudo R-squared
sex_3a_t.R2 = (sum(training_meta_article_md$sigma2)-sum(sex_3a_t$sigma2))/sum(training_meta_article_md$sigma2)*100
sex_3a_t.R2 = if_else(sex_3a_t.R2<0, 0, sex_3a_t.R2)
```
Exact model p values: `r sex_3a_t$pval` | 
Exact Q test of moderators p value: `r sex_3a_t$QMp` | 
Total I2: `r mlm.variance.distribution(sex_3a_t)$totalI2`
```{r}
age_3a_t = rma.mv(yi=yi, V=vi, data = training_md, method = "REML", random = ~1|ID/Comparison.Code, mods = training_md$Age_mean)
summary(age_3a_t)
#pseudo R-squared
age_3a_t.R2 = (sum(training_meta_article_md$sigma2)-sum(age_3a_t$sigma2))/sum(training_meta_article_md$sigma2)*100
age_3a_t.R2 = if_else(age_3a_t.R2<0, 0, age_3a_t.R2)
```
Exact model p values: `r age_3a_t$pval` | 
Exact Q test of moderators p value: `r age_3a_t$QMp` | 
Total I2: `r mlm.variance.distribution(age_3a_t)$totalI2`
```{r}
housing_3a_t = rma.mv(yi=yi, V=vi, data = training_md, method = "REML", random = ~1|ID/Comparison.Code, mods = training_md$Housing)
summary(housing_3a_t)
#pseudo R-squared
housing_3a_t.R2 = (sum(training_meta_article_md$sigma2)-sum(housing_3a_t$sigma2))/sum(training_meta_article_md$sigma2)*100
housing_3a_t.R2 = if_else(housing_3a_t.R2<0, 0, housing_3a_t.R2)
```
Exact model p values: `r housing_3a_t$pval` | 
Exact Q test of moderators p value: `r housing_3a_t$QMp` | 
Total I2: `r mlm.variance.distribution(housing_3a_t)$totalI2`
```{r}
training_md %>% count(Active.Drug)
drug_3a_t = rma.mv(yi=yi, V=vi, data = training_md, method = "REML", random = ~1|ID/Comparison.Code, mods = ~relevel(factor(training_md$Active.Drug), ref = "Anisomycin"))
summary(drug_3a_t)
#pseudo R-squared
drug_3a_t.R2 = (sum(training_meta_article_md$sigma2)-sum(drug_3a_t$sigma2))/sum(training_meta_article_md$sigma2)*100
drug_3a_t.R2 = if_else(drug_3a_t.R2<0, 0, drug_3a_t.R2)
```
Exact model p values: `r drug_3a_t$pval` | 
Exact Q test of moderators p value: `r drug_3a_t$QMp` | 
Total I2: `r mlm.variance.distribution(drug_3a_t)$totalI2`
```{r}
training_md %>% count(Site.injection.other)
sites_3a_t = rma.mv(yi=yi, V=vi, data = training_md, method = "REML", random = ~1|ID/Comparison.Code, mods = ~relevel(factor(training_md$Site.injection.other), ref = "Systemic"))
summary(sites_3a_t)
#pseudo R-squared
sites_3a_t.R2 = (sum(training_meta_article_md$sigma2)-sum(sites_3a_t$sigma2))/sum(training_meta_article_md$sigma2)*100
sites_3a_t.R2 = if_else(sites_3a_t.R2<0, 0, sites_3a_t.R2)
```
Exact model p values: `r sites_3a_t$pval` | 
Exact Q test of moderators p value: `r sites_3a_t$QMp` | 
Total I2: `r mlm.variance.distribution(sites_3a_t)$totalI2`
```{r}
#dose_3a_t = rma.mv(yi=yi, V=vi, data = training_md, method = "REML", random = ~1|ID/Comparison.Code, mods = training_md$dose_zscore)
#summary(dose_3a_t)
#pseudo R-squared
#dose_3a_t.R2 = (sum(training_meta_article_md$sigma2)-sum(dose_3a_t$sigma2))/sum(training_meta_article_md$sigma2)*100
#dose_3a_t.R2 = if_else(dose_3a_t.R2<0, 0, dose_3a_t.R2)
```

```{r}
dose_3a_t_s = rma.mv(yi=yi, V=vi, data = training_md, method = "REML", random = ~1|ID/Comparison.Code, mods = training_md$dose_zscore_s)
summary(dose_3a_t_s)
#pseudo R-squared
dose_3a_t.R2_s = (sum(training_meta_article_md$sigma2)-sum(dose_3a_t_s$sigma2))/sum(training_meta_article_md$sigma2)*100
dose_3a_t.R2_s = if_else(dose_3a_t.R2_s<0, 0, dose_3a_t.R2_s)
```
Exact model p values: `r dose_3a_t_s$pval` | 
Exact Q test of moderators p value: `r dose_3a_t_s$QMp` | 
Total I2: `r mlm.variance.distribution(dose_3a_t_s)$totalI2`

```{r}
training_md %>% count(Randomization)
randomization_3a_t = rma.mv(yi=yi, V=vi, data = training_md, method = "REML", random = ~1|ID/Comparison.Code, mods = ~factor(training_md$Randomization))
summary(randomization_3a_t)
#pseudo R-squared
randomization_3a_t.R2 = (sum(training_meta_article_md$sigma2)-sum(randomization_3a_t$sigma2))/sum(training_meta_article_md$sigma2)*100
randomization_3a_t.R2 = if_else(randomization_3a_t.R2<0, 0, randomization_3a_t.R2)
```
Exact model p values: `r randomization_3a_t$pval` | 
Exact Q test of moderators p value: `r randomization_3a_t$QMp` | 
Total I2: `r mlm.variance.distribution(randomization_3a_t)$totalI2`
```{r}
training_md %>% count(Blinding)
blinding_3a_t = rma.mv(yi=yi, V=vi, data = training_md, method = "REML", random = ~1|ID/Comparison.Code, mods = ~factor(training_md$Blinding))
summary(blinding_3a_t)
#pseudo R-squared
blinding_3a_t.R2 = (sum(training_meta_article_md$sigma2)-sum(blinding_3a_t$sigma2))/sum(training_meta_article_md$sigma2)*100
blinding_3a_t.R2 = if_else(blinding_3a_t.R2<0, 0, blinding_3a_t.R2)
```
Exact model p values: `r blinding_3a_t$pval` | 
Exact Q test of moderators p value: `r blinding_3a_t$QMp` | 
Total I2: `r mlm.variance.distribution(blinding_3a_t)$totalI2`
```{r}
training_md %>% count(RegulatoryReq)
requirements_3a_t = rma.mv(yi=yi, V=vi, data = training_md, method = "REML", random = ~1|ID/Comparison.Code, mods = ~factor(training_md$RegulatoryReq))
summary(requirements_3a_t)
#pseudo R-squared
requirements_3a_t.R2 = (sum(training_meta_article_md$sigma2)-sum(requirements_3a_t$sigma2))/sum(training_meta_article_md$sigma2)*100
requirements_3a_t.R2 = if_else(requirements_3a_t.R2<0, 0, requirements_3a_t.R2)
```
Exact model p values: `r requirements_3a_t$pval` | 
Exact Q test of moderators p value: `r requirements_3a_t$QMp` | 
Total I2: `r mlm.variance.distribution(requirements_3a_t)$totalI2`
```{r}
training_md %>% count(ConflictInterest)
conflict_3a_t = rma.mv(yi=yi, V=vi, data = training_md, method = "REML", random = ~1|ID/Comparison.Code, mods = ~factor(training_md$ConflictInterest))
summary(conflict_3a_t)
#pseudo R-squared
conflict_3a_t.R2 = (sum(training_meta_article_md$sigma2)-sum(conflict_3a_t$sigma2))/sum(training_meta_article_md$sigma2)*100
conflict_3a_t.R2 = if_else(conflict_3a_t.R2<0, 0, conflict_3a_t.R2)
```
Exact model p values: `r conflict_3a_t$pval` | 
Exact Q test of moderators p value: `r conflict_3a_t$QMp` | 
Total I2: `r mlm.variance.distribution(conflict_3a_t)$totalI2`
```{r}
impact_3a_t = rma.mv(yi=yi, V=vi, data = training_md, method = "REML", random = ~1|ID/Comparison.Code, mods = training_md$Impact.Factor)
summary(impact_3a_t)
#pseudo R-squared
impact_3a_t.R2 = (sum(training_meta_article_md$sigma2)-sum(impact_3a_t$sigma2))/sum(training_meta_article_md$sigma2)*100
impact_3a_t.R2 = if_else(impact_3a_t.R2<0, 0, impact_3a_t.R2)
```
Exact model p values: `r impact_3a_t$pval` | 
Exact Q test of moderators p value: `r impact_3a_t$QMp` | 
Total I2: `r mlm.variance.distribution(impact_3a_t)$totalI2`
```{r}
citations_3a_t = rma.mv(yi=yi, V=vi, data = training_md, method = "REML", random = ~1|ID/Comparison.Code, mods = training_md$Citations.per.year)
summary(citations_3a_t)
#pseudo R-squared
citations_3a_t.R2 = (sum(training_meta_article_md$sigma2)-sum(citations_3a_t$sigma2))/sum(training_meta_article_md$sigma2)*100
citations_3a_t.R2 = if_else(citations_3a_t.R2<0, 0, citations_3a_t.R2)
```
Exact model p values: `r citations_3a_t$pval` | 
Exact Q test of moderators p value: `r citations_3a_t$QMp` | 
Total I2: `r mlm.variance.distribution(citations_3a_t)$totalI2`
```{r}
training_md %>% count(Region.of.origin)
region_3a_t = rma.mv(yi=yi, V=vi, data = training_md, method = "REML", random = ~1|ID/Comparison.Code, mods = ~relevel(factor(training_md$Region.of.origin), ref="North America"))
summary(region_3a_t)
#pseudo R-squared
region_3a_t.R2 = (sum(training_meta_article_md$sigma2)-sum(region_3a_t$sigma2))/sum(training_meta_article_md$sigma2)*100
region_3a_t.R2 = if_else(region_3a_t.R2<0, 0, region_3a_t.R2)
```
Exact model p values: `r region_3a_t$pval` | 
Exact Q test of moderators p value: `r region_3a_t$QMp` | 
Total I2: `r mlm.variance.distribution(region_3a_t)$totalI2`

Reactivation dataset:
(Suppl Table 18)
```{r}
reactivation_md %>% count(Intervention.Session)
session_3a_r = rma.mv(yi=yi, V=vi, data = reactivation_md, method = "REML", random = ~1|ID/Comparison.Code, mods = ~relevel(factor(reactivation_md$Intervention.Session), ref="Reactivation"))
summary(session_3a_r)
#pseudo R-squared
session_3a_r.R2 = (sum(reactivation_meta_article_md$sigma2)-sum(session_3a_r$sigma2))/sum(reactivation_meta_article_md$sigma2)*100
session_3a_r.R2 = if_else(session_3a_r.R2<0, 0, session_3a_r.R2)
```
Exact model p values: `r session_3a_r$pval` | 
Exact Q test of moderators p value: `r session_3a_r$QMp` | 
Total I2: `r mlm.variance.distribution(session_3a_r)$totalI2`
```{r}
reactivation_md %>% count(ConditioningType)
conditioning_3a_r = rma.mv(yi=yi, V=vi, data = reactivation_md, method = "REML", random = ~1|ID/Comparison.Code, mods = ~factor(reactivation_md$ConditioningType))
summary(conditioning_3a_r)
#pseudo R-squared
conditioning_3a_r.R2 = (sum(reactivation_meta_article_md$sigma2)-sum(conditioning_3a_r$sigma2))/sum(reactivation_meta_article_md$sigma2)*100
conditioning_3a_r.R2 = if_else(conditioning_3a_r.R2<0, 0, conditioning_3a_r.R2)
```
Exact model p values: `r conditioning_3a_r$pval` | 
Exact Q test of moderators p value: `r conditioning_3a_r$QMp` | 
Total I2: `r mlm.variance.distribution(conditioning_3a_r)$totalI2`
```{r}
reactivation_md %>% count(Habituation)
habituation_3a_r = rma.mv(yi=yi, V=vi, data = reactivation_md, method = "REML", random = ~1|ID/Comparison.Code, mods = ~factor(reactivation_md$Habituation))
summary(habituation_3a_r)
#pseudo R-squared
habituation_3a_r.R2 = (sum(reactivation_meta_article_md$sigma2)-sum(habituation_3a_r$sigma2))/sum(reactivation_meta_article_md$sigma2)*100
habituation_3a_r.R2 = if_else(habituation_3a_r.R2<0, 0, habituation_3a_r.R2)
```
Exact model p values: `r habituation_3a_r$pval` | 
Exact Q test of moderators p value: `r habituation_3a_r$QMp` | 
Total I2: `r mlm.variance.distribution(habituation_3a_r)$totalI2`
```{r}
reactivation_md %>% count(Handling)
handling_3a_r = rma.mv(yi=yi, V=vi, data = reactivation_md, method = "REML", random = ~1|ID/Comparison.Code, mods = ~factor(reactivation_md$Handling))
summary(handling_3a_r)
#pseudo R-squared
handling_3a_r.R2 = (sum(reactivation_meta_article_md$sigma2)-sum(handling_3a_r$sigma2))/sum(reactivation_meta_article_md$sigma2)*100
handling_3a_r.R2 = if_else(handling_3a_r.R2<0, 0, handling_3a_r.R2)
```
Exact model p values: `r handling_3a_r$pval` | 
Exact Q test of moderators p value: `r handling_3a_r$QMp` | 
Total I2: `r mlm.variance.distribution(handling_3a_r)$totalI2`
```{r}
shocks_3a_r = rma.mv(yi=yi, V=vi, data = reactivation_md, method = "REML", random = ~1|ID/Comparison.Code, mods = reactivation_md$NumberShock)
summary(shocks_3a_r)
#pseudo R-squared
shocks_3a_r.R2 = (sum(reactivation_meta_article_md$sigma2)-sum(shocks_3a_r$sigma2))/sum(reactivation_meta_article_md$sigma2)*100
shocks_3a_r.R2 = if_else(shocks_3a_r.R2<0, 0, shocks_3a_r.R2)
```
Exact model p values: `r shocks_3a_r$pval` | 
Exact Q test of moderators p value: `r shocks_3a_r$QMp` | 
Total I2: `r mlm.variance.distribution(shocks_3a_r)$totalI2`
```{r}
intensity_3a_r = rma.mv(yi=yi, V=vi, data = reactivation_md, method = "REML", random = ~1|ID/Comparison.Code, mods = reactivation_md$ShockIntensity)
summary(intensity_3a_r)
#pseudo R-squared
intensity_3a_r.R2 = (sum(reactivation_meta_article_md$sigma2)-sum(intensity_3a_r$sigma2))/sum(reactivation_meta_article_md$sigma2)*100
intensity_3a_r.R2 = if_else(intensity_3a_r.R2<0, 0, intensity_3a_r.R2)
```
Exact model p values: `r intensity_3a_r$pval` | 
Exact Q test of moderators p value: `r intensity_3a_r$QMp` | 
Total I2: `r mlm.variance.distribution(intensity_3a_r)$totalI2`
```{r}
time_intervention_3a_r = rma.mv(yi=yi, V=vi, data = reactivation_md, method = "REML", random = ~1|ID/Comparison.Code, mods = reactivation_md$InterventionTime)
summary(time_intervention_3a_r)
#pseudo R-squared
time_intervention_3a_r.R2 = (sum(reactivation_meta_article_md$sigma2)-sum(time_intervention_3a_r$sigma2))/sum(reactivation_meta_article_md$sigma2)*100
time_intervention_3a_r.R2 = if_else(time_intervention_3a_r.R2<0, 0, time_intervention_3a_r.R2)
```
Exact model p values: `r time_intervention_3a_r$pval` | 
Exact Q test of moderators p value: `r time_intervention_3a_r$QMp` | 
Total I2: `r mlm.variance.distribution(time_intervention_3a_r)$totalI2`
```{r}
time_reexposure_3a_r = rma.mv(yi=yi, V=vi, data = reactivation_md, method = "REML", random = ~1|ID/Comparison.Code, mods = reactivation_md$Time.to.reexposure)
summary(time_reexposure_3a_r)
#pseudo R-squared
time_reexposure_3a_r.R2 = (sum(reactivation_meta_article_md$sigma2)-sum(time_reexposure_3a_r$sigma2))/sum(reactivation_meta_article_md$sigma2)*100
time_reexposure_3a_r.R2 = if_else(time_reexposure_3a_r.R2<0, 0, time_reexposure_3a_r.R2)
```
Exact model p values: `r time_reexposure_3a_r$pval` | 
Exact Q test of moderators p value: `r time_reexposure_3a_r$QMp` | 
Total I2: `r mlm.variance.distribution(time_reexposure_3a_r)$totalI2`
```{r}
duration_3a_r = rma.mv(yi=yi, V=vi, data = reactivation_md, method = "REML", random = ~1|ID/Comparison.Code, mods = reactivation_md$duration_zscore)
summary(duration_3a_r)
#pseudo R-squared
duration_3a_r.R2 = (sum(reactivation_meta_article_md$sigma2)-sum(duration_3a_r$sigma2))/sum(reactivation_meta_article_md$sigma2)*100
duration_3a_r.R2 = if_else(duration_3a_r.R2<0, 0, duration_3a_r.R2)
```
Exact model p values: `r duration_3a_r$pval` | 
Exact Q test of moderators p value: `r duration_3a_r$QMp` | 
Total I2: `r mlm.variance.distribution(duration_3a_r)$totalI2`
```{r}
time_test_3a_r = rma.mv(yi=yi, V=vi, data = reactivation_md, method = "REML", random = ~1|ID/Comparison.Code, mods = reactivation_md$TimetoTest)
summary(time_test_3a_r)
#pseudo R-squared
time_test_3a_r.R2 = (sum(reactivation_meta_article_md$sigma2)-sum(time_test_3a_r$sigma2))/sum(reactivation_meta_article_md$sigma2)*100
time_test_3a_r.R2 = if_else(time_test_3a_r.R2<0, 0, time_test_3a_r.R2)
```
Exact model p values: `r time_test_3a_r$pval` | 
Exact Q test of moderators p value: `r time_test_3a_r$QMp` | 
Total I2: `r mlm.variance.distribution(time_test_3a_r)$totalI2`
```{r}
reactivation_md %>% count(Species)
species_3a_r = rma.mv(yi=yi, V=vi, data = reactivation_md, method = "REML", random = ~1|ID/Comparison.Code, mods = ~factor(reactivation_md$Species))
summary(species_3a_r)
#pseudo R-squared
species_3a_r.R2 = (sum(reactivation_meta_article_md$sigma2)-sum(species_3a_r$sigma2))/sum(reactivation_meta_article_md$sigma2)*100
species_3a_r.R2 = if_else(species_3a_r.R2<0, 0, species_3a_r.R2)
```
Exact model p values: `r species_3a_r$pval` | 
Exact Q test of moderators p value: `r species_3a_r$QMp` | 
Total I2: `r mlm.variance.distribution(species_3a_r)$totalI2`
```{r}
reactivation_md %>% count(Sex)
sex_3a_r = rma.mv(yi=yi, V=vi, data = reactivation_md, method = "REML", random = ~1|ID/Comparison.Code, mods = ~factor(reactivation_md$Sex))
summary(sex_3a_r)
#pseudo R-squared
sex_3a_r.R2 = (sum(reactivation_meta_article_md$sigma2)-sum(sex_3a_r$sigma2))/sum(reactivation_meta_article_md$sigma2)*100
sex_3a_r.R2 = if_else(sex_3a_r.R2<0, 0, sex_3a_r.R2)
```
Exact model p values: `r sex_3a_r$pval` | 
Exact Q test of moderators p value: `r sex_3a_r$QMp` | 
Total I2: `r mlm.variance.distribution(sex_3a_r)$totalI2`
```{r}
age_3a_r = rma.mv(yi=yi, V=vi, data = reactivation_md, method = "REML", random = ~1|ID/Comparison.Code, mods = reactivation_md$Age_mean)
summary(age_3a_r)
#pseudo R-squared
age_3a_r.R2 = (sum(reactivation_meta_article_md$sigma2)-sum(age_3a_r$sigma2))/sum(reactivation_meta_article_md$sigma2)*100
age_3a_r.R2 = if_else(age_3a_r.R2<0, 0, age_3a_r.R2)
```
Exact model p values: `r age_3a_r$pval` | 
Exact Q test of moderators p value: `r age_3a_r$QMp` | 
Total I2: `r mlm.variance.distribution(age_3a_r)$totalI2`
```{r}
housing_3a_r = rma.mv(yi=yi, V=vi, data = reactivation_md, method = "REML", random = ~1|ID/Comparison.Code, mods = reactivation_md$Housing)
summary(housing_3a_r)
#pseudo R-squared
housing_3a_r.R2 = (sum(reactivation_meta_article_md$sigma2)-sum(housing_3a_r$sigma2))/sum(reactivation_meta_article_md$sigma2)*100
housing_3a_r.R2 = if_else(housing_3a_r.R2<0, 0, housing_3a_r.R2)
```
Exact model p values: `r housing_3a_r$pval` | 
Exact Q test of moderators p value: `r housing_3a_r$QMp` | 
Total I2: `r mlm.variance.distribution(housing_3a_r)$totalI2`
```{r}
reactivation_md %>% count(Active.Drug)
drug_3a_r = rma.mv(yi=yi, V=vi, data = reactivation_md, method = "REML", random = ~1|ID/Comparison.Code, mods = ~relevel(factor(reactivation_md$Active.Drug), ref = "Anisomycin"))
summary(drug_3a_r)
#pseudo R-squared
drug_3a_r.R2 = (sum(reactivation_meta_article_md$sigma2)-sum(drug_3a_r$sigma2))/sum(reactivation_meta_article_md$sigma2)*100
drug_3a_r.R2 = if_else(drug_3a_r.R2<0, 0, drug_3a_r.R2)
```
Exact model p values: `r drug_3a_r$pval` | 
Exact Q test of moderators p value: `r drug_3a_r$QMp` | 
Total I2: `r mlm.variance.distribution(drug_3a_r)$totalI2`
```{r}
reactivation_md %>% count(Site.injection.other)
sites_3a_r = rma.mv(yi=yi, V=vi, data = reactivation_md, method = "REML", random = ~1|ID/Comparison.Code, mods = ~relevel(factor(reactivation_md$Site.injection.other), ref = "Systemic"))
summary(sites_3a_r)
#pseudo R-squared
sites_3a_r.R2 = (sum(reactivation_meta_article_md$sigma2)-sum(sites_3a_r$sigma2))/sum(reactivation_meta_article_md$sigma2)*100
sites_3a_r.R2 = if_else(sites_3a_r.R2<0, 0, sites_3a_r.R2)
```
Exact model p values: `r sites_3a_r$pval` | 
Exact Q test of moderators p value: `r sites_3a_r$QMp` | 
Total I2: `r mlm.variance.distribution(sites_3a_r)$totalI2`
```{r}
#dose_3a_r = rma.mv(yi=yi, V=vi, data = reactivation_md, method = "REML", random = ~1|ID/Comparison.Code, mods = reactivation_md$dose_zscore)
#summary(dose_3a_r)
#pseudo R-squared
#dose_3a_r.R2 = (sum(reactivation_meta_article_md$sigma2)-sum(dose_3a_r$sigma2))/sum(reactivation_meta_article_md$sigma2)*100
#dose_3a_r.R2 = if_else(dose_3a_r.R2<0, 0, dose_3a_r.R2)
```

```{r}
dose_3a_r_s = rma.mv(yi=yi, V=vi, data = reactivation_md, method = "REML", random = ~1|ID/Comparison.Code, mods = reactivation_md$dose_zscore_s)
summary(dose_3a_r_s)
#pseudo R-squared
dose_3a_r.R2_s = (sum(reactivation_meta_article_md$sigma2)-sum(dose_3a_r_s$sigma2))/sum(reactivation_meta_article_md$sigma2)*100
dose_3a_r.R2_s = if_else(dose_3a_r.R2_s<0, 0, dose_3a_r.R2_s)
```
Exact model p values: `r dose_3a_r_s$pval` | 
Exact Q test of moderators p value: `r dose_3a_r_s$QMp` | 
Total I2: `r mlm.variance.distribution(dose_3a_r_s)$totalI2`
```{r}
reactivation_md %>% count(Randomization)
randomization_3a_r = rma.mv(yi=yi, V=vi, data = reactivation_md, method = "REML", random = ~1|ID/Comparison.Code, mods = ~factor(reactivation_md$Randomization))
summary(randomization_3a_r)
#pseudo R-squared
randomization_3a_r.R2 = (sum(reactivation_meta_article_md$sigma2)-sum(randomization_3a_r$sigma2))/sum(reactivation_meta_article_md$sigma2)*100
randomization_3a_r.R2 = if_else(randomization_3a_r.R2<0, 0, randomization_3a_r.R2)
```
Exact model p values: `r randomization_3a_r$pval` | 
Exact Q test of moderators p value: `r randomization_3a_r$QMp` | 
Total I2: `r mlm.variance.distribution(randomization_3a_r)$totalI2`
```{r}
reactivation_md %>% count(Blinding)
blinding_3a_r = rma.mv(yi=yi, V=vi, data = reactivation_md, method = "REML", random = ~1|ID/Comparison.Code, mods = ~factor(reactivation_md$Blinding))
summary(blinding_3a_r)
#pseudo R-squared
blinding_3a_r.R2 = (sum(reactivation_meta_article_md$sigma2)-sum(blinding_3a_r$sigma2))/sum(reactivation_meta_article_md$sigma2)*100
blinding_3a_r.R2 = if_else(blinding_3a_r.R2<0, 0, blinding_3a_r.R2)
```
Exact model p values: `r blinding_3a_r$pval` | 
Exact Q test of moderators p value: `r blinding_3a_r$QMp` | 
Total I2: `r mlm.variance.distribution(blinding_3a_r)$totalI2`
```{r}
reactivation_md %>% count(SampleSizeCalc)
samplesize_3a_r = rma.mv(yi=yi, V=vi, data = reactivation_md, method = "REML", random = ~1|ID/Comparison.Code, mods = ~factor(reactivation_md$SampleSizeCalc))
summary(samplesize_3a_r)
#pseudo R-squared
samplesize_3a_r.R2 = (sum(reactivation_meta_article_md$sigma2)-sum(samplesize_3a_r$sigma2))/sum(reactivation_meta_article_md$sigma2)*100
samplesize_3a_r.R2 = if_else(samplesize_3a_r.R2<0, 0, samplesize_3a_r.R2)
```
Exact model p values: `r samplesize_3a_r$pval` | 
Exact Q test of moderators p value: `r samplesize_3a_r$QMp` | 
Total I2: `r mlm.variance.distribution(samplesize_3a_r)$totalI2`
```{r}
reactivation_md %>% count(RegulatoryReq)
requirements_3a_r = rma.mv(yi=yi, V=vi, data = reactivation_md, method = "REML", random = ~1|ID/Comparison.Code, mods = ~factor(reactivation_md$RegulatoryReq))
summary(requirements_3a_r)
#pseudo R-squared
requirements_3a_r.R2 = (sum(reactivation_meta_article_md$sigma2)-sum(requirements_3a_r$sigma2))/sum(reactivation_meta_article_md$sigma2)*100
requirements_3a_r.R2 = if_else(requirements_3a_r.R2<0, 0, requirements_3a_r.R2)
```
Exact model p values: `r requirements_3a_r$pval` | 
Exact Q test of moderators p value: `r requirements_3a_r$QMp` | 
Total I2: `r mlm.variance.distribution(requirements_3a_r)$totalI2`
```{r}
reactivation_md %>% count(ConflictInterest)
conflict_3a_r = rma.mv(yi=yi, V=vi, data = reactivation_md, method = "REML", random = ~1|ID/Comparison.Code, mods = ~relevel(factor(reactivation_md$ConflictInterest), ref = "Not informed"))
summary(conflict_3a_r)
#pseudo R-squared
conflict_3a_r.R2 = (sum(reactivation_meta_article_md$sigma2)-sum(conflict_3a_r$sigma2))/sum(reactivation_meta_article_md$sigma2)*100
conflict_3a_r.R2 = if_else(conflict_3a_r.R2<0, 0, conflict_3a_r.R2)
```
Exact model p values: `r conflict_3a_r$pval` | 
Exact Q test of moderators p value: `r conflict_3a_r$QMp` | 
Total I2: `r mlm.variance.distribution(conflict_3a_r)$totalI2`
```{r}
impact_3a_r = rma.mv(yi=yi, V=vi, data = reactivation_md, method = "REML", random = ~1|ID/Comparison.Code, mods = reactivation_md$Impact.Factor)
summary(impact_3a_r)
#pseudo R-squared
impact_3a_r.R2 = (sum(reactivation_meta_article_md$sigma2)-sum(impact_3a_r$sigma2))/sum(reactivation_meta_article_md$sigma2)*100
impact_3a_r.R2 = if_else(impact_3a_r.R2<0, 0, impact_3a_r.R2)
```
Exact model p values: `r impact_3a_r$pval` | 
Exact Q test of moderators p value: `r impact_3a_r$QMp` | 
Total I2: `r mlm.variance.distribution(impact_3a_r)$totalI2`
```{r}
citations_3a_r = rma.mv(yi=yi, V=vi, data = reactivation_md, method = "REML", random = ~1|ID/Comparison.Code, mods = reactivation_md$Citations.per.year)
summary(citations_3a_r)
#pseudo R-squared
citations_3a_r.R2 = (sum(reactivation_meta_article_md$sigma2)-sum(citations_3a_r$sigma2))/sum(reactivation_meta_article_md$sigma2)*100
citations_3a_r.R2 = if_else(citations_3a_r.R2<0, 0, citations_3a_r.R2)
```
Exact model p values: `r citations_3a_r$pval` | 
Exact Q test of moderators p value: `r citations_3a_r$QMp` | 
Total I2: `r mlm.variance.distribution(citations_3a_r)$totalI2`
```{r}
reactivation_md %>% count(Region.of.origin)
region_3a_r = rma.mv(yi=yi, V=vi, data = reactivation_md, method = "REML", random = ~1|ID/Comparison.Code, mods = ~relevel(factor(reactivation_md$Region.of.origin), ref="North America"))
summary(region_3a_r)
#pseudo R-squared
region_3a_r.R2 = (sum(reactivation_meta_article_md$sigma2)-sum(region_3a_r$sigma2))/sum(reactivation_meta_article_md$sigma2)*100
region_3a_r.R2 = if_else(region_3a_r.R2<0, 0, region_3a_r.R2)
```
Exact model p values: `r region_3a_r$pval` | 
Exact Q test of moderators p value: `r region_3a_r$QMp` | 
Total I2: `r mlm.variance.distribution(region_3a_r)$totalI2`

Fig 3 B:
```{r}
dataset = c('Complete, by article')
moderadores = c('Conditioning Type', 'Habituation reported', 'Handling reported', 'Number of shocks', 'Shock intensity', 'Intervention time', 'Time to reexposure', 'Reexposure duration', 'Time to test', 'Species', 'Sex', 'Mean age', 'Housing', 'Active drug', 'Injection site', 'Dose', 'Randomization', 'Blinding', 'Sample size calculation', 'Regulatory requirements', 'Conflict of interest statement', 'Impact factor', 'Citations per year', 'Region of origin')
rquadrado = c(conditioning_3a.R2, habituation_3a.R2, handling_3a.R2, shocks_3a.R2, intensity_3a.R2, time_intervention_3a.R2, NA, NA, time_test_3a.R2, species_3a.R2, sex_3a.R2, age_3a.R2, housing_3a.R2, drug_3a.R2, sites_3a.R2, dose_3a.R2_s, randomization_3a.R2, blinding_3a.R2, samplesize_3a.R2, requirements_3a.R2, conflict_3a.R2, impact_3a.R2, citations_3a.R2, region_3a.R2)
pvalor = c(conditioning_3a$QMp, habituation_3a$QMp, handling_3a$QMp, shocks_3a$QMp, intensity_3a$QMp, time_intervention_3a$QMp, NA, NA, time_test_3a$QMp, species_3a$QMp, sex_3a$QMp, age_3a$QMp, housing_3a$QMp, drug_3a$QMp, sites_3a$QMp, dose_3a_s$QMp, randomization_3a$QMp, blinding_3a$QMp, samplesize_3a$QMp, requirements_3a$QMp, conflict_3a$QMp, impact_3a$QMp, citations_3a$QMp, region_3a$QMp)

moderators_3a = data.frame(dataset, moderadores, rquadrado, pvalor)
moderators_3a = moderators_3a %>% mutate(pval_cat = if_else(pvalor>0.05, ">0.05", if_else(pvalor<=0.05&pvalor>0.01, "<0.05", if_else(pvalor<=0.01&pvalor>0.001, "<0.01", "<0.001"))))
```
```{r}
dataset = c('Training, by article')
rquadrado = c(conditioning_3a_t.R2, habituation_3a_t.R2, handling_3a_t.R2, shocks_3a_t.R2, intensity_3a_t.R2, time_intervention_3a_t.R2, NA, NA, time_test_3a_t.R2, species_3a_t.R2, sex_3a_t.R2, age_3a_t.R2, housing_3a_t.R2, drug_3a_t.R2, sites_3a_t.R2, dose_3a_t.R2_s, randomization_3a_t.R2, blinding_3a_t.R2, NA, requirements_3a_t.R2, conflict_3a_t.R2, impact_3a_t.R2, citations_3a_t.R2, region_3a_t.R2)
pvalor = c(conditioning_3a_t$QMp, habituation_3a_t$QMp, handling_3a_t$QMp, shocks_3a_t$QMp, intensity_3a_t$QMp, time_intervention_3a_t$QMp, NA, NA, time_test_3a_t$QMp, species_3a_t$QMp, sex_3a_t$QMp, age_3a_t$QMp, housing_3a_t$QMp, drug_3a_t$QMp, sites_3a_t$QMp, dose_3a_t_s$QMp, randomization_3a_t$QMp, blinding_3a_t$QMp, NA, requirements_3a_t$QMp, conflict_3a_t$QMp, impact_3a_t$QMp, citations_3a_t$QMp, region_3a_t$QMp)

moderators_3a_t = data.frame(dataset, moderadores, rquadrado, pvalor)
moderators_3a_t = moderators_3a_t %>% mutate(pval_cat = if_else(pvalor>0.05, ">0.05", if_else(pvalor<=0.05&pvalor>0.01, "<0.05", if_else(pvalor<=0.01&pvalor>0.001, "<0.01", "<0.001"))))
```
```{r}
dataset = c('Reactivation, by article')
rquadrado = c(conditioning_3a_r.R2, habituation_3a_r.R2, handling_3a_r.R2, shocks_3a_r.R2, intensity_3a_r.R2, time_intervention_3a_r.R2, time_reexposure_3a_r.R2, duration_3a_r.R2, time_test_3a_r.R2, species_3a_r.R2, sex_3a_r.R2, age_3a_r.R2, housing_3a_r.R2, drug_3a_r.R2, sites_3a_r.R2, dose_3a.R2_s, randomization_3a_r.R2, blinding_3a_r.R2, samplesize_3a_r.R2, requirements_3a_r.R2, conflict_3a_r.R2, impact_3a_r.R2, citations_3a_r.R2, region_3a_r.R2)
pvalor = c(conditioning_3a_r$QMp, habituation_3a_r$QMp, handling_3a_r$QMp, shocks_3a_r$QMp, intensity_3a_r$QMp, time_intervention_3a_r$QMp, time_reexposure_3a_r$QMp, duration_3a_r$QMp, time_test_3a_r$QMp, species_3a_r$QMp, sex_3a_r$QMp, age_3a_r$QMp, housing_3a_r$QMp, drug_3a_r$QMp, sites_3a_r$QMp, dose_3a_r_s$QMp, randomization_3a_r$QMp, blinding_3a_r$QMp, samplesize_3a_r$QMp, requirements_3a_r$QMp, conflict_3a_r$QMp, impact_3a_r$QMp, citations_3a_r$QMp, region_3a_r$QMp)

moderators_3a_r = data.frame(dataset, moderadores, rquadrado, pvalor)
moderators_3a_r = moderators_3a_r %>% mutate(pval_cat = if_else(pvalor>0.05, ">0.05", if_else(pvalor<=0.05&pvalor>0.01, "<0.05", if_else(pvalor<=0.01&pvalor>0.001, "<0.01", "<0.001"))))
```
```{r}
all_moderators_3a = bind_rows(moderators_3a, moderators_3a_t, moderators_3a_r)
all_moderators_3a$moderadores = factor(all_moderators_3a$moderadores, levels = rev(factor(moderadores)))
all_moderators_3a$dataset = factor(all_moderators_3a$dataset, levels = c("Complete, by article", "Training, by article", "Reactivation, by article"))
labels_3a = c("Complete,\nby article","Training,\nby article", "Reactivation,\nby article")

all_mods_3a = ggplot(all_moderators_3a, aes(x=dataset, y=moderadores, fill=pval_cat)) + 
  geom_tile() + 
  scale_fill_manual(values=c("#3182bd","#6baed6","#bdd7e7","#eff3ff"), na.value = "white", name = "Q test for moderators \np value") + 
  geom_text(aes(x=dataset, y=moderadores, label = if_else(condition = rquadrado>=0, true = paste(round(rquadrado, 1), "%"), false = " ")), color = "black", size = 6) + 
  scale_x_discrete(position = "top", labels=labels_3a) + 
  theme(axis.title.x = element_blank(), axis.text.x.top = element_text(angle = 0, size = 14), axis.title.y = element_blank(), panel.background = element_blank(),  axis.ticks = element_blank(), legend.position = "bottom", axis.text = element_text(size=15), legend.text = element_text(size=12), legend.title = element_text(size=12))
all_mods_3a
```

**Univariate 3-level meta-regressions (research group)**

Complete dataset:
(Suppl Table 19)
```{r}
full_data_md %>% count(Intervention.Session)
session_3g = rma.mv(yi=yi, V=vi, data = full_data_md, method = "REML", random = ~1|group/Comparison.Code, mods = ~relevel(factor(full_data_md$Intervention.Session), ref="Training"))
summary(session_3g)
#pseudo R-squared
session_3g.R2 = (sum(full_data_meta_article_md$sigma2)-sum(session_3g$sigma2))/sum(full_data_meta_article_md$sigma2)*100
session_3g.R2 = if_else(session_3g.R2<0, 0, session_3g.R2)
```
Exact model p values: `r session_3g$pval` | 
Exact Q test of moderators p value: `r session_3g$QMp` | 
Total I2: `r mlm.variance.distribution(session_3g)$totalI2`
```{r}
full_data_md %>% count(ConditioningType)
conditioning_3g = rma.mv(yi=yi, V=vi, data = full_data_md, method = "REML", random = ~1|group/Comparison.Code, mods = ~factor(full_data_md$ConditioningType))
summary(conditioning_3g)
#pseudo R-squared
conditioning_3g.R2 = (sum(full_data_meta_article_md$sigma2)-sum(conditioning_3g$sigma2))/sum(full_data_meta_article_md$sigma2)*100
conditioning_3g.R2 = if_else(conditioning_3g.R2<0, 0, conditioning_3g.R2)
```
Exact model p values: `r conditioning_3g$pval` | 
Exact Q test of moderators p value: `r conditioning_3g$QMp` | 
Total I2: `r mlm.variance.distribution(conditioning_3g)$totalI2`
```{r}
full_data_md %>% count(Habituation)
habituation_3g = rma.mv(yi=yi, V=vi, data = full_data_md, method = "REML", random = ~1|group/Comparison.Code, mods = ~factor(full_data_md$Habituation))
summary(habituation_3g)
#pseudo R-squared
habituation_3g.R2 = (sum(full_data_meta_article_md$sigma2)-sum(habituation_3g$sigma2))/sum(full_data_meta_article_md$sigma2)*100
habituation_3g.R2 = if_else(habituation_3g.R2<0, 0, habituation_3g.R2)
```
Exact model p values: `r habituation_3g$pval` | 
Exact Q test of moderators p value: `r habituation_3g$QMp` | 
Total I2: `r mlm.variance.distribution(habituation_3g)$totalI2`
```{r}
full_data_md %>% count(Handling)
handling_3g = rma.mv(yi=yi, V=vi, data = full_data_md, method = "REML", random = ~1|group/Comparison.Code, mods = ~factor(full_data_md$Handling))
summary(handling_3g)
#pseudo R-squared
handling_3g.R2 = (sum(full_data_meta_article_md$sigma2)-sum(handling_3g$sigma2))/sum(full_data_meta_article_md$sigma2)*100
handling_3g.R2 = if_else(handling_3g.R2<0, 0, handling_3g.R2)
```
Exact model p values: `r handling_3g$pval` | 
Exact Q test of moderators p value: `r handling_3g$QMp` | 
Total I2: `r mlm.variance.distribution(handling_3g)$totalI2`
```{r}
shocks_3g = rma.mv(yi=yi, V=vi, data = full_data_md, method = "REML", random = ~1|group/Comparison.Code, mods = full_data_md$NumberShock)
summary(shocks_3g)
#pseudo R-squared
shocks_3g.R2 = (sum(full_data_meta_article_md$sigma2)-sum(shocks_3g$sigma2))/sum(full_data_meta_article_md$sigma2)*100
shocks_3g.R2 = if_else(shocks_3g.R2<0, 0, shocks_3g.R2)
```
Exact model p values: `r shocks_3g$pval` | 
Exact Q test of moderators p value: `r shocks_3g$QMp` | 
Total I2: `r mlm.variance.distribution(shocks_3g)$totalI2`
```{r}
intensity_3g = rma.mv(yi=yi, V=vi, data = full_data_md, method = "REML", random = ~1|group/Comparison.Code, mods = full_data_md$ShockIntensity)
summary(intensity_3g)
#pseudo R-squared
intensity_3g.R2 = (sum(full_data_meta_article_md$sigma2)-sum(intensity_3g$sigma2))/sum(full_data_meta_article_md$sigma2)*100
intensity_3g.R2 = if_else(intensity_3g.R2<0, 0, intensity_3g.R2)
```
Exact model p values: `r intensity_3g$pval` | 
Exact Q test of moderators p value: `r intensity_3g$QMp` | 
Total I2: `r mlm.variance.distribution(intensity_3g)$totalI2`
```{r}
time_intervention_3g = rma.mv(yi=yi, V=vi, data = full_data_md, method = "REML", random = ~1|group/Comparison.Code, mods = full_data_md$InterventionTime)
summary(time_intervention_3g)
#pseudo R-squared
time_intervention_3g.R2 = (sum(full_data_meta_article_md$sigma2)-sum(time_intervention_3g$sigma2))/sum(full_data_meta_article_md$sigma2)*100
time_intervention_3g.R2 = if_else(time_intervention_3g.R2<0, 0, time_intervention_3g.R2)
```
Exact model p values: `r time_intervention_3g$pval` | 
Exact Q test of moderators p value: `r time_intervention_3g$QMp` | 
Total I2: `r mlm.variance.distribution(time_intervention_3g)$totalI2`
```{r}
time_test_3g = rma.mv(yi=yi, V=vi, data = full_data_md, method = "REML", random = ~1|group/Comparison.Code, mods = full_data_md$TimetoTest)
summary(time_test_3g)
#pseudo R-squared
time_test_3g.R2 = (sum(full_data_meta_article_md$sigma2)-sum(time_test_3g$sigma2))/sum(full_data_meta_article_md$sigma2)*100
time_test_3g.R2 = if_else(time_test_3g.R2<0, 0, time_test_3g.R2)
```
Exact model p values: `r time_test_3g$pval` | 
Exact Q test of moderators p value: `r time_test_3g$QMp` | 
Total I2: `r mlm.variance.distribution(time_test_3g)$totalI2`
```{r}
full_data_md %>% count(Species)
species_3g = rma.mv(yi=yi, V=vi, data = full_data_md, method = "REML", random = ~1|group/Comparison.Code, mods = ~factor(full_data_md$Species))
summary(species_3g)
#pseudo R-squared
species_3g.R2 = (sum(full_data_meta_article_md$sigma2)-sum(species_3g$sigma2))/sum(full_data_meta_article_md$sigma2)*100
species_3g.R2 = if_else(species_3g.R2<0, 0, species_3g.R2)
```
Exact model p values: `r species_3g$pval` | 
Exact Q test of moderators p value: `r species_3g$QMp` | 
Total I2: `r mlm.variance.distribution(species_3g)$totalI2`
```{r}
full_data_md %>% count(Sex)
sex_3g = rma.mv(yi=yi, V=vi, data = full_data_md, method = "REML", random = ~1|group/Comparison.Code, mods = ~factor(full_data_md$Sex))
summary(sex_3g)
#pseudo R-squared
sex_3g.R2 = (sum(full_data_meta_article_md$sigma2)-sum(sex_3g$sigma2))/sum(full_data_meta_article_md$sigma2)*100
sex_3g.R2 = if_else(sex_3g.R2<0, 0, sex_3g.R2)
```
Exact model p values: `r sex_3g$pval` | 
Exact Q test of moderators p value: `r sex_3g$QMp` | 
Total I2: `r mlm.variance.distribution(sex_3g)$totalI2`
```{r}
age_3g = rma.mv(yi=yi, V=vi, data = full_data_md, method = "REML", random = ~1|group/Comparison.Code, mods = full_data_md$Age_mean)
summary(age_3g)
#pseudo R-squared
age_3g.R2 = (sum(full_data_meta_article_md$sigma2)-sum(age_3g$sigma2))/sum(full_data_meta_article_md$sigma2)*100
age_3g.R2 = if_else(age_3g.R2<0, 0, age_3g.R2)
```
Exact model p values: `r age_3g$pval` | 
Exact Q test of moderators p value: `r age_3g$QMp` | 
Total I2: `r mlm.variance.distribution(age_3g)$totalI2`
```{r}
housing_3g = rma.mv(yi=yi, V=vi, data = full_data_md, method = "REML", random = ~1|group/Comparison.Code, mods = full_data_md$Housing)
summary(housing_3g)
#pseudo R-squared
housing_3g.R2 = (sum(full_data_meta_article_md$sigma2)-sum(housing_3g$sigma2))/sum(full_data_meta_article_md$sigma2)*100
housing_3g.R2 = if_else(housing_3g.R2<0, 0, housing_3g.R2)
```
Exact model p values: `r housing_3g$pval` | 
Exact Q test of moderators p value: `r housing_3g$QMp` | 
Total I2: `r mlm.variance.distribution(housing_3g)$totalI2`
```{r}
full_data_md %>% count(Active.Drug)
drug_3g = rma.mv(yi=yi, V=vi, data = full_data_md, method = "REML", random = ~1|group/Comparison.Code, mods = ~relevel(factor(full_data_md$Active.Drug), ref = "Anisomycin"))
summary(drug_3g)
#pseudo R-squared
drug_3g.R2 = (sum(full_data_meta_article_md$sigma2)-sum(drug_3g$sigma2))/sum(full_data_meta_article_md$sigma2)*100
drug_3g.R2 = if_else(drug_3g.R2<0, 0, drug_3g.R2)
```
Exact model p values: `r drug_3g$pval` | 
Exact Q test of moderators p value: `r drug_3g$QMp` | 
Total I2: `r mlm.variance.distribution(drug_3g)$totalI2`
```{r}
full_data_md %>% count(Site.injection.other)
sites_3g = rma.mv(yi=yi, V=vi, data = full_data_md, method = "REML", random = ~1|group/Comparison.Code, mods = ~relevel(factor(full_data_md$Site.injection.other), ref = "Systemic"))
summary(sites_3g)
#pseudo R-squared
sites_3g.R2 = (sum(full_data_meta_article_md$sigma2)-sum(sites_3g$sigma2))/sum(full_data_meta_article_md$sigma2)*100
sites_3g.R2 = if_else(sites_3g.R2<0, 0, sites_3g.R2)
```
Exact model p values: `r sites_3g$pval` | 
Exact Q test of moderators p value: `r sites_3g$QMp` | 
Total I2: `r mlm.variance.distribution(sites_3g)$totalI2`
```{r}
#dose_3g = rma.mv(yi=yi, V=vi, data = full_data_md, method = "REML", random = ~1|group/Comparison.Code, mods = full_data_md$dose_zscore)
#summary(dose_3g)
#pseudo R-squared
#dose_3g.R2 = (sum(full_data_meta_article_md$sigma2)-sum(dose_3g$sigma2))/sum(full_data_meta_article_md$sigma2)*100
#dose_3g.R2 = if_else(dose_3g.R2<0, 0, dose_3g.R2)
```

```{r}
dose_3g_s = rma.mv(yi=yi, V=vi, data = full_data_md, method = "REML", random = ~1|group/Comparison.Code, mods = full_data_md$dose_zscore_s)
summary(dose_3g_s)
#pseudo R-squared
dose_3g.R2_s = (sum(full_data_meta_article_md$sigma2)-sum(dose_3g_s$sigma2))/sum(full_data_meta_article_md$sigma2)*100
dose_3g.R2_s = if_else(dose_3g.R2_s<0, 0, dose_3g.R2_s)
```
Exact model p values: `r dose_3g_s$pval` | 
Exact Q test of moderators p value: `r dose_3g_s$QMp` | 
Total I2: `r mlm.variance.distribution(dose_3g_s)$totalI2`

```{r}
full_data_md %>% count(Randomization)
randomization_3g = rma.mv(yi=yi, V=vi, data = full_data_md, method = "REML", random = ~1|group/Comparison.Code, mods = ~factor(full_data_md$Randomization))
summary(randomization_3g)
#pseudo R-squared
randomization_3g.R2 = (sum(full_data_meta_article_md$sigma2)-sum(randomization_3g$sigma2))/sum(full_data_meta_article_md$sigma2)*100
randomization_3g.R2 = if_else(randomization_3g.R2<0, 0, randomization_3g.R2)
```
Exact model p values: `r randomization_3g$pval` | 
Exact Q test of moderators p value: `r randomization_3g$QMp` | 
Total I2: `r mlm.variance.distribution(randomization_3g)$totalI2`
```{r}
full_data_md %>% count(Blinding)
blinding_3g = rma.mv(yi=yi, V=vi, data = full_data_md, method = "REML", random = ~1|group/Comparison.Code, mods = ~factor(full_data_md$Blinding))
summary(blinding_3g)
#pseudo R-squared
blinding_3g.R2 = (sum(full_data_meta_article_md$sigma2)-sum(blinding_3g$sigma2))/sum(full_data_meta_article_md$sigma2)*100
blinding_3g.R2 = if_else(blinding_3g.R2<0, 0, blinding_3g.R2)
```
Exact model p values: `r blinding_3g$pval` | 
Exact Q test of moderators p value: `r blinding_3g$QMp` | 
Total I2: `r mlm.variance.distribution(blinding_3g)$totalI2`
```{r}
full_data_md %>% count(SampleSizeCalc)
samplesize_3g = rma.mv(yi=yi, V=vi, data = full_data_md, method = "REML", random = ~1|group/Comparison.Code, mods = ~factor(full_data_md$SampleSizeCalc))
summary(samplesize_3g)
#pseudo R-squared
samplesize_3g.R2 = (sum(full_data_meta_article_md$sigma2)-sum(samplesize_3g$sigma2))/sum(full_data_meta_article_md$sigma2)*100
samplesize_3g.R2 = if_else(samplesize_3g.R2<0, 0, samplesize_3g.R2)
```
Exact model p values: `r samplesize_3g$pval` | 
Exact Q test of moderators p value: `r samplesize_3g$QMp` | 
Total I2: `r mlm.variance.distribution(samplesize_3g)$totalI2`
```{r}
full_data_md %>% count(RegulatoryReq)
requirements_3g = rma.mv(yi=yi, V=vi, data = full_data_md, method = "REML", random = ~1|group/Comparison.Code, mods = ~factor(full_data_md$RegulatoryReq))
summary(requirements_3g)
#pseudo R-squared
requirements_3g.R2 = (sum(full_data_meta_article_md$sigma2)-sum(requirements_3g$sigma2))/sum(full_data_meta_article_md$sigma2)*100
requirements_3g.R2 = if_else(requirements_3g.R2<0, 0, requirements_3g.R2)
```
Exact model p values: `r requirements_3g$pval` | 
Exact Q test of moderators p value: `r requirements_3g$QMp` | 
Total I2: `r mlm.variance.distribution(requirements_3g)$totalI2`
```{r}
full_data_md %>% count(ConflictInterest)
conflict_3g = rma.mv(yi=yi, V=vi, data = full_data_md, method = "REML", random = ~1|group/Comparison.Code, mods = ~relevel(factor(full_data_md$ConflictInterest), ref = "Not informed"))
summary(conflict_3g)
#pseudo R-squared
conflict_3g.R2 = (sum(full_data_meta_article_md$sigma2)-sum(conflict_3g$sigma2))/sum(full_data_meta_article_md$sigma2)*100
conflict_3g.R2 = if_else(conflict_3g.R2<0, 0, conflict_3g.R2)
```
Exact model p values: `r conflict_3g$pval` | 
Exact Q test of moderators p value: `r conflict_3g$QMp` | 
Total I2: `r mlm.variance.distribution(conflict_3g)$totalI2`
```{r}
impact_3g = rma.mv(yi=yi, V=vi, data = full_data_md, method = "REML", random = ~1|group/Comparison.Code, mods = full_data_md$Impact.Factor)
summary(impact_3g)
#pseudo R-squared
impact_3g.R2 = (sum(full_data_meta_article_md$sigma2)-sum(impact_3g$sigma2))/sum(full_data_meta_article_md$sigma2)*100
impact_3g.R2 = if_else(impact_3g.R2<0, 0, impact_3g.R2)
```
Exact model p values: `r impact_3g$pval` | 
Exact Q test of moderators p value: `r impact_3g$QMp` | 
Total I2: `r mlm.variance.distribution(impact_3g)$totalI2`
```{r}
citations_3g = rma.mv(yi=yi, V=vi, data = full_data_md, method = "REML", random = ~1|group/Comparison.Code, mods = full_data_md$Citations.per.year)
summary(citations_3g)
#pseudo R-squared
citations_3g.R2 = (sum(full_data_meta_article_md$sigma2)-sum(citations_3g$sigma2))/sum(full_data_meta_article_md$sigma2)*100
citations_3g.R2 = if_else(citations_3g.R2<0, 0, citations_3g.R2)
```
Exact model p values: `r citations_3g$pval` | 
Exact Q test of moderators p value: `r citations_3g$QMp` | 
Total I2: `r mlm.variance.distribution(citations_3g)$totalI2`
```{r}
full_data_md %>% count(Region.of.origin)
region_3g = rma.mv(yi=yi, V=vi, data = full_data_md, method = "REML", random = ~1|group/Comparison.Code, mods = ~relevel(factor(full_data_md$Region.of.origin), ref="North America"))
summary(region_3g)
region_3g$pval
mlm.variance.distribution(region_3g)$totalI2
#pseudo R-squared
region_3g.R2 = (sum(full_data_meta_article_md$sigma2)-sum(region_3g$sigma2))/sum(full_data_meta_article_md$sigma2)*100
region_3g.R2 = if_else(region_3g.R2<0, 0, region_3g.R2)
```
Exact model p values: `r region_3g$pval` | 
Exact Q test of moderators p value: `r region_3g$QMp` | 
Total I2: `r mlm.variance.distribution(region_3g)$totalI2`

Training dataset:
(Suppl Table 20)
```{r}
training_md %>% count(ConditioningType)
conditioning_3g_t = rma.mv(yi=yi, V=vi, data = training_md, method = "REML", random = ~1|group/Comparison.Code, mods = ~factor(training_md$ConditioningType))
summary(conditioning_3g_t)
#pseudo R-squared
conditioning_3g_t.R2 = (sum(training_meta_article_md$sigma2)-sum(conditioning_3g_t$sigma2))/sum(training_meta_article_md$sigma2)*100
conditioning_3g_t.R2 = if_else(conditioning_3g_t.R2<0, 0, conditioning_3g_t.R2)
```
Exact model p values: `r conditioning_3g_t$pval` | 
Exact Q test of moderators p value: `r conditioning_3g_t$QMp` | 
Total I2: `r mlm.variance.distribution(conditioning_3g_t)$totalI2`
```{r}
training_md %>% count(Habituation)
habituation_3g_t = rma.mv(yi=yi, V=vi, data = training_md, method = "REML", random = ~1|group/Comparison.Code, mods = ~factor(training_md$Habituation))
summary(habituation_3g_t)
#pseudo R-squared
habituation_3g_t.R2 = (sum(training_meta_article_md$sigma2)-sum(habituation_3g_t$sigma2))/sum(training_meta_article_md$sigma2)*100
habituation_3g_t.R2 = if_else(habituation_3g_t.R2<0, 0, habituation_3g_t.R2)
```
Exact model p values: `r habituation_3g_t$pval` | 
Exact Q test of moderators p value: `r habituation_3g_t$QMp` | 
Total I2: `r mlm.variance.distribution(habituation_3g_t)$totalI2`
```{r}
training_md %>% count(Handling)
handling_3g_t = rma.mv(yi=yi, V=vi, data = training_md, method = "REML", random = ~1|group/Comparison.Code, mods = ~factor(training_md$Handling))
summary(handling_3g_t)
#pseudo R-squared
handling_3g_t.R2 = (sum(training_meta_article_md$sigma2)-sum(handling_3g_t$sigma2))/sum(training_meta_article_md$sigma2)*100
handling_3g_t.R2 = if_else(handling_3g_t.R2<0, 0, handling_3g_t.R2)
```
Exact model p values: `r handling_3g_t$pval` | 
Exact Q test of moderators p value: `r handling_3g_t$QMp` | 
Total I2: `r mlm.variance.distribution(handling_3g_t)$totalI2`
```{r}
shocks_3g_t = rma.mv(yi=yi, V=vi, data = training_md, method = "REML", random = ~1|group/Comparison.Code, mods = training_md$NumberShock)
summary(shocks_3g_t)
#pseudo R-squared
shocks_3g_t.R2 = (sum(training_meta_article_md$sigma2)-sum(shocks_3g_t$sigma2))/sum(training_meta_article_md$sigma2)*100
shocks_3g_t.R2 = if_else(shocks_3g_t.R2<0, 0, shocks_3g_t.R2)
```
Exact model p values: `r shocks_3g_t$pval` | 
Exact Q test of moderators p value: `r shocks_3g_t$QMp` | 
Total I2: `r mlm.variance.distribution(shocks_3g_t)$totalI2`
```{r}
intensity_3g_t = rma.mv(yi=yi, V=vi, data = training_md, method = "REML", random = ~1|group/Comparison.Code, mods = training_md$ShockIntensity)
summary(intensity_3g_t)
#pseudo R-squared
intensity_3g_t.R2 = (sum(training_meta_article_md$sigma2)-sum(intensity_3g_t$sigma2))/sum(training_meta_article_md$sigma2)*100
intensity_3g_t.R2 = if_else(intensity_3g_t.R2<0, 0, intensity_3g_t.R2)
```
Exact model p values: `r intensity_3g_t$pval` | 
Exact Q test of moderators p value: `r intensity_3g_t$QMp` | 
Total I2: `r mlm.variance.distribution(intensity_3g_t)$totalI2`
```{r}
time_intervention_3g_t = rma.mv(yi=yi, V=vi, data = training_md, method = "REML", random = ~1|group/Comparison.Code, mods = training_md$InterventionTime)
summary(time_intervention_3g_t)
#pseudo R-squared
time_intervention_3g_t.R2 = (sum(training_meta_article_md$sigma2)-sum(time_intervention_3g_t$sigma2))/sum(training_meta_article_md$sigma2)*100
time_intervention_3g_t.R2 = if_else(time_intervention_3g_t.R2<0, 0, time_intervention_3g_t.R2)
```
Exact model p values: `r time_intervention_3g_t$pval` | 
Exact Q test of moderators p value: `r time_intervention_3g_t$QMp` | 
Total I2: `r mlm.variance.distribution(time_intervention_3g_t)$totalI2`
```{r}
time_test_3g_t = rma.mv(yi=yi, V=vi, data = training_md, method = "REML", random = ~1|group/Comparison.Code, mods = training_md$TimetoTest)
summary(time_test_3g_t)
#pseudo R-squared
time_test_3g_t.R2 = (sum(training_meta_article_md$sigma2)-sum(time_test_3g_t$sigma2))/sum(training_meta_article_md$sigma2)*100
time_test_3g_t.R2 = if_else(time_test_3g_t.R2<0, 0, time_test_3g_t.R2)
```
Exact model p values: `r time_test_3g_t$pval` | 
Exact Q test of moderators p value: `r time_test_3g_t$QMp` | 
Total I2: `r mlm.variance.distribution(time_test_3g_t)$totalI2`
```{r}
training_md %>% count(Species)
species_3g_t = rma.mv(yi=yi, V=vi, data = training_md, method = "REML", random = ~1|group/Comparison.Code, mods = ~factor(training_md$Species))
summary(species_3g_t)
#pseudo R-squared
species_3g_t.R2 = (sum(training_meta_article_md$sigma2)-sum(species_3g_t$sigma2))/sum(training_meta_article_md$sigma2)*100
species_3g_t.R2 = if_else(species_3g_t.R2<0, 0, species_3g_t.R2)
```
Exact model p values: `r species_3g_t$pval` | 
Exact Q test of moderators p value: `r species_3g_t$QMp` | 
Total I2: `r mlm.variance.distribution(species_3g_t)$totalI2`
```{r}
training_md %>% count(Sex)
sex_3g_t = rma.mv(yi=yi, V=vi, data = training_md, method = "REML", random = ~1|group/Comparison.Code, mods = ~factor(training_md$Sex))
summary(sex_3g_t)
#pseudo R-squared
sex_3g_t.R2 = (sum(training_meta_article_md$sigma2)-sum(sex_3g_t$sigma2))/sum(training_meta_article_md$sigma2)*100
sex_3g_t.R2 = if_else(sex_3g_t.R2<0, 0, sex_3g_t.R2)
```
Exact model p values: `r sex_3g_t$pval` | 
Exact Q test of moderators p value: `r sex_3g_t$QMp` | 
Total I2: `r mlm.variance.distribution(sex_3g_t)$totalI2`
```{r}
age_3g_t = rma.mv(yi=yi, V=vi, data = training_md, method = "REML", random = ~1|group/Comparison.Code, mods = training_md$Age_mean)
summary(age_3g_t)
mlm.variance.distribution(age_3g_t)$totalI2
#pseudo R-squared
age_3g_t.R2 = (sum(training_meta_article_md$sigma2)-sum(age_3g_t$sigma2))/sum(training_meta_article_md$sigma2)*100
age_3g_t.R2 = if_else(age_3g_t.R2<0, 0, age_3g_t.R2)
```
Exact model p values: `r age_3g_t$pval` | 
Exact Q test of moderators p value: `r age_3g_t$QMp` | 
Total I2: `r mlm.variance.distribution(age_3g_t)$totalI2`
```{r}
housing_3g_t = rma.mv(yi=yi, V=vi, data = training_md, method = "REML", random = ~1|group/Comparison.Code, mods = training_md$Housing)
summary(housing_3g_t)
#pseudo R-squared
housing_3g_t.R2 = (sum(training_meta_article_md$sigma2)-sum(housing_3g_t$sigma2))/sum(training_meta_article_md$sigma2)*100
housing_3g_t.R2 = if_else(housing_3g_t.R2<0, 0, housing_3g_t.R2)
```
Exact model p values: `r housing_3g_t$pval` | 
Exact Q test of moderators p value: `r housing_3g_t$QMp` | 
Total I2: `r mlm.variance.distribution(housing_3g_t)$totalI2`
```{r}
training_md %>% count(Active.Drug)
drug_3g_t = rma.mv(yi=yi, V=vi, data = training_md, method = "REML", random = ~1|group/Comparison.Code, mods = ~relevel(factor(training_md$Active.Drug), ref = "Anisomycin"))
summary(drug_3g_t)
#pseudo R-squared
drug_3g_t.R2 = (sum(training_meta_article_md$sigma2)-sum(drug_3g_t$sigma2))/sum(training_meta_article_md$sigma2)*100
drug_3g_t.R2 = if_else(drug_3g_t.R2<0, 0, drug_3g_t.R2)
```
Exact model p values: `r drug_3g_t$pval` | 
Exact Q test of moderators p value: `r drug_3g_t$QMp` | 
Total I2: `r mlm.variance.distribution(drug_3g_t)$totalI2`
```{r}
training_md %>% count(Site.injection.other)
sites_3g_t = rma.mv(yi=yi, V=vi, data = training_md, method = "REML", random = ~1|group/Comparison.Code, mods = ~relevel(factor(training_md$Site.injection.other), ref = "Systemic"))
summary(sites_3g_t)
#pseudo R-squared
sites_3g_t.R2 = (sum(training_meta_article_md$sigma2)-sum(sites_3g_t$sigma2))/sum(training_meta_article_md$sigma2)*100
sites_3g_t.R2 = if_else(sites_3g_t.R2<0, 0, sites_3g_t.R2)
```
Exact model p values: `r sites_3g_t$pval` | 
Exact Q test of moderators p value: `r sites_3g_t$QMp` | 
Total I2: `r mlm.variance.distribution(sites_3g_t)$totalI2`
```{r}
#dose_3g_t = rma.mv(yi=yi, V=vi, data = training_md, method = "REML", random = ~1|group/Comparison.Code, mods = training_md$dose_zscore)
#summary(dose_3g_t)
#pseudo R-squared
#dose_3g_t.R2 = (sum(training_meta_article_md$sigma2)-sum(dose_3g_t$sigma2))/sum(training_meta_article_md$sigma2)*100
#dose_3g_t.R2 = if_else(dose_3g_t.R2<0, 0, dose_3g_t.R2)
```

```{r}
dose_3g_t_s = rma.mv(yi=yi, V=vi, data = training_md, method = "REML", random = ~1|group/Comparison.Code, mods = training_md$dose_zscore_s)
summary(dose_3g_t_s)
#pseudo R-squared
dose_3g_t.R2_s = (sum(training_meta_article_md$sigma2)-sum(dose_3g_t_s$sigma2))/sum(training_meta_article_md$sigma2)*100
dose_3g_t.R2_s = if_else(dose_3g_t.R2_s<0, 0, dose_3g_t.R2_s)
```
Exact model p values: `r dose_3g_t_s$pval` | 
Exact Q test of moderators p value: `r dose_3g_t_s$QMp` | 
Total I2: `r mlm.variance.distribution(dose_3g_t_s)$totalI2`
```{r}
training_md %>% count(Randomization)
randomization_3g_t = rma.mv(yi=yi, V=vi, data = training_md, method = "REML", random = ~1|group/Comparison.Code, mods = ~factor(training_md$Randomization))
summary(randomization_3g_t)
#pseudo R-squared
randomization_3g_t.R2 = (sum(training_meta_article_md$sigma2)-sum(randomization_3g_t$sigma2))/sum(training_meta_article_md$sigma2)*100
randomization_3g_t.R2 = if_else(randomization_3g_t.R2<0, 0, randomization_3g_t.R2)
```
Exact model p values: `r randomization_3g_t$pval` | 
Exact Q test of moderators p value: `r randomization_3g_t$QMp` | 
Total I2: `r mlm.variance.distribution(randomization_3g_t)$totalI2`
```{r}
training_md %>% count(Blinding)
blinding_3g_t = rma.mv(yi=yi, V=vi, data = training_md, method = "REML", random = ~1|group/Comparison.Code, mods = ~factor(training_md$Blinding))
summary(blinding_3g_t)
#pseudo R-squared
blinding_3g_t.R2 = (sum(training_meta_article_md$sigma2)-sum(blinding_3g_t$sigma2))/sum(training_meta_article_md$sigma2)*100
blinding_3g_t.R2 = if_else(blinding_3g_t.R2<0, 0, blinding_3g_t.R2)
```
Exact model p values: `r blinding_3g_t$pval` | 
Exact Q test of moderators p value: `r blinding_3g_t$QMp` | 
Total I2: `r mlm.variance.distribution(blinding_3g_t)$totalI2`
```{r}
training_md %>% count(RegulatoryReq)
requirements_3g_t = rma.mv(yi=yi, V=vi, data = training_md, method = "REML", random = ~1|group/Comparison.Code, mods = ~factor(training_md$RegulatoryReq))
summary(requirements_3g_t)
#pseudo R-squared
requirements_3g_t.R2 = (sum(training_meta_article_md$sigma2)-sum(requirements_3g_t$sigma2))/sum(training_meta_article_md$sigma2)*100
requirements_3g_t.R2 = if_else(requirements_3g_t.R2<0, 0, requirements_3g_t.R2)
```
Exact model p values: `r requirements_3g_t$pval` | 
Exact Q test of moderators p value: `r requirements_3g_t$QMp` | 
Total I2: `r mlm.variance.distribution(requirements_3g_t)$totalI2`
```{r}
training_md %>% count(ConflictInterest)
conflict_3g_t = rma.mv(yi=yi, V=vi, data = training_md, method = "REML", random = ~1|group/Comparison.Code, mods = ~factor(training_md$ConflictInterest))
summary(conflict_3g_t)
#pseudo R-squared
conflict_3g_t.R2 = (sum(training_meta_article_md$sigma2)-sum(conflict_3g_t$sigma2))/sum(training_meta_article_md$sigma2)*100
conflict_3g_t.R2 = if_else(conflict_3g_t.R2<0, 0, conflict_3g_t.R2)
```
Exact model p values: `r conflict_3g_t$pval` | 
Exact Q test of moderators p value: `r conflict_3g_t$QMp` | 
Total I2: `r mlm.variance.distribution(conflict_3g_t)$totalI2`
```{r}
impact_3g_t = rma.mv(yi=yi, V=vi, data = training_md, method = "REML", random = ~1|group/Comparison.Code, mods = training_md$Impact.Factor)
summary(impact_3g_t)
#pseudo R-squared
impact_3g_t.R2 = (sum(training_meta_article_md$sigma2)-sum(impact_3g_t$sigma2))/sum(training_meta_article_md$sigma2)*100
impact_3g_t.R2 = if_else(impact_3g_t.R2<0, 0, impact_3g_t.R2)
```
Exact model p values: `r impact_3g_t$pval` | 
Exact Q test of moderators p value: `r impact_3g_t$QMp` | 
Total I2: `r mlm.variance.distribution(impact_3g_t)$totalI2`
```{r}
citations_3g_t = rma.mv(yi=yi, V=vi, data = training_md, method = "REML", random = ~1|group/Comparison.Code, mods = training_md$Citations.per.year)
summary(citations_3g_t)
#pseudo R-squared
citations_3g_t.R2 = (sum(training_meta_article_md$sigma2)-sum(citations_3g_t$sigma2))/sum(training_meta_article_md$sigma2)*100
citations_3g_t.R2 = if_else(citations_3g_t.R2<0, 0, citations_3g_t.R2)
```
Exact model p values: `r citations_3g_t$pval` | 
Exact Q test of moderators p value: `r citations_3g_t$QMp` | 
Total I2: `r mlm.variance.distribution(citations_3g_t)$totalI2`
```{r}
training_md %>% count(Region.of.origin)
region_3g_t = rma.mv(yi=yi, V=vi, data = training_md, method = "REML", random = ~1|group/Comparison.Code, mods = ~relevel(factor(training_md$Region.of.origin), ref="North America"))
summary(region_3g_t)
#pseudo R-squared
region_3g_t.R2 = (sum(training_meta_article_md$sigma2)-sum(region_3g_t$sigma2))/sum(training_meta_article_md$sigma2)*100
region_3g_t.R2 = if_else(region_3g_t.R2<0, 0, region_3g_t.R2)
```
Exact model p values: `r region_3g_t$pval` | 
Exact Q test of moderators p value: `r region_3g_t$QMp` | 
Total I2: `r mlm.variance.distribution(region_3g_t)$totalI2`

Reactivation dataset:
(Suppl Table 21)
```{r}
reactivation_md %>% count(Intervention.Session)
session_3g_r = rma.mv(yi=yi, V=vi, data = reactivation_md, method = "REML", random = ~1|group/Comparison.Code, mods = ~relevel(factor(reactivation_md$Intervention.Session), ref="Reactivation"))
summary(session_3g_r)
#pseudo R-squared
session_3g_r.R2 = (sum(reactivation_meta_article_md$sigma2)-sum(session_3g_r$sigma2))/sum(reactivation_meta_article_md$sigma2)*100
session_3g_r.R2 = if_else(session_3g_r.R2<0, 0, session_3g_r.R2)
```
Exact model p values: `r session_3g_r$pval` | 
Exact Q test of moderators p value: `r session_3g_r$QMp` | 
Total I2: `r mlm.variance.distribution(session_3g_r)$totalI2`
```{r}
reactivation_md %>% count(ConditioningType)
conditioning_3g_r = rma.mv(yi=yi, V=vi, data = reactivation_md, method = "REML", random = ~1|group/Comparison.Code, mods = ~factor(reactivation_md$ConditioningType))
summary(conditioning_3g_r)
#pseudo R-squared
conditioning_3g_r.R2 = (sum(reactivation_meta_article_md$sigma2)-sum(conditioning_3g_r$sigma2))/sum(reactivation_meta_article_md$sigma2)*100
conditioning_3g_r.R2 = if_else(conditioning_3g_r.R2<0, 0, conditioning_3g_r.R2)
```
Exact model p values: `r conditioning_3g_r$pval` | 
Exact Q test of moderators p value: `r conditioning_3g_r$QMp` | 
Total I2: `r mlm.variance.distribution(conditioning_3g_r)$totalI2`
```{r}
reactivation_md %>% count(Habituation)
habituation_3g_r = rma.mv(yi=yi, V=vi, data = reactivation_md, method = "REML", random = ~1|group/Comparison.Code, mods = ~factor(reactivation_md$Habituation))
summary(habituation_3g_r)
#pseudo R-squared
habituation_3g_r.R2 = (sum(reactivation_meta_article_md$sigma2)-sum(habituation_3g_r$sigma2))/sum(reactivation_meta_article_md$sigma2)*100
habituation_3g_r.R2 = if_else(habituation_3g_r.R2<0, 0, habituation_3g_r.R2)
```
Exact model p values: `r habituation_3g_r$pval` | 
Exact Q test of moderators p value: `r habituation_3g_r$QMp` | 
Total I2: `r mlm.variance.distribution(habituation_3g_r)$totalI2`
```{r}
reactivation_md %>% count(Handling)
handling_3g_r = rma.mv(yi=yi, V=vi, data = reactivation_md, method = "REML", random = ~1|group/Comparison.Code, mods = ~factor(reactivation_md$Handling))
summary(handling_3g_r)
#pseudo R-squared
handling_3g_r.R2 = (sum(reactivation_meta_article_md$sigma2)-sum(handling_3g_r$sigma2))/sum(reactivation_meta_article_md$sigma2)*100
handling_3g_r.R2 = if_else(handling_3g_r.R2<0, 0, handling_3g_r.R2)
```
Exact model p values: `r handling_3g_r$pval` | 
Exact Q test of moderators p value: `r handling_3g_r$QMp` | 
Total I2: `r mlm.variance.distribution(handling_3g_r)$totalI2`
```{r}
shocks_3g_r = rma.mv(yi=yi, V=vi, data = reactivation_md, method = "REML", random = ~1|group/Comparison.Code, mods = reactivation_md$NumberShock)
summary(shocks_3g_r)
#pseudo R-squared
shocks_3g_r.R2 = (sum(reactivation_meta_article_md$sigma2)-sum(shocks_3g_r$sigma2))/sum(reactivation_meta_article_md$sigma2)*100
shocks_3g_r.R2 = if_else(shocks_3g_r.R2<0, 0, shocks_3g_r.R2)
```
Exact model p values: `r shocks_3g_r$pval` | 
Exact Q test of moderators p value: `r shocks_3g_r$QMp` | 
Total I2: `r mlm.variance.distribution(shocks_3g_r)$totalI2`
```{r}
intensity_3g_r = rma.mv(yi=yi, V=vi, data = reactivation_md, method = "REML", random = ~1|group/Comparison.Code, mods = reactivation_md$ShockIntensity)
summary(intensity_3g_r)
#pseudo R-squared
intensity_3g_r.R2 = (sum(reactivation_meta_article_md$sigma2)-sum(intensity_3g_r$sigma2))/sum(reactivation_meta_article_md$sigma2)*100
intensity_3g_r.R2 = if_else(intensity_3g_r.R2<0, 0, intensity_3g_r.R2)
```
Exact model p values: `r intensity_3g_r$pval` | 
Exact Q test of moderators p value: `r intensity_3g_r$QMp` | 
Total I2: `r mlm.variance.distribution(intensity_3g_r)$totalI2`
```{r}
time_reexposure_3g_r = rma.mv(yi=yi, V=vi, data = reactivation_md, method = "REML", random = ~1|group/Comparison.Code, mods = reactivation_md$Time.to.reexposure)
summary(time_reexposure_3g_r)
#pseudo R-squared
time_reexposure_3g_r.R2 = (sum(reactivation_meta_article_md$sigma2)-sum(time_reexposure_3g_r$sigma2))/sum(reactivation_meta_article_md$sigma2)*100
time_reexposure_3g_r.R2 = if_else(time_reexposure_3g_r.R2<0, 0, time_reexposure_3g_r.R2)
```
Exact model p values: `r time_reexposure_3g_r$pval` | 
Exact Q test of moderators p value: `r time_reexposure_3g_r$QMp` | 
Total I2: `r mlm.variance.distribution(time_reexposure_3g_r)$totalI2`
```{r}
time_intervention_3g_r = rma.mv(yi=yi, V=vi, data = reactivation_md, method = "REML", random = ~1|group/Comparison.Code, mods = reactivation_md$InterventionTime)
summary(time_intervention_3g_r)
#pseudo R-squared
time_intervention_3g_r.R2 = (sum(reactivation_meta_article_md$sigma2)-sum(time_intervention_3g_r$sigma2))/sum(reactivation_meta_article_md$sigma2)*100
time_intervention_3g_r.R2 = if_else(time_intervention_3g_r.R2<0, 0, time_intervention_3g_r.R2)
```
Exact model p values: `r time_intervention_3g_r$pval` | 
Exact Q test of moderators p value: `r time_intervention_3g_r$QMp` | 
Total I2: `r mlm.variance.distribution(time_intervention_3g_r)$totalI2`
```{r}
duration_3g_r = rma.mv(yi=yi, V=vi, data = reactivation_md, method = "REML", random = ~1|group/Comparison.Code, mods = reactivation_md$duration_zscore)
summary(duration_3g_r)
#pseudo R-squared
duration_3g_r.R2 = (sum(reactivation_meta_article_md$sigma2)-sum(duration_3g_r$sigma2))/sum(reactivation_meta_article_md$sigma2)*100
duration_3g_r.R2 = if_else(duration_3g_r.R2<0, 0, duration_3g_r.R2)
```
Exact model p values: `r duration_3g_r$pval` | 
Exact Q test of moderators p value: `r duration_3g_r$QMp` | 
Total I2: `r mlm.variance.distribution(duration_3g_r)$totalI2`
```{r}
time_test_3g_r = rma.mv(yi=yi, V=vi, data = reactivation_md, method = "REML", random = ~1|group/Comparison.Code, mods = reactivation_md$TimetoTest)
summary(time_test_3g_r)
#pseudo R-squared
time_test_3g_r.R2 = (sum(reactivation_meta_article_md$sigma2)-sum(time_test_3g_r$sigma2))/sum(reactivation_meta_article_md$sigma2)*100
time_test_3g_r.R2 = if_else(time_test_3g_r.R2<0, 0, time_test_3g_r.R2)
```
Exact model p values: `r time_test_3g_r$pval` | 
Exact Q test of moderators p value: `r time_test_3g_r$QMp` | 
Total I2: `r mlm.variance.distribution(time_test_3g_r)$totalI2`
```{r}
reactivation_md %>% count(Species)
species_3g_r = rma.mv(yi=yi, V=vi, data = reactivation_md, method = "REML", random = ~1|group/Comparison.Code, mods = ~factor(reactivation_md$Species))
summary(species_3g_r)
#pseudo R-squared
species_3g_r.R2 = (sum(reactivation_meta_article_md$sigma2)-sum(species_3g_r$sigma2))/sum(reactivation_meta_article_md$sigma2)*100
species_3g_r.R2 = if_else(species_3g_r.R2<0, 0, species_3g_r.R2)
```
Exact model p values: `r species_3g_r$pval` | 
Exact Q test of moderators p value: `r species_3g_r$QMp` | 
Total I2: `r mlm.variance.distribution(species_3g_r)$totalI2`
```{r}
reactivation_md %>% count(Sex)
sex_3g_r = rma.mv(yi=yi, V=vi, data = reactivation_md, method = "REML", random = ~1|group/Comparison.Code, mods = ~factor(reactivation_md$Sex))
summary(sex_3g_r)
#pseudo R-squared
sex_3g_r.R2 = (sum(reactivation_meta_article_md$sigma2)-sum(sex_3g_r$sigma2))/sum(reactivation_meta_article_md$sigma2)*100
sex_3g_r.R2 = if_else(sex_3g_r.R2<0, 0, sex_3g_r.R2)
```
Exact model p values: `r sex_3g_r$pval` | 
Exact Q test of moderators p value: `r sex_3g_r$QMp` | 
Total I2: `r mlm.variance.distribution(sex_3g_r)$totalI2`
```{r}
age_3g_r = rma.mv(yi=yi, V=vi, data = reactivation_md, method = "REML", random = ~1|group/Comparison.Code, mods = reactivation_md$Age_mean)
summary(age_3g_r)
#pseudo R-squared
age_3g_r.R2 = (sum(reactivation_meta_article_md$sigma2)-sum(age_3g_r$sigma2))/sum(reactivation_meta_article_md$sigma2)*100
age_3g_r.R2 = if_else(age_3g_r.R2<0, 0, age_3g_r.R2)
```
Exact model p values: `r age_3g_r$pval` | 
Exact Q test of moderators p value: `r age_3g_r$QMp` | 
Total I2: `r mlm.variance.distribution(age_3g_r)$totalI2`
```{r}
housing_3g_r = rma.mv(yi=yi, V=vi, data = reactivation_md, method = "REML", random = ~1|group/Comparison.Code, mods = reactivation_md$Housing)
summary(housing_3g_r)
#pseudo R-squared
housing_3g_r.R2 = (sum(reactivation_meta_article_md$sigma2)-sum(housing_3g_r$sigma2))/sum(reactivation_meta_article_md$sigma2)*100
housing_3g_r.R2 = if_else(housing_3g_r.R2<0, 0, housing_3g_r.R2)
```
Exact model p values: `r housing_3g_r$pval` | 
Exact Q test of moderators p value: `r housing_3g_r$QMp` | 
Total I2: `r mlm.variance.distribution(housing_3g_r)$totalI2`
```{r}
reactivation_md %>% count(Active.Drug)
drug_3g_r = rma.mv(yi=yi, V=vi, data = reactivation_md, method = "REML", random = ~1|group/Comparison.Code, mods = ~relevel(factor(reactivation_md$Active.Drug), ref = "Anisomycin"))
summary(drug_3g_r)
#pseudo R-squared
drug_3g_r.R2 = (sum(reactivation_meta_article_md$sigma2)-sum(drug_3g_r$sigma2))/sum(reactivation_meta_article_md$sigma2)*100
drug_3g_r.R2 = if_else(drug_3g_r.R2<0, 0, drug_3g_r.R2)
```
Exact model p values: `r drug_3g_r$pval` | 
Exact Q test of moderators p value: `r drug_3g_r$QMp` | 
Total I2: `r mlm.variance.distribution(drug_3g_r)$totalI2`
```{r}
reactivation_md %>% count(Site.injection.other)
sites_3g_r = rma.mv(yi=yi, V=vi, data = reactivation_md, method = "REML", random = ~1|group/Comparison.Code, mods = ~relevel(factor(reactivation_md$Site.injection.other), ref = "Systemic"))
summary(sites_3g_r)
#pseudo R-squared
sites_3g_r.R2 = (sum(reactivation_meta_article_md$sigma2)-sum(sites_3g_r$sigma2))/sum(reactivation_meta_article_md$sigma2)*100
sites_3g_r.R2 = if_else(sites_3g_r.R2<0, 0, sites_3g_r.R2)
```
Exact model p values: `r sites_3g_r$pval` | 
Exact Q test of moderators p value: `r sites_3g_r$QMp` | 
Total I2: `r mlm.variance.distribution(sites_3g_r)$totalI2`
```{r}
#dose_3g_r = rma.mv(yi=yi, V=vi, data = reactivation_md, method = "REML", random = ~1|group/Comparison.Code, mods = reactivation_md$dose_zscore)
#summary(dose_3g_r)
#pseudo R-squared
#dose_3g_r.R2 = (sum(reactivation_meta_article_md$sigma2)-sum(dose_3g_r$sigma2))/sum(reactivation_meta_article_md$sigma2)*100
#dose_3g_r.R2 = if_else(dose_3g_r.R2<0, 0, dose_3g_r.R2)
```

```{r}
dose_3g_r_s = rma.mv(yi=yi, V=vi, data = reactivation_md, method = "REML", random = ~1|group/Comparison.Code, mods = reactivation_md$dose_zscore_s)
summary(dose_3g_r_s)
#pseudo R-squared
dose_3g_r.R2_s = (sum(reactivation_meta_article_md$sigma2)-sum(dose_3g_r_s$sigma2))/sum(reactivation_meta_article_md$sigma2)*100
dose_3g_r.R2_s = if_else(dose_3g_r.R2_s<0, 0, dose_3g_r.R2_s)
```
Exact model p values: `r dose_3g_r_s$pval` | 
Exact Q test of moderators p value: `r dose_3g_r_s$QMp` | 
Total I2: `r mlm.variance.distribution(dose_3g_r_s)$totalI2`
```{r}
reactivation_md %>% count(Randomization)
randomization_3g_r = rma.mv(yi=yi, V=vi, data = reactivation_md, method = "REML", random = ~1|group/Comparison.Code, mods = ~factor(reactivation_md$Randomization))
summary(randomization_3g_r)
#pseudo R-squared
randomization_3g_r.R2 = (sum(reactivation_meta_article_md$sigma2)-sum(randomization_3g_r$sigma2))/sum(reactivation_meta_article_md$sigma2)*100
randomization_3g_r.R2 = if_else(randomization_3g_r.R2<0, 0, randomization_3g_r.R2)
```
Exact model p values: `r randomization_3g_r$pval` | 
Exact Q test of moderators p value: `r randomization_3g_r$QMp` | 
Total I2: `r mlm.variance.distribution(randomization_3g_r)$totalI2`
```{r}
reactivation_md %>% count(Blinding)
blinding_3g_r = rma.mv(yi=yi, V=vi, data = reactivation_md, method = "REML", random = ~1|group/Comparison.Code, mods = ~factor(reactivation_md$Blinding))
summary(blinding_3g_r)
#pseudo R-squared
blinding_3g_r.R2 = (sum(reactivation_meta_article_md$sigma2)-sum(blinding_3g_r$sigma2))/sum(reactivation_meta_article_md$sigma2)*100
blinding_3g_r.R2 = if_else(blinding_3g_r.R2<0, 0, blinding_3g_r.R2)
```
Exact model p values: `r blinding_3g_r$pval` | 
Exact Q test of moderators p value: `r blinding_3g_r$QMp` | 
Total I2: `r mlm.variance.distribution(blinding_3g_r)$totalI2`
```{r}
reactivation_md %>% count(SampleSizeCalc)
samplesize_3g_r = rma.mv(yi=yi, V=vi, data = reactivation_md, method = "REML", random = ~1|group/Comparison.Code, mods = ~factor(reactivation_md$SampleSizeCalc))
summary(samplesize_3g_r)
#pseudo R-squared
samplesize_3g_r.R2 = (sum(reactivation_meta_article_md$sigma2)-sum(samplesize_3g_r$sigma2))/sum(reactivation_meta_article_md$sigma2)*100
samplesize_3g_r.R2 = if_else(samplesize_3g_r.R2<0, 0, samplesize_3g_r.R2)
```
Exact model p values: `r samplesize_3g_r$pval` | 
Exact Q test of moderators p value: `r samplesize_3g_r$QMp` | 
Total I2: `r mlm.variance.distribution(samplesize_3g_r)$totalI2`
```{r}
reactivation_md %>% count(RegulatoryReq)
requirements_3g_r = rma.mv(yi=yi, V=vi, data = reactivation_md, method = "REML", random = ~1|group/Comparison.Code, mods = ~factor(reactivation_md$RegulatoryReq))
summary(requirements_3g_r)
#pseudo R-squared
requirements_3g_r.R2 = (sum(reactivation_meta_article_md$sigma2)-sum(requirements_3g_r$sigma2))/sum(reactivation_meta_article_md$sigma2)*100
requirements_3g_r.R2 = if_else(requirements_3g_r.R2<0, 0, requirements_3g_r.R2)
```
Exact model p values: `r requirements_3g_r$pval` | 
Exact Q test of moderators p value: `r requirements_3g_r$QMp` | 
Total I2: `r mlm.variance.distribution(requirements_3g_r)$totalI2`
```{r}
reactivation_md %>% count(ConflictInterest)
conflict_3g_r = rma.mv(yi=yi, V=vi, data = reactivation_md, method = "REML", random = ~1|group/Comparison.Code, mods = ~relevel(factor(reactivation_md$ConflictInterest), ref = "Not informed"))
summary(conflict_3g_r)
#pseudo R-squared
conflict_3g_r.R2 = (sum(reactivation_meta_article_md$sigma2)-sum(conflict_3g_r$sigma2))/sum(reactivation_meta_article_md$sigma2)*100
conflict_3g_r.R2 = if_else(conflict_3g_r.R2<0, 0, conflict_3g_r.R2)
```
Exact model p values: `r conflict_3g_r$pval` | 
Exact Q test of moderators p value: `r conflict_3g_r$QMp` | 
Total I2: `r mlm.variance.distribution(conflict_3g_r)$totalI2`
```{r}
impact_3g_r = rma.mv(yi=yi, V=vi, data = reactivation_md, method = "REML", random = ~1|group/Comparison.Code, mods = reactivation_md$Impact.Factor)
summary(impact_3g_r)
#pseudo R-squared
impact_3g_r.R2 = (sum(reactivation_meta_article_md$sigma2)-sum(impact_3g_r$sigma2))/sum(reactivation_meta_article_md$sigma2)*100
impact_3g_r.R2 = if_else(impact_3g_r.R2<0, 0, impact_3g_r.R2)
```
Exact model p values: `r impact_3g_r$pval` | 
Exact Q test of moderators p value: `r impact_3g_r$QMp` | 
Total I2: `r mlm.variance.distribution(impact_3g_r)$totalI2`
```{r}
citations_3g_r = rma.mv(yi=yi, V=vi, data = reactivation_md, method = "REML", random = ~1|group/Comparison.Code, mods = reactivation_md$Citations.per.year)
summary(citations_3g_r)
#pseudo R-squared
citations_3g_r.R2 = (sum(reactivation_meta_article_md$sigma2)-sum(citations_3g_r$sigma2))/sum(reactivation_meta_article_md$sigma2)*100
citations_3g_r.R2 = if_else(citations_3g_r.R2<0, 0, citations_3g_r.R2)
```
Exact model p values: `r citations_3g_r$pval` | 
Exact Q test of moderators p value: `r citations_3g_r$QMp` | 
Total I2: `r mlm.variance.distribution(citations_3g_r)$totalI2`
```{r}
reactivation_md %>% count(Region.of.origin)
region_3g_r = rma.mv(yi=yi, V=vi, data = reactivation_md, method = "REML", random = ~1|group/Comparison.Code, mods = ~relevel(factor(reactivation_md$Region.of.origin), ref="North America"))
summary(region_3g_r)
region_3g_r$pval
mlm.variance.distribution(region_3g_r)$totalI2
#pseudo R-squared
region_3g_r.R2 = (sum(reactivation_meta_article_md$sigma2)-sum(region_3g_r$sigma2))/sum(reactivation_meta_article_md$sigma2)*100
region_3g_r.R2 = if_else(region_3g_r.R2<0, 0, region_3g_r.R2)
```
Exact model p values: `r region_3g_r$pval` | 
Exact Q test of moderators p value: `r region_3g_r$QMp` | 
Total I2: `r mlm.variance.distribution(region_3g_r)$totalI2`

Fig 3C:
```{r}
dataset = c('Complete, by group')
moderadores = c('Conditioning Type', 'Habituation reported', 'Handling reported', 'Number of shocks', 'Shock intensity', 'Intervention time', 'Time to reexposure', 'Reexposure duration', 'Time to test', 'Species', 'Sex', 'Mean age', 'Housing', 'Active drug', 'Injection site', 'Dose', 'Randomization', 'Blinding', 'Sample size calculation', 'Regulatory requirements', 'Conflict of interest statement', 'Impact factor', 'Citations per year', 'Region of origin')
rquadrado = c(conditioning_3g.R2, habituation_3g.R2, handling_3g.R2, shocks_3g.R2, intensity_3g.R2, time_intervention_3g.R2, NA, NA, time_test_3g.R2, species_3g.R2, sex_3g.R2, age_3g.R2, housing_3g.R2, drug_3g.R2, sites_3g.R2, dose_3g.R2_s, randomization_3g.R2, blinding_3g.R2, samplesize_3g.R2, requirements_3g.R2, conflict_3g.R2, impact_3g.R2, citations_3g.R2, region_3g.R2)
pvalor = c(conditioning_3g$QMp, habituation_3g$QMp, handling_3g$QMp, shocks_3g$QMp, intensity_3g$QMp, time_intervention_3g$QMp, NA, NA, time_test_3g$QMp, species_3g$QMp, sex_3g$QMp, age_3g$QMp, housing_3g$QMp, drug_3g$QMp, sites_3g$QMp, dose_3g_s$QMp, randomization_3g$QMp, blinding_3g$QMp, samplesize_3g$QMp, requirements_3g$QMp, conflict_3g$QMp, impact_3g$QMp, citations_3g$QMp, region_3g$QMp)

moderators_3g = data.frame(dataset, moderadores, rquadrado, pvalor)
moderators_3g = moderators_3g %>% mutate(pval_cat = if_else(pvalor>0.05, ">0.05", if_else(pvalor<=0.05&pvalor>0.01, "<0.05", if_else(pvalor<=0.01&pvalor>0.001, "<0.01", "<0.001"))))
```
```{r}
dataset = c('Training, by group')
rquadrado = c(conditioning_3g_t.R2, habituation_3g_t.R2, handling_3g_t.R2, shocks_3g_t.R2, intensity_3g_t.R2, time_intervention_3g_t.R2, NA, NA, time_test_3g_t.R2, species_3g_t.R2, sex_3g_t.R2, age_3g_t.R2, housing_3g_t.R2, drug_3g_t.R2, sites_3g_t.R2, dose_3g_t.R2_s, randomization_3g_t.R2, blinding_3g_t.R2, NA, requirements_3g_t.R2, conflict_3g_t.R2, impact_3g_t.R2, citations_3g_t.R2, region_3g_t.R2)
pvalor = c(conditioning_3g_t$QMp, habituation_3g_t$QMp, handling_3g_t$QMp, shocks_3g_t$QMp, intensity_3g_t$QMp, time_intervention_3g_t$QMp, NA, NA, time_test_3g_t$QMp, species_3g_t$QMp, sex_3g_t$QMp, age_3g_t$QMp, housing_3g_t$QMp, drug_3g_t$QMp, sites_3g_t$QMp, dose_3g_t_s$QMp, randomization_3g_t$QMp, blinding_3g_t$QMp, NA, requirements_3g_t$QMp, conflict_3g_t$QMp, impact_3g_t$QMp, citations_3g_t$QMp, region_3g_t$QMp)

moderators_3g_t = data.frame(dataset, moderadores, rquadrado, pvalor)
moderators_3g_t = moderators_3g_t %>% mutate(pval_cat = if_else(pvalor>0.05, ">0.05", if_else(pvalor<=0.05&pvalor>0.01, "<0.05", if_else(pvalor<=0.01&pvalor>0.001, "<0.01", "<0.001"))))
```
```{r}
dataset = c('Reactivation, by group')
rquadrado = c(conditioning_3g_r.R2, habituation_3g_r.R2, handling_3g_r.R2, shocks_3g_r.R2, intensity_3g_r.R2, time_intervention_3g_r.R2, time_reexposure_3g_r.R2, duration_3g_r.R2, time_test_3g_r.R2, species_3g_r.R2, sex_3g_r.R2, age_3g_r.R2, housing_3g_r.R2, drug_3g_r.R2, sites_3g_r.R2, dose_3g_r.R2_s, randomization_3g_r.R2, blinding_3g_r.R2, samplesize_3g_r.R2, requirements_3g_r.R2, conflict_3g_r.R2, impact_3g_r.R2, citations_3g_r.R2, region_3g_r.R2)
pvalor = c(conditioning_3g_r$QMp, habituation_3g_r$QMp, handling_3g_r$QMp, shocks_3g_r$QMp, intensity_3g_r$QMp, time_intervention_3g_r$QMp, time_reexposure_3g_r$QMp, duration_3g_r$QMp, time_test_3g_r$QMp, species_3g_r$QMp, sex_3g_r$QMp, age_3g_r$QMp, housing_3g_r$QMp, drug_3g_r$QMp, sites_3g_r$QMp, dose_3g_r_s$QMp, randomization_3g_r$QMp, blinding_3g_r$QMp, samplesize_3g_r$QMp, requirements_3g_r$QMp, conflict_3g_r$QMp, impact_3g_r$QMp, citations_3g_r$QMp, region_3g_r$QMp)

moderators_3g_r = data.frame(dataset, moderadores, rquadrado, pvalor)
moderators_3g_r = moderators_3g_r %>% mutate(pval_cat = if_else(pvalor>0.05, ">0.05", if_else(pvalor<=0.05&pvalor>0.01, "<0.05", if_else(pvalor<=0.01&pvalor>0.001, "<0.01", "<0.001"))))
```
```{r}
all_moderators_3g = bind_rows(moderators_3g, moderators_3g_t, moderators_3g_r)
all_moderators_3g$moderadores = factor(all_moderators_3g$moderadores, levels = rev(factor(moderadores)))
all_moderators_3g$dataset = factor(all_moderators_3g$dataset, levels = c("Complete, by group", "Training, by group", "Reactivation, by group"))
labels_3g = c("Complete,\nby group","Training,\nby group", "Reactivation,\nby group")

all_mods_3g = ggplot(all_moderators_3g, aes(x=dataset, y=moderadores, fill=pval_cat)) + geom_tile() + scale_fill_manual(values=c("#3182bd","#6baed6","#bdd7e7","#eff3ff"), na.value = "white", name = "Q test for moderators \np value") + geom_text(aes(x=dataset, y=moderadores, label = if_else(condition = rquadrado>=0, true = paste(round(rquadrado, 1), "%"), false = " ")), color = "black", size = 6) + scale_x_discrete(position = "top", labels=labels_3g) + theme(axis.title.x = element_blank(), axis.text.x.top = element_text(angle = 0, size=14), axis.title.y = element_blank(), panel.background = element_blank(),  axis.ticks = element_blank(), legend.position = "bottom", axis.text = element_text(size=15), legend.text = element_text(size=12), legend.title = element_text(size=12))
all_mods_3g
```

Complete figure 3:
```{r}
plot_grid(all_mods, all_mods_3a, all_mods_3g, ncol = 3, labels = LETTERS)
ggsave("FIG_3.png", width = 60, height = 40, units = "cm")
```

**Meta-regression visualizations**

Suppl. Fig. 9:
```{r}
bubble_t_time3 = ggplot(training_md, aes(x=InterventionTime, y=yi, size = 1/sqrt(vi))) + 
  geom_point(shape = 1) + 
  labs(y = "Absolute mean difference", x = "Time (min) between injection and training") + 
  theme_classic() + 
  scale_size_continuous(guide = "none") + 
  geom_abline(slope = 0, linetype = "dashed", color = "grey") +
  geom_abline(intercept = time_intervention_t$beta[1,1], slope = time_intervention_t$beta[2,1], size=0.1)
bubble_t_time3
ggsave("FIG_S9.png", width = 15, height = 10, units = "cm")
```

Suppl. Fig. 10:
```{r}
bubble_r_conditioning = ggplot(reactivation_md, aes(x=ConditioningType, y=yi, size = 1/sqrt(vi), color = Intervention.Session)) + 
  geom_point(shape = 1, position =  position_jitterdodge(dodge.width = 0), stroke = 1) + 
  labs(y = "Absolute mean difference", x = "Type of conditioning") + 
  theme_classic() + 
  scale_size_continuous(guide = "none") + 
  theme(legend.position = "top") + 
  scale_color_manual(values = c("#1b9e77", "#7570b3", "#d95f02")) + 
  geom_abline(yintercept = 0, slope = 0, linetype = "dashed", color = "grey")

bubble_r_duration_context = ggplot((reactivation_md %>% filter(ConditioningType=="Contextual" | ConditioningType=="Contextual background")), aes(x = ReexposureDuration_min, y=yi, size = 1/sqrt(vi), color = Intervention.Session)) + 
  geom_point(shape = 1, stroke = 1) + 
  labs(y = "Absolute mean difference", x = "Reexposure duration (min)") + 
  theme_classic() + scale_size_continuous(guide = "none") + 
  theme(legend.position = "none") + 
  geom_abline(slope = 0, linetype = "dashed", color = "grey") + 
  scale_color_manual(values = c("#1b9e77", "#7570b3", "#d95f02")) +
  geom_abline(intercept = duration_r_context$beta[1,1], slope = duration_r_context$beta[2,1], size=0.1)

bubble_r_duration_tone = ggplot((reactivation_md %>% filter(ConditioningType=="Tone" | ConditioningType=="Tone-Trace")), aes(x = ReexposureDuration_number, y=yi, size = 1/sqrt(vi), color = Intervention.Session)) + 
  geom_point(shape = 1, stroke = 1) + 
  labs(y = "Absolute mean difference", x = "Reexposure duration (tones)") + theme_classic() + 
  scale_size_continuous(guide = "none") + 
  theme(legend.position = "none") + 
  geom_abline(slope = 0, linetype = "dashed", color = "grey") + 
  scale_color_manual(values = c("#1b9e77", "#7570b3", "#d95f02")) +
  geom_abline(intercept = duration_r_tone$beta[1,1], slope = duration_r_tone$beta[2,1], size=0.1)

plot_grid(bubble_r_duration_context, bubble_r_duration_tone, bubble_r_conditioning, ncol = 1, labels = LETTERS)
ggsave("FIG_S10.png", height = 30, units = "cm")
```

Suppl Fig 11
```{r}
bubble_r_number_shocks_new = ggplot(reactivation_md, aes(x = NumberShock, y=yi, size = 1/sqrt(vi), color = Intervention.Session)) + 
  geom_point(shape = 1, stroke = 1) + 
  labs(y = "Absolute mean difference", x = "Number of shocks") + 
  theme_classic() + 
  scale_size_continuous(guide = "none") + 
  theme(legend.position = "bottom") + 
  geom_abline(slope = 0, linetype = "dashed", color = "grey") + 
  scale_color_manual(values = c("#1b9e77", "#7570b3", "#d95f02")) + 
  scale_y_continuous(n.breaks = 7) + 
  scale_x_continuous(n.breaks = 7) +
  geom_abline(intercept = shocks_r$beta[1,1], slope = shocks_r$beta[2,1], size=0.1)

bubble_r_time_reexp_d_new = ggplot(reactivation_md, aes(x = Time.to.reexposure/24, y=yi, size = 1/sqrt(vi), color = Intervention.Session)) + 
  geom_point(shape = 1, stroke = 1) + 
  labs(y = "Absolute mean difference", x = "Time between training and reexposure (days)") + 
  theme_classic() + 
  scale_size_continuous(guide = "none") + 
  theme(legend.position = "bottom") + 
  geom_abline(slope = 0, linetype = "dashed", color = "grey") + 
  scale_color_manual(values = c("#1b9e77", "#7570b3", "#d95f02")) + 
  scale_y_continuous(n.breaks = 7) + 
  scale_x_continuous(n.breaks = 11) +
  geom_abline(intercept = time_reexposure_r$beta[1,1], slope = time_reexposure_r$beta[2,1], size=0.1)

plot_grid(bubble_r_number_shocks_new, bubble_r_time_reexp_d_new, ncol = 1, labels = LETTERS)
ggsave("FIG_S11.png", height = 20, units = "cm")
```
```{r}
reactivation_md %>% count(Intervention.Session)
```

**Multivariate meta-regressions**
Following the tutorial presented at http://www.metafor-project.org/doku.php/tips:model_selection_with_glmulti_and_mumin
```{r mv-functions}
rma.glmulti <- function(formula, data, ...)
   rma(formula, vi, data=data, method="ML", ...)
rma.mv.glmulti <- function(formula, data, ...)
   rma.mv(formula, vi, data=data, method="ML", random = ~1|ID/Comparison.Code, verbose = FALSE, ...)
rma.mv.glmulti2 <- function(formula, data, ...)
   rma.mv(formula, vi, data=data, method="ML", random = ~1|group/Comparison.Code, ...)
```

Functions to decompose R2 (3-level, article)
```{r}
# Roda todos os modelos, par a par, full x reduced, a partir das fÃ³rmulas, e faz a diferença de R2
all_model_comparisons_3a = function (comparison_list) {
  map2_dfr(comparison_list$reduced_models, comparison_list$full_models, function (m0s, m1s) {
    if (m0s == "") {
      # Se nÃ£o tem preditor, Ã© o modelo sem moderadores
      m0 = rma(yi, vi, data=dat, method="ML", control = list(maxiter = 3000))
    } else {
      # Se tem preditores, usa a fÃ³rmula que vem da outra funÃ§Ã£o
      m0f = as.formula(m0s)
      m0 = rma.mv(yi, vi, random = ~1|ID/Comparison.Code, mods = m0f, data=dat, method="ML", control = list(maxiter = 3000))
    }
    
    m1f = as.formula(m1s)
    m1 = rma.mv(yi, vi, random = ~1|ID/Comparison.Code, mods = m1f, data=dat, method="ML", control = list(maxiter = 3000))
   
    m2 = rma.mv(yi=yi, V=vi, data = dat, method = "REML", random = ~1|ID/Comparison.Code, control = list(maxiter = 3000)) 
    
    reduced_model_size = length(all.vars(m1f)) - 1
    
    tibble(reduced = m0s, full = m1s, R2 = ((sum(m2$sigma2)-sum(m1$sigma2))/sum(m2$sigma2)*100)-((sum(m2$sigma2)-sum(m0$sigma2))/sum(m2$sigma2)*100), permutations = factorial(reduced_model_size))
  })
}

# Faz todo o processo pra um Ãºnico preditor e resume na mÃ©dia dos R2
get_R2_for_single_predictor_3a = function (todos_preditores, preditor_interesse) {
  comparison_list = build_model_combinations(todos_preditores, preditor_interesse)
  
  r = all_model_comparisons_3a(comparison_list)
  s = mean(r$R2, na.rm = T)
  # Pra essa, o peso Ã© o nÃºmero de permutaÃ§Ãµes do modelo reduzido (se a variÃ¡vel de interesse Ã© 4a, tem 3! maneiras possÃ­veis dela ser a 4a, porque tem 3 variÃ¡veis antes dela pra permutar ... faz sentido?)
  ws = weighted.mean(r$R2, r$permutations, na.rm = T)
  
  list(all_predictors = todos_preditores, predictor = preditor_interesse, results = r, mean = s, wmean = ws)
}

# Faz todo o processo em separado pra cada um dos preditores da lista
get_R2_for_all_predictors_3a = function (todos_preditores) {
  df = map(todos_preditores, function (preditor_interesse) {
    print(paste("Running models for", preditor_interesse))
    get_R2_for_single_predictor_3a(todos_preditores, preditor_interesse)
  })
  s = tibble(mean_R2 = map_dbl(df, "mean"), weighted_mean_R2 = map_dbl(df, "wmean"), predictor = map_chr(df, "predictor"))
  list(full_results = df, summary = s)
}
```

Functions to decompose R2 (3-level, group)
```{r}
# Roda todos os modelos, par a par, full x reduced, a partir das fÃ³rmulas, e faz a diferença de R2
all_model_comparisons_3g = function (comparison_list) {
  map2_dfr(comparison_list$reduced_models, comparison_list$full_models, function (m0s, m1s) {
    if (m0s == "") {
      # Se nÃ£o tem preditor, Ã© o modelo sem moderadores
      m0 = rma(yi, vi, data=dat, method="ML", control = list(maxiter = 3000))
    } else {
      # Se tem preditores, usa a fÃ³rmula que vem da outra funÃ§Ã£o
      m0f = as.formula(m0s)
      m0 = rma.mv(yi, vi, random = ~1|group/Comparison.Code, mods = m0f, data=dat, method="ML", control = list(maxiter = 3000))
    }
    
    m1f = as.formula(m1s)
    m1 = rma.mv(yi, vi, random = ~1|group/Comparison.Code, mods = m1f, data=dat, method="ML", control = list(maxiter = 3000))
   
    m2 = rma.mv(yi=yi, V=vi, data = dat, method = "REML", random = ~1|group/Comparison.Code, control = list(maxiter = 3000)) 
    
    reduced_model_size = length(all.vars(m1f)) - 1
    
    tibble(reduced = m0s, full = m1s, R2 = ((sum(m2$sigma2)-sum(m1$sigma2))/sum(m2$sigma2)*100)-((sum(m2$sigma2)-sum(m0$sigma2))/sum(m2$sigma2)*100), permutations = factorial(reduced_model_size))
  })
}

# Faz todo o processo pra um Ãºnico preditor e resume na mÃ©dia dos R2
get_R2_for_single_predictor_3g = function (todos_preditores, preditor_interesse) {
  comparison_list = build_model_combinations(todos_preditores, preditor_interesse)
  
  r = all_model_comparisons_3g(comparison_list)
  s = mean(r$R2, na.rm = T)
  # Pra essa, o peso Ã© o nÃºmero de permutaÃ§Ãµes do modelo reduzido (se a variÃ¡vel de interesse Ã© 4a, tem 3! maneiras possÃ­veis dela ser a 4a, porque tem 3 variÃ¡veis antes dela pra permutar ... faz sentido?)
  ws = weighted.mean(r$R2, r$permutations, na.rm = T)
  
  list(all_predictors = todos_preditores, predictor = preditor_interesse, results = r, mean = s, wmean = ws)
}

# Faz todo o processo em separado pra cada um dos preditores da lista
get_R2_for_all_predictors_3g = function (todos_preditores) {
  df = map(todos_preditores, function (preditor_interesse) {
    print(paste("Running models for", preditor_interesse))
    get_R2_for_single_predictor_3g(todos_preditores, preditor_interesse)
  })
  s = tibble(mean_R2 = map_dbl(df, "mean"), weighted_mean_R2 = map_dbl(df, "wmean"), predictor = map_chr(df, "predictor"))
  list(full_results = df, summary = s)
}
```

Complete dataset:
```{r}
full_data_complete <- full_data_md
full_data_complete <-  full_data_complete[!apply(full_data_complete[,c("Intervention.Session", "Site.injection.other", "ConditioningType", "Habituation", "Handling", "NumberShock", "ShockIntensity", "InterventionTime", "TimetoTest", "Species", "Sex", "Age_mean", "Housing", "Active.Drug", "Site.of.injection", "dose_zscore_s", "Randomization", "Blinding", "SampleSizeCalc", "RegulatoryReq", "ConflictInterest", "Impact.Factor", "Citations.per.year", "Region.of.origin")], 1, anyNA),]

full_data_selected <- full_data_md
full_data_selected <-  full_data_selected[!apply(full_data_selected[,c("Intervention.Session", "Site.injection.other", "Active.Drug", "dose_zscore_s", "TimetoTest", "Species", "NumberShock", "ShockIntensity", "Sex", "ConditioningType", "InterventionTime")], 1, anyNA),]
```
Considering all 24 possible variables, we'd have `r nrow(full_data_complete)` experiments available (i.e. `r nrow(full_data_md)-nrow(full_data_complete)` exclusions due to missing data). Including only the 11 variables selected, we have `r nrow(full_data_selected)` experiments (i.e. `r nrow(full_data_md)-nrow(full_data_selected)` exclusions due to missing data) and `r 2^11` possible models.

2-level, All possible models:
```{r results='hide'}
res <- glmulti(yi ~ relevel(factor(Intervention.Session), ref = "Training") + relevel(factor(Site.injection.other), ref = "Systemic") + Active.Drug + dose_zscore_s + TimetoTest + Species + NumberShock + ShockIntensity + Sex + relevel(factor(ConditioningType), ref = "Contextual") + InterventionTime, data = full_data_selected, level=1, fitfunction=rma.glmulti, crit="aicc", confsetsize=(2^11), plotty = F)
```
```{r}
print(res)
```

(Suppl. Table 22)
Looking at the best models (within 2 IC units of the best model):
```{r}
top <- weightable(res)
top <- top[top$aicc <= min(top$aicc) + 2,]
top
```

Getting the details for each of the top 10 models:

Model1:
```{r}
summary(res@objects[[1]])
```
Exact model p values: `r res@objects[[1]]$pval` | 
Exact Q test for moderators p value: `r res@objects[[1]]$QMp`

Decomposing R2 for the best model:
```{r}
dat = full_data_selected
meus_preditores = c("relevel(factor(Intervention.Session), ref = 'Training')", "relevel(factor(Site.injection.other), ref = 'Systemic')", "relevel(factor(ConditioningType), ref = 'Contextual')", "Active.Drug", "NumberShock", "InterventionTime")
res.best = get_R2_for_all_predictors(meus_preditores)
res.best.summary = res.best$summary %>% mutate(dataset = "Complete")
all_moderators_multi = res.best.summary
all_moderators_multi = all_moderators_multi %>% mutate(pvalor = 0)
all_moderators_multi[1,5] = anova(res@objects[[1]], btt = 2:4)$QMp
all_moderators_multi[2,5] = anova(res@objects[[1]], btt = 5:8)$QMp
all_moderators_multi[3,5] = anova(res@objects[[1]], btt = 9:11)$QMp
all_moderators_multi[4,5] = anova(res@objects[[1]], btt = 12:13)$QMp
all_moderators_multi[5,5] = anova(res@objects[[1]], btt = 14)$QMp
all_moderators_multi[6,5] = anova(res@objects[[1]], btt = 15)$QMp
```

Model2:
```{r}
summary(res@objects[[2]])
```
Exact model p values: `r res@objects[[2]]$pval` | 
Exact Q test for moderators p value: `r res@objects[[2]]$QMp`

Model3:
```{r}
summary(res@objects[[3]])
```
Exact model p values: `r res@objects[[3]]$pval` | 
Exact Q test for moderators p value: `r res@objects[[3]]$QMp`

Model4:
```{r}
summary(res@objects[[4]])
```
Exact model p values: `r res@objects[[4]]$pval` | 
Exact Q test for moderators p value: `r res@objects[[4]]$QMp`

Model5:
```{r}
summary(res@objects[[5]])
```
Exact model p values: `r res@objects[[5]]$pval` | 
Exact Q test for moderators p value: `r res@objects[[5]]$QMp`

Model6:
```{r}
summary(res@objects[[6]])
```
Exact model p values: `r res@objects[[6]]$pval` | 
Exact Q test for moderators p value: `r res@objects[[6]]$QMp`

Model7:
```{r}
summary(res@objects[[7]])
```
Exact model p values: `r res@objects[[7]]$pval` | 
Exact Q test for moderators p value: `r res@objects[[7]]$QMp`

Model8:
```{r}
summary(res@objects[[8]])
```
Exact model p values: `r res@objects[[8]]$pval` | 
Exact Q test for moderators p value: `r res@objects[[8]]$QMp`

Model9:
```{r}
summary(res@objects[[9]])
```
Exact model p values: `r res@objects[[9]]$pval` | 
Exact Q test for moderators p value: `r res@objects[[9]]$QMp`

Model10:
```{r}
summary(res@objects[[10]])
```
Exact model p values: `r res@objects[[10]]$pval` | 
Exact Q test for moderators p value: `r res@objects[[10]]$QMp`

(Suppl. Table 23)
3-level (article), complete dataset:
All possible models:
```{r results='hide'}
res2 <- glmulti(yi ~ relevel(factor(Intervention.Session), ref = "Training") + relevel(factor(Site.injection.other), ref = "Systemic") + Active.Drug + dose_zscore_s + TimetoTest + Species + NumberShock + ShockIntensity + Sex + relevel(factor(ConditioningType), ref = "Contextual") + InterventionTime, data = full_data_selected, level=1, fitfunction=rma.mv.glmulti, crit="aicc", confsetsize=(2^11), plotty = F)
```
```{r}
print(res2)
```

Looking at the best models (within 2 IC units of the best model):
```{r}
top2 <- weightable(res2)
top2 <- top2[top2$aicc <= min(top2$aicc) + 2,]
top2
```

Getting the details for each of the top 9 models:

Model1:
```{r}
summary(res2@objects[[1]])
```
Exact model p values: `r res2@objects[[1]]$pval` | 
Exact Q test for moderators p value: `r res2@objects[[1]]$QMp` | 
Total I2: `r mlm.variance.distribution(res2@objects[[1]])$totalI2` | 
Pseudo R2: `r (sum(full_data_meta_article_md$sigma2)-sum(res2@objects[[1]]$sigma2))/sum(full_data_meta_article_md$sigma2)*100`

Decomposing R2 for the best model:
```{r}
dat = full_data_selected
meus_preditores = c("relevel(factor(Intervention.Session), ref = 'Training')", "relevel(factor(Site.injection.other), ref = 'Systemic')", "Active.Drug", "NumberShock", "InterventionTime")
res2.best = get_R2_for_all_predictors_3a(meus_preditores)
res2.best.summary = res2.best$summary %>% mutate(dataset = "Complete, by article")
all_moderators_multi_3a = res2.best.summary
all_moderators_multi_3a = all_moderators_multi_3a %>% mutate(pvalor = 0)
all_moderators_multi_3a[1,5] = anova(res2@objects[[1]], btt = 2:4)$QMp
all_moderators_multi_3a[2,5] = anova(res2@objects[[1]], btt = 5:8)$QMp
all_moderators_multi_3a[3,5] = anova(res2@objects[[1]], btt = 9:10)$QMp
all_moderators_multi_3a[4,5] = anova(res2@objects[[1]], btt = 11)$QMp
all_moderators_multi_3a[5,5] = anova(res2@objects[[1]], btt = 12)$QMp
```

Model2:
```{r}
summary(res2@objects[[2]])
```
Exact model p values: `r res2@objects[[2]]$pval` | 
Exact Q test for moderators p value: `r res2@objects[[2]]$QMp` | 
Total I2: `r mlm.variance.distribution(res2@objects[[2]])$totalI2` | 
Pseudo R2: `r (sum(full_data_meta_article_md$sigma2)-sum(res2@objects[[2]]$sigma2))/sum(full_data_meta_article_md$sigma2)*100`

Model3:
```{r}
summary(res2@objects[[3]])
```
Exact model p values: `r res2@objects[[3]]$pval` | 
Exact Q test for moderators p value: `r res2@objects[[3]]$QMp` | 
Total I2: `r mlm.variance.distribution(res2@objects[[3]])$totalI2` | 
Pseudo R2: `r (sum(full_data_meta_article_md$sigma2)-sum(res2@objects[[3]]$sigma2))/sum(full_data_meta_article_md$sigma2)*100`

Model4:
```{r}
summary(res2@objects[[4]])
```
Exact model p values: `r res2@objects[[4]]$pval` | 
Exact Q test for moderators p value: `r res2@objects[[4]]$QMp` | 
Total I2: `r mlm.variance.distribution(res2@objects[[4]])$totalI2` | 
Pseudo R2: `r (sum(full_data_meta_article_md$sigma2)-sum(res2@objects[[4]]$sigma2))/sum(full_data_meta_article_md$sigma2)*100`

Model5:
```{r}
summary(res2@objects[[5]])
```
Exact model p values: `r res2@objects[[5]]$pval` | 
Exact Q test for moderators p value: `r res2@objects[[5]]$QMp` | 
Total I2: `r mlm.variance.distribution(res2@objects[[5]])$totalI2` | 
Pseudo R2: `r (sum(full_data_meta_article_md$sigma2)-sum(res2@objects[[5]]$sigma2))/sum(full_data_meta_article_md$sigma2)*100`

Model6:
```{r}
summary(res2@objects[[6]])
```
Exact model p values: `r res2@objects[[6]]$pval` | 
Exact Q test for moderators p value: `r res2@objects[[6]]$QMp` | 
Total I2: `r mlm.variance.distribution(res2@objects[[6]])$totalI2` | 
Pseudo R2: `r (sum(full_data_meta_article_md$sigma2)-sum(res2@objects[[6]]$sigma2))/sum(full_data_meta_article_md$sigma2)*100`

Model7:
```{r}
summary(res2@objects[[7]])
```
Exact model p values: `r res2@objects[[7]]$pval` | 
Exact Q test for moderators p value: `r res2@objects[[7]]$QMp` | 
Total I2: `r mlm.variance.distribution(res2@objects[[7]])$totalI2` | 
Pseudo R2: `r (sum(full_data_meta_article_md$sigma2)-sum(res2@objects[[7]]$sigma2))/sum(full_data_meta_article_md$sigma2)*100`

Model8:
```{r}
summary(res2@objects[[8]])
```
Exact model p values: `r res2@objects[[8]]$pval` | 
Exact Q test for moderators p value: `r res2@objects[[8]]$QMp` | 
Total I2: `r mlm.variance.distribution(res2@objects[[8]])$totalI2` | 
Pseudo R2: `r (sum(full_data_meta_article_md$sigma2)-sum(res2@objects[[8]]$sigma2))/sum(full_data_meta_article_md$sigma2)*100`

Model9:
```{r}
summary(res2@objects[[9]])
```
Exact model p values: `r res2@objects[[9]]$pval` | 
Exact Q test for moderators p value: `r res2@objects[[9]]$QMp` | 
Total I2: `r mlm.variance.distribution(res2@objects[[9]])$totalI2` | 
Pseudo R2: `r (sum(full_data_meta_article_md$sigma2)-sum(res2@objects[[9]]$sigma2))/sum(full_data_meta_article_md$sigma2)*100`

(Suppl Table 24)
3-level (group), complete dataset:
```{r results='hide'}
res3 <- glmulti(yi ~ relevel(factor(Intervention.Session), ref = "Training") + relevel(factor(Site.injection.other), ref = "Systemic") + Active.Drug + dose_zscore_s + TimetoTest + Species + NumberShock + ShockIntensity + Sex + relevel(factor(ConditioningType), ref = "Contextual") + InterventionTime, data = full_data_selected, level=1, fitfunction=rma.mv.glmulti2, crit="aicc", confsetsize=(2^11), plotty = F)
```
```{r}
print(res3)
```

Looking at the best models (within 2 IC units of the best model):
```{r}
top3 <- weightable(res3)
top3 <- top3[top3$aicc <= min(top3$aicc) + 2,]
top3
```

Getting the details for each of the top 3 models:

Model1:
```{r}
summary(res3@objects[[1]])
```
Exact model p values: `r res3@objects[[1]]$pval` | 
Exact Q test for moderators p value: `r res3@objects[[1]]$QMp` | 
Total I2: `r mlm.variance.distribution(res3@objects[[1]])$totalI2` | 
Pseudo R2: `r (sum(full_data_meta_group_md$sigma2)-sum(res3@objects[[1]]$sigma2))/sum(full_data_meta_group_md$sigma2)*100`

Decomposing R2 for the best model:
```{r}
dat = full_data_selected
meus_preditores = c("relevel(factor(Intervention.Session), ref = 'Training')", "relevel(factor(Site.injection.other), ref = 'Systemic')", "Active.Drug", "NumberShock", "InterventionTime")
res3.best = get_R2_for_all_predictors_3g(meus_preditores)
res3.best.summary = res3.best$summary %>% mutate(dataset = "Complete, by group")
all_moderators_multi_3g = res3.best.summary
all_moderators_multi_3g = all_moderators_multi_3g %>% mutate(pvalor = 0)
all_moderators_multi_3g[1,5] = anova(res3@objects[[1]], btt = 2:4)$QMp
all_moderators_multi_3g[2,5] = anova(res3@objects[[1]], btt = 5:8)$QMp
all_moderators_multi_3g[3,5] = anova(res3@objects[[1]], btt = 9:10)$QMp
all_moderators_multi_3g[4,5] = anova(res3@objects[[1]], btt = 11)$QMp
all_moderators_multi_3g[5,5] = anova(res3@objects[[1]], btt = 12)$QMp
```

Model2:
```{r}
summary(res3@objects[[2]])
```
Exact model p values: `r res3@objects[[2]]$pval` | 
Exact Q test for moderators p value: `r res3@objects[[2]]$QMp` | 
Total I2: `r mlm.variance.distribution(res3@objects[[2]])$totalI2` | 
Pseudo R2: `r (sum(full_data_meta_group_md$sigma2)-sum(res3@objects[[2]]$sigma2))/sum(full_data_meta_group_md$sigma2)*100`

Model3:
```{r}
summary(res3@objects[[3]])
```
Exact model p values: `r res3@objects[[3]]$pval` | 
Exact Q test for moderators p value: `r res3@objects[[3]]$QMp` | 
Total I2: `r mlm.variance.distribution(res3@objects[[3]])$totalI2` | 
Pseudo R2: `r (sum(full_data_meta_group_md$sigma2)-sum(res3@objects[[3]]$sigma2))/sum(full_data_meta_group_md$sigma2)*100`

Sample sizes:
```{r}
full_data_selected %>% count(Intervention.Session)
full_data_selected %>% count(Site.injection.other)
full_data_selected %>% count(ConditioningType)
full_data_selected %>% count(Active.Drug)
full_data_selected %>% count(Species)
full_data_selected %>% count(Sex)
```

Training dataset
```{r}
training_complete <- training_md
training_complete <-  training_complete[!apply(training_complete[,c("Site.injection.other", "ConditioningType", "Habituation", "Handling", "NumberShock", "ShockIntensity", "InterventionTime", "TimetoTest", "Species", "Sex", "Age_mean", "Housing", "Active.Drug", "dose_zscore_s", "Randomization", "Blinding", "SampleSizeCalc", "RegulatoryReq", "ConflictInterest", "Impact.Factor", "Citations.per.year", "Region.of.origin")], 1, anyNA),]

training_selected_plusage <- training_md
training_selected_plusage <-  training_selected_plusage[!apply(training_selected_plusage[,c("Site.injection.other", "Active.Drug", "dose_zscore_s", "TimetoTest", "Species", "NumberShock", "ShockIntensity", "Sex", "Age_mean", "ConditioningType", "InterventionTime")], 1, anyNA),]

training_selected <- training_md
training_selected <-  training_selected[!apply(training_selected[,c("Site.injection.other", "Active.Drug", "dose_zscore_s", "TimetoTest", "Species", "NumberShock", "ShockIntensity", "Sex", "ConditioningType", "InterventionTime")], 1, anyNA),]
```
Considering all 22 possible variables, we'd have `r nrow(training_complete)` experiments available (i.e. `r nrow(training_md)-nrow(training_complete)` exclusions due to missing data). Including only the 10 variables selected, we have `r nrow(training_selected)` experiments (i.e. `r nrow(training_md)-nrow(training_selected)` exclusions due to missing data) and `r 2^10` possible models.

(Suppl Table 25)
2-level, training dataset:
```{r results='hide'}
res4 <- glmulti(yi ~ relevel(factor(Site.injection.other), ref = "Systemic") + Active.Drug + dose_zscore_s + TimetoTest + Species + NumberShock + ShockIntensity + Sex + relevel(factor(ConditioningType), ref = "Contextual") + InterventionTime, data = training_selected, level=1, fitfunction=rma.glmulti, crit="aicc", confsetsize=(2^10), plotty = F)
```
```{r}
print(res4)
```

Looking at the best models (within 2 IC units of the best model):
```{r}
top4 <- weightable(res4)
top4 <- top4[top4$aicc <= min(top4$aicc) + 2,]
top4
```

Getting the details for each of the top 12 models:

Model1:
```{r}
summary(res4@objects[[1]])
```
Exact model p values: `r res4@objects[[1]]$pval` | 
Exact Q test for moderators p value: `r res4@objects[[1]]$QMp`

Decomposing R2 for the best model:
```{r}
dat = training_selected
meus_preditores = c("relevel(factor(Site.injection.other), ref = 'Systemic')", "InterventionTime")
res4.best = get_R2_for_all_predictors(meus_preditores)
res4.best.summary = res4.best$summary %>% mutate(dataset = "Training")
all_moderators_multi = bind_rows(all_moderators_multi, res4.best.summary)
all_moderators_multi[7,5] = anova(res4@objects[[1]], btt = 2:5)$QMp
all_moderators_multi[8,5] = anova(res4@objects[[1]], btt = 6)$QMp
```

Model2:
```{r}
summary(res4@objects[[2]])
```
Exact model p values: `r res4@objects[[2]]$pval` | 
Exact Q test for moderators p value: `r res4@objects[[2]]$QMp`

Model3:
```{r}
summary(res4@objects[[3]])
```
Exact model p values: `r res4@objects[[3]]$pval` | 
Exact Q test for moderators p value: `r res4@objects[[3]]$QMp`

Model4:
```{r}
summary(res4@objects[[4]])
```
Exact model p values: `r res4@objects[[4]]$pval` | 
Exact Q test for moderators p value: `r res4@objects[[4]]$QMp`

Model5:
```{r}
summary(res4@objects[[5]])
```
Exact model p values: `r res4@objects[[5]]$pval` | 
Exact Q test for moderators p value: `r res4@objects[[5]]$QMp`

Model6:
```{r}
summary(res4@objects[[6]])
```
Exact model p values: `r res4@objects[[6]]$pval` | 
Exact Q test for moderators p value: `r res4@objects[[6]]$QMp`

Model7:
```{r}
summary(res4@objects[[7]])
```
Exact model p values: `r res4@objects[[7]]$pval` | 
Exact Q test for moderators p value: `r res4@objects[[7]]$QMp`

Model8:
```{r}
summary(res4@objects[[8]])
```
Exact model p values: `r res4@objects[[8]]$pval` | 
Exact Q test for moderators p value: `r res4@objects[[8]]$QMp`

Model9:
```{r}
summary(res4@objects[[9]])
```
Exact model p values: `r res4@objects[[9]]$pval` | 
Exact Q test for moderators p value: `r res4@objects[[9]]$QMp`

Model10:
```{r}
summary(res4@objects[[10]])
```
Exact model p values: `r res4@objects[[10]]$pval` | 
Exact Q test for moderators p value: `r res4@objects[[10]]$QMp`

Model11:
```{r}
summary(res4@objects[[11]])
```
Exact model p values: `r res4@objects[[11]]$pval` | 
Exact Q test for moderators p value: `r res4@objects[[11]]$QMp`

Model12:
```{r}
summary(res4@objects[[12]])
```
Exact model p values: `r res4@objects[[12]]$pval` | 
Exact Q test for moderators p value: `r res4@objects[[12]]$QMp`

(Suppl Table 26)
3-level (article), training dataset:
```{r results='hide'}
res5 <- glmulti(yi ~ relevel(factor(Site.injection.other), ref = "Systemic") + Active.Drug + dose_zscore_s + TimetoTest + Species + NumberShock + ShockIntensity + Sex + relevel(factor(ConditioningType), ref = "Contextual") + InterventionTime, data = training_selected, level=1, fitfunction=rma.mv.glmulti, crit="aicc", confsetsize=(2^10), plotty = F)
```
```{r}
print(res5)
```

Looking at the best models (within 2 IC units of the best model):
```{r}
top5 <- weightable(res5)
top5 <- top5[top5$aicc <= min(top5$aicc) + 2,]
top5
```

Getting the details for each of the top 11 models:

Model1:
```{r}
summary(res5@objects[[1]])
```
Exact model p values: `r res5@objects[[1]]$pval` | 
Exact Q test for moderators p value: `r res5@objects[[1]]$QMp` | 
Total I2: `r mlm.variance.distribution(res5@objects[[1]])$totalI2` | 
Pseudo R2: `r (sum(training_meta_article_md$sigma2)-sum(res5@objects[[1]]$sigma2))/sum(training_meta_article_md$sigma2)*100`

Decomposing R2 for the best model:
```{r}
dat = training_selected
meus_preditores = c("relevel(factor(Site.injection.other), ref = 'Systemic')", "InterventionTime")
res5.best = get_R2_for_all_predictors_3a(meus_preditores)
res5.best.summary = res5.best$summary %>% mutate(dataset = "Training, by article")
all_moderators_multi_3a = bind_rows(all_moderators_multi_3a, res5.best.summary)
all_moderators_multi_3a[6,5] = anova(res5@objects[[1]], btt = 2:5)$QMp
all_moderators_multi_3a[7,5] = anova(res5@objects[[1]], btt = 6)$QMp
```

Model2:
```{r}
summary(res5@objects[[2]])
```
Exact model p values: `r res5@objects[[2]]$pval` | 
Exact Q test for moderators p value: `r res5@objects[[2]]$QMp`
Total I2: `r mlm.variance.distribution(res5@objects[[2]])$totalI2` | 
Pseudo R2: `r (sum(training_meta_article_md$sigma2)-sum(res5@objects[[2]]$sigma2))/sum(training_meta_article_md$sigma2)*100`

Model3:
```{r}
summary(res5@objects[[3]])
```
Exact model p values: `r res5@objects[[3]]$pval` | 
Exact Q test for moderators p value: `r res5@objects[[3]]$QMp` | 
Total I2: `r mlm.variance.distribution(res5@objects[[3]])$totalI2` | 
Pseudo R2: `r (sum(training_meta_article_md$sigma2)-sum(res5@objects[[3]]$sigma2))/sum(training_meta_article_md$sigma2)*100`

Model4:
```{r}
#summary(res5@objects[[4]])
```
Exact model p values: `r res5@objects[[4]]$pval` | 
Exact Q test for moderators p value: `r res5@objects[[4]]$QMp` | 
Total I2: `r mlm.variance.distribution(res5@objects[[4]])$totalI2` | 
Pseudo R2: `r (sum(training_meta_article_md$sigma2)-sum(res5@objects[[4]]$sigma2))/sum(training_meta_article_md$sigma2)*100`

Model5:
```{r}
summary(res5@objects[[5]])
```
Exact model p values: `r res5@objects[[5]]$pval` | 
Exact Q test for moderators p value: `r res5@objects[[5]]$QMp` | 
Total I2: `r mlm.variance.distribution(res5@objects[[5]])$totalI2` | 
Pseudo R2: `r (sum(training_meta_article_md$sigma2)-sum(res5@objects[[5]]$sigma2))/sum(training_meta_article_md$sigma2)*100`

Model6:
```{r}
summary(res5@objects[[6]])
```
Exact model p values: `r res5@objects[[6]]$pval` | 
Exact Q test for moderators p value: `r res5@objects[[6]]$QMp` | 
Total I2: `r mlm.variance.distribution(res5@objects[[6]])$totalI2` | 
Pseudo R2: `r (sum(training_meta_article_md$sigma2)-sum(res5@objects[[6]]$sigma2))/sum(training_meta_article_md$sigma2)*100`

Model7:
```{r}
summary(res5@objects[[7]])
```
Exact model p values: `r res5@objects[[7]]$pval` | 
Exact Q test for moderators p value: `r res5@objects[[7]]$QMp` | 
Total I2: `r mlm.variance.distribution(res5@objects[[7]])$totalI2` | 
Pseudo R2: `r (sum(training_meta_article_md$sigma2)-sum(res5@objects[[7]]$sigma2))/sum(training_meta_article_md$sigma2)*100`

Model8:
```{r}
summary(res5@objects[[8]])
```
Exact model p values: `r res5@objects[[8]]$pval` | 
Exact Q test for moderators p value: `r res5@objects[[8]]$QMp` | 
Total I2: `r mlm.variance.distribution(res5@objects[[8]])$totalI2` | 
Pseudo R2: `r (sum(training_meta_article_md$sigma2)-sum(res5@objects[[8]]$sigma2))/sum(training_meta_article_md$sigma2)*100`

Model9:
```{r}
summary(res5@objects[[9]])
```
Exact model p values: `r res5@objects[[9]]$pval` | 
Exact Q test for moderators p value: `r res5@objects[[9]]$QMp` | 
Total I2: `r mlm.variance.distribution(res5@objects[[9]])$totalI2` | 
Pseudo R2: `r (sum(training_meta_article_md$sigma2)-sum(res5@objects[[9]]$sigma2))/sum(training_meta_article_md$sigma2)*100`

Model10:
```{r}
summary(res5@objects[[10]])
```
Exact model p values: `r res5@objects[[10]]$pval` | 
Exact Q test for moderators p value: `r res5@objects[[10]]$QMp` | 
Total I2: `r mlm.variance.distribution(res5@objects[[10]])$totalI2` | 
Pseudo R2: `r (sum(training_meta_article_md$sigma2)-sum(res5@objects[[10]]$sigma2))/sum(training_meta_article_md$sigma2)*100`

Model11:
```{r}
summary(res5@objects[[11]])
```
Exact model p values: `r res5@objects[[11]]$pval` | 
Exact Q test for moderators p value: `r res5@objects[[11]]$QMp` | 
Total I2: `r mlm.variance.distribution(res5@objects[[11]])$totalI2` | 
Pseudo R2: `r (sum(training_meta_article_md$sigma2)-sum(res5@objects[[11]]$sigma2))/sum(training_meta_article_md$sigma2)*100`

(Suppl Table 27)
3-level (group), training dataset:
```{r results='hide'}
res6 <- glmulti(yi ~ relevel(factor(Site.injection.other), ref = "Systemic") + Active.Drug + dose_zscore_s + TimetoTest + Species + NumberShock + ShockIntensity + Sex + relevel(factor(ConditioningType), ref = "Contextual") + InterventionTime, data = training_selected, level=1, fitfunction=rma.mv.glmulti2, crit="aicc", confsetsize=(2^10), plotty = F)
```
```{r}
print(res6)
```

Looking at the best models (within 2 IC units of the best model):
```{r}
top6 <- weightable(res6)
top6 <- top6[top6$aicc <= min(top6$aicc) + 2,]
top6
```

Getting the details for each of the top 11 models:

Model1:
```{r}
summary(res6@objects[[1]])
```
Exact model p values: `r res6@objects[[1]]$pval` | 
Exact Q test for moderators p value: `r res6@objects[[1]]$QMp` | 
Total I2: `r mlm.variance.distribution(res6@objects[[1]])$totalI2` | 
Pseudo R2: `r (sum(training_meta_group_md$sigma2)-sum(res6@objects[[1]]$sigma2))/sum(training_meta_group_md$sigma2)*100`

Decomposing R2 for the best model:
```{r}
dat = training_selected
meus_preditores = c("relevel(factor(Site.injection.other), ref = 'Systemic')", "InterventionTime")
res6.best = get_R2_for_all_predictors_3g(meus_preditores)
res6.best.summary = res6.best$summary %>% mutate(dataset = "Training, by group")
all_moderators_multi_3g = bind_rows(all_moderators_multi_3g, res6.best.summary)
all_moderators_multi_3g[6,5] = anova(res6@objects[[1]], btt = 2:5)$QMp
all_moderators_multi_3g[7,5] = anova(res6@objects[[1]], btt = 6)$QMp
```

Model2:
```{r}
summary(res6@objects[[2]])
```
Exact model p values: `r res6@objects[[2]]$pval` | 
Exact Q test for moderators p value: `r res6@objects[[2]]$QMp` | 
Total I2: `r mlm.variance.distribution(res6@objects[[2]])$totalI2` | 
Pseudo R2: `r (sum(training_meta_group_md$sigma2)-sum(res6@objects[[2]]$sigma2))/sum(training_meta_group_md$sigma2)*100`

Model3:
```{r}
summary(res6@objects[[3]])
```
Exact model p values: `r res6@objects[[3]]$pval` | 
Exact Q test for moderators p value: `r res6@objects[[3]]$QMp` | 
Total I2: `r mlm.variance.distribution(res6@objects[[3]])$totalI2` | 
Pseudo R2: `r (sum(training_meta_group_md$sigma2)-sum(res6@objects[[3]]$sigma2))/sum(training_meta_group_md$sigma2)*100`

Model4:
```{r}
summary(res6@objects[[4]])
```
Exact model p values: `r res6@objects[[4]]$pval` | 
Exact Q test for moderators p value: `r res6@objects[[4]]$QMp` | 
Total I2: `r mlm.variance.distribution(res6@objects[[4]])$totalI2` | 
Pseudo R2: `r (sum(training_meta_group_md$sigma2)-sum(res6@objects[[4]]$sigma2))/sum(training_meta_group_md$sigma2)*100`

Model5:
```{r}
summary(res6@objects[[5]])
```
Exact model p values: `r res6@objects[[5]]$pval` | 
Exact Q test for moderators p value: `r res6@objects[[5]]$QMp` | 
Total I2: `r mlm.variance.distribution(res6@objects[[5]])$totalI2` | 
Pseudo R2: `r (sum(training_meta_group_md$sigma2)-sum(res6@objects[[5]]$sigma2))/sum(training_meta_group_md$sigma2)*100`

Model6:
```{r}
summary(res6@objects[[6]])
```
Exact model p values: `r res6@objects[[6]]$pval` | 
Exact Q test for moderators p value: `r res6@objects[[6]]$QMp` | 
Total I2: `r mlm.variance.distribution(res6@objects[[6]])$totalI2` | 
Pseudo R2: `r (sum(training_meta_group_md$sigma2)-sum(res6@objects[[6]]$sigma2))/sum(training_meta_group_md$sigma2)*100`

Model7:
```{r}
summary(res6@objects[[7]])
```
Exact model p values: `r res6@objects[[7]]$pval` | 
Exact Q test for moderators p value: `r res6@objects[[7]]$QMp` | 
Total I2: `r mlm.variance.distribution(res6@objects[[7]])$totalI2` | 
Pseudo R2: `r (sum(training_meta_group_md$sigma2)-sum(res6@objects[[7]]$sigma2))/sum(training_meta_group_md$sigma2)*100`

Model8:
```{r}
summary(res6@objects[[8]])
```
Exact model p values: `r res6@objects[[8]]$pval` | 
Exact Q test for moderators p value: `r res6@objects[[8]]$QMp` | 
Total I2: `r mlm.variance.distribution(res6@objects[[8]])$totalI2` | 
Pseudo R2: `r (sum(training_meta_group_md$sigma2)-sum(res6@objects[[8]]$sigma2))/sum(training_meta_group_md$sigma2)*100`

Model9:
```{r}
summary(res6@objects[[9]])
```
Exact model p values: `r res6@objects[[9]]$pval` | 
Exact Q test for moderators p value: `r res6@objects[[9]]$QMp` | 
Total I2: `r mlm.variance.distribution(res6@objects[[9]])$totalI2` | 
Pseudo R2: `r (sum(training_meta_group_md$sigma2)-sum(res6@objects[[9]]$sigma2))/sum(training_meta_group_md$sigma2)*100`

Model10:
```{r}
summary(res6@objects[[10]])
```
Exact model p values: `r res6@objects[[10]]$pval` | 
Exact Q test for moderators p value: `r res6@objects[[10]]$QMp` | 
Total I2: `r mlm.variance.distribution(res6@objects[[10]])$totalI2` | 
Pseudo R2: `r (sum(training_meta_group_md$sigma2)-sum(res6@objects[[10]]$sigma2))/sum(training_meta_group_md$sigma2)*100`

Model11:
```{r}
summary(res6@objects[[11]])
```
Exact model p values: `r res6@objects[[11]]$pval` | 
Exact Q test for moderators p value: `r res6@objects[[11]]$QMp` | 
Total I2: `r mlm.variance.distribution(res6@objects[[11]])$totalI2` | 
Pseudo R2: `r (sum(training_meta_group_md$sigma2)-sum(res6@objects[[11]]$sigma2))/sum(training_meta_group_md$sigma2)*100`

Sample sizes:
```{r}
training_selected %>% count(Site.injection.other)
training_selected %>% count(Species)
training_selected %>% count(Active.Drug)
```

Reactivation dataset
```{r}
reactivation_complete <- reactivation_md
reactivation_complete <-  reactivation_complete[!apply(reactivation_complete[,c("Site.injection.other", "ConditioningType", "Habituation", "Handling", "NumberShock", "ShockIntensity", "InterventionTime", "TimetoTest", "Species", "Sex", "Age_mean", "Housing", "Active.Drug", "dose_zscore_s", "Randomization", "Blinding", "SampleSizeCalc", "RegulatoryReq", "ConflictInterest", "Impact.Factor", "Citations.per.year", "Region.of.origin", "duration_zscore", "Time.to.reexposure")], 1, anyNA),]

reactivation_selected_plusage <- reactivation_md
reactivation_selected_plusage <-  reactivation_selected_plusage[!apply(reactivation_selected_plusage[,c("Site.injection.other", "duration_zscore", "Active.Drug", "dose_zscore_s", "TimetoTest", "Time.to.reexposure", "Species", "NumberShock", "ShockIntensity", "Sex", "Age_mean")], 1, anyNA),]

reactivation_selected <- reactivation_md
reactivation_selected <-  reactivation_selected[!apply(reactivation_selected[,c("Site.injection.other", "duration_zscore", "Active.Drug", "dose_zscore_s", "TimetoTest", "Time.to.reexposure", "Species", "NumberShock", "ShockIntensity", "Sex")], 1, anyNA),]
```
Considering all 22 possible variables, we'd have `r nrow(reactivation_complete)` experiments available (i.e. `r nrow(reactivation_md)-nrow(reactivation_complete)` exclusions due to missing data). Including only the 10 variables selected, we have `r nrow(reactivation_selected)` experiments (i.e. `r nrow(reactivation_md)-nrow(reactivation_selected)` exclusions due to missing data) and `r 2^10` possible models.

(Suppl Table 28)
2-level, reactivation dataset:
```{r results='hide'}
res7 <- glmulti(yi ~ relevel(factor(Site.injection.other), ref = "Systemic") + duration_zscore + Active.Drug + dose_zscore_s + TimetoTest + Time.to.reexposure + Species + NumberShock + ShockIntensity + Sex, data = reactivation_selected, level=1, fitfunction=rma.glmulti, crit="aicc", confsetsize=(2^10), plotty = F)
```
```{r}
print(res7)
```

Looking at the best models (within 2 IC units of the best model):
```{r}
top7 <- weightable(res7)
top7 <- top7[top7$aicc <= min(top7$aicc) + 2,]
top7
```

Getting the details for each of the top 2 models:

Model1:
```{r}
summary(res7@objects[[1]])
```
Exact model p values: `r res7@objects[[1]]$pval` | 
Exact Q test for moderators p value: `r res7@objects[[1]]$QMp`

Decomposing R2 for the best model:
```{r}
dat = reactivation_selected
meus_preditores = c("relevel(factor(Site.injection.other), ref = 'Systemic')", "duration_zscore", "NumberShock")
res7.best = get_R2_for_all_predictors(meus_preditores)
res7.best.summary = res7.best$summary %>% mutate(dataset = "Reactivation")
all_moderators_multi = bind_rows(all_moderators_multi, res7.best.summary)
all_moderators_multi[9,5] = anova(res7@objects[[1]], btt = 2:5)$QMp
all_moderators_multi[10,5] = anova(res7@objects[[1]], btt = 6)$QMp
all_moderators_multi[11,5] = anova(res7@objects[[1]], btt = 7)$QMp
```

Model2:
```{r}
summary(res7@objects[[2]])
```
Exact model p values: `r res7@objects[[2]]$pval` | 
Exact Q test for moderators p value: `r res7@objects[[2]]$QMp`

(Suppl Table 29)
3-level (article), reactivation dataset:
```{r}
res8 <- glmulti(yi ~ relevel(factor(Site.injection.other), ref = "Systemic") + duration_zscore + Active.Drug + dose_zscore_s + TimetoTest + Time.to.reexposure + Species + NumberShock + ShockIntensity + Sex, data = reactivation_selected, level=1, fitfunction=rma.mv.glmulti, crit="aicc", confsetsize=(2^10), plotty = F)
```
```{r}
print(res8)
```

Looking at the best models (within 2 IC units of the best model):
```{r}
top8 <- weightable(res8)
top8 <- top8[top8$aicc <= min(top8$aicc) + 2,]
top8
```

Getting the details for each of the top 2 models:

Model1:
```{r}
summary(res8@objects[[1]])
```
Exact model p values: `r res8@objects[[1]]$pval` | 
Exact Q test for moderators p value: `r res8@objects[[1]]$QMp` | 
Total I2: `r mlm.variance.distribution(res8@objects[[1]])$totalI2` | 
Pseudo R2: `r (sum(reactivation_meta_article_md$sigma2)-sum(res8@objects[[1]]$sigma2))/sum(reactivation_meta_article_md$sigma2)*100`

Decomposing R2 for the best model:
```{r}
dat = reactivation_selected
meus_preditores = c("relevel(factor(Site.injection.other), ref = 'Systemic')", "duration_zscore", "NumberShock")
res8.best = get_R2_for_all_predictors_3a(meus_preditores)
res8.best.summary = res8.best$summary %>% mutate(dataset = "Reactivation, by article")
all_moderators_multi_3a = bind_rows(all_moderators_multi_3a, res8.best.summary)
all_moderators_multi_3a[8,5] = anova(res8@objects[[1]], btt = 2:5)$QMp
all_moderators_multi_3a[9,5] = anova(res8@objects[[1]], btt = 6)$QMp
all_moderators_multi_3a[10,5] = anova(res8@objects[[1]], btt = 7)$QMp
```

Model2:
```{r}
summary(res8@objects[[2]])
```
Exact model p values: `r res8@objects[[2]]$pval` | 
Exact Q test for moderators p value: `r res8@objects[[2]]$QMp` | 
Total I2: `r mlm.variance.distribution(res8@objects[[2]])$totalI2` | 
Pseudo R2: `r (sum(reactivation_meta_article_md$sigma2)-sum(res8@objects[[2]]$sigma2))/sum(reactivation_meta_article_md$sigma2)*100`

(Suppl Table 30)
3-level (group), reactivation dataset:
```{r results='hide'}
res9 <- glmulti(yi ~ relevel(factor(Site.injection.other), ref = "Systemic") + duration_zscore + Active.Drug + dose_zscore_s + TimetoTest + Time.to.reexposure + Species + NumberShock + ShockIntensity + Sex, data = reactivation_selected, level=1, fitfunction=rma.mv.glmulti2, crit="aicc", confsetsize=(2^10), plotty = F)
```
```{r}
print(res9)
```

Looking at the best models (within 2 IC units of the best model):
```{r}
top9 <- weightable(res9)
top9 <- top9[top9$aicc <= min(top9$aicc) + 2,]
top9
```

Getting the details for each of the top 2 models:

Model1:
```{r}
summary(res9@objects[[1]])
```
Exact model p values: `r res9@objects[[1]]$pval` | 
Exact Q test for moderators p value: `r res9@objects[[1]]$QMp` | 
Total I2: `r mlm.variance.distribution(res9@objects[[1]])$totalI2` | 
Pseudo R2: `r (sum(reactivation_meta_group_md$sigma2)-sum(res9@objects[[1]]$sigma2))/sum(reactivation_meta_group_md$sigma2)*100`

Decomposing R2 for the best model:
```{r}
dat = reactivation_selected
meus_preditores = c("relevel(factor(Site.injection.other), ref = 'Systemic')", "duration_zscore", "NumberShock")
res9.best = get_R2_for_all_predictors_3g(meus_preditores)
res9.best.summary = res9.best$summary %>% mutate(dataset = "Reactivation, by group")
all_moderators_multi_3g = bind_rows(all_moderators_multi_3g, res9.best.summary)
all_moderators_multi_3g[8,5] = anova(res9@objects[[1]], btt = 2:5)$QMp
all_moderators_multi_3g[9,5] = anova(res9@objects[[1]], btt = 6)$QMp
all_moderators_multi_3g[10,5] = anova(res9@objects[[1]], btt = 7)$QMp
```

Model2:
```{r}
summary(res9@objects[[2]])
```
Exact model p values: `r res9@objects[[2]]$pval` | 
Exact Q test for moderators p value: `r res9@objects[[2]]$QMp` | 
Total I2: `r mlm.variance.distribution(res9@objects[[2]])$totalI2` | 
Pseudo R2: `r (sum(reactivation_meta_group_md$sigma2)-sum(res9@objects[[2]]$sigma2))/sum(reactivation_meta_group_md$sigma2)*100`

Sample sizes:
```{r}
reactivation_selected %>% count(Site.injection.other)
reactivation_selected %>% count(Sex)
```

Variable importance - Suppl. fig 12:
```{r}
#Complete
plot(res, type="s")
plot(res2, type="s")
plot(res3, type="s")
#Training
plot(res4, type="s")
plot(res5, type="s")
plot(res6, type="s")
#Reactivation
plot(res7, type="s")
plot(res8, type="s")
plot(res9, type="s")
```

Figure with summary of best multivariate models
Fig 4A
```{r}
all_moderators_multi$dataset = factor(all_moderators_multi$dataset, levels = c("Complete", "Training", "Reactivation"))
all_moderators_multi = all_moderators_multi %>% mutate(pval_cat = if_else(pvalor>0.05, ">0.05", if_else(pvalor<=0.05&pvalor>0.01, "<0.05", if_else(pvalor<=0.01&pvalor>0.001, "<0.01", "<0.001"))))
all_moderators_multi[all_moderators_multi=="relevel(factor(Intervention.Session), ref = 'Training')"] = "Intervention session"
all_moderators_multi[all_moderators_multi=="relevel(factor(Site.injection.other), ref = 'Systemic')"] = "Injection site"
all_moderators_multi[all_moderators_multi=="relevel(factor(ConditioningType), ref = 'Contextual')"] = "Conditioning type"
all_moderators_multi[all_moderators_multi=="Active.Drug"] = "Active drug"
all_moderators_multi[all_moderators_multi=="NumberShock"] = "Number of shocks"
all_moderators_multi[all_moderators_multi=="duration_zscore"] = "Reexposure duration"
all_moderators_multi[all_moderators_multi=="InterventionTime"] = "Intervention time"

all_moderators_multi$predictor = factor(all_moderators_multi$predictor, level = c("Injection site", "Active drug", "Reexposure duration", "Intervention time", "Number of shocks", "Intervention session", "Conditioning type"))

all_mods_multi = ggplot(all_moderators_multi, aes(x=dataset, y=predictor, fill=pval_cat)) + 
  geom_tile() + scale_fill_manual(values=c("#3182bd","#6baed6","#bdd7e7","#eff3ff"), na.value = "#CACACA", name = "Q test for moderators \np value") +
  geom_text(aes(x=dataset, y=predictor, label = if_else(condition = weighted_mean_R2>=0, true = paste(round(weighted_mean_R2, 1), "%"), false = " ")), color = "black", size = 6) +
  scale_x_discrete(position = "top", labels=labels_2l) + 
  theme(axis.title.x = element_blank(), axis.text.x.top = element_text(angle = 0, size=14), axis.title.y = element_blank(), panel.background = element_blank(),  axis.ticks = element_blank(), legend.position = "bottom", axis.text=element_text(size=15), legend.text=element_text(size=12), legend.title=element_text(size=12))
all_mods_multi
```
Fig 4B
```{r}
all_moderators_multi_3a$dataset = factor(all_moderators_multi_3a$dataset, levels = c("Complete, by article", "Training, by article", "Reactivation, by article"))
all_moderators_multi_3a = all_moderators_multi_3a %>% mutate(pval_cat = if_else(pvalor>0.05, ">0.05", if_else(pvalor<=0.05&pvalor>0.01, "<0.05", if_else(pvalor<=0.01&pvalor>0.001, "<0.01", "<0.001"))))
all_moderators_multi_3a[all_moderators_multi_3a=="relevel(factor(Intervention.Session), ref = 'Training')"] = "Intervention session"
all_moderators_multi_3a[all_moderators_multi_3a=="relevel(factor(Site.injection.other), ref = 'Systemic')"] = "Injection site"
all_moderators_multi_3a[all_moderators_multi_3a=="relevel(factor(ConditioningType), ref = 'Contextual')"] = "Conditioning type"
all_moderators_multi_3a[all_moderators_multi_3a=="Active.Drug"] = "Active drug"
all_moderators_multi_3a[all_moderators_multi_3a=="NumberShock"] = "Number of shocks"
all_moderators_multi_3a[all_moderators_multi_3a=="duration_zscore"] = "Reexposure duration"
all_moderators_multi_3a[all_moderators_multi_3a=="InterventionTime"] = "Intervention time"
all_moderators_multi_3a$predictor = factor(all_moderators_multi_3a$predictor, level = c("Injection site", "Active drug", "Reexposure duration", "Intervention time", "Number of shocks", "Intervention session", "Conditioning type"))
all_moderators_multi_3a = all_moderators_multi_3a %>% mutate(R2_mod = abs(weighted_mean_R2))
all_moderators_multi_3a = rbind(all_moderators_multi_3a, c(NA, NA, "Conditioning type", "Complete, by article", NA, NA, NA))

all_mods_multi_3a_mod = ggplot(all_moderators_multi_3a, aes(x=dataset, y=predictor, fill=pval_cat)) + 
  geom_tile() + scale_fill_manual(values=c("#3182bd","#6baed6","#bdd7e7","#eff3ff"), na.value = "white", name = "Q test for moderators \np value") +
  geom_text(aes(x=dataset, y=predictor, label =  if_else(condition = R2_mod>=0, true = paste(round(as.numeric(R2_mod), 1), "%"), false = " ")), color = "black", size = 6) +
  scale_x_discrete(position = "top", labels=labels_3a) + 
  theme(axis.title.x = element_blank(), axis.text.x.top = element_text(angle = 0, size=14), axis.title.y = element_blank(), panel.background = element_blank(),  axis.ticks = element_blank(), legend.position = "bottom", legend.text = element_text(size = 12), legend.title = element_text(size=12), axis.text=element_text(size=15))
all_mods_multi_3a_mod
```
Fig 4C
```{r}
all_moderators_multi_3g$dataset = factor(all_moderators_multi_3g$dataset, levels = c("Complete, by group", "Training, by group", "Reactivation, by group"))
all_moderators_multi_3g = all_moderators_multi_3g %>% mutate(pval_cat = if_else(pvalor>0.05, ">0.05", if_else(pvalor<=0.05&pvalor>0.01, "<0.05", if_else(pvalor<=0.01&pvalor>0.001, "<0.01", "<0.001"))))
all_moderators_multi_3g[all_moderators_multi_3g=="relevel(factor(Intervention.Session), ref = 'Training')"] = "Intervention session"
all_moderators_multi_3g[all_moderators_multi_3g=="relevel(factor(Site.injection.other), ref = 'Systemic')"] = "Injection site"
all_moderators_multi_3g[all_moderators_multi_3g=="relevel(factor(ConditioningType), ref = 'Contextual')"] = "Conditioning type"
all_moderators_multi_3g[all_moderators_multi_3g=="Active.Drug"] = "Active drug"
all_moderators_multi_3g[all_moderators_multi_3g=="NumberShock"] = "Number of shocks"
all_moderators_multi_3g[all_moderators_multi_3g=="duration_zscore"] = "Reexposure duration"
all_moderators_multi_3g[all_moderators_multi_3g=="InterventionTime"] = "Intervention time"

all_moderators_multi_3g$predictor = factor(all_moderators_multi_3g$predictor, level = c("Injection site", "Active drug", "Reexposure duration", "Intervention time", "Number of shocks", "Intervention session", "Conditioning type"))
all_moderators_multi_3g = all_moderators_multi_3g %>% mutate(R2_mod = abs(weighted_mean_R2))
all_moderators_multi_3g = rbind(all_moderators_multi_3g, c(NA, NA, "Conditioning type", "Complete, by group", NA, NA, NA))

all_mods_multi_3g_mod = ggplot(all_moderators_multi_3g, aes(x=dataset, y=predictor, fill=pval_cat)) + 
  geom_tile() + scale_fill_manual(values=c("#3182bd","#6baed6","#bdd7e7","#eff3ff"), na.value = "white", name = "Q test for moderators \np value") +
  geom_text(aes(x=dataset, y=predictor, label = if_else(condition = R2_mod>=0, true = paste(round(as.numeric(R2_mod), 1), "%"), false = " ")), color = "black", size = 6) +
  scale_x_discrete(position = "top", labels = labels_3g) + 
  theme(axis.title.x = element_blank(), axis.text.x.top = element_text(angle = 0, size = 14), axis.title.y = element_blank(), panel.background = element_blank(),  axis.ticks = element_blank(), legend.position = "bottom", axis.text = element_text(size=15), legend.text = element_text(size=12), legend.title = element_text(size=12))
all_mods_multi_3g_mod
```

Complete Fig 4
```{r}
plot_grid(all_mods_multi,all_mods_multi_3a_mod, all_mods_multi_3g_mod, ncol = 3, labels=LETTERS)
ggsave("FIG_4.png", width = 60, height = 30, units = "cm")
```

```{r}

```
````
